# *De novo* assembly of 64 haplotype-resolved human genomes of diverse ancestry and integrated analysis of structural variation

**DOI:** 10.1101/2020.12.16.423102

**Authors:** Peter Ebert, Peter A. Audano, Qihui Zhu, Bernardo Rodriguez-Martin, David Porubsky, Marc Jan Bonder, Arvis Sulovari, Jana Ebler, Weichen Zhou, Rebecca Serra Mari, Feyza Yilmaz, Xuefang Zhao, PingHsun Hsieh, Joyce Lee, Sushant Kumar, Jiadong Lin, Tobias Rausch, Yu Chen, Jingwen Ren, Martin Santamarina, Wolfram Höps, Hufsah Ashraf, Nelson T. Chuang, Xiaofei Yang, Katherine M. Munson, Alexandra P. Lewis, Susan Fairley, Luke J. Tallon, Wayne E. Clarke, Anna O. Basile, Marta Byrska-Bishop, André Corvelo, Mark J.P. Chaisson, Junjie Chen, Chong Li, Harrison Brand, Aaron M. Wenger, Maryam Ghareghani, William T. Harvey, Benjamin Raeder, Patrick Hasenfeld, Allison Regier, Haley Abel, Ira Hall, Paul Flicek, Oliver Stegle, Mark B. Gerstein, Jose M.C. Tubio, Zepeng Mu, Yang I. Li, Xinghua Shi, Alex R. Hastie, Kai Ye, Zechen Chong, Ashley D. Sanders, Michael C. Zody, Michael E. Talkowski, Ryan E. Mills, Scott E. Devine, Charles Lee, Jan O. Korbel, Tobias Marschall, Evan E. Eichler

**Affiliations:** Heinrich Heine University, Medical Faculty, Institute for Medical Biometry and Bioinformatics, Moorenstr. 20, 40225 Düsseldorf, Germany; Department of Genome Sciences, University of Washington School of Medicine, 3720 15th Ave NE, Seattle, WA 98195-5065, USA; The Jackson Laboratory for Genomic Medicine, 10 Discovery Dr, Farmington, CT 06030, USA; European Molecular Biology Laboratory (EMBL), Genome Biology Unit, Meyerhofstr. 1, 69117 Heidelberg, Germany; Division of Computational Genomics and Systems Genetics, German Cancer Research Center (DKFZ), 69120 Heidelberg, Germany; Department of Computational Medicine & Bioinformatics, University of Michigan, 500 S. State Street, Ann Arbor, MI 48109, USA; Center for Genomic Medicine, Massachusetts General Hospital, Department of Neurology, Harvard Medical School, Boston, MA 02114, USA; Bionano Genomics, San Diego, CA 92121, USA; Program in Computational Biology and Bioinformatics, Yale University, BASS 432&437, 266 Whitney Avenue, New Haven, CT 06520, USA; School of Automation Science and Engineering, Faculty of Electronic and Information Engineering, Xi’an Jiaotong University, Xi’an, Shaanxi, 710049, China; Department of Genetics and Informatics Institute, School of Medicine, University of Alabama at Birmingham, Birmingham, AL 35294, USA; Molecular and Computational Biology, University of Southern California, Los Angeles, CA 90089, USA; Genomes and Disease, Centre for Research in Molecular Medicine and Chronic Diseases (CIMUS), Universidade de Santiago de Compostela, Santiago de Compostela, Spain; Department of Zoology, Genetics and Physical Anthropology, Universidade de Santiago de Compostela, Santiago de Compostela, Spain; Institute for Genome Sciences, University of Maryland School of Medicine, 670 W Baltimore Street, Baltimore, MD 21201, USA; School of Computer Science and Technology, Faculty of Electronic and Information Engineering, Xi’an Jiaotong University, Xi’an, Shaanxi, 710049, China; European Molecular Biology Laboratory, European Bioinformatics Institute, Wellcome Genome Campus, Hinxton, Cambridge, CB10 1SD, United Kingdom; New York Genome Center, New York, NY 10013, USA; Department of Computer & Information Sciences, Temple University, Philadelphia, PA 19122, USA; Pacific Biosystems of California, Inc., Menlo Park, CA 94025, USA; Max Planck Institute for Informatics, Saarland Informatics Campus E1.4, 66123 Saarbrücken, Germany; Washington University, St. Louis, MO 63108, USA; Genetics, Genomics, and Systems Biology, University of Chicago, Chicago, IL 60637 USA; Section of Genetic Medicine, Department of Medicine, University of Chicago, Chicago, IL 60637 USA; Precision Medicine Center, The First Affiliated Hospital of Xi’an Jiaotong University, 277 West Yanta Rd., Xi’an, 710061, Shaanxi, China; Howard Hughes Medical Institute, University of Washington, Seattle, WA 98195, USA

**Author notes:** Correspondence should be addressed to (E.E.E.), (T.M.), (J.O.K.) and (C.L.). These authors contributed equally to this work (Peter Ebert, Peter A. Audano, Qihui Zhu, Bernardo Rodriguez-Martin). Joint senior authors (Charles Lee, Jan O. Korbel, Tobias Marschall, Evan E. Eichler).

## Abstract

Long-read and strand-specific sequencing technologies together facilitate the *de novo* assembly of high-quality haplotype-resolved human genomes without parent–child trio data. We present 64 assembled haplotypes from 32 diverse human genomes. These highly contiguous haplotype assemblies (average contig N50: 26 Mbp) integrate all forms of genetic variation across even complex loci such as the major histocompatibility complex. We focus on 107,590 structural variants (SVs), of which 68% are inaccessible by short-read sequencing. We identify new SV hotspots (spanning megabases of gene-rich sequence), characterize 130 of the most active mobile element source elements, and find that 63% of all SVs arise by homology-mediated mechanisms—a twofold increase from previous studies. Our resource now enables reliable graph-based genotyping from short reads of up to 50,340 SVs, resulting in the identification of 1,525 expression quantitative trait loci (SV-eQTLs) as well as SV candidates for adaptive selection within the human population.

## INTRODUCTION

Advances in long-read sequencing, coupled with orthogonal genome-wide mapping technologies, have made it possible to fully resolve and assemble both haplotypes of a human genome (*1*–*3*). While such phased human genome assemblies generally improve variant discovery compared to Illumina or “squashed” long-read genome assemblies (*4*), the largest gains in sensitivity have been among structural variants (SVs)—inversions, deletions, duplications, translocations, and insertions ≥50 bp in length. Typical Illumina-based discovery approaches identify only 5,000–10,000 SVs (*1, 5, 6*) in contrast to long-read genome analyses that now routinely detect >20,000 (*1, 3, 4, 7*). Among the different classes of SVs, the greatest gains in sensitivity have been noted specifically for insertions where >85% of the variation has been reported as novel (*1*). In addition, repeat-mediated alterations within SV classes, such as variable number of tandem repeats (VNTRs) and short tandem repeats (STRs), have been challenging to delineate from short-read sequencing technologies and are underrepresented in the reference genome and often collapsed in unphased genome assemblies (*8*). The integration of long-read sequencing with new technologies such as single-cell strand sequencing (Strand-seq) has further catalyzed the unambiguous confirmation of both heterozygous and homozygous inverted configurations in a genome (*1, 9*). Long-read phased genome assemblies (*1*) also better resolve larger full-length mobile element insertions (MEIs) providing an opportunity to systematically investigate their origins, distribution, and the mutational processes underlying their mobilization within more complex regions of the genome, including transductions (*10, 11*).

The Human Genome Structural Variation Consortium (HGSVC) recently developed a method for phased genome assembly that combines long-read PacBio whole-genome sequencing (WGS) and Strand-seq data to produce fully phased diploid genome assemblies without dependency on parent–child trio data (**Fig. 1A**) (*3*). These phased assemblies enable a near-complete sequence-resolved representation of variation in human genomes. Here, we present a resource consisting of phased genome assemblies, corresponding to 70 haplotypes (64 unrelated and 6 children) from a diverse panel of human genomes. We focus specifically on the discovery of novel SVs performing extensive orthogonal validation using supporting technologies with the goal of comprehensively understanding the full complexity of SVs, including regions that cannot yet be resolved by long-read sequencing (**fig. S1**). Further, we genotype these newly defined SVs using a pangenome graph framework (*12*–*14*) into a diversity panel of human genomes now deeply sequenced (>30-fold) with short-read data from the 1000 Genomes Project (1000GP) (*15*). These findings allow us to establish their population frequency, identify ancestral haplotypes, and discover new associations with respect to gene expression and candidate disease loci. The work provides fundamental new insights into the structure, variation, and mutation of the human genome providing a framework for more systematic analyses of thousands of human genomes going forward.

**Fig 1.**
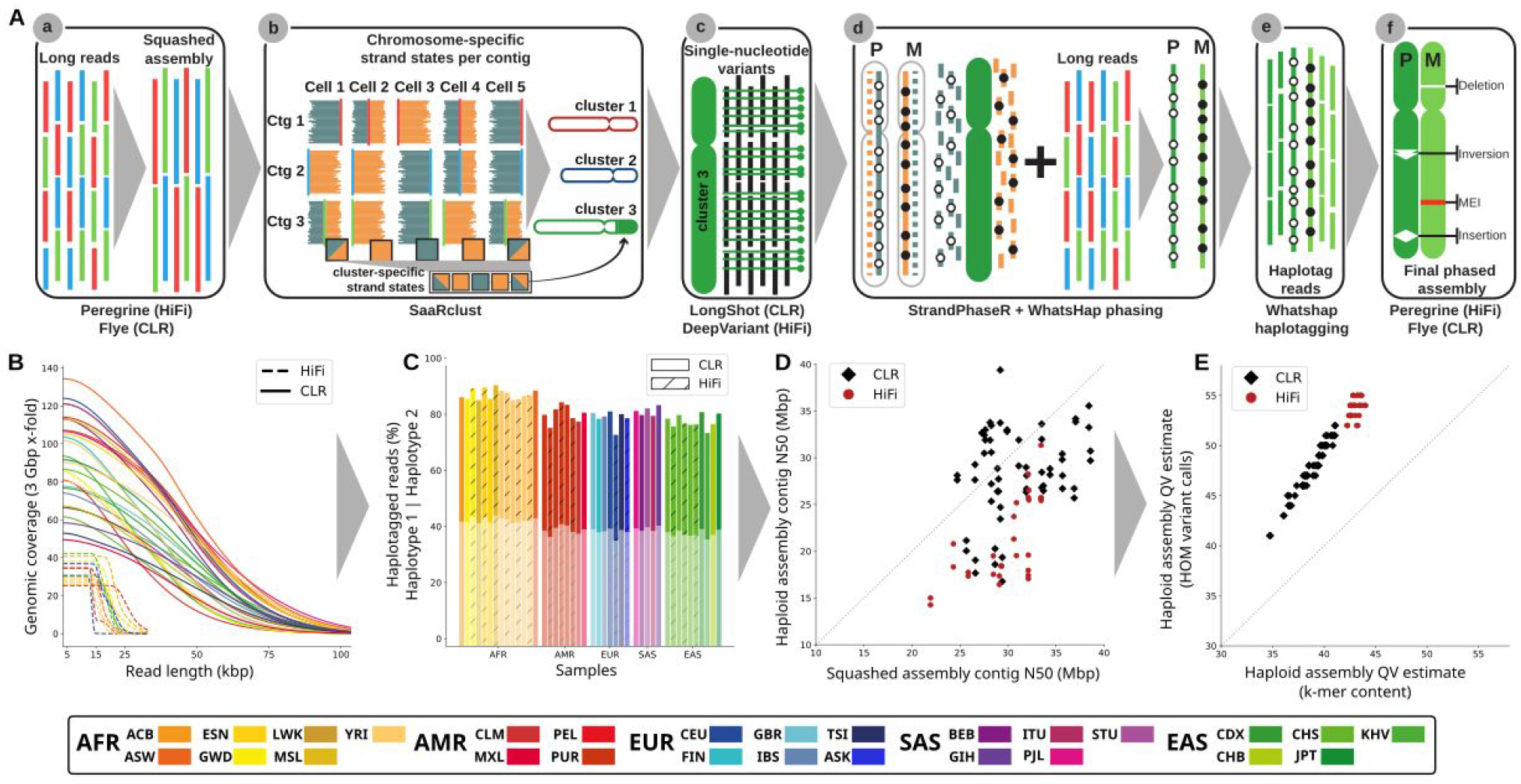
Trio-free phased diploid genome assembly using Strand-seq (PGAS). (**A**) A schematic of the key steps of the PGAS pipeline (*3*): (a) generation of a non-haplotype-resolved (“squashed”) long-read assembly; (b) clustering of assembled contigs into “chromosome” clusters based on Strand-seq Watson/Crick signal; (c) calling of single-nucleotide variants (SNVs) relative to the clustered squashed assembly; (d) integrative phasing combines local (SNV) and global (Strand-seq) haplotype information for chromosome-wide phasing; (e) tagging of input long reads by haplotype; (f) phased genome assembly based on haplotagged long reads and subsequent variant calling (Supplementary Information). (**B**) Genomic coverage (y-axis) as function of the long-read sequence read length (x-axis). (**C**) Fraction of reads that can be assigned (“haplotagged”) to either haplotype 1 (semitransparent) or haplotype 2 for HiFi (hatched) and CLR (solid) datasets. (**D**) Contig-level N50 values for squashed (x-axis) and haploid assemblies (y-axis) for CLR (black diamonds) and HiFi (red circles) samples. (**E**) Haploid assembly QV estimates computed from unique and shared k-mers (x-axis) based on homozygous Illumina variant calls (y-axis). Samples colored according to the 1000GP population color scheme (*15*) with exception of the added Ashkenazim individual NA24385/HG002 (Coriell family ID 3140) (ASK).

## RESULTS

### Sequencing and phased assembly of human genomes

We initially selected 34 unrelated individual genomes for *de novo* sequencing, with the goal of at least one representative from each of the 26 1000GP populations, of which 30 samples passed initial QC (**tables S1 and S2**). We additionally sequenced three previously studied child samples completing three parent–child trios, and we included for analysis publicly available sequencing data for two samples, NA12878 and HG002/NA24385, generated as part of the Genome in a Bottle effort (*16*). The complete set of 35 genomes includes 19 females and 16 males of African (AFR, n=11), American (AMR, n=5), East Asian (EAS, n=7), European (EUR, n=7) and South Asian (SAS, n=5; table S1) descent. All genomes were sequenced (Methods) using continuous long-read (CLR) sequencing (n=30) to an excess of 40-fold coverage or high-fidelity (HiFi) sequencing (n=12) to an excess of 20-fold coverage (**Fig. 1B, table S1**; see Data and materials availability). As a control for phasing and platform differences, we sequenced nine overlapping samples with both CLR as well as HiFi sequence data corresponding to the three parent–child trios (**tables S1 and S2**) that had been studied extensively for SVs previously by the HGSVC (*1*). For the purpose of phasing, we generated corresponding Strand-seq data (74-183 cells) for each of the samples. We used these data to successfully produce 70 (64 unrelated) phased and assembled human haplotypes (5.7 to 6.1 Gbp in length for the diploid sequence, **table S1**) using a reference-free assembly approach (**Fig. 1A**) (*3*), which works in the absence of parent–child trio information.

We find that the phased genomes are accurate at the base-pair level (QV > 40) and highly contiguous (contig N50 > 25 Mbp, **Fig. 1C-E, table S1**) with low switch error rates (median 0.12%, table S8) providing, for the first time, a diversity panel of physically resolved and fully phased single-nucleotide variant (SNV) and indel haplotypes flanking sequence-resolved SVs (**table S28**). Using two different metrics based on variant calling and k-mer content methods, respectively (**Fig. 1E**), we find that sequence accuracy is higher for human genome assemblies generated by HiFi (median QV = 54 [hom. var.] / 43 [k-mer], **Fig. 1E**) when compared to CLR (median QV = 48 [hom. var.] / 39 [k-mer], **Fig. 1E**) sequencing. Considering only accessible regions of the genome (Methods), the MAPQ60 contig coverage of HiFi and CLR genomes are similar (95.43% and 95.12%, **table S9**). CLR assemblies, however, are more contiguous (HiFi median contig N50 was 19.5 vs. 28.6 Mbp for CLR; p-value <10e-9, t-test). Fifteen of our assembled haplotypes exceed a contig N50 of 32 Mbp, all of which were based on CLR sequencing where insert libraries are much larger and sequence coverage is higher with half the number of single-molecule, real-time (SMRT) cells (**Fig. 1D, fig. S3, table S5**). Comparing Strand-seq phasing accuracy for six samples where parent–child trio data are available (**table S8, figs. S4, S5**; see Methods in (*3*)), we estimate on average 99.86% of all 1 Mbp segments are correctly phased from telomere-to-telomere (average switch error rate of 0.18% and Hamming distance of 0.21%, **table S8**). Predictably (*3*), remaining assembly gaps are enriched (Methods) in regions of segmental duplications (SDs) and acrocentric and centromeric regions of human chromosomes (**figs. S8, S9, table S10**). As a final QC of assembly quality, we analyzed Bionano Genomics optical mapping data for 32 genomes and found a median concordance of >97% between the optical map and the phased genome assemblies (**figs. S8, S9, table S11**).

### Phased variant discovery and distribution

Unlike previous population surveys of structural variation (*1, 4, 17*–*19*), which mapped reads or unphased contigs to the human reference genome, we developed the Phased Assembly Variant (PAV) caller (https://github.com/EichlerLab/pav) to discover genetic variants based on a direct comparison between the two sequence-assembled haplotypes and the human reference genome, GRCh38 (Methods). In the end, each human genome is rendered into two haplotype-resolved assemblies (each 2.9 Gbp) where all variants are physically linked (**table S28**). We classify variants as SNVs, indels (insertions and deletions 1-49 bp), and SVs (≥50 bp), which includes copy number variants (CNVs) and balanced inversion polymorphisms. After filtering (Methods), our nonredundant callset contains 107,590 insertion/deletion SVs, 316 inversions, 2.3 million indels, and 15.8 million SNVs. We observe the expected 2 bp periodicity for indels (dinucleotide repeats) and modes at 300 bp and 6 kbp for Alu and L1 MEIs, respectively (**Fig. 2A**), with only a small fraction intersecting functional elements (*20*) (**Fig. 2B**). PAV readily flags all reference-based artefacts or minor alleles by pinpointing regions where the 64 phased human genomes consistently differ from GRCh38 (1,573 SVs, 18,630 indels, and 91,537 SNVs, “shared variants”) (**Fig. 2C**, Methods). The greater haplotype diversity allows us to reclassify 50% previously annotated shared SVs (*4*) as minor alleles and correct the coding sequence annotation of five genes with tandem repeats (*RRBP1, ZNF676, MUC2, STOX1*) or extreme GC content (*SAMD1*) (**table S32**). We estimate an FDR of 5-7% for SVs based on support from sequence-read-based callers, as well as an independent alignment method (**table S30**, Methods). Similarly, we estimate a 6% FDR for indels and 4% for SNVs based on an assessment of Mendelian transmission error from the HiFi and CLR parent–child trios (**table S31**, Methods). We find that 42% of the SVs are novel when compared to recent long-read surveys of human genomes (*1, 4, 17*–*19*) (**fig. S44**). The addition of African samples more than doubles the rate of new variant discovery when compared to non-Africans for all classes of variation (2.21× SVs (809 vs. 366), 3.70× indels (11,514 vs. 3,109), and 2.97× SNVs (160,232 vs. 54,006) for the 64th haplotype (**Fig. 2C, table S29**, Methods). On average, we detect 24,653 SVs (14,914 insertions, 9,622 deletions, 117 inversions), 794,406 indels (407,693 insertions, 386,713 deletions), and 3,895,274 SNVs per diploid human genome (**table S28**).

**Fig 2.**
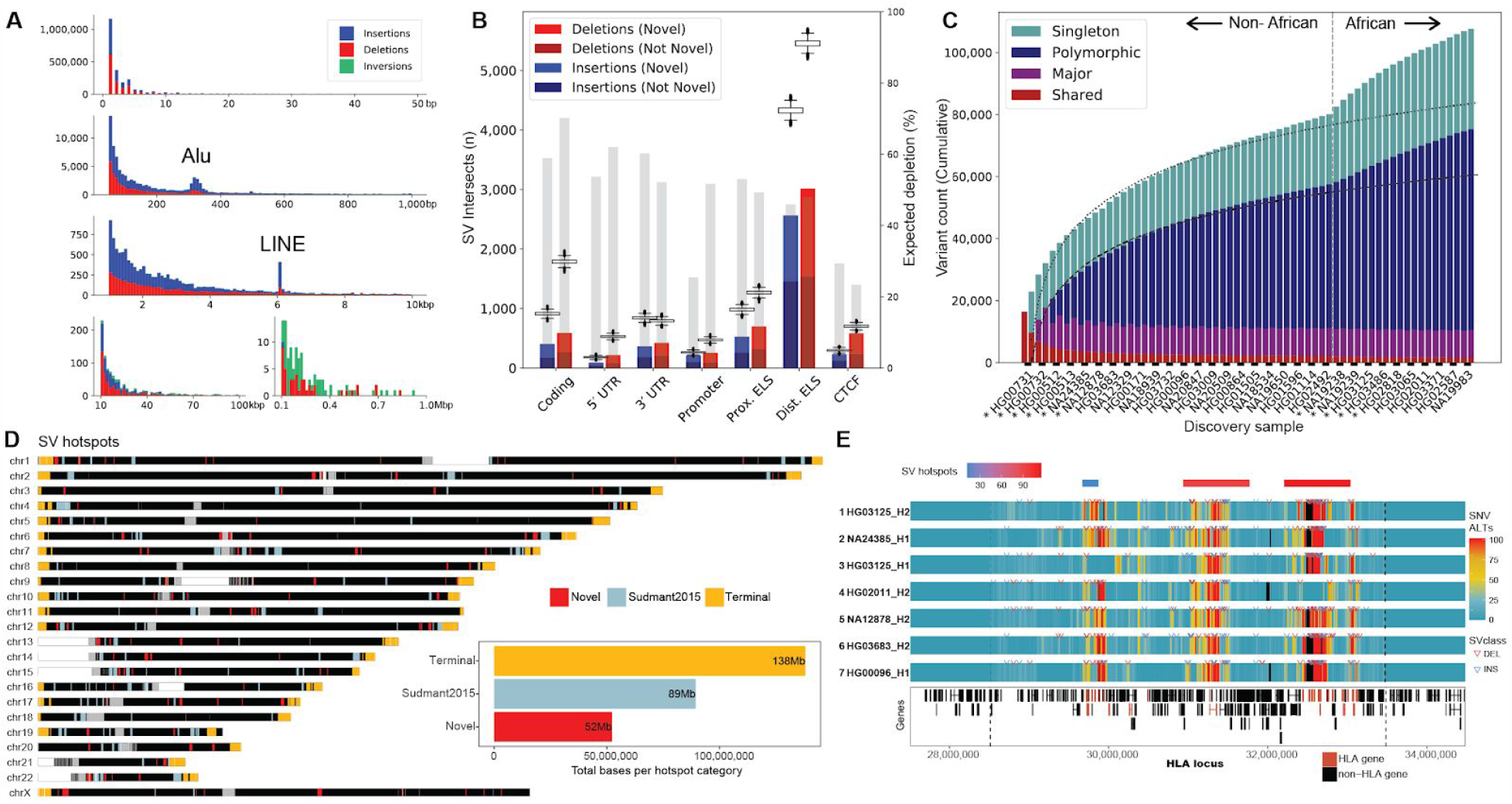
Variant discovery and distribution. (**A**) Size distribution of indels and SVs from 64 unrelated reference genomes shows expected 2 bp periodicity for indels, 300 bp peak for Alu insertions (second row), and 6 kbp peak for L1 MEIs. (**B**) The number of SVs intersecting functional elements (horizontal axis) compared to randomly permuting SV locations (box plots). Gray bars depict percent depletion (right axis scale). ELS: Enhancer-like signature. CTCF: CCCTC-binding factor. (**C**) The rate of SV discovery slows with each new haplotype (regression lines); however, the addition of haplotypes of African origin (dashed line) increases SV yield. We distinguish SVs as shared among all human haplotypes and not present in GRCh38 (red), major allele variants (AF≥50%, purple), polymorphisms (≥2 haplotypes, blue) from singletons (teal). (**D**) Genome-wide distribution of SV hotspots divided in three categories: last 5 Mbp of chromosomes (yellow), overlapping (light blue), and novel (red) when compared to short-read SV analysis of 1000GP (*21*).The total sequence length is represented by each hotspot category (inset). (**E**) Heatmap of seven SV haplotypes for 4 Mbp MHC region (chr6:28510,120-33,480,577 dashed lines) comparing regions of high SNV (red) and low diversity (blue) regions based on the number of alternate SNVs compared to the reference (GRCh38; alignment bin size 10 kbp, step 1 kbp). Phased SV insertions (blue arrows) and deletions (red arrows) are mapped above each haplotype. The most diverse regions correspond to SV hotspots (red/blue bars top row) and cluster with HLA genes (red bottom track).

SVs are particularly clustered and we identify 278 SV hotspots (**Fig. 2D, table S33**, Methods) spanning ∼279 Mbp of the genome (**Fig. 2D, fig. S48**). We find that 30.6% (32,222/105,327) of SVs on autosomes and chromosome X map within the last 5 Mbp of chromosome arms, corresponding to a ∼4-fold enrichment (p=0.001, z-score=301.3), with few notable exceptions—the long arm of the X chromosome and the short arms of chromosomes 3 and 20 (**Fig. 2D, fig. S46A**). Focusing on SVs > 5 Mbp from chromosome ends (73,105) (Methods), we identify 221 hotspots (**fig. S46B**). Of these, 49% (109/221) have not been previously detected based on short-read analysis of the 1000GP data (*21*). These interstitial hotspots are enriched 6.6-fold (p=0.001, z-score=26.6) for SDs consistent with homologous recombination and frequently correspond to gene-rich regions of exceptional diversity among human populations. For example, we identify three distinct hotspots mapping to the major histocompatibility complex (MHC) region that distinguish seven diverse structural haplotypes (**Fig. 2E, table S34**). Our analysis indicates that a majority (98.85%) of this 4 Mbp region has been sequence resolved at the base-pair level (29 of the assemblies are a single assembled contig and 18 have a single gap). The most structurally diverse regions also correspond to HLA (human leukocyte antigen) genes also enriched for single-nucleotide polymorphisms (SNPs) and indel polymorphisms.

A detailed analysis of the SVs with unambiguous breakpoint locations provided an opportunity to examine mechanisms of SV formation (*22*–*24*). Excluding MEIs and SVs with ambiguous breakpoints, we assessed 52,974 insertions and 30,467 deletions (**table S38**). We find 58% of insertions and 70% of deletions, including SVs in VNTRs, are flanked by at least 50 bp of homologous sequence suggesting formation by homology-directed repair (HDR) processes or non-allelic homologous recombination (NAHR). Amongst those, 15% of insertions and 25% of deletions showed greater than 200 bp flanking homology and are more likely mediated by NAHR. VNTRs with short repeat units (<50 bp) account for a smaller number of events (1.6% insertions and 0.4% deletions) and suggest replication slippage-mediated expansion and contraction. Additionally, 40% of insertions and 29% of deletions show blunt-ended breakpoints or microhomology (<50 bp flanking sequence identity), consistent with non-homologous end joining, microhomology-mediated end joining, or microhomology-mediated break-induced replication (*25*). Homology-associated SVs are twofold more frequent than expected based on previous reports using short reads (*22*–*24*), and when considering Illumina sequencing-based SV calls from the same samples, only 2% of insertions and 19% of deletions appear to be NAHR-mediated SVs with ≥200 bp flanking homology (p-value <2.2e-16; Fisher’s exact test; **table S38**).

Breakpoints and SVs more generally are depleted within protein-coding sequences and other functional elements with the exception of specific gene families where variability in the length of amino acid sequences relates to the function of the molecule (lipoprotein (A), mucins, zinc finger genes, etc.; **table S49**). We identify 9.4% of all SV breakpoints that intersect functional elements, such as exons (n=993), untranslated regions (UTRs; n=1,097), promoters (n=466), and enhancer-like elements (n=6,796) (**Fig. 2B, table S45**). When we consider structural polymorphisms that arise from perfect triplet repeats, expansions outnumber contractions 3 to 1 (271 expansions, 88 contractions) consistent with such regions being systematically underrepresented in the original reference (*8, 26*). Over the 64 haplotypes, there are six such SVs per haplotype and we identify a total of 106 nonredundant loci (**tables S46, S47**). Of note, 5/7 of the largest insertions of uninterrupted CTG or CGG repeat insertions mapping within exons correspond to genes already associated with triplet repeat instability diseases or fragile sites. For example, we identify a 21-copy CTG repeat expansion in *ATXN3* (Machado-Joseph disease), a 17-copy gain of CAG in *HTT* (Huntington’s disease), a 21-copy gain of a CGG repeat in *ZNF713* (Fragile site 4A), and a 36-copy CGG gain in *DIP2B* (Fragile site 12A) (Methods). The discovery of these perfect repeat insertion alleles with respect to the human reference provides an important reference for future investigations of triplet repeat instability.

### Mobile element insertions

Based on the phased genome assemblies, we identified the largest collection (n=9,453) of fully sequence-resolved non-reference MEIs, including 7,743 Alus, 1,170 L1Hs, and 540 SVAs (Supplementary Methods) and used sequence content of the elements and their flanking sequences to provide insight into their origin and mechanisms of retrotransposition. Full-length L1 (FL-L1) elements are an especially relevant source of genetic variation since they continue to mutagenize germline and somatic cells and can lead to gene disruptions that cause human disease (*27, 28*). While a minority (28%; 329/1,170) of L1s are full-length (**fig. S35, table S24**), we find that 78% of FL-L1s (257/329) possess two intact open reading frames (ORF1 and ORF2), encoding the proteins that drive L1, Alu, SVA, and processed pseudogene mobilization. Indeed, 23% (76/329) of these sequences show evidence of activity as they are part of a database of 198 FL-L1s known to be active *in vitro* (*29, 30*), in human populations (*31*), and in cancers (*32*–*34*). Most active copies (72%; 142/198) are either in our callset or present in the reference genome and are now fully sequence resolved (**table S25**). We note that 19% (27/142) of the active FL-L1s have at least one ORF disrupted, which includes a hot element at 9q32 reported to be highly active in diverse tumors (*32*). Finally, using L1 *Pan troglodytes* (L1Pt) as an outgroup, we construct a phylogeny of active human L1s and estimate their age in millions years (Myr) (**Fig. 3A, fig. S36**). As expected, Ta-1 copies are the youngest (mean = 1.00 [95% CI: 0.88-1.13]), followed by Ta-0 (mean = 1.63 [95% CI: 1.49-1.77]) and pre-Ta (mean = 2.15 [95% CI: 1.91-2.40]). Notably, the evolutionary age correlates with L1 features such as subfamily, level of activity, and allele frequency—with Ta-1 sequences being highly polymorphic and active. Indeed, three out of the four youngest FL-L1s, namely 2q24.1 (age estimate = 0.20 Myr), 6p24.1 (0.39) and 6p22.1-2 (0.45), are Ta-1 copies reported to be extremely active in cancer genomes (*32*). In contrast, 1p12 is a fixed Pre-Ta insertion that despite integrating into the human genome approximately 1.8 Myr ago, remains highly active both in the germline (*31*) and somatically associated with tumors (*32*–*34*). This indicates that a small set of pre-Ta representatives possibly remain very active in the human genome.

**Fig 3.**
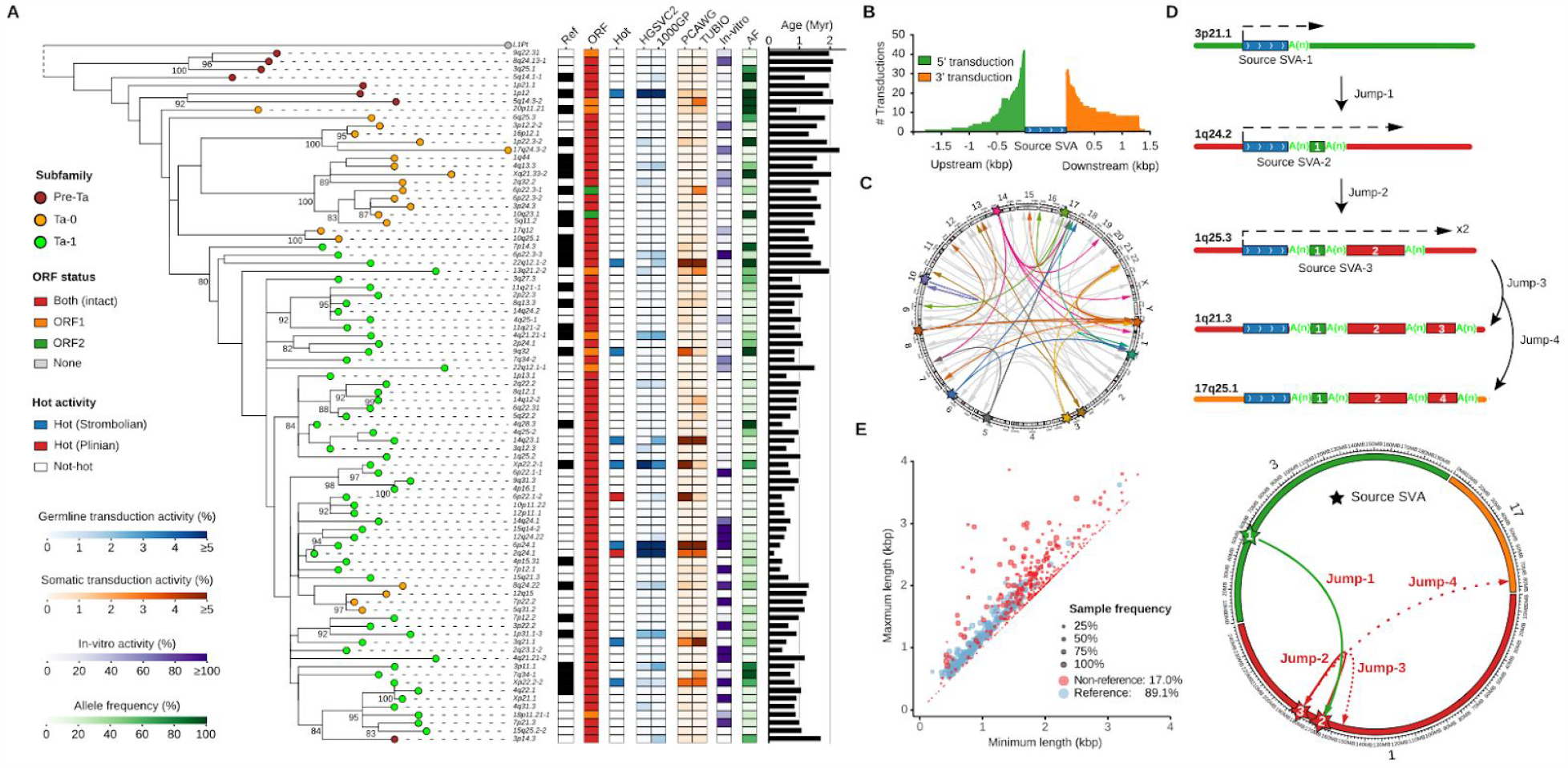
Mobile element insertions. (**A**) Maximum-likelihood phylogenetic tree for active sequence-resolved FL-L1s annotated by subfamily designation, presence/absence on the reference, ORF content, and hot activity profile (*32*–*34*) (bootstrap values >80% shown). Consensus sequence for L1 *Pan troglodytes* (L1Pt) is included as an outgroup. Heatmaps represent allele frequency (AF) based on the assembly discovery set, activity estimates based on *in vitro* assays (*29, 30*) and the number of transduction events detected in human populations (*31*) or cancer studies (*32*–*34*). (Phylogeny does not include those FL-L1s with low activity; see **fig. S36** for a complete representation.) (**B**) Size distribution and number of 5’ and 3’ SVA-mediated transductions based on the analysis of flanking sequences. (**C**) Circos plot of SVA transductions and source SVA loci. Source elements mediating multiple transductions are highlighted with a star. Transductions derived from these copies are colored according to their source, while those derived from single-transduction source SVAs are in gray. (**D**) Schematic and circos representation for serial SVA-mediated transduction events. Dashed arrows indicate SVA transcription initiation and end. Transduced sequences are shown as colored boxes with their length proportional to transduction size. (**E**) Distributions of VNTR length (x-axis: the minimum, y-axis: the maximum) of reference and non-reference SVA elements. Reference SVAs are shown as blue dots and non-reference SVAs as red dots. The dot size represents the sample frequency of SVAs among discovery samples in the HGSVC.

SVA source elements are able to produce 5’ and 3’ transductions through alternative transcription start sites or bypassing of normal poly(A) sites during retrotransposition (*10, 11*). We detected 77 transduced non-repetitive DNA sequences at SVA insertions ends (**table S26**). Interestingly, 5’ transductions are more abundant (58%, 45/77) than 3’ transductions (**Fig. 3B**), as opposed to L1s, which primarily mediate 3’ transduction events (95%, 89/94). We used these unique transduced sequences to trace the origin of all 77 SVAs to 54 source SVA elements (**fig. S37, table S27**). A majority of source loci (87%, 47/54) belong to the youngest human-specific SVA-E and SVA-F subfamilies (*35*), and only 11 source elements generate 38% (29/77) of the offspring insertions (**Fig. 3C**). SVA transductions can occasionally shuffle coding sequences as illustrated by the mobilization of a complete exon of *HGSNAT* by an intronic SVA in antisense orientation (**fig. S38**). In addition, one SVA source element appears to have caused three sequential mobilization events as indicated by nested transductions flanked by poly(A) tails (**Fig. 3D, fig. S39**). Finally, SVA elements harbor CpG-rich VNTRs in their interior regions that can expand and contract and have been associated with changes in local gene expression (*8*). Examining the fully sequence-resolved copy number differences, we find that non-reference SVAs show significantly greater variability in VNTR copy number compared to those present in the reference (p-value < 10e-5, student’s t-test, two-sided, **Fig. 3E**). Non-reference SVAs are significantly rarer (17.0%, p-value < 10e-5, student’s t-test, two-sided) when compared to reference SVAs (89.1%) in the discovery set, suggesting a more recent origin.

### Inversions

Copy number neutral inversions are among the most difficult SVs to detect and validate (*1*). We applied multiple approaches integrating Strand-seq, Bionano optical mapping, and PAV-based variant discovery to generate a comprehensive and orthogonally validated and manually curated set of inversions. PAV specifically increases inversion detection sensitivity for smaller events (**fig. S30B**) by including a novel k-mer density assessment to resolve inner and outer breakpoints of flanking repeats, which does not rely on alignment breaks to identify inversion sites (Supplementary Methods). PAV identifies an additional 43 inversions, on average, increasing sensitivity >2-fold compared to previous phased assembly callsets (*2*). In total, we discover on average 117 inversions per sample (316 nonredundant calls) (**fig. S30**, Methods). As expected, inversions flanked by SDs tend to be larger than those in unique regions of the genome (*36*) (Wilcoxon rank sum test (one-sided, greater), p-value: 3.2 x 10^−13^, **fig. S31**). We focus on one complex region mapping to chromosome 16p12 where we observed a large number of polymorphic inversions flanked by SDs (*9*) (**fig. S32A**). The region harbors 11 different inversions (red and gray arrows) distinguishing 22 different structural configurations that span a ∼2.5 Mbp gene-rich region of chromosome 16p (up to 13 protein-coding genes are flipped in orientation depending on human haplotypes) (**Fig. 4A**, Supplementary Methods). These configurations are distributed among human populations, but do not correspond to unique haplotypes (**Fig. 4A**). For example, an analysis of the flanking sequence shows that at least five of the inversions occur in multiple haplotype backgrounds, indicative of recurrent inversion toggling (*36, 37*) between a direct and inverted state (**fig. S33**, Supplementary Methods). Although Strand-seq data allow us to unambiguously identify the inversion status of the unique regions, most of the breakpoints themselves are not yet fully sequence resolved due to the presence of large repeats (*3*) (**Fig. 4A, fig. S32B**).

**Fig 4.**
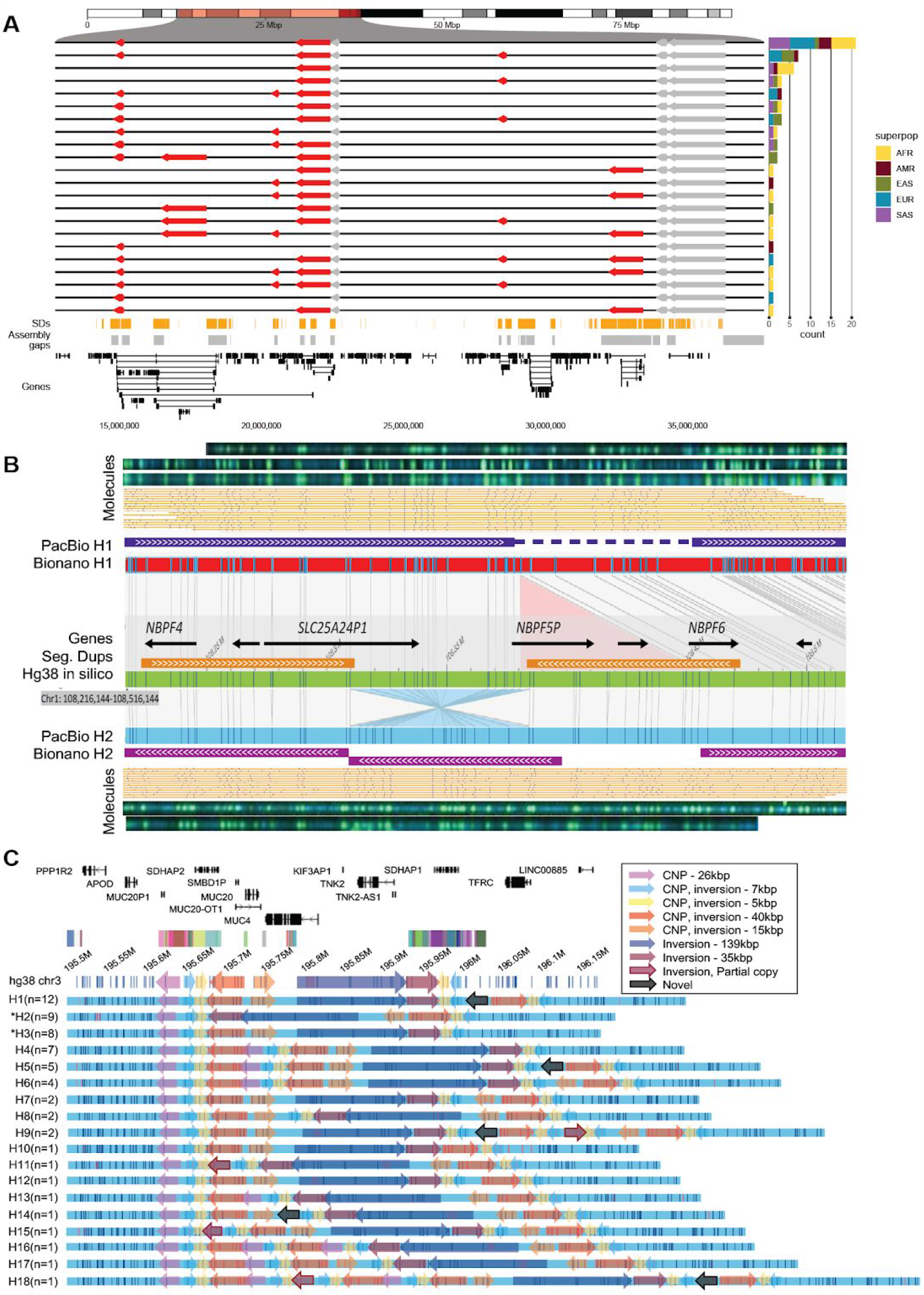
Complex patterns of structural variation. (**A**) An inversion hotspot mapping to a 2.5 Mbp gene-rich region of chromosome 16p12 (highlighted portion of ideogram). Haplotype structure of inversions (red arrows) are compared to the GRCh38 reference orientation (black lines) as well as additional inversions (gray), which could not be haplotype integrated because of uninformative markers. A barplot (right panel) enumerates the frequency of each distinct inversion configuration (n=22) by superpopulation for the 64 phased genomes. Bottom panels: Shows distribution of SDs (orange), assembly gaps (gray), and genes (black) in a given region. **(B)** A partially resolved complex SV locus (HG00733 at chr1:108,216,144-108,516,144). Optical maps generated by *DLE1* digestion predict a deletion (red bar, Bionano H1) and an inversion (blue bar, Bionano H2) when compared to GRCh38 (green bar). Haplotype structures are strongly supported by extracted single molecules (beige) and raw images (green dots). Phased assembly correctly resolves hap1 deletion (purple top) and Strand-seq detects the inversion (blue) but misses the flanking SD, which is a gap in the H2 assembly (gap). (**C**) Haplotype structural complexity at chromosome 3q29. Optical mapping of a gene-rich 410 kbp region (chr3:195,607,154-196,027,006) predicts 18 distinct structural haplotypes (H1-H8) that vary in abundance (n=1 to 12) and differ by at least 9 copy number SDs and associated inversion polymorphisms (see colored arrows). This hotspot leads to changes in gene copy and order (GENCODE v34 top panel). 26 haplotypes are fully resolved by phased assembly (21 CLR, 5 HiFi) and the median MAP60 contig coverage of the region is 96.1%.

### Complex structural variation events

We investigated the remaining gaps in our assemblies that map near or within centromeres, acrocentric regions, and SDs (**figs. S6, S7, table S10**). Because such repetitive regions have long been known to be enriched in complex variation (*38*) refractory to sequence assembly even with long-read data (*1*), we re-examined the genome-wide optical maps to assess additional regions of structural variation. In 30 samples, we find that 72% (14,231/19,821, summed across samples) of the large insertions and deletions (≥5 kbp) discovered by optical mapping are completely sequence resolved and concordant with the assembly (**table S17**) but the remainder show additional complexity. As an example, our analysis of the Puerto Rican phased genome assembly (HG00733) originally identified a 75 kbp deletion between the two haplotypes at chromosome 1p13.3, but a comparison with Bionano Genomics data shows a more complex pattern than a single deletion event: An inversion of 75 kbp is found in the alternate allele flanked by inverted SDs of 100 kbp involving *NBPF* genes (**Fig. 4B**). Interestingly, such discrepant regions appear to cluster in the genome. A comparison between the phased assemblies and the Bionano Genomics maps reveals 3,453 Bionano-unique insertions and 2,137 Bionano-unique deletions, corresponding to 1,657 nonredundant clusters (**table S19**) where a cluster might have PacBio support in one sample but not in other samples that have the same variant. If we restrict the analysis to clusters that are fully unresolved in the phased assemblies, we identify 1,175 regions (697 insertion clusters and 478 deletion clusters) (**table S20**). A majority of these (630/1,175 [383 insertion and 247 deletion clusters]) localize to SDs and overlap genes. We estimate that there are still ∼35 unresolved regions per human genome that are greater than 50 kbp in length where there are five or more distinct SV haplotypes in the human population. On chromosome 3q29, for example (**Fig. 4C**), we identify 18 distinct structural haplotypes involving at least nine copy number and inversion polymorphisms affecting hundreds of kilobases of gene-rich sequence (min. 375 kbp, max. 690 kbp) (**Fig. 4C**). This extraordinary pattern of structural diversity maps to the proximal breakpoint of the chromosome 3q29 microdeletion and microduplication syndrome rearrangement (chr3:195,999,954-197,617,802) associated with developmental delay and adult neuropsychiatric disease (*39*).

### Short-read vs. long-read SV discovery

Previous comparisons between long-read and short-read datasets have been limited by differences in samples and sequence coverage. We deeply sequenced 3,202 samples from the 1000GP (34.5-fold) (Supplementary Information) and discovered SVs using three state-of-the-art callers: GATK-SV (*5*), SVTools (*6*) and Absinthe (github.com/nygenome/absinthe). We focused on the 34 samples with matching PacBio long-read sequences detecting 9,338 SVs per genome (FDR <5%) resulting in the discovery and genotyping of 34,061 loci, including 15,565 deletions, 3,197 duplications, 260 CNVs displaying multiple copy states (mCNVs), 14,164 insertions, 194 inversions, and 681 complex SVs (**fig. S21A-C**). Compared to the long-read callset, we find 62.8% of deletions and 74.9% of insertions are not captured by short-read sequencing (**fig. S21D**). Most SVs specific to long-read sequencing localize to highly repetitive simple repeat (SR) and SD sequences (83% of deletions and 81% of insertions, **table S16**). The greatest added value in long-read sequencing is observed from increased sensitivity among deletions less than 250 bp and insertions under 10 kbp (**Fig. 5A, fig. S21E**). While recent human population SV resources published from short-read Illumina WGS from tens of thousands of human genomes, such as the Centers for Common Disease Genomics (CCDG) (*6*) and Genome Aggregation Database (gnomAD) (*5*), have successfully discovered SVs down to very low allele frequencies, a comparison of common SVs (allele frequency [AF]≥5%) emphasizes the substantial increase in sensitivity for our long-read-based callsets for relatively small insertions and deletions (**Fig. 5B, fig S22**). Nevertheless, read-depth approaches have the ability to detect larger copy number differences that have yet to be fully sequence resolved (*40, 41*). This is especially the case for duplications and mCNVs displaying three or more copy states (**fig. S21F**). Among the 31 samples compared, we detect 210 large CNVs (>5 kbp) (67 deletions, 41 duplications and 102 mCNVs) by read-depth analysis from the short-read data (**figs. S21F, S23**) and at least one-third of these correspond to the complex SVs highlighted above. While variation in these loci can be detected by read-depth analyses, their sequence structure and location in the genome are still unknown.

**Fig 5.**
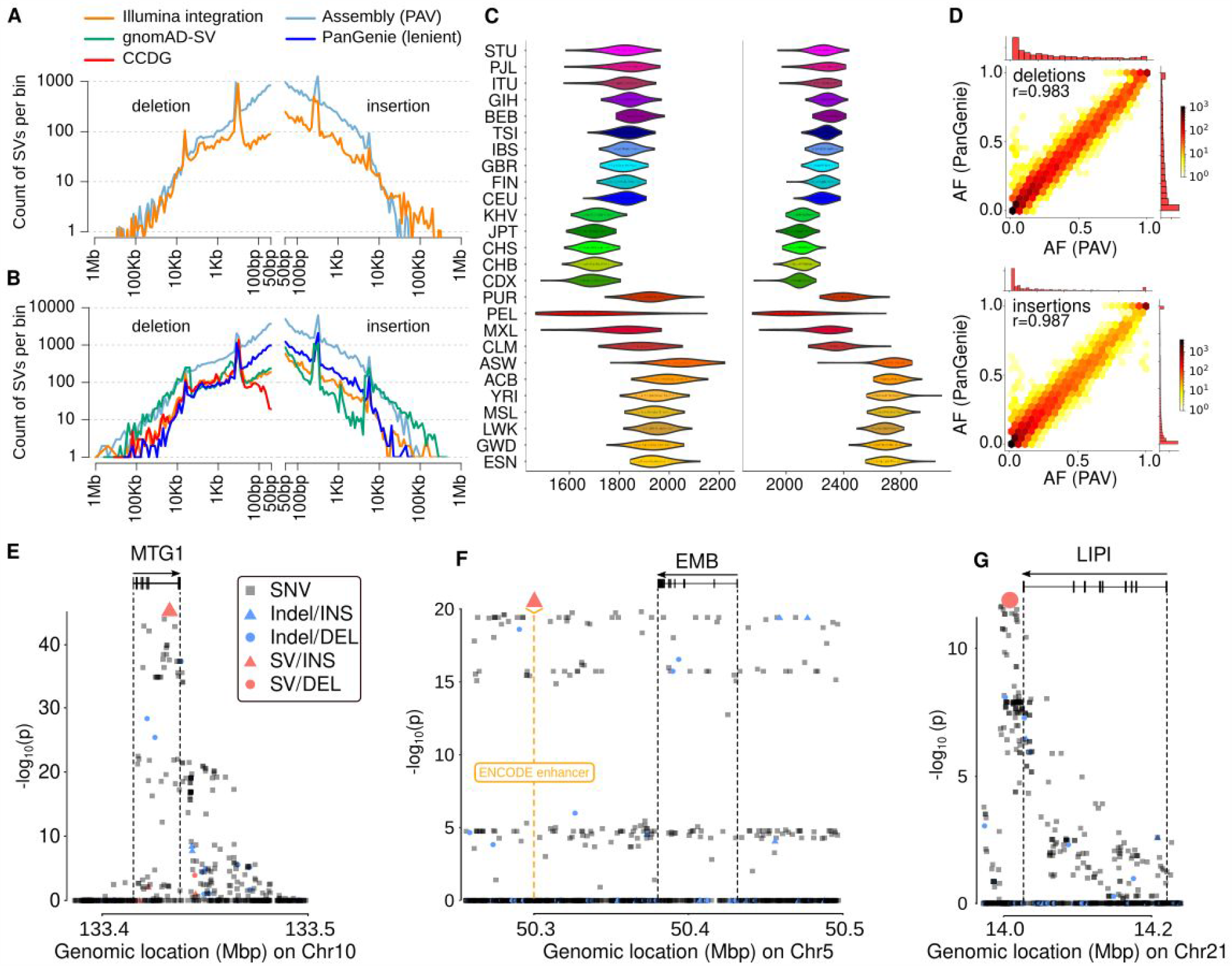
SV genotyping and eQTL analysis. (**A**) Length distribution of SVs in an average genome discovered from the PacBio assemblies using PAV and Illumina using an integrated callset from multiple tools. (**B**) Length distribution of total number of common SVs (AF>5%) represented in assembly-based callset, genotypable using PanGenie, CCDG and gnomAD-SV. **(C)** Distribution of heterozygous SV counts per diploid genome broken down by population, based on PanGenie genotypes passing strict filters (see **fig. S40** for unfiltered set). (**D**) Concordance of allele frequency (AF) estimates from the assembly-based PAV discovery callset and AF estimates from genotyping unrelated Illumina genomes (n=2,504) with PanGenie (strict genotype set of 24,107 SVs); marginal histograms are in linear scale. (**E-G**) Examples of lead SV-eQTLs (large symbols) in context of their respective genes, overlapping regulatory annotation, and other variants (small symbols). (**E**) An 89-base insertion (chr10-133415975-INS-89) is linked to decreased expression of *MTGI* (q-value:2.23e-11, Beta: -0.55 [-0.51 — -0.59]). (**F**) A 186-base insertion (chr5-50299995-INS-186), overlapping an ENCODE enhancer mark (orange), is the lead variant associated with decreased expression of *EMB* (q-value = 2.82e-06, Beta: -0.44 [-0.39 — -0.49]). (**G**) A 1,069-base deletion (chr21-14088468-DEL-1069) downstream of *LIPI* is linked to increased expression of *LIPI* (q-value = 0.0032, Beta=0.44 [0.38—0.50]).

### Genotyping

We applied PanGenie (*42*), a method designed to leverage a panel of assembly-based reference haplotypes threaded through a graph representation of genetic variation that takes advantage of the linkage disequilibrium inherent in the phased genomes. We initially performed this genotyping step using a reference set of 15.5M SNVs, 1.03M indels (1-49 bp), and 96.1k SVs (where there was less than 20% allelic dropout; **fig. S1, table S39**) and genotype these variants into the 1000GP short-read Illumina sequencing dataset (Supplementary Information) observing expected patterns of diversity (**Fig. 5C, figs. S76, S77**). As one measure of genotyping quality, we compare the allele frequencies derived from assembly-based PAV calls across the 64 reference haplotypes to short-read-based allele frequencies obtained from PanGenie for the 2,504 unrelated individuals. From the raw output of PanGenie, we observe an allele frequency correlation (Pearson’s) of 0.98 for SNVs, 0.95 for indels, and 0.85 for SVs. To further improve SV genotyping, we filter the variants by assessing Mendelian consistency, the ability to detect the non-reference allele, genotype qualities, and concordance to assembly-based calls in a leave-out-one experiment into account (Supplementary Information). Using these criteria, we define a subset of strict and lenient SVs for genotyping containing 24,107 SVs (25%) and 50,340 SVs (52%), respectively, with excellent allele frequency correlation of 0.99 (strict, **Fig. 5D**) and 0.95 (lenient, **fig. S74**). Performance metrics for deletions and insertions are comparable (strict set: SV deletions, r=0.98; SV insertions, r=0.99; **Fig. 5B**), highlighting the value of sequence-resolved insertion alleles being part of our reference panel, as well as the algorithm’s ability to leverage it (**fig. S72**). Beyond SVs, 12,283,650 SNVs (79%) and 705,893 indels (68%) met strict filter criteria (note: given this larger fraction, we did not define a lenient set for these variant classes). Importantly, we find that 42.5% (strict) and 59.9% (lenient) of our genotypable SVs are absent from our integrated SV callset for the same 3,202 short-read sequenced genomes (Supplementary Information). This ability to genotype variation typically not detected in Illumina callsets is also reflected in increased numbers of common SVs (AF>5%), particularly deletions below 250 bp and insertions, genotyped by PanGenie compared to CCDG and gnomAD-SV (**Fig. 5B**).

### eQTL analyses

We applied PanGenie genotypes (strict set) to systematically discover expression quantitative trait loci associated with structural variation (SV-eQTLs). First, we performed deep RNA-seq (>200M fragments) of the corresponding 34 lymphoblastoid cell lines and integrated these data with 397 transcriptomes of 1000GP samples from GEUVADIS (*43*). We pursued *cis*-eQTL mapping across the merged set of 427 donors, using a window of 1 Mbp centered around the transcription start site of a gene, testing all variants with a minor allele frequency of ≥1% and at Hardy-Weinberg equilibrium (P ≤ 0.0001). We considered 23,866 expressed genes, 15,452 of which were protein-coding. Using this design, we identify 57,953 indel-eQTLs (linked to 6,743 unique genes) and 2,108 SV-eQTLs (linked to 1,525 unique genes; **table S42**), at an FDR of 5%. The set includes 713 lead indel-eQTLs and 34 lead SV-eQTLs at distinct genes, respectively (**table S42**). In line with prior studies (*21, 44*), lead eQTL hits are enriched for SVs (Fisher’s exact p-value = 0.0436, OR = 1.8 [1.0 - 3.5]) as well as smaller indels (p-value = 1.92e-6, OR = 1.33 [95% CI: 1.2-1.5]), whereas they are depleted for SNVs (p-value = 2.7e-7, OR = 0.74 [0.7-0.8]).

We overlapped lead SV-eQTLs with our Illumina-based discovery callset (Supplementary Information) and a recent large-scale SV study of 17,795 genomes (*6*) and find that 48% (16 out of 33 SVs) of the lead eQTL associations reported here are novel. Of these previously inaccessible SVs, 12 (75%) correspond to insertions (2 Alu MEIs, 3 tandem duplications, and 7 repeat expansions)—SV classes typically under-ascertained in short-read datasets (*1*). For example, one of our top novel lead SVs is a 89 bp VNTR insertion in the terminal intron of the mitochondrial ribosome-associated GTPase 1 gene (*MTG1;* **Fig. 5E**) and is seen in conjunction with decreased expression. Similarly, we identify a 186 bp insertion in an ENCODE enhancer for B-cell lymphomas, which is associated with reduced expression of the immunoglobulin superfamily gene embigin (*EMB;* **Fig. 5F**). In contrast, we sequence resolve a 1,069 bp deletion located in an SD region downstream of the Lipase I gene (*LIPI;* **Fig. 5G**) and find that it is associated with increased gene expression of *LIPI*. SNPs at this locus have been linked to heart rate in patients with heart failure with reduced ejection fraction in a previous genome-wide association study (GWAS, p-value: 9.0e-06) (*45*).

### Ancestry and population genetic analyses

The availability of haplotype-phased assemblies provides an opportunity to explore the ancestry and population genetic properties of the genomes and SVs at multiple levels. We applied a machine-learning method (*46*) and developed a hidden Markov model to identify ancestry-informative SNVs and to assign ancestral segments per block based on population genetic data from the Simons Genome Diversity Project (SGDP) (*47*) (Supplementary Information). The two methods as well as the different sequencing platforms produce highly concordant results (>90%, **fig. S83**). At the family level, we can accurately assign paternal and maternal haplotypes and distinguish recombination crossover events in the child compared to parental haplotypes (**Fig. 6A**). At the population level, on average 87.2% of the assembled sequence can be assigned ancestry. 1000GP samples originating from the African continent show the largest tracts of uniform ancestry (mean length = 23.6 cM, **Fig. 6B, fig. S24**) in contrast to North and South American populations (mean length=2.65 cM, **Fig. 6B, fig. S24**) and South Asians (mean length=4.38 cM, **Fig. 6B**), consistent with recent and more ancient admixture. For example, the African American, African Caribbean, and admixed American 1000GP samples show the greatest diversity of ancestral segments (**Fig. 6B, figs. S24, S25**) most likely as a result of the transatlantic slave trade and colonial era migration (*48*).

**Fig 6.**
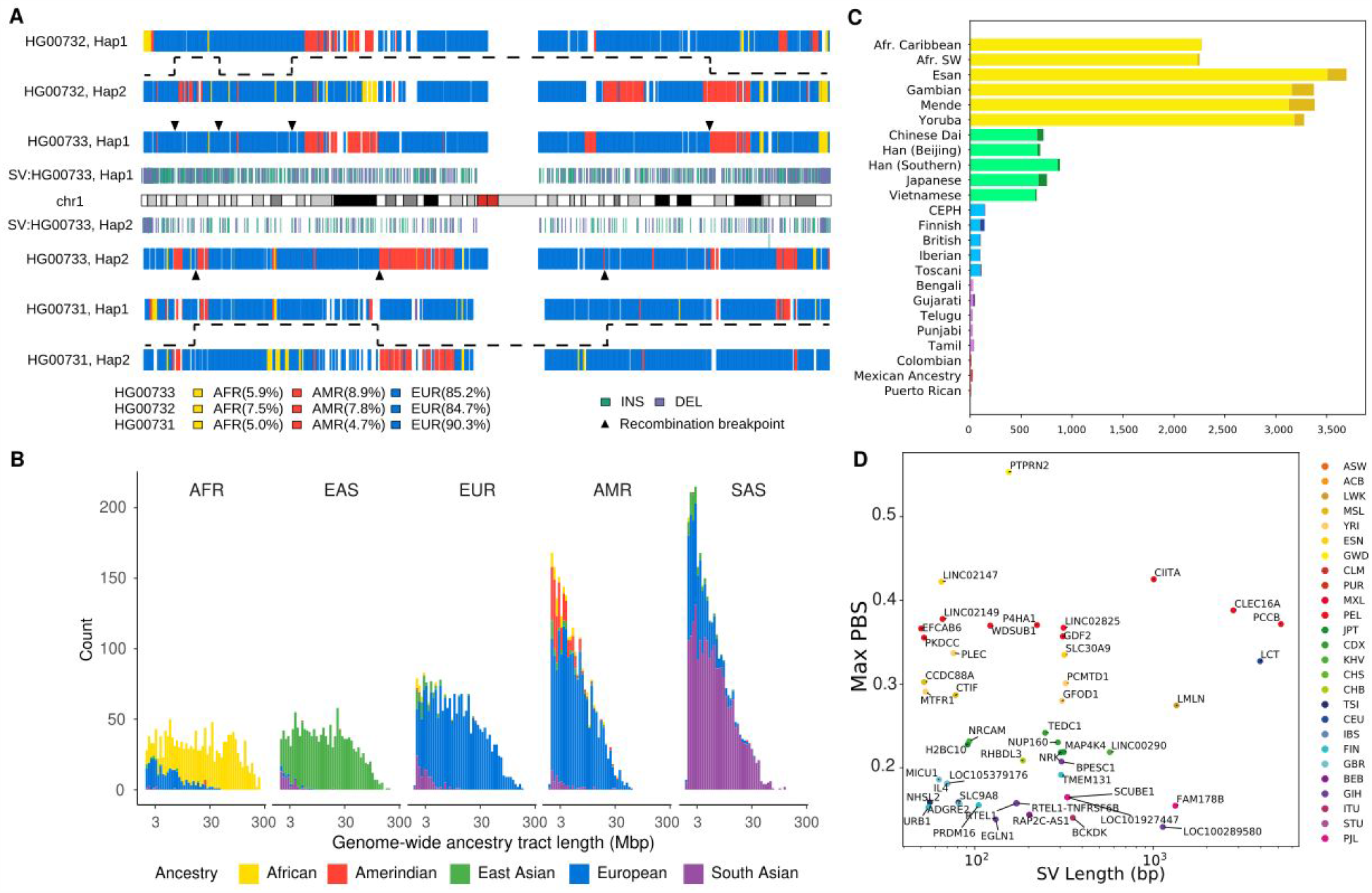
Ancestry and population differentiation inferences using haplotype-phased diploid assemblies. (**A**) Inferred local ancestries (based on an SGDP reference panel) for maternal (upper) and paternal (bottom) haplotypes of a Puerto Rican 1000GP genome (HG00733) are compared to parental haplotypes (maternal: HG00732, paternal: HG00731). Ancestral segments are colored (AFR: yellow, AMR: red, and EUR: blue) and are consistent with the recent demographic history of the island (Methods). HG0733 SVs (≥50 bp; insertion: green, deletion: purple), inferred recombination breakpoints (triangles), and transmission of recombinant parental haplotypes (dashed lines) are shown. (**B**) Length distribution (log10 scale) of ancestry tracts among the 64 genomes assigned to five superpopulations shows evidence of recent (AMR) and more ancient (SAS) admixture. (**C**) Top population-specific Fst variants (dark color) and top superpopulation-specific Fst variants (light color). The number of stratified SVs differs by orders of magnitude depending on population. (**D**) Top SV PBS values within 5 kbp of genes identifies SV candidates for selection and disease.

Focusing on our more comprehensive genotyping of SVs, we searched for population-stratified variants since these are potential candidates for local adaptation (*49, 50*) that could not have been characterized in the original study of 1000GP populations (*15*). Using Fst as a metric, we find that the number of such population-stratified variants varies widely among different groups likely as a consequence of ancestral diversity (Africans), population bottlenecks (East Asians), and admixture (South Asians) (**Fig. 6C**). Restricting our analysis to SVs located within 5 kbp of genes, we identify 117 stratified SVs (population branch statistic or PBS greater than 3 s.d. (*50*)) (**table S44, S48**) and further characterize these by the number of base pairs deleted or inserted per locus (**Fig. 6D**). The greatest outlier is a 4.0 kbp insertion within the first intron of the *LCT* (lactase gene) originally reported based on fosmid sequencing from European samples (*47*). We determine that the corresponding insertion is ancestral (i.e., the human reference genome carries the derived deleted allele), the insertion harbors 11 predicted transcription factor binding sites, and the deletion likely occurred as a result of an Alu-mediated NAHR event ∼520,000 years ago (**Fig. S87**). *LCT* variation is one of the most well-known genes under adaptive evolution among Europeans. Notably, the reported causal, derived allele of lactase persistence in Europeans (−13910*T; rs4988235) is in complete linkage disequilibrium (D′=1) with the reference allele of this SV, and it will be interesting to determine the functional roles of these two mutations in lactase persistence (*51*). In other cases, the population-stratified variants are nested among known regulatory elements or intersect them directly, such as a 76 bp tandem repeat expansion in a PLEC intron, a cytoskeleton component, seen only in Africans (AF=0.82) and Americans (AF=0.06). Similarly, we identify a 2.8 kbp insertion mapping near potential repressor binding sites in a *CLEC16A* intron, a gene associated with type 1 diabetes when disrupted (*52*). This variant shows a high frequency in American populations (AF=0.28) with the highest PBS signal among Peruvians (AF=0.39) but is rarely observed in other populations (AF≤0.04). Further studies would be needed to confirm functional effect; however, it is interesting to note that type 1 diabetes in Peruvians is among the highest in the world (*53*).

## DISCUSSION

We have generated a diversity panel of phased long-read human genome assemblies that has significantly improved SV discovery and will serve as the basis to construct new population-specific references. The work begins to fill an important gap in our understanding of normal patterns of human genetic variation. Previous large-scale efforts have largely been inferential and biased when it comes to the detection of SVs. Here, we develop a method to discover all forms of genetic variation (PAV) directly by comparison of assembled human genomes. In contrast, SV discovery from the 1000GP was indirect and limited given the frequent proximity of SVs to repeat sequences inaccessible to short reads (*15, 21*). The 1000GP, for example, reported 69,000 SVs based on the analysis of 2,504 short-read sequenced genomes. In contrast, our analysis of 32 genomes (64 unrelated haplotypes) recovers 107,136 SVs. Recent large-scale studies of extended short-read-based cohorts (*5, 6*), interrogating tens of thousands of samples for SVs, typically reporting 5,000 to 10,000 SVs per sample, while our assembly-based SV calls identify 23,000 to 28,000 SVs per sample. This lack of sensitivity for SV discovery from short reads also affects common variation (AF>5%) and we increase the amount of common SVs by 2.6-fold. Notably, all forms of genetic variation we discover are physically phased with their flanking SNVs allowing haplotypes to be constructed where SVs are fully integrated. We take advantage of this information to apply a graph-based approach (PanGenie) to generate extremely accurate SV genotypes across the 1000GP to discover novel eQTLs and associations with human genetic disease and loci for adaptive evolution.

Application of both HiFi and CLR long-read technologies on the same samples allowed us to compare their performance. While HiFi sequencing was more expensive (on average 6 HiFi vs. 2 SMRT cells in this study), HiFi assemblies are more accurate. CLR-based assemblies are more contiguous and resolve the complex 3q29 locus (**Fig. 4C**) more often (35% or 21/60) than HiFi-based assemblies (18% or 5/28). As sequencing technology and assembly algorithms continue to evolve (*54, 55*), HiFi sequencing has been predicted to predominate because accuracy and higher depth will afford access to even more previously inaccessible regions (*54, 55*). Nevertheless, orthogonal technologies such as optical maps and Strand-seq data are still required. Strand-seq, for example, was critical to achieve chromosome-length phasing in the absence of parental sequencing data (*3*). Given the challenges of broadly obtaining parental material from populations as well as patients, trio-free haplotype-resolved assembly will be a major asset for further expanding human genome diversity and the discovery of more complex forms of pathogenic variation (*56, 57*).

Complete sequence resolution of SVs provides new insights into their mechanism of formation. Compared to previous reports based on short-read sequencing (*22*–*24*), a surprising finding has been the larger fraction of SVs (63%) now assigned to homology-based (>50 bp) mutation mechanisms, including HDR, NAHR and VNTR. Breakpoint characterization with short-read data apparently biased early reports toward relatively unique regions concluding that <30% of SVs were driven by homology-based mutational mechanisms (*22*–*24*). Since a majority of unresolved structural variation still maps to large repeats, including centromeres and SDs subject to NAHR, we conclude that homology-based mutational mechanisms will contribute even further and are, therefore, the most predominant mode shaping the SV germline mutational landscape. Notwithstanding, access to fully assembled retrotransposons and their flanking sequence provides the largest collection of annotated source elements for both L1 and SVA mobile elements. We find that 14% of SVA insertions are associated with transductions compared to 8% of L1s—a difference driven in part by the proclivity of SVAs to transduce sequences at their 5’ and 3’ ends. We find a surprisingly large number of L1 source elements (19%) with defective ORFs suggesting either trans-complementation (*58*) or polymorphisms leading to the recent demise of these active source elements. Of note, some of the youngest L1 copies (e.g., 6p22.1-1 and 2q24.1) have been reported to be rare polymorphisms able to mediate massive bursts of somatic retrotransposition in cancer genomes (*59*). This suggests that recently acquired hot L1s, which have not yet reached an equilibrium with our species, contribute disproportionately to disease-causing variation (*60*).

Genome-wide eQTL scans can bridge the gap between molecular and clinical phenotypes and serve as a proxy for functional effects mediated by genetic variant classes (*21, 43, 61*). Taking advantage of the fully phased sequence-resolved genetic variation, we demonstrate this by applying PanGenie, a new pangenome-based genotyping method, to 3,202 1000GP genomes, resulting in reliable genotype calls for 705,893 indels and up to 50,340 SVs (lenient genotype set). Of these, 59.9% are presently missed in multi-algorithm short-read discovery callsets and the majority (68.2%) of these novel SVs are insertions. Our work, thus, provides a framework for the discovery of eQTLs and disease-associated variants with the potential to discriminate among SNVs, indels, and SVs as the most likely causal variants (lead variants) associated with human genetic traits. The fact that 31.9% of SV-eQTLs and 48% of lead SV-eQTLs are rendered accessible to short reads only through the availability of our panel of haplotype-resolved assemblies testifies to the importance of this resource for future GWAS. Once again, among the lead SV-eQTLs, 75% are insertions although there are also promising deletion eQTLs. For example, we identify a 1,069 bp deletion eQTL near *LIPI*, a GWAS disease locus for cardiac failure (*45*). Indeed, summary-data-based Mendelian randomization analysis (SMR, (*62*)) suggests that this SV-eQTLs of *LIPI* may be driving this association (SMR p-value adj.: 5.6e-4). Further analysis (Supplementary Information) using known GWAS summary statistics (p-value <= 1.0e-6 extracted from the GWAS Catalog (*63*), PhenoScanner (*64, 65*), Pan-UKB project, and UK Biobank (Pan-UKB team https://pan.ukbb.broadinstitute.org2020) highlights 1,178 genes associated with GWAS traits whose expression changes are significantly associated with SVs and 4,494 genes are associated with indels. We note that 17 of these genes have an SV as the lead eQTL and 377 genes have a lead indel-eQTL and represent promising candidates for future exploration of disease association.

Haplotype-resolved SVs with accurate genotypes will also facilitate evolutionary and population genetic studies of SVs, including estimations of the rates of recurrent mutation, population stratification, and selective sweeps. As part of this analysis, we identify 117 loci associated with genes where allele frequencies differ radically between populations and are candidates for local adaptation (*49, 50*). Ancestral reconstructions of haplotype-resolved SVs can be further extended to identify introgressed SVs from Neandertals and Denisova (*66*). While archaic SNV haplotypes have been identified in modern-day humans, little is known regarding SV content given the degraded nature of ancient DNA. Combined with coalescent estimates of evolutionary age, it should now be possible to systematically identify associated introgressed SVs and assess them for signatures of adaptive evolution as was recently demonstrated (*67*). Even though we estimate that 96% of SVs with an allele frequency above 2% have been theoretically discovered, a greater diversity of human genomes are required and our findings clearly indicate that genomes of African ancestry represent the deepest reservoir of untapped structural variation. Ongoing efforts from the HGSVC, *All of Us*, and the Human Pangenome Reference Consortium (HPRC, https://humanpangenome.org) exploring the normal pattern of structural variation using long-read sequences over the next few years will be critical in better understanding of human genetic variation.

Currently, our understanding of the full spectrum of structural variation is not yet complete, despite the advances presented here. There are two important limitations. First, comparison with optical mapping data identifies hundreds of gene-rich regions near and within SDs harboring more complex forms of SVs that are still not fully resolved by whole-genome long-read sequencing. The remaining gaps in human genomes cluster and a subset represent complex SV differences between human haplotypes. Second, only ∼50% of our long-read discovery set of SVs can, at present, be reliably genotyped in short-read data using PanGenie. Expanding the number of assembly-based haplotypes available as pangenomic reference will likely mitigate this, but multiallelic VNTRs/STRs as well as SVs embedded in larger repeats such as SDs and centromeres are particularly problematic and novel methods are needed to characterize these. Recent advances coupling both HiFi and ultra-long-read Oxford Nanopore data show promise in resolving the sequence of these more complex regions from both haploid (*68*) and diploid human genome assemblies (*69*). Once a larger number of such complex regions are haplotype resolved across diversity panels of human genomes—and algorithms continue to evolve to exploit this information—we expect larger portions of the human genome to become amenable to genotyping and association with human traits.

## METHODS (short)

Libraries were prepared from high-molecular-weight DNA from lymphoblast lines (Coriell Institute). Long-read CLR and HiFi sequencing data (25-50X) were generated on the Sequel II platform (Pacific Biosciences) using 15-hour (CLR) or 30-hour (HiFi) movie times. Strand-seq data were produced from the same samples and used to identify and phase heterozygous SNVs (LongShot (*70*) and DeepVariant (*71*)) from the squashed genome assemblies (Peregrine or Flye). StrandphaseR (*72*), SaaRclust (*73*) and WhatsHap (*74, 75*) partitioned long reads into haplotypes to generate phased genome assemblies (PGAS). MAPQ60 phased assembly contig coverage is estimated for autosomes (chr 1-22) and the X chromosome to balance male and female comparisons, excluding regions of heterochromatin (Giemsa pos./var. staining) and unresolved reference sequence (N-gaps). We generated optical maps for 30 of the 32 samples based on *DLE*1 digestion (Bionano Genomics). PAV was used to characterize SNVs, indels, and SVs compared to the human reference GRCh38. Inversions were detected using Strand-seq (*1, 9, 36*), optical mapping data (Bionano Solve v3.5) and PAV, which detects inversion signatures using a novel k-mer density approach to identify inner and outer breakpoints of flanking repeats without relying on alignment truncation. The diploid callset is created by merging two independent haploid callsets. We removed variants in collapses by SDA (*76*) and misaligned contig clusters then merged variants from all samples to create a nonredundant callset that was subsequently filtered by additional support (Supplementary Methods). SVs required support from at least one of seven other sources including read-based callers (MELT, PBSV, PALMER) (*31, 77*), optical mapping data, breakpoint k-mer analysis, and PAV replication with LRA (github.com/ChaissonLab/LRA) (Supplementary Methods). Indels required support from at least two of four sources and SNVs required support from at least two of five sources. In the PAV callset, we fully sequence resolved 9,950 non-reference MEIs, including 8,110 Alus, 1,248 L1s, 589 SVAs, and three HERV-Ks. Applying read-based callers (MELT and PALMER), we discovered an additional 1,932 putative MEIs (although not all were fully sequence resolved). Combining all methods, we discovered 11,882 MEIs, including 9,516 Alus, 1,646 L1Hs, 688 SVAs, and 32 HERV-Ks. We estimated functional element depletion for SVs by simulation permuting SVs within their 1 Mbp bin 100,000 times and recording functional element hits for insertions and deletions for each functional category (CDS, 5’ UTR, 3’ UTR, promoter, proximal enhancer, distal enhancer, CTCF, and intron). SV hotspots were defined by searching for regions of increased SV density using kernel density estimation implemented with the ‘hotspotter’ function from the primatR package (*36, 78*). Illumina WGS short reads (250 bp paired end) were generated (34.5-fold) (Supplementary Information) from 1000GP samples (2,504 unrelated individuals and additional samples from children to form 602 trios). SVs were called from an ensemble of three methods: GATK-SV (*5*), SVTools (*6*) and Absinthe (github.com/nygenome/absinthe) and detailed comparisons between long-read and short-read data were performed for the 34 matched samples (Supplementary Methods). We genotyped all 3,202 genomes using PanGenie, which determines k-mer abundances from an input set of unaligned short reads and infers the genotypes of this short-read sample at all loci represented in the reference set. The method exploits both the linkage disequilibrium structure inherent to the reference haplotypes and the sequence resolution they provide, and hence makes full use of the haplotype resource provided. RNA-seq data QC was conducted with Trim Galore! (*79*) and mapped to the reference genome using STAR (*80*), followed by gene-level quantification using FeatureCounts (*81*). We mapped the effect of genetic variation on expression levels using an eQTL mapping pipeline based on a linear mixed model implemented in LIMIX (*82*–*84*). We combined our eQTL statistics with published GWAS associations to assess the link among genetic variation, gene expression and associated traits using SMR (*62*). To identify population-stratified SVs in the 26 populations, we computed the FST-based PBS statistic (Supplementary Information). For each focal population, we constructed population triplets by choosing sister- and out-groups inside and outside the continent where the focal population resides, respectively. For each focal population, we selected the maximum PBS per gene for all possible PBS triplets and selected the subset that are at least 3 standard deviations (Z transformation) beyond the PBS mean as potential targets of selection.

## Acknowledgements

We thank T. Brown for assistance in editing this manuscript and K. Hoekzema and C. Baker for the preparation of DNA from cell lines. We also recognize the computational support (P.H. Rehs and C. Siebert) and infrastructure provided by the Centre for Information and Media Technology (ZIM) at the University of Düsseldorf, the EMBL IT Services, and additional computational analyses (C. Alkan, F. Hormozdiari, D.S. Gordon and S. Murali). We thank M. Paulsen from the EMBL Flow Cytometry Core Facility, as well as J. Zimmermann and V. Benes from the EMBL Genomics Core Facility for assisting in Strand-seq sample preparation and sequencing. We thank the Human Pangenome Reference Consortium for use of the publicly available GIAB sequence data for the Ashkenazim benchmark sample HG002/NA24385. We are grateful to the people who generously contributed samples as part of the 1000 Genomes Project (1000GP). We thank the Pan-UKB project and UK Biobank for making the GWAS results available.

## Authors contributions

PacBio production sequencing: K.M.M., A.P.L., Q.Z., L.J.T., S.E.D. Strand-seq production: A.D.S., B.R., P.H., J.O.K. Phased genome assembly: P.E., P.A.A., D.P., Q.Z., F.Y., W.T.H., T.M. Assembly analysis: P.E. Assembly-based variant calling: P.A. Variant QC, merging, and annotation: P.A.A., T.R., M.J.P.C., J.R., Z.C., Y.C., K.Y., J.L., X.Y., J.O.K. Assembly scaffolding: F.Y., D.P., P.E. Additional long-read callsets: P.A.A., Y.C., Z.C., W.T.H., J.R., A.M.W. Short-read SV calling and merging: X.Z., Q.Z., H.A., H.B., N.T.C., W.E.C., A.C., S.E.D., I.H., W.T.H., A.R., M.C.Z., M.E.T. Bionano Genomics SV discovery and analysis: F.Y., J.L., A.R.H. Strand-seq inversion detection and genotyping: D.P., W.T.H., H.A., M.G., T.M., A.D.S., J.O.K. MEI discovery and integration: B.R.-M., W.Z., M.S., N.T.C., J.M.C.T., J.O.K., R.E.M., S.E.D. Variant hotspot analysis: D.P., E.E.E. Breakpoint analysis: S.K., J.L., X.Y., M.G., K.Y., J.O.K. PanGenie genotyping: J.E., T.M. Illumina genotype analysis: J.E., X.Z., W.E.C., P.E., T.R., P.A.A., H.B., J.O.K., M.E.T., M.C.Z., T.M. RNA-seq and eQTL analysis: M.J.B., A.S., Z.M., J.C., C.L., M.B.-B., A.O.B., O.S., Y.I.L., X.S., M.C.Z., J.O.K. Ancestry and population genetic analyses: PH.H., R.S.M., P.A.A., T.M., E.E.E. Data archiving: S.F., P.A.A., K.M.M., P.F. Organization of supplementary materials: Q.Z. and C.L. Display items: P.A.A., P.E., J.E., A.R.H., PH.H., R.S.M., T.M., D.P., T.R., B.R.-M., M.S., F.Y., X.Z., W.Z. Manuscript writing: P.A.A., P.E., B.R.-M., A.S., D.P., PH.H., Q.Z., F.Y., A.R.H., J.L., M.T., M.J.B., X.S., S.E.D., J.O.K., T.M., E.E.E. HGSVC Co-chairs: C.L., J.O.K., E.E.E.

## Data and materials availability

All data generated was made immediately publicly available via the International Genome Sample Resource (IGSR) (www.internationalgenome.org) at ftp.1000genomes.ebi.ac.uk/vol1/ftp/data_collections/HGSVC2/. Data are available at INSDC under the following accessions and project IDs: Illumina high-coverage genomic sequence (PRJEB37677), HiC and RNA-seq (ERP123231), Bionano Genomics (ERP124807), PacBio (PRJEB36100 and pending accession), and Strand-seq (PRJEB39750). The merged callsets are available via Zenodo (10.5281/zenodo.4268828). URLs for all data are available in **table S4**. The following cell lines/DNA samples were obtained from the NIGMS Human Genetic Cell Repository at the Coriell Institute for Medical Research: [NA06984, NA06985, NA06986, NA06989, NA06991, NA06993, NA06994, NA06995, NA06997, NA07000, NA07014, NA07019, NA07022, NA07029, NA07031, NA07034, NA07037, NA07045, NA07048, NA07051, NA07055, NA07056, NA07340, NA07345, NA07346, NA07347, NA07348, NA07349, NA07357, NA07435, NA10830, NA10831, NA10835, NA10836, NA10837, NA10838, NA10839, NA10840, NA10842, NA10843, NA10845, NA10846, NA10847, NA10850, NA10851, NA10852, NA10853, NA10854, NA10855, NA10856, NA10857, NA10859, NA10860, NA10861, NA10863, NA10864, NA10865, NA11829, NA11830, NA11831, NA11832, NA11839, NA11840, NA11843, NA11881, NA11882, NA11891, NA11892, NA11893, NA11894, NA11917, NA11918, NA11919, NA11920, NA11930, NA11931, NA11932, NA11933, NA11992, NA11993, NA11994, NA11995, NA12003, NA12004, NA12005, NA12006, NA12043, NA12044, NA12045, NA12046, NA12056, NA12057, NA12058, NA12144, NA12145, NA12146, NA12154, NA12155, NA12156, NA12234, NA12239, NA12248, NA12249, NA12264, NA12272, NA12273, NA12274, NA12275, NA12282, NA12283, NA12286, NA12287, NA12329, NA12335, NA12336, NA12340, NA12341, NA12342, NA12343, NA12344, NA12347, NA12348, NA12375, NA12376, NA12383, NA12386, NA12399, NA12400, NA12413, NA12414, NA12485, NA12489, NA12546, NA12707, NA12708, NA12716, NA12717, NA12718, NA12739, NA12740, NA12748, NA12749, NA12750, NA12751, NA12752, NA12753, NA12760, NA12761, NA12762, NA12763, NA12766, NA12767, NA12775, NA12776, NA12777, NA12778, NA12801, NA12802, NA12812, NA12813, NA12814, NA12815, NA12817, NA12818, NA12827, NA12828, NA12829, NA12830, NA12832, NA12842, NA12843, NA12864, NA12865, NA12872, NA12873, NA12874, NA12875, NA12877, NA12878, NA12889, NA12890, NA12891, NA12892].

## Supplementary Materials

Materials and Methods

Table S1 – S49

Fig S1 – S87

References (X – Y)

