## Supplemental Methods for "*De novo* assembly of 64 haplotype-resolved human genomes of diverse ancestry and integrated analysis of structural variation"

#### METHODS and SUPPLEMENTARY MATERIAL

Table of Contents

**DATA PRODUCTION**

**1. Samples and data availability** 4

**2. Long-read, whole-genome sequence production** 5

**3. Strand-seq production** 7

**4. Illumina sequencing** 8

**5. Bionano production** 8

**6. Hi-C data generation** 8

**7. RNA-seq data generation** 9

**VARIANT CALLING**

**8. Phased long-read genome assembly** 9
8.1. K-mer-based analysis of phased assemblies 11
8.2. Reference-based analysis of phased assemblies 12
8.3. Assembly Scaffolding 13
8.4. Analysis of phased assemblies 14

**9. Reference and global merging strategy** 17
9.1. Genome references 17
9.2. Variant merging strategies 18

**10. Phased assembly variant discovery** 18
10.1. Contig alignment and trimming. 18
10.2. Interalignment and alignment truncating variants 19
10.3. Inversion detection 20
10.4. Callset finishing 21
10.5. Post-PAV callset filters 22

**11. Variant discovery with aligned reads** 22
11.1. PBSV 22
11.2. DeepVariant 23
11.3. DeBreak 23

**12. Illumina SV calling** 24
**12.1. SV discovery from individual algorithms** 24
**12.2. SV integration** 25
12.2.1. FusorSV 25

12.2.2. GATK-SV 26
12.2.2.2 SV discovery from raw algorithms 27

**13. Bionano Genomics discovery 28**
13.1. Bionano Genomics de novo assembly and structural variant calling 28
13.2. Bionano Genomics discovery of large, complex structural variants 29

**14. Strand-seq Inversion detection and genotyping 30**

**15. MEI discovery and integration 32**
15.1. Callsets across different platforms 33
15.1.1. MELT 33
15.1.2. PALMER 33
15.1.3. MEIGA 34
15.2. MEI integration and PAV annotation 35
15.3. Characteristics of MEIs and analysis 36
15.3.1. Sequence-resolved full-length L1s 36
15.3.2. Phylogenetic analysis and age estimation for active sequence-resolved L1s 38
15.3.3. A map of active source SVA loci in diverse populations 38
15.3.4. VNTR distributions in SVAs 39
15.3.5. Distributions of poly(A) tract and EN cleavage site sequences 41

**16. Nonredundant callsets, filtering, and properties 41**
16.1. Nonredundant callset merging 41
16.2. Post-merge filtering 43
16.3. Callset quality estimates 44
16.4. Variant frequency distributions 44
16.5. Identify SV clusters - hotspot analysis 44
16.6. HLA analysis 45
16.7 Shared variants 45

**VALIDATION**

**17. SV validation by reads and contigs 46**
17.1. Subseq validations with raw reads 46
17.2. LRA read and contig alignment support 46
17.3. Comparison of phase 1 and phase 2 callsets. 47
17.4. Variant concordance by reads, contigs and assemblies (Inspector) 48
17.5. Raw read variant and breakpoint concordance. 48
17.6. Breakpoint analysis 49

**18. Analyzing non-reference k-mers using 3,202 genomes 51**

#### GENOTYPING AND ASSOCIATION ANALYSIS

**19. Genotyping Illumina Genomes** **52**
19.1. Genotyping Paragraph 52
19.2. Genotyping PanGenie and graph construction 52

**20. RNA-seq analysis** **56**
20.1. Read QC and mapping 56
20.2. Expression and splicing quantification and normalization 57

**21. eQTL and GWAS** **57**
21.1. eQTL and sQTL analyses 57
21.2. GWAS intersection and enrichment analysis 59
21.3. GWAS and eQTL co-localization analysis 60

**22. Ancestry Analysis** **61**
22.1 Local ancestry 61
22.2 Variant age estimations 63
22.3 Ancestral state determination from primate assemblies 63
22.4 Population Stratification and Population Branch Statistics 64

**23. Functional Annotations** **64**
23.1. Functional variant annotations 64
23.2 Triplet repeat expansions 64

**24. References** **65**

### DATA PRODUCTION

#### 1. Samples and data availability

**(*Contributors: Katy Munson, Qihui Zhu and Scott Devine*)**

Samples were selected from the 1000 Genomes Project (1000GP) Diversity Panel (*1*) and at least one representative was selected from each of 26 populations (Table S1). Cell lines were obtained from Coriell for data production. Existing sequencing data for NA12878 and NA24385/HG002 from the Genome in a Bottle (GIAB) effort (*2*) were also added to the final dataset. Table S2 provides the sample statistics of the project and Table S3 provides a list of samples generated from each technology.

The links to the raw data generated in this study can be found in the Table S4.

The data flowchart can be found in Fig. S1.

***Table S1. PacBio sequencing and phased assembly summary***

Summary statistics for sequencing datasets input to phased assembly pipeline and for "squashed" and haplotype assemblies per sample (family trios listed for both HiFi and CLR). All assembly statistics pertain to the pipeline parameterization v12. Column mean and median are given per long-read technology and for the entire dataset (bottom).

| Sample Information | | | | | Sequencing Statistics * | | | | Strand-seq ‡ | Assembly Statistics | | | | | |
|---|---|---|---|---|---|---|---|---|---|---|---|---|---|---|---|
| | | | | | Yield | Read Length | | Read Quality † | | Squashed | | Phased | | | |
| Sample ID | Family ID | Population | Super Population | Gender | Gbp | Mean (kbp) | N50 (kbp) | Median (Q) | Libraries (Count) | Length (Gbp) | N50 (Mbp) | H1 Length (Gbp) | H1 N50 (Mbp) | H2 Length (Gbp) | H2 N50 (Mbp) |
| HiFi | | | | | | | | | | | | | | | |
| NA19238 | Y117 | YRI | AFR | Female | 78 | 12.4 | 12.2 | 30 | 172 | 3.03 | 32.1 | 3.01 | 18.0 | 3.00 | 17.4 |
| NA19239 | Y117 | YRI | AFR | Male | 81 | 14.0 | 14.1 | 29 | 183 | 3.07 | 29.1 | 3.08 | 16.4 | 3.08 | 17.4 |
| NA19240 | Y117 | YRI | AFR | Female | 89 | 12.9 | 12.9 | 30 | 117 | 3.03 | 30.6 | 3.03 | 23.7 | 3.02 | 21.3 |
| HG00731 | PR05 | PUR | AMR | Male | 103 | 11.1 | 10.6 | 32 | 122 | 3.04 | 28.5 | 3.06 | 19.5 | 3.05 | 17.5 |
| HG00732 | PR05 | PUR | AMR | Female | 76 | 22.9 | 22.9 | 28 | 94 | 3.11 | 30.9 | 3.08 | 19.5 | 3.07 | 25.2 |
| HG00733 | PR05 | PUR | AMR | Female | 104 | 13.6 | 13.5 | 30 | 115 | 3.03 | 33.4 | 3.05 | 25.7 | 3.05 | 25.7 |
| HG00512 | SH032 | CHS | EAS | Male | 93 | 15.4 | 15.6 | 30 | 111 | 3.07 | 32.2 | 3.10 | 25.5 | 3.08 | 26.5 |
| HG00513 | SH032 | CHS | EAS | Female | 93 | 15.4 | 15.6 | 30 | 74 | 3.05 | 32.1 | 3.05 | 25.8 | 3.05 | 28.2 |
| HG00514 | SH032 | CHS | EAS | Female | 76 | 17.0 | 17.1 | 29 | 76 | 3.04 | 29.3 | 3.04 | 18.4 | 3.05 | 18.4 |
| NA24385 | 3140 | Ashk. | EUR | Male | 111 | 15.0 | 14.7 | 33 | 65 | 3.08 | 33.4 | 3.09 | 31.3 | 3.09 | 25.4 |
| NA12878 | 1463 | CEU | EUR | Female | 90 | 10.0 | 10.0 | 33 | 132 | 2.98 | 24.3 | 2.98 | 18.3 | 2.98 | 20.8 |
| HG02818 | GB66 | GWD | AFR | Female | 114 | 16.5 | 16.8 | 31 | 70 | 3.08 | 21.9 | 3.08 | 15.0 | 3.08 | 14.3 |
| HG03125 | NG34 | ESN | AFR | Female | 85 | 17.8 | 17.4 | 30 | 56 | 3.07 | 32.1 | 3.05 | 19.6 | 3.05 | 17.1 |
| HG03486 | SL61 | MSL | AFR | Female | 122 | 19.1 | 18.7 | 30 | 62 | 3.11 | 25.8 | 3.10 | 17.7 | 3.10 | 17.3 |
| | | | | **Mean:** | 94 | 15.2 | 15.1 | 30 | 104 | 3.06 | 29.7 | 3.06 | 21.0 | 3.05 | 20.9 |
| | | | | **Median:** | 91 | 15.2 | 15.1 | 30 | 103 | 3.06 | 30.7 | 3.05 | 19.5 | 3.05 | 19.6 |
| CLR | | | | | | | | | | | | | | | |
| NA19238 | Y117 | YRI | AFR | Female | 271 | 30.1 | 49.2 | - | 172 | 2.84 | 35.7 | 2.86 | 28.1 | 2.85 | 26.7 |
| NA19239 | Y117 | YRI | AFR | Male | 305 | 21.2 | 33.8 | - | 183 | 2.83 | 24.7 | 2.86 | 28.1 | 2.86 | 27.6 |
| NA19240 | Y117 | YRI | AFR | Female | 241 | 19.7 | 31.6 | - | 117 | 2.84 | 31.9 | 2.85 | 26.7 | 2.85 | 28.1 |
| HG00731 | PR05 | PUR | AMR | Male | 363 | 25.8 | 40.7 | - | 122 | 2.84 | 28.3 | 2.86 | 33.5 | 2.86 | 30.6 |
| HG00732 | PR05 | PUR | AMR | Female | 342 | 28.6 | 44.9 | - | 94 | 2.84 | 38.6 | 2.75 | 30.7 | 2.84 | 29.7 |
| HG00733 | PR05 | PUR | AMR | Female | 321 | 30.6 | 45.8 | - | 115 | 2.84 | 38.4 | 2.84 | 33.3 | 2.84 | 35.6 |
| HG00512 | SH032 | CHS | EAS | Male | 199 | 26.6 | 46.4 | - | 111 | 2.83 | 29.9 | 2.86 | 32.8 | 2.86 | 33.0 |
| HG00513 | SH032 | CHS | EAS | Female | 260 | 24.6 | 42.4 | - | 74 | 2.81 | 27.3 | 2.85 | 32.6 | 2.85 | 32.6 |
| HG00514 | SH032 | CHS | EAS | Female | 185 | 19.3 | 34.3 | - | 76 | 2.84 | 28.6 | 2.83 | 20.3 | 2.84 | 18.6 |
| HG02011 | BB13 | ACB | AFR | Male | 321 | 28.9 | 45.4 | - | 56 | 2.84 | 28.1 | 2.86 | 31.9 | 2.86 | 33.8 |
| HG02587 | GB24 | GWD | AFR | Female | 257 | 16.6 | 31.1 | - | 54 | 2.81 | 25.7 | 2.84 | 20.0 | 2.85 | 21.1 |
| HG03065 | SL05 | MSL | AFR | Male | 339 | 27.1 | 43.0 | - | 57 | 2.83 | 27.5 | 2.86 | 32.4 | 2.86 | 30.5 |
| HG03371 | NG98 | ESN | AFR | Male | 315 | 28.0 | 44.9 | - | 62 | 2.84 | 27.6 | 2.86 | 31.9 | 2.86 | 33.0 |
| NA19983 | 2436 | ASW | AFR | Female | 403 | 28.4 | 43.9 | - | 47 | 2.82 | 29.2 | 2.84 | 33.7 | 2.84 | 39.4 |
| HG01114 | CLM03 | CLM | AMR | Female | 243 | 15.6 | 24.9 | - | 53 | 2.84 | 29.4 | 2.83 | 16.8 | 2.83 | 19.3 |
| NA19650 | M001 | MXL | AMR | Male | 149 | 24.7 | 39.6 | - | 56 | 2.86 | 29.2 | 2.85 | 23.4 | 2.85 | 24.7 |
| HG00864 | - | CDX | EAS | Female | 230 | 26.5 | 42.2 | - | 43 | 2.84 | 33.7 | 2.85 | 27.0 | 2.85 | 26.5 |
| HG01596 | - | KHV | EAS | Male | 281 | 17.8 | 28.4 | - | 53 | 2.85 | 26.6 | 2.85 | 19.0 | 2.86 | 17.7 |
| NA18534 | - | CHB | EAS | Male | 202 | 23.3 | 39.7 | - | 57 | 2.85 | 36.9 | 2.85 | 25.7 | 2.85 | 26.7 |
| NA18939 | - | JPT | EAS | Female | 275 | 26.8 | 42.8 | - | 43 | 2.82 | 31.1 | 2.85 | 29.1 | 2.85 | 31.9 |
| HG00096 | - | GBR | EUR | Male | 233 | 27.6 | 44.8 | - | 69 | 2.85 | 34.4 | 2.87 | 28.8 | 2.87 | 28.2 |
| HG00171 | - | FIN | EUR | Female | 310 | 21.0 | 33.2 | - | 50 | 2.82 | 26.6 | 2.84 | 29.3 | 2.85 | 27.6 |
| HG01505 | IBS002 | IBS | EUR | Male | 222 | 21.9 | 40.2 | - | 64 | 2.85 | 33.5 | 2.87 | 26.8 | 2.86 | 28.5 |
| NA12329 | 1328 | CEU | EUR | Female | 372 | 25.6 | 40.9 | - | 61 | 2.85 | 35.6 | 2.85 | 30.4 | 2.85 | 29.8 |
| NA20509 | - | TSI | EUR | Male | 159 | 23.4 | 43.3 | - | 41 | 2.86 | 28.3 | 2.85 | 27.2 | 2.85 | 25.3 |
| HG02492 | PK06 | PJL | SAS | Male | 320 | 30.0 | 46.9 | - | 41 | 2.85 | 37.0 | 2.86 | 33.8 | 2.86 | 34.2 |
| HG03009 | BD01 | BEB | SAS | Male | 339 | 24.2 | 40.2 | - | 57 | 2.84 | 29.0 | 2.85 | 27.7 | 2.85 | 26.4 |
| HG03683 | ST12 | STU | SAS | Female | 363 | 25.9 | 40.4 | - | 51 | 2.84 | 33.5 | 2.85 | 32.0 | 2.85 | 33.6 |
| HG03732 | IT003 | ITU | SAS | Male | 200 | 25.6 | 41.4 | - | 50 | 2.83 | 31.9 | 2.85 | 29.0 | 2.85 | 26.3 |
| NA20847 | - | GIH | SAS | Female | 175 | 29.1 | 47.4 | - | 58 | 2.82 | 28.9 | 2.84 | 28.8 | 2.84 | 26.4 |
| | | | | **Mean:** | 273 | 24.8 | 40.5 | | 73 | 2.84 | 30.9 | 2.85 | 28.4 | 2.85 | 28.4 |
| | | | | **Median:** | 273 | 25.7 | 41.8 | | 57 | 2.84 | 29.3 | 2.85 | 28.9 | 2.85 | 28.1 |
| | | | | **Mean:** | 216 | 21.8 | 32.4 | | 83 | 2.91 | 30.5 | 2.91 | 26.0 | 2.92 | 26.0 |
| | | | | **Median:** | 230 | 23.3 | 39.7 | | 64 | 2.84 | 29.4 | 2.86 | 27.2 | 2.86 | 26.5 |

* Data reported is HiFi reads (estimated QV ≥ Q20) for HiFi data sets and raw subreads for CLR data sets

† Read quality is not reported for raw CLR subreads

‡ The same Strand-seq libraries were used for both HiFi and CLR assemblies for samples where both data types were generated

### 2. Long-read, whole-genome sequence production

We generated and analyzed PacBio High-Fidelity (HiFi)/circular consensus sequencing (CCS) for 9/38 samples and continuous long-read (CLR) data for 30/38 samples across three centers. Among them, three trios were sequenced by both HiFi/CCS and CLR.

Sample quality control (QC) and statistics can be found in Table S5. In addition, existing datasets for five samples sequenced as part of the GIAB or Human Genome Reference consortium were obtained from publicly available sources (Table S6).

(***Contributors: Katy Munson, Qihui Zhu, Scott Devine and Alex Lewis***)

***University of Washington*** — Genomic DNA was isolated as previously described (*3*) and evaluated for purity and quantity using UV-Vis (Nanodrop 1000, Thermo Fisher) and fluorometric (Qubit, Thermo Fisher) assays. DNA sizing was checked on the FEMTO Pulse (Agilent) using the Genomic DNA 165 kb kit on extended mode. Samples all exhibited mode size above 50 kbp (most above 100 kbp) and were considered good candidates for PacBio sequencing. For HiFi/CCS, three parent–child trios (9 individuals) were selected and DNA was sheared using gTUBEs (Covaris) to a mode size ~15 kbp using 3200 RPM for four passes on an Eppendorf 5424R centrifuge. The sheared material was subjected to SMRTbell library preparation using the Template Prep Kit v1 (PacBio). After checking for size and quantity, the material was size fractionated on the SageELF instrument (Sage Science) using the protocol "0.75% 1-18kb v2", size-based separation mode, and target value 3400 in well #12. Fractions were checked via fluorometric quantitation (Qubit) and pulse-field sizing (FEMTO Pulse). Additional libraries were prepared by g-TUBE shearing to 20 kbp using 4000 rpm (1500 rcf) in an Eppendorf 5415R centrifuge for six passes, followed by SMRTbell prep using the SMRTbell Express Template Prep Kit 2.0 kit and Enzyme Cleanup Kit (PacBio). These libraries were size fractionated on SageELF using a custom protocol "Waveform 250-100" timed mode, 3-hour run time. All cells were sequenced on a Sequel II instrument (PacBio) using 30-hour movie times. Fractions averaging roughly 11 and 14 kbp were prepared using version 1.0 sequencing chemistry and 2-hour pre-extension. Fractions averaging roughly 18 kbp were prepared with version 2EA sequencing chemistry and 4-hour pre-extension. Fractions averaging roughly 20 kbp were prepared with version 2.0 sequencing chemistry and 4-hour pre-extension. HiFi/CCS analysis was performed using SMRT Link v7.0, v7.1, v8.0, or v9.0, using filters of three full passes and an estimated read-quality value of 0.99. For CLR sequencing, without further shearing, isolated gDNA was SMRTbell library prepped using the Express Kit v2 (PacBio) and subjected to size selection on a BluePippin instrument (Sage Science) with a 40 kbp size cutoff. Libraries were loaded on a Sequel II using v2.0 binding and v2.0 sequencing kits, no pre-extension, and 15-hour movie times.

***The Jackson Laboratory*** — High-molecular-weight DNA was extracted from frozen pelleted cells using the phenol-chloroform approach as previously described (*4*). Purified gDNA was assessed using fluorometric (Qubit, Thermo Fisher) assays for quantity and FEMTO Pulse (Agilent) on quality. For HiFi/CCS sequencing, samples exhibiting a mode size above 50 kbp were considered good candidates. DNA was sheared using gTUBEs (Covaris) to target 12 kbp fragments using the centrifugation settings at 5500 RPM for 60s. The sheared material was subjected to SMRTbell library preparation using the Template Prep Kit v1 (PacBio). After checking for size and quantity, the material was size fractionated on the SageELF (Sage Science) using the protocol "0.75% 1-18kb v2", size-based separation mode and 3400 in

Targae. Fractions were checked via fluorometric quantitation (Qubit) and pulse-field sizing (FEMTO Pulse). All libraries were sequenced on a Sequel II (PacBio) using 30-hour movie times. Fractions between 9 to 13 kbp were selected for sequencing using sequencing chemistry v1.0, 2-hour pre-extension and 30-hour movie times. HiFi/CCS analysis was performed using SMRT Link v7.0 or v7.1 by an estimated read-quality cutoff of 0.99. For CLR sequencing, samples exhibiting a mode size above 50 kbp were considered good candidates to proceed for sequencing. gDNA was sheared with passing through a needle, and the SMRTbell library was prepped using the Express Kit v2 (PacBio) and subjected to size selection on a BluePippin (Sage Science) with a 15 and 30 kbp size cutoff. Libraries were loaded on a Sequel II using v1.0/v2.0 binding and v1.0/v2.0 sequencing kits, no pre-extension, and 20-hour movie times.

***University of Maryland*** — High-molecular-weight genomic DNA (gDNA) was prepared from cell pellets provided by Coriell Institute. The cells were lysed in a buffer containing proteinase K at 50°C overnight, followed by phenol/chloroform extraction and ethanol precipitation. The resulting gDNA typically showed a major peak at 165 kbp on a FEMTO pulsed-field run. For CLR sequencing, DNA samples were purified with SPRIselect beads (Beckman Coulter, Brea, CA) to remove small fragments and impurities prior to library preparation. Prior to sequencing, libraries were bound to polymerase with the Sequel II Binding Kit 2.0 (PacBio, Menlo Park, CA), and then sequenced with the Sequel II Sequencing Kit 2.0 and two 8M SMRT Cells on the Sequel II (PacBio, Menlo Park, CA) with 15-hour movie time. For HiFi/CCS sequencing, libraries were constructed using the SMRTbell Express Template Prep Kit 2.0 (PacBio, Menlo Park, CA) and fractionated on the SageELF (Sage Science, Beverly, MA) into narrow library fractions. Prior to sequencing, library fractions were bound to polymerase with the Sequel II Binding Kit 2.0 (PacBio, Menlo Park, CA), and then sequenced with the Sequel II Sequencing Kit 2.0 and 5-6 8M SMRT Cells on the Sequel II (PacBio, Menlo Park, CA) with 30-hour movie times.

### 3. Strand-seq production

**(*Contributors: Ashley Sanders)***

We generated Strand-seq data for 38/38 samples targeted in this study (Table S2). EBV-transformed lymphoblastoid cell lines (Coriell Institute) were cultured in BrdU (100 uM final concentration; Sigma, B5002) for 18 or 24 hours and single isolated nuclei (0.1% NP-40 lysis buffer (*5*)) were sorted into 96-well plates using the BD FACSMelody cell sorter. In each sorted plate, 94 single cells plus one 100-cell positive control and one 0-cell negative control were deposited. Strand-specific DNA sequencing libraries were generated using the previously described Strand-seq protocol (*5*, *6*) and automated on the Beckman Coulter Biomek FX P liquid handling robotic system (*7*). Following 15 rounds of PCR amplification, 288 individually barcoded libraries (amounting to three 96-well plates) were pooled for sequencing on the Illumina NextSeq5000 platform (MID-mode, 75 bp paired-end protocol). The demultiplexed FASTQ files were aligned to the GRCh38 reference assembly (BWA 0.7.15) for standard library selection. Low-quality libraries were excluded from future analyses if they showed low read counts, uneven coverage, or an excess of ‘background reads’ yielding noisy single-cell data, as previously described (Fig. S2) (*5*). Selected FASTQ files were used directly to guide the *de novo* reference-free assemblies, and the aligned BAM files were used for structural variant (SV) detection.

### 4. Illumina sequencing

**(*Contributor: Wayne Clarke and Michael Zody*)**

We generated and analyzed Illumina whole-genome sequencing (WGS) data for 38/38 samples (Table S2). WGS libraries were prepared using the TruSeq DNA PCR-Free Library Preparation Kit (Illumina) in accordance with the manufacturer’s instructions. Briefly, 1 ug of DNA was sheared using a Covaris LE220 sonicator (adaptive focused acoustics). DNA fragments underwent bead-based size selection and were subsequently end-repaired, adenylated, and ligated to Illumina sequencing adapters. Final libraries were evaluated using fluorescent-based assays, including qPCR with the Universal KAPA Library Quantification Kit and Fragment Analyzer (Advanced Analytics) or BioAnalyzer (Agilent 2100). Libraries were sequenced on an Illumina NovaSeq 6000 sequencer using 2 x 150 bp cycles to a minimum depth of 30X.

### 5. Bionano production

**(*Contributor: Alex Hastie)***

We generated and analyzed Bionano Genomics Optical Mapping data for 33/38 samples. Bionano data were previously generated for NA12878 and HG002, and not available for the three samples HG02818, HG03125 and HG03486 (Table S2). Cell lines were obtained from Coriell and grown in RPMI 1640 media with 15% FBS, supplemented with L-glutamine and penicillin/streptomycin, at 37°C and 5% $CO_2$. Ultra-high-molecular-weight DNA was extracted according to the Bionano Prep SP Fresh Cells DNA Isolation protocol, revision C (Document number 30257), using a Bionano SP Blood & Cell DNA Isolation Kit (catalog #80030). In short, 1.5 million cells were centrifuged and resuspended in a solution containing detergents, proteinase K, and RNase A. DNA was bound to a silica disk, washed, eluted, and homogenized via 1-hour end-over-end rotation at 15 rpm, followed by an overnight rest at room temperature. Isolated DNA was fluorescently tagged at motif CTTAAG by the enzyme DLE-1 and counter-stained using a Bionano Prep™ DNA Labeling Kit – DLS (catalog # 8005) according to the Bionano Prep Direct Label and Stain (DLS) Protocol, revision F (Document number 30206). Data collection was performed using Saphyr 2nd generation instruments (Part number 60325) and Instrument Control Software (ICS) version 4.9.19316.1.

### 6. Hi-C data generation

**(*Contributor: Qihui Zhu*)**

We generated and analyzed Hi-C data for 33/38 samples. Hi-C data were not available for the following five samples: NA12878, HG002, HG02818, HG03125 and HG03486 (Table S2). Lymphoblastoid cell lines were obtained from Coriell Cell Repositories and cultured in RPMI 1650 supplemented with 15% fetal bovine serum. Cells were maintained at 37°C in an atmosphere containing 5% carbon dioxide. Hi-C was performed with Phase Genomics Proximo Hi-C kits v3.0 following the manufacturer's protocol using 1.5 M cells as input. Libraries were sequenced at New York Genome Center (NYGC) on an Illumina NovaSeq 6000 in a paired-end 150 bp format.

#### 7. RNA-seq data generation

**(*Contributor: Qihui Zhu*)**

We generated and analyzed RNA-seq data for 33/38 samples. RNAseq data were not available for five samples: NA12878, HG002, HG02818, HG03125 and HG03486 (Table S2). Total RNA of cell pellets were isolated simultaneously using QIAGEN RNeasy Mini Kit according to the manufacturer's instructions. Briefly, each cell pellet (1 million cells) was homogenized and lysed in Buffer RLT Plus, supplemented with 1% $\beta$-mercaptoethanol. The lysate-containing RNA was purified using an RNeasy spin column, followed by an in-column DNase I treatment by incubating for 10 min at room temperature, and then washed. Finally, total RNA was eluted in 50 uL RNase-free water. RNA-seq libraries were prepared with 300 ng total RNA using KAPA RNA Hyperprep with RiboErase (Roche) according to manufacturer's instruction. First, ribosomal RNA was depleted using RiboErase. Purified RNA was then fragmented at 85°C for 6 mins, targeting fragments ranging 250-300 bp. Fragmented RNA was reverse transcribed with an incubation of 25°C for 10 mins, 42°C for 15 mins, and an inactivation step at 70°C for 15 mins. This was followed by a second strand synthesis and A-tailing at 16°C for 30 mins, 62°C for 10 min. The double-stranded cDNA A-tailed fragments were ligated with Illumina unique dual index adapters. Adapter-ligated cDNA fragments were then purified by washing with AMPure XP beads (Beckman). This was followed by 10 cycles of PCR amplification. The final library was cleaned up using AMPure XP beads. Quantification of libraries was performed using real-time qPCR (Thermo Fisher). Sequencing was performed on an Illumina NovaSeq platform generating paired end reads of 100 bp at The Jackson Laboratory for Genomic Medicine. The RNA-seq QC statistics can be found in Table S7.

### VARIANT CALLING

#### 8. Phased long-read genome assembly

**(*Contributor: Peter Ebert*)**

We applied a recently developed computational pipeline for phased genome assembly using Strand-seq (PGAS, Fig. 1; github.com/ptrebert/project-diploid-assembly; (*8*)) with minor modifications to produce fully phased diploid genome assemblies without dependency on parent-child trio data for both CLR and HiFi data sets:

*Filtering Strand-seq libraries*. Differences in the experimental protocols for preparing Strand-seq libraries between the three family trios (CHS, PUR, and YRI; Strand-seq data available under ENA accession PRJEB12849) and the other 26 individuals in this study required an initial filtering step for the Strand-seq libraries. Based on a previously described methodology (*5*), we excluded control probes, technical dropouts, and libraries of medium quality if the total read count was less than 50,000, which provided little information to guide the assembly clustering process (see Methods in (*8*)). The complete list of excluded libraries is available in Table S12.

*Generating squashed and phased assemblies*. The initial non-haplotype resolved ("squashed") and the final phased assemblies for PacBio HiFi/CCS reads were generated as previously described (*8*). For the squashed assembly of PacBio CLR reads, we used Flye v2.6 (*9*) or, in cases where Flye v2.6 could not assemble the input reads, Flye v2.7 was applied with otherwise identical parameters. The phased assemblies for all CLR samples were generated with Flye v2.7. Following developer recommendations, we set the parameter "`--asm-coverage`" to 50 to lower Flye's memory consumption during the initial assembly steps; notably, despite this setting, all reads are used for the final assembly. The target genome size was set to the genome size of GRCh38 for the squashed assemblies, and to the total length of all sequences contained in the squashed-assembly cluster for the per-cluster haplotype assembly (see Methods in (*8*)):

```
flye --pacbio-raw {reads} --asm-coverage 50
-g {genome_size} -t {threads} --out-dir {output}
```

*Polishing phased assemblies*. The final phased assemblies for PacBio HiFi/CCS reads were polished as previously described ((*8*)). For the PacBio CLR assemblies, we performed a single pass of polishing with gcpp v1.9.0 (https://github.com/PacificBiosciences/gcpp) and used pbmm2 v1.1.0 (https://github.com/PacificBiosciences/pbmm2) to generate the alignments of long reads to the assembled sequence for each cluster (see Methods in (*8*)):

```
pbmm2 align --log-level INFO --sort --sort-memory {sort_memory}M
--no-bai --alignment-threads {align_threads} --sort-threads
{sort_threads} --preset SUBREAD --min-length 5000 --sample
{individual} {input_contigs} {input_reads} {output}

gcpp --num-threads {threads} --algorithm=arrow  --log-level INFO
--log-file {log} --reference {input_contigs} --output
{out_fasta},{out_gff} {input_alignments}
```

*Generating haplotype read coverage tracks*. For each sample, we generated coverage tracks for all three fractions of haplotagged reads (haplotypes 1 and 2, or unassigned) indicating haploid read depth along the human reference GRCh38. We aligned long reads to the human reference with pbmm2 (as above, preset “CCS” for PacBio HiFi) and generated bedGraph coverage tracks with BEDTools v2.29.0 (*10*). Coverage tracks were subsequently converted into the binary bigWig format using bedGraphToBigWig v377 (*11*) to reduce data transfer volumes:

```
bedtools genomecov -bg -ibam {input_alignments} |
LC_COLLATE=C sort --buffer-size={sort_buffer}M
--parallel={threads} -k1,1 -k2,2n > {output_bedgraph}

bedGraphToBigWig {input_bedgraph} {hg38_chroms} {output_bigwig}
```

#### *8.1. K-mer-based analysis of phased assemblies*

**(*Contributors: Peter Ebert*)**

The basic data structure for the k-mer-based analysis of the phased assemblies is a colored de Bruijn graph constructed with Bifrost (*12*). To benefit from CPU architecture optimizations, we compiled the Bifrost executable from source (git commit hash ab43065). For each phased assembly, a colored de Bruijn graph was constructed as follows:

```
Bifrost build --input-seq-file {Illumina_reads} --input-ref-file
{phased_assemblies, GRCh38 reference} --output-file {graph} --threads
{threads} --colors --kmer-length 31
```

The Illumina short reads were quality trimmed with Trim Galore v0.6.5 ([www.bioinformatics.babraham.ac.uk/projects/trim_galore](www.bioinformatics.babraham.ac.uk/projects/trim_galore)) and error corrected with Lighter v1.1.2 (*13*) to ensure that predominant reads consisting of high-quality genomic k-mers were used in the graph construction process:

```
trim_galore --quality 20 --length 51 --trim-n --output_dir {outdir}
--cores 4 --paired {Illumina.pair-mate1} {Illumina.pair-mate2}

lighter -r {Illumina.pair-mate1.trim} -r {Illumina.pair-mate2.trim}
-k 31 3100000000 {alpha} -od {outdir} -t {threads} -zlib 6
```

The parameters for Lighter were set to match the Bifrost graph construction process (k = 31, genome size of GRCh38 3.1 Gbp), and “`alpha`” was computed following developer recommendations as 7 divided by the coverage of the Illumina short reads relative to the GRCh38 reference. Presence of regulatory sequences in the phased assemblies was assessed by querying the de Bruijn graph with k-mers generated from regions annotated in the Ensembl Regulatory Build v98 (*14*). Regions shorter than the k-mer size of 31 bp were discarded (167 regions of type “transcription factor binding site” [TFBS] with a combined length of 2600 bp, approximately 0.0002% of all bases in TFBS regions). A query regulatory sequence is counted as present in the haploid assembly if 99% of its constituent k-mers are matched with at most one mismatch or indel error (Bifrost parameter "`--inexact`"):

```
Bifrost query --input-graph-file {graph} --input-color-file
{graph_colors} --input-query-file {input.queries} --output-file
{output} --threads {threads} --inexact --ratio-kmers 0.99
```

#### *8.2. Reference-based analysis of phased assemblies*

**(*Contributors:* Peter Ebert)**

We aligned all phased assembly contigs to the GRCh38 reference genome as previously described (DOI:10.1038/s41587-020-0719-5) and examined aligned contig coverage for several annotation data tracks. If not indicated otherwise, alignments were filtered to only retain aligned contigs with the highest alignment quality (MAPQ = 60), and potentially overlapping contig alignments were merged using BEDTools' "`merge`" command (*10*). GRCh38 reference annotation tracks for cytogenetic bands, SDs, assembly gaps, repeat elements, and functional elements were downloaded from the UCSC Table Browser (*15*–*17*). GRCh38 locations of centromeres and of unresolved sequence issues (type "Gap" or "Unknown") were downloaded from the Genome Reference Consortium (*18*) on 2020-07-23. All issue annotations without sequence coordinates or with an indicated fix version (minor GRCh38.p14 or major GRCh39 release) were discarded. Raw and processed versions of the issue annotation are available in the assembly pipeline repository under "annotation/grch38" (see Code Availability). Illumina short-read genome coverage tracks indicating "Illumina-accessible" regions generated as part of the 1000 Genomes Project (*1*, *19*) were downloaded for both the "pilot" (lenient) and strict annotation (ftp.1000genomes.ebi.ac.uk/vol1/ftp/data_collections/1000_genomes_project/working/20160622_genome_mask_GRCh38). If applicable, the GRCh38 assembly was modified to define accessible regions of the genome by excluding Giemsa-stained positive or variable regions, roughly corresponding to heterochromatin, and by subtracting N gaps (see Table S9).

*Phased assembly gaps relative to GRCh38.* For the analysis of common gaps in the haploid assemblies, we identified candidate regions at several levels of stringency: we selected all regions where none of the assemblies had any contig alignment with minimum mapping quality of 0, 10 and 20, and excluded regions smaller than 10 kbp. The resulting list of common assembly break candidate regions was then subjected to a LOLA v1.12 enrichment analysis (*20*) to reveal underlying sequence features that may cause these assembly breaks. The necessary control set for the LOLA enrichment analysis was derived in a way similar to the candidate region set, but by selecting all regions where at least one assembly did not have a contig alignment with the required minimum mapping quality. The LOLA analysis was performed with standard parameters using the above listed UCSC annotation tracks.

#### *8.3. Assembly Scaffolding*

(***Contributors: Peter Ebert, Feyza Yilmaz, David Porubsky***)

Bionano cmaps were aligned to phased assembly haplotypes to identify haplotype 1 and haplotype 2 cmaps using Bionano RefAligner.

```
python2.7 Solve3.5.1_01142020/Pipeline/1.0/runCharacterize.py -t
Solve3.5.1_01142020/RefAligner/1.0/RefAligner -q {querymap} -r
{referencemap} -o {outputfolder} -p Solve3.5.1_01142020/Pipeline/1.0/
-a
Solve3.5.1_01142020/RefAligner/1.0/optArguments_haplotype_DLE1_saphyr
_human.xml -n 64 1>alignmentstatistics.out
```

Bionano genome maps that aligned to phased assembly haplotype 1 were assigned as haplotype 1, and genome maps that aligned to phased assembly haplotype 2 were assigned as haplotype 2. To obtain scaffolds, phased assemblies were mapped to *de novo* assembly of Bionano cmaps using the Bionano Solve3.5.1_01142020 hybrid scaffolding pipeline.

```
perl Solve3.5.1_01142020/HybridScaffold/12162019/hybridScaffold.pl -n
{inputNGSFASTA} -b {inputBionanocmap} -c
Solve3.5.1_01142020/HybridScaffold/12162019/hybridScaffold_DLE1_confi
g.xml -r Solve3.5.1_01142020/RefAligner/1.0/RefAligner -o
{outputFolder} -f -B 2 -N 2 -x -y -m {inputBionanoMoleculesbnx} -p
{inputDeNovoAssemblyPipelineDirectory} -q
Solve3.5.1_01142020/RefAligner/1.0/optArguments_nonhaplotype_DLE1_sap
hyr_human.xml -e {inputDeNovoAssemblyNoiseParameter}
```

We combined the information from the scaffolded Bionano hybrid assemblies with the contig-level phased assemblies as follows: we established the correspondence between Bionano scaffold and phased assembly cluster, i.e., human chromosome, based on the amount of mapping quality-weighted contig sequence aligned to the GRCh38 reference chromosomes. For cases where either less than 50% of the scaffolded sequence aligned to a single chromosome, or more than 50% pertained to unplaced sequence (“chrUn”), the respective scaffold was marked as entirely “unplaced”. We used the scaffold-to-chromosome assignment to characterize misassemblies detected and corrected in the Bionano hybrid scaffolding process (see Section 8.4). Next, we re-aligned all scaffolded phased assembly contigs to the matched chromosome (chromosomes X and Y for male samples), and all “unplaced” scaffolded sequences to the full GRCh38 reference (chromosomes 1-22, X; plus chromosome Y for male samples). All unsupported sequences, i.e., those parts of our phased assemblies that could not be scaffolded by Bionano, were split into 500 bp reads and aligned to GRCh38 using minimap2's "-sr" preset. The resulting read alignments were aggregated in genomic bins of 500 kbp and averaged over all samples before plotting.

For the realigned scaffolded contigs, we derived confidence levels as follows: scaffolds with aligned contigs scattered over at least two chromosomes were assigned low confidence. Scaffolds comprising a single aligned contig were assigned high confidence. For all other cases, a custom script was used to select all alignments within a region of the estimated scaffold length (plus 5% on both ends). The region center was chosen as alignment-weighted midpoint over all contig alignments, thus giving more weight to larger contigs that are part of the scaffold. In case no alignment was selected, and the scaffold comprised at least three contigs, the contig selection was adjusted by discarding the contig with maximal average distance to all other contigs, and subsequently recalculating the region midpoint. This strategy was limited to a single attempt to avoid placing scaffolds based on marginal evidence collected from few contig alignments. If no alignment could be selected to place the scaffold, e.g., because contig alignments were scattered along the entire chromosome, all contigs were assigned low confidence. If all contigs belonging to a scaffold were selected, and their alignment order was identical to the scaffolded order, the contigs were labeled as high confidence, and medium confidence otherwise.

#### *8.4. Analysis of phased assemblies*

**(*Contributors:* Peter Ebert)**

The design of our study enabled us to directly compare the results of our phased genome assembly pipeline for two types of PacBio long reads: less accurate, but on average, longer CLR reads and highly accurate CCS reads (HiFi; Figure 1b). We report statistics from different stages of our phased genome assembly process (Figure 1a) that suggest varying dependency between pipeline performance and type of the long reads being inputed. The initial step of creating a non-haplotype-resolved ("squashed") assembly using all input reads resulted in highly contiguous assemblies with a mean contig-level N50 of 30.5 Mbp (Figure 1d). However, only CLR-based assemblies exceeded N50 values of 35 Mbp in 20% of all cases, presumably due to the longer insert sizes of the sequencing libraries (Fig. S3, Table S5). These differences in assembly contiguity are less relevant for the subsequent step of clustering contigs into roughly chromosome-scale clusters using the Strand-seq data (see Methods in (*8*)). Consequently, on average, 98.4% of the sequence in one cluster aligns to a single GRCh38 reference chromosome (1-22, X). The pipeline step of identifying heterozygous (HET) single-nucleotide variants (SNVs) to obtain local phasing information indicates differences between CLR- and HiFi-based assemblies with an average of 2.32 and 2.66 million HET SNVs, respectively (Fig. S10). These numbers are, however, dominated by the high-diversity African population with a technology-agnostic average number of approximately 2.92 million HET SNVs per assembly, compared to 2.2 million HET SNVs for all other populations combined. These discrepancies seem to have little bearing on variant phasing or haplotagging efficiencies. In general, our assembly pipeline phases more than 99.9% of all HET SNVs irrespective of long-read technology or sample population of origin (Fig. S10). This information is then used to haplotag an average of 84.3% of all input long reads, with African samples reaching mean haplotagging rates of 89.8% versus 81.7% for non-African samples (Figure 1c). The assemblies built from the haplotagged read sets reach mean contig-level N50 values of 28.4 Mbp for CLR and 21 Mbp for HiFi reads (Figure 1d). Notably, 38.3% of all CLR haplotype assemblies exceeded a contig-level N50 of 30 Mbp—with a maximum of 39.4 Mbp—compared to only 3.6% for the HiFi-based haplotype assemblies. We computed base quality (QV) estimates for all polished haplotype assemblies in two orthogonal ways: first, we used Illumina short reads to identify homozygous variants as potential sequence errors (see Methods in DOI:10.1038/s41587-020-0719-5), which resulted in an average QV estimate of 49.6 (HiFi 53.8, CLR 47.7; Figure 1e). Second, we used 31-mer counts either unique to the assembly or supported by Illumina short reads or by the GRCh38 reference sequence to calculate an average assembly QV value of 40.4 (HiFi 43.1, CLR 38.8; Figure 1e). The assembly 31-mer counts supported by Illumina short reads can also serve as an estimate for the amount of sequence that is not present in the current GRCh38 reference. On average, we count 50 million homozygous and 38 million haplotype-specific 31-mers with Illumina support per sample (Fig. S11). The differences in haplotype-specific k-mer counts between CLR and HiFi assemblies are small (37.2 to 39.6 million) when contrasted with the comparison of African to non-African samples (46 to 34.2 million).

We further evaluated the completeness of our phased assemblies relative to various annotations for the GRCh38 genome reference (Methods). We plotted the contig coverage along the cytogenetic bands as the mean fraction of covered bases in each chromosomal segment (Fig. S6). This high-level view indicated that the majority of gaps in our assemblies coincides with (peri-) centromeric regions or occurs near the end of chromosomes (see also assembly break analysis in next paragraph). This observation is confirmed by an average 2.3% haploid assembly contig coverage in centromeres, with the exception of chromosome 20, where up to 38.7% of the centromere is covered by aligned contigs for HiFi assemblies (Fig. S7). Overall, we computed a median contig coverage of our haploid assemblies of >95% in accessible regions of the GRCh38 chromosomes 1-22 and X (Methods; Table S9). Next, we performed the same coverage analysis with a set of 98 GRCh38 regions annotated as "unresolved issues" by the Genome Reference Consortium (Methods; issue type "gap" partially coincides with centromeres). Our phased assemblies have, on average, assembled sequences for 67% of the determined bases in those regions, with no substantial difference between CLR and HiFi assemblies (66% to 68.3% coverage, Fig. S12). Other genomic regions that are prone to coverage dropouts in assemblies are often enriched in segmental duplications (SDs). For the UCSC SD annotation track (Methods), we observed a clear difference in contig alignment coverage between CLR and HiFi assemblies of 65% to 73.2%. We also examined the presence of known regulatory elements annotated in the Ensembl Regulatory Build v98 (*14*) based on their constituent k-mers (Methods). This analysis did not reveal any sizable differences between CLR or HiFi assemblies: approximately 90% of the sequences annotated as regulatory regions is detectable in both haplotype assemblies, and 3.2% is specific to only one haplotype (Fig. S13).

To systematically characterize regions that we identified as common assembly breaks (Methods), we performed enrichment analyses at various thresholds of contig alignment quality to account for potential alignment artifacts. At a minimum size of 10 kbp, we identified between 230 and 236 regions across all haploid assemblies as candidates for common assembly breaks. As expected, the results of our enrichment analysis include regions of unresolved sequence in GRCh38 ("N gaps"), which cannot be aligned and thus provide no mechanistically relevant insight. Besides, enrichment in genomic annotations such as RNA repeats (OR > 7), segmental duplications (OR > 3) and microsatellite repeats (OR > 2) suggests that genome assembly in these sequence contexts is still a challenge (Table S10). While it is expected that difficult-to-assemble regions such as centromeres lead to genuine gaps in the haploid assemblies, GRCh38-relative haploid contig coverage can only approximate the actual completeness of the *de novo* assemblies due to the unresolved N gaps in GRCh38.

As an orthogonal method for assessing assembly completeness that does not rely on aligning contigs to a reference, we scaffolded our contig-level phased assemblies for 32 out of the 35 individuals for which Bionano optical maps were available (Methods). This enabled us to detect and characterize various types of misassemblies (Fig. S8), the most frequent one being a contig break due to missing support from Bionano. Despite these assembly breaks (median of 65 misassemblies per haploid assembly), the median concordance between our phased assemblies and the Bionano maps is >97% (CLR median 98.2%, HiFi median 92.0%; Table S11). We aligned the unscaffolded sequence back to GRCh38 and observed a tendency for elevated read coverage in 500 kbp genomic bins around centromeres and in acrocentric regions, suggesting that at least some of the unscaffolded sequence may originate from those regions (Fig. S9).

Next, we realigned the scaffolded contigs to GRCh38 and classified contig alignments as high, medium or low confidence (Methods). Across all assemblies, a median of 69.01% of the aligned sequence is part of a high-confidence scaffold, with CLR samples showing an overall lower percentage (median 65.69%) compared to HiFi (median 79.03%). The majority of the remaining sequence pertains to contig alignments within medium-confidence scaffolds (median CLR 31.74%; HiFi 20.37%). For the remaining scaffolds, the majority of the contig alignments are scattered across one or more chromosomes, making it impossible to reasonably infer an approximate genomic location for the respective scaffold. Consequently, we consider the remaining contig alignments as low confidence (median CLR 0.62%; HiFi 0.3%), reflecting the view that the scaffolded assembled sequence cannot be adequately represented via GRCh38-relative alignments.

Finally, we combined the alignment information from contig-level and scaffolded assemblies to estimate the amount of sequence that can be interrogated with our phased assemblies. As a standard of comparison, we selected Illumina coverage tracks generated by the 1000GP (Methods) that define between ~2.29 Gbp (strict) and ~2.75 Gbp (lenient) of the human genome as "accessible" to short-read-based analyses (excluding N gaps in the GRCh38 reference, restricted to chromosomes 1-22, X and Y). We defined an analogous set of regions in two ways:

first, we selected all regions where at least one phased assembly had a MAPQ 60 contig alignment; three samples (HG03486, HG02818, HG03125) were excluded from this procedure because no corresponding Bionano data were available for hybrid scaffolding of these samples. Second, we selected all regions with contig alignments that were part of at least one high- or medium-confidence scaffold, i.e., the second set of regions does not enforce a threshold on the contig alignment MAPQ, thus being more permissive in that regard. Notably, we do not consider our scaffold-derived definition of "assembly accessible" regions as necessarily more lenient than the MAPQ-based regions because we included orthogonal information from Bionano supporting contig alignments that may be below a stringent MAPQ 60 threshold. Both resulting region sets cover approximately 2.87 Gbp of sequence, with as few as 250 and 243 regions, respectively (Fig. S14). The median region size is ~397 kbp (N50 ~58 Mbp) and ~480 kbp (N50 ~61 Mbp), compared to 79 bp (N50 ~2 kbp) and 240 bp (N50 ~20 kbp) for the strict and lenient Illumina region sets. Our definitions include ~582 Mbp more sequence relative to the strict definition of Illumina-accessible regions, and ~123 Mbp relative to the lenient definition. However, we also find that approximately 848 kbp and 121 kbp of the sequence in the Illumina strict set is not covered by our MAPQ60 or scaffold-derived coverage tracks, respectively. The lower amount of sequence that is missing from the scaffold-derived region set suggests that thresholding on the alignment quality excludes many regions that may be difficult to align to. We thus first removed all Illumina-exclusive strict regions overlapping centromeres (~2 kbp), which we already know not to be accessible by our assemblies (see above), and then annotated all remaining Illumina strict regions with their distance to the closest SD. As expected, we observe a median distance of 40 kbp (mean: 53.5 kbp) between the Illumina region and the closest SD for regions missing from our MAPQ60 coverage track, and a median distance of 0 kbp (mean: 5.9 kbp), i.e., overlapping, for Illumina regions missing from our scaffold-derived coverage track. We observed a similar behavior when checking how many of our SV calls based on the phased contig-level assemblies are located outside of our region sets. For our MAPQ60 assembly coverage track, we found 1,222 SV calls to be uncovered. This number was reduced to 113 for the coverage track based on the scaffolded assemblies, which in turn does not include regions covered by contig alignments as part of low-confidence scaffolds, irrespective of the MAPQ value of the individual contig alignments.

### 9. Reference and global merging strategy

**(*Contributors: Peter Audano*)**

#### *9.1. Genome references*

*GRCh38 No-ALT reference.* Long reads and contigs were aligned to the GRCh38 primary assembly only, which includes chromosome scaffolds, unplaced contigs, and unlocalized contigs. No ALTs, patches, or decoys were included, which were constructed for short-read mapping and would confound long-read and assembly analysis. This reference was used by multiple variant discovery pipelines including PAV and PBSV (see below).

The No-ALT reference is available on the IGSR FTP site:
ftp://ftp.1000genomes.ebi.ac.uk/vol1/ftp/data_collections/HGSVC2/technical/reference/20200513_hg38_NoALT/

##### 9.2. Variant merging strategies

Variant merging was done for several analyses in this study, including merging haplotypes into a diploid callset, merging calls into a nonredundant set, and comparing SVs with other callsets. Merging variants by 50% reciprocal overlap (RO) has been used for numerous SV papers (*3*, *21*, *22*), but it often under-merges small SVs and indels.

To address this problem, we adopt a three-step approach for SVs and indels. First, exact match variants are intersected (same size and location for insertions and deletions). Second, the RO is applied to variant calls for variants with intersecting reference coordinates. For insertions, the end position is the sum of the start position and variant length. Since an RO-only approach often misses smaller variants, we finally merge variants within 200 bp and 50% overlap by size (i.e., maximum RO if variants were shifted) (Fig. S15). For the 200 bp distance, we take the minimum of the start and end position offsets. SNVs are only intersected by exact match (same position and alternate base). In all cases, only the same variant classes are considered for matches (INS with INS, DEL with DEL, etc.). During this merging process, a variant in a new sample could support a call if it intersects the lead variant of a merged set (i.e., it is not matched against other supporting variant calls). The code implementing this approach is now released by HGSVC as SV-Pop (https://github.com/EichlerLab/svpop).

#### 10. Phased assembly variant discovery

(***Contributors: Peter Audano***)

Variants including SVs were called by directly comparing the haplotype-resolved assemblies to the human genome reference, GRCh38. This process is implemented in the Phased Assembly Variant (PAV) discovery tool described in this section (https://github.com/EichlerLab/pav).

##### 10.1. Contig alignment and trimming.

*Contig alignment.* For each haplotype, contigs were aligned to the GRCh38 No-ALT reference with minimap2 2.17 with parameters “-x asm20 -m 10000 -z 10000,50 -r 50000 --end-bonus=100 --secondary=no -a -t 20 --eqx -Y -O 5,56 -E 4,1 -B 5” (*23*). The callset was generated using minimap2 alignments. For validations, we also ran the pipeline with LRA (https://github.com/ChaissonLab/LRA) alignments using parameters “-CONTIG -p s -t” (PAV-LRA).

*Alignment trimming per haplotype.* Minimap often makes redundant alignments where the same part of a contig is aligned to more than one location. Alternatively, a location of the reference may be covered by more than one alignment record, and these overlaps often occur around SVs that truncate alignments (i.e., one contig in multiple alignment records). For example, for a large deletion flanked by repeats, the single contig copy of the repeat is often mapped to both reference copies. This obscures the size of the SV and introduces artifacts in the flanking repeats that appear to be SNVs, indels, and small SVs. A similar anomaly was observed for tandem duplications where multiple contig copies were aligned to the same reference copy. To address these multiple-mapping issues, we trimmed alignment records to resolve multiply-mapped contig bases followed by a second round to trim multiply-mapped reference bases. For a pair of alignment records, trimming is performed with a dynamic programming algorithm that attempts to maximize the number of variant events removed, conditioned on zero overlap of trimmed alignment. One variant event is defined as a single SNV, insertion, or deletion regardless of size. In short, it takes one contig and finds the cut-site if only that contig were trimmed. It then traverses the CIGAR operations of both alignment records using the zero-overlap and maximum-event conditions to guide its progress without considering more cut-site combinations than necessary (Fig. S16). The trimmed bases are soft-clipped. Lastly, whole alignment records less than 1 kbp before or after trimming, or if it is completely contained within another alignment record. One haplotype can be processed within approximately 1-2 core minutes. After alignment trimming, each assembly base maps to, at most, one reference base and vice versa. The alignment-truncating variant caller in PAV (see below) then uses these refined breakpoints to make an SV call.

The variants in Fig. S16 were aligned by LRA without fragmenting assemblies, and the SVs were called directly from the CIGAR string. Both minimap2+alignment trimming and LRA place breakpoints at nearly identical locations and call an SV of a similar size.

#### *10.2. Interalignment and alignment truncating variants*

*Interalignment variant calling per haplotype.* Most variant calls are contained within alignment records and can be obtained by examining the alignment CIGAR string. For this, PAV requires alignments with “=” and “X” CIGAR operations (an error is produced if there are any “M” CIGAR operations). These variant calls rely on the aligner for correct placement.

*Alignment-truncating variants.* SVs often cause contig alignments to break, leaving more than one alignment record for the same contig. Because alignments are trimmed, the insertion and deletion breakpoints are mapped at the ends of the alignment records and can be identified if they leave an excess of reference bases (deletion) or contig bases (insertions). Both alignment records must be in the same orientation or SV discovery is not attempted. The variants are detected by finding alignment records matching the same chromosome and contig. The reference and contig size gap between the two alignment records is compared to the minimum alignment length of two records (lesser of the number aligned bases in each alignment). If the minimum alignment length is greater than the contig gap size or greater than three times the reference gap size, the pair are ignored and no SV call is attempted.

#### *10.3. Inversion detection*

*Flagging inversion signatures.* Inversions in assemblies either fragment alignments into alternating forward and reverse oriented records, or they cause aberrant variant patterns if contigs are aligned through them without inverting. We define these as inter-alignment and intra-alignment inversions, respectively. Intra-alignment inversions often result in matched insertion and deletion events of a similar size, and they are often accompanied by clusters of false SNVs and indels (Fig. S17-a). We take advantage of this by using matched SVs and indels to flag regions that may contain an inversion. Inter-alignment inversions are flagged by alignment-truncating events (Fig. S18-a).

*Inversion detection with k-mer density.* Each intra- and inter-alignment flagged site is a reference region (contig, start position, and end position). It is initially expanded by 4 kbp because flagged sites are often much smaller than the inversion. K-mers of size 31 (31-mers) are extracted from the reference region and counted. If a reference 31-mer appears more than 100 times in that region or if the region contains N bases, inversion discovery terminates to avoid excessive CPU time spent on unresolvable loci.

Using the alignments, the reference region is lifted to a contig region. If the lift fails or the reference and contig region sizes are not within 60%, inversion detection is terminated. The contig region is extracted and translated into a list of k-mers in the order they appear in the contig region. Reference k-mers are reverse complemented if contig alignment records suggest that the contig is in the opposite orientation of the reference. Using the reference k-mers, each contig k-mer is assigned to one of three states: FWD (reference k-mer set has the k-mer), REV (reference k-mer set has the reverse-complement of the k-mer), or FWDREV (reference k-mer set has both FWD and REV k-mers). Contig k-mers with no representation in the reference k-mer set are dropped. If any state (FWD, REV, FWDREV) has fewer than 20 k-mers, those k-mers are also dropped because they cause large false spikes in density calculations that confound analysis. If this results in fewer than 2,000 k-mers, inversion detection is terminated.

A density function is constructed to guide inversion detection (Fig. S17-b and Fig. S18-b). First, the remaining informative k-mers in FWD, REV, and FWDREV states are reindexed consecutively from 1 (1 ... nth k-mer), which reduces biases in the density function if an inversion contains variants (e.g., Alu insertion), but the original index is retained so coordinates can be accurately translated back to the contig region. A Gaussian-kernel density function is then computed over these indices. Density bandwidth is computed using Scott's rule with one-dimensional data using all informative k-mers, and the density function is constructed with Numpy 1.16.2 (https://docs.scipy.org/doc/scipy/reference/generated/scipy.stats.gaussian_kde.html). One density function for each state is then computed. When the density function is applied to a k-mer index, it is subsequently scaled by the number of k-mers with the same state (FWD, REV,

FWDREV) so that density-smoothed values for each state approach 1 regardless of the number of k-mers in each state, which is needed for maximum state computations.

Density calculations are initially computed on one of every 20 informative k-mers and the blocks are subsequently filled in by interpolation (to save CPU cycles) or full density computation. If there are no state changes within the block and the density value between each side of the block changes by less than 0.005, the block is filled in by interpolating with the density values on each side, otherwise, a full density computation is performed for each index in the block.

Finally, each k-mer is assigned a new state using the maximum density, which yields smoothed runs of FWD, REV, and FWDREV states. An inversion is detected if the contig region begins and ends with reference oriented k-mers and contains inverted k-mers. The outer breakpoints of the inversion are placed at the first and last non-FWD states in the state list, and these outer breakpoints are the reported breakpoints for the inversion event. Often, k-mers are flanked by inverted duplications, so a typical inversion state progression is “FWD, FWDREV, REV, FWDREV, FWD”. In this case, inner breakpoints are placed at the first and last REV state(s) and reported as an annotation with the inversion call. Finally, the breakpoints identified in contig coordinates are lifted to the reference (Fig. S17-c and Fig. S18-c) and a variant call is constructed with inner- and outer-breakpoint annotations in contig and reference coordinates.

If inverted k-mers are identified in the region but density calculations did not yield smoothed states indicating an inversion (FWD states on flanks with REV states between), then the region is expanded by 50% and the density is recomputed. If one flank has FWD k-mer states, then the expansion is biased to add more bases on the non-FWD-state end. If only FWD k-mers are found after three expansions (including the initial expansion), inversion detection terminates. There is currently no upper limit, so expansion could continue until alignment records for the contig are exhausted.

##### *10.4. Callset finishing*

*In-PAV filtering.* Variant calls intersecting inversions are removed. Variants occurring within inter-alignment deletions are also removed. We observed that incomplete phasing generated false contigs that fit into large heterozygous deletions, which was likely created by reads from the other haplotype. Ignoring variants inside deletions on the same haplotype removes false calls from these phantom contigs and from misalignments.

*Merging haplotypes.* An independent callset is first generated for each haplotype and they are merged using the three-step strategy (see Section 9.2). As before, SNVs are intersected only if they match exactly (position and alternate base). PAV merges SVs and indels as one large set of variants and then splits them into SV/indel variant classes post-merge. For all homozygous variants, PAV chooses the variant representation from haplotype 1 as a default by the merging process.

##### *10.5. Post-PAV callset filters*

*SDA.* Many high-identity repeats are still not resolvable by assemblers and result in assembly collapses, which carry a mix of paralog-specific variants (PSVs) from multiple sites and may or may not align to the best reference copy. We identified collapsed regions with Segmental Duplication Assembler (SDA) (*24*) and filtered PAV calls if they intersect a collapse on either haplotype. To avoid over-filtering inversion and deletion SVs, we require at least 50% of the SV to intersect a collapse before removing. Indels and SNVs are removed if they intersect collapses by 1 bp or more (Table S14).

*Misclustered contigs.* Using cluster IDs generated by the assembler, which were embedded in the contig names, PAV assigns the best reference chromosome for each cluster if 85% or more of the mapped bases within contigs 1 Mbp or greater are assigned to the same reference chromosome. For chromosomes where a max was defined, variants called on contigs belonging to a different cluster are filtered out. This prevents misaligned contigs (or parts of contigs) from inflating the callset.

*Pericentromeric filter.* A centromere and pericentromeric region filter (*3*, *21*) is applied to address regions that are difficult to replicate among callsets.

#### 11. Variant discovery with aligned reads

To increase sensitivity, we also applied methods that call variants based on mapping of the underlying raw long-read data against the human reference genome (GRCh38). These methods have been optimized for particular SV classes (e.g., PBSV for SVs and DeepVariant for indels and SNVs).

##### *11.1. PBSV*

(***Contributors: Aaron Wenger and Peter Audano***)

Reads were aligned to GRCh38-NoALT (Supplementary section 9.1) using pbmm2 version 1.2.1 (https://github.com/PacificBiosciences/pbmm2) with "--sort --preset CCS -L 0.1 -c 0" for CCS and “--sort --median-filter” for CLR. Variants were called using pbsv version 2.3.0 (https://github.com/PacificBiosciences/pbsv). The pbsv workflow was executed separately per sample (not jointly) and per chromosome. First, SV signatures were discovered with "pbsv discover --tandem-repeats <SR.bed> -r <CHROM>" where SR.bed is the UCSC simpleRepeats track (http://hgdownload.soe.ucsc.edu/goldenPath/hg38/database/simpleRepeats.txt.gz) with regions within 200 bp merged ("bedtools merge -d 200"). Next, variants were called with "pbsv call" with "--ccs -O 2 -P 20 -m 10" for CCS and "-m 10" for CLR. For each sample, the per-chromosome calls were concatenated, sorted, and compressed with bcftools 1.9 (http://samtools.github.io/bcftools/bcftools.html).

#### *11.2. DeepVariant*

(***Contributors: William Harvey***)

CCS reads were aligned to the human reference genome GRCh38 with pbmm2 (https://github.com/PacificBiosciences/pbmm2) with "--preset CCS --min-length 5000". DeepVariant v0.9.0 (*25*) uses a deep neural network with a pre-trained model (--model_type=PACBIO) specifically for PacBio CCS reads. The pipeline consists of three separate steps: make_examples, call_variants, and postprocess_variants. The first step utilizes the existing alignments to make a six-channel (read base, quality score, mapping quality score, read strand, read allele support, read match reference) TensorFlow object, which is then processed by the neural network using the pre-trained model to make variant calls. The variant calls are processed and combined into a VCF.

#### *11.3. DeBreak*

(***Contributors: Zechen Chong, Yu Chen***)

Reads were aligned to human reference genome GRCh38 with minimap2 (*26*) version 2.15-r905 with the option "--secondary=no" for both CLR and CCS samples. DeBreak version v1.0.2 (https://github.com/ChongLab/DeBreak) was executed to call variants from read-to-reference alignments for each sample independently with option "--poa --ref hg38.fa". For each sample, DeBreak scanned all read alignments for both intra- and inter-alignment SV signals. Smaller indels can be detected within a single alignment. For larger indels, inversions, duplications, and translocations (potential mobile element insertions or MEIs), DeBreak utilized split-read information to infer SV breakpoint positions and lengths. All raw SV calls were sorted and clustered with density-based clustering to generate SV candidates. Auto-adaptive filters were applied to remove noises and artifacts. Only variants that passed all filters were retained, while variants with "HighCov" or "LowMapQ" filters were discarded to ensure accuracy of SV callsets. Partial order alignment (*27*) was carried out using variant-supporting reads for each variant to refine SV breakpoints. SVs located in ATL contigs were also removed from the callset (Table S15). The DeBreak SV callset was compared with the PAV callset for each sample with 50% RO (Fig. S19). For CLR data, the numbers of DeBreak deletions overlapping with PAV ranged from 6,882 to 8,738, while insertions ranged from 10,595 to 12,913. For CCS data, the numbers of DeBreak deletions overlapping with PAV ranged from 7,076 to 9,040, while insertions ranged from 10,907 to 13,181. DeBreak also identified unique calls. For CLR data, the numbers of unique DeBreak deletions ranged from 991 to 1,495, while unique insertions ranged from 2,385 to 3,560. For CCS data, the numbers of unique DeBreak deletions ranged from 1,307 to 2,057, while unique insertions ranged from 1,691 to 3,501.

### 12. Illumina SV calling

**(*Contributor: Mike Talkowski, Xuefang Zhao and Qihui Zhu*)**

#### *12.1. SV discovery from individual algorithms*

SV discovery from Illumina short-read whole-genome sequences (srWGS) on the 34 samples in this study were done in two batches: 15 samples (HG00096, HG00171, HG00513, HG00731, HG00732, HG00864, HG01596, HG03009, NA12878, NA18534, NA18939, NA19238, NA19239, NA20509, NA20847) were processed in the first batch that included 2,504 genomes, which we refer to as 'batch 1' in the following text. The remaining 19 samples (HG00512, HG00514, HG00733, HG01114, HG01505, HG02011, HG02492, HG02587, HG02818, HG03065, HG03125, HG03371, HG03486, HG03683, HG03732, NA12329, NA19240, NA19650, NA19983) were processed together with another 679 samples in 'batch 2'. SV discovery methods were mostly the same on the two batches with slight differences in technical details.

**Manta**
(***Contributor: Allison Regier and Xuefang Zhao***)

The Manta VCFs for batch 1 samples were produced in a docker container with the image halllab/manta_samtools@sha256:6c8dfccfd3124ebf902ac6f0303e6f09b02a15e2c09963354620740788c407d0 using Manta v1.4.0 (*28*) with default parameters.

Manta V1.5.0 (*28*) were applied to batch 2 samples in single-sample mode on Terra (previously FireCloud) with default parameters. SVs were only discovered from canonical chromosomes (chromosome 1 to 22, X and Y) and mitochondrial DNA.

**Wham**
(***Contributor: William Harvey and Xuefang Zhao***)

Wham v1.7(*29*) was applied to the 15 samples in batch 1 in single-sample mode with default parameters and a mapping quality filter of 15. All SVs were required to be called as a 'PASS' by whamg genotyping and have a confidence of variant mapping >0.2. Additionally, DELs and INVs smaller than 50 bp and larger than 1 Mbp were removed. For DUPs, the size threshold was 300 bp and 1 Mbp, respectively. Finally, calls inside of SDs, tandem repeats, and centromere/peri-centromere regions were removed.

Wham v1.7(*29*) was applied to batch 2 samples in single-sample mode on Terra with default parameters. A customized whitelist that excluded N-masked genomic regions to keep the computing cost under reasonable scale. This whitelist is publically available at: ("gs://gatk-sv-resources/resources/wham_whitelist.bed")

**MELT**
(***Contributor: Scott Devine***)

MELT v. 2.1.5 (*30*) was used to discover MEIs (Alu, L1, SVA, HERV-K) from all samples in both batches using the MELT_Split mode. Trace size (-t) was set to 150 bp, coverage (-c) was set to 30X, and human reference genome GRCh38 was used as the reference in this analysis.

**Lumpy**
(***Contributor: Allison Regier***)

Lumpy (v0.2.13) (*31*) was run on all batch 1 and 2 samples using the Smoove wrapper (v0.2.3) in a docker container with the image brentp/smoove@sha256:c839ed223462a1c1ae26e7acc27f28f0f67b4581d80a06823895f295ad2bdaf4. The command `smoove call` was run with the `--noextrafilters` and `--genotype` parameters set. The `--exclude` parameter was set to blacklist regions in gs://human-b38/GRCh38DH/annotations/exclude.cnvnator_100bp.GRCh38.20170403.bed.

**CNVnator**
(***Contributor: Allison Regier***)

CNVnator (v0.3.3) (*32*) was run on all batch 1 samples using the wrapper script cnvnator_wrapper.py contained in the docker image halllab/cnvnator@sha256:8bf4fa64a288c5647a9a6b1ea90d14e76f48a3e16c5bf98c63419bb7d81c8938. Depth histograms were generated with a window size of 100 bp.

#### *12.2. SV integration*

##### 12.2.1. FusorSV
(***Contributors: Qihui Zhu***)

Existing SV detection algorithms are typically optimized for the discovery of specific types of SVs, thus their performance varies by type, size and genomic location of SVs. Therefore, it is impossible to achieve comprehensive discovery over the entire SV spectrum with a single algorithm. To mitigate this issue, we applied FusorSV (*33*), a unique data-mining method, to integrate SV calls from six different algorithms including CNVnator (*32*), Delly (*34*), Manta (*28*), Lumpy (*31*), WHAM (*29*) and MELT (*30*). Tool versions were described above in the section 12.1.

FusorSV takes SV calls from different algorithms and integrates them in a manner that minimizes false positives and maximizes discovery. FusorSV includes two phases: the training and the discovery phase. In the training phase, we use the Illumina SV calls from NA19238, NA19239, HG00731, HG00732 and HG00513 reported by Chaisson et al. (*3*) as the ground truth to train a FusorSV model. The performance of each of the six algorithms was compared to the ground truth, in addition to calculating the pairwise performances across all algorithms.

Combinations of algorithms that work more comprehensively will be promoted. Then the score for every possible combination of algorithms will be calculated and the performance value will be determined to a FusorSV fusion model (30). In the discovery phase, the FusorSV fusion model was used for comprehensive SV detection in all 33 test samples. This fusion model integrated individual algorithm performances with pairwise similarity of algorithms across SV types, which allowed FusorSV to select subsets of algorithms for each SV type that are more comprehensive, with maximum discovery and minimal subset false positives.

FusorSV provides a unique data-mining method that intelligently takes SV calls from different algorithms and combines them in a manner that minimizes false positives and maximizes discovery. Using per-algorithm performance information and similarity between algorithms, the smallest set of SV callers can be selected using the concept of mutual exclusion, which makes our method both more accurate and comprehensive than other approaches merging SV calls based on consensus or other heuristics.

##### 12.2.2. GATK-SV

(**Contributors: Xuefang Zhao**)

GATK-SV is a multi-module SV discovery and refinement pipeline for srWGS data. This method was previously adopted by the Genome Aggregation Database (gnomAD) for SV discovery, and the technical details have been described by Collins et al. (*35*). In this study, GATK-SV was applied to all 3,202 samples from both batches for SV discovery, genotyping, complex resolution, and final refinement. Moreover, we applied a machine-learning method (*36*) to further refine the callset. Most parts of the method were the same as in Collins et al. (*37*), while processes that were unique to this study are described here. An overview of this method is represented in Fig. S20a.

###### 12.2.2.1 Sample batching

The samples were clustered into batches of ~200 samples for raw algorithm processing, then merged for complex resolution and filtering. The batching strategy depends on the gender, dosage bias score ($\partial$), and median read coverage of each sample. $\partial$ and median read coverage were calculated to determine genomes with an unusual distribution of reads across segments of the genome using the same method as in Collins et al. (*35*). Samples were first separated by gender to form groups of 1,599 male samples and 1,603 female samples. Each group was then divided into four subgroups by the rank of median coverage, and each subgroup was further split into four groups by the rank of $\partial$. The resulting 32 batches of approximately 100 samples of unique gender were matched by their rank of median coverage and $\partial$ to form 16 batches of 200 samples with equal numbers of male and female samples. A brief summary of the batching schema is represented in Fig. S20b.

12.2.2.2 SV discovery from raw algorithms

SVs discovered from Manta (*28*), Wham (*29*), MELT (*30*), cn.MOPS (*38*) and GATK-gCNV were integrated in this pipeline. Tool versions and run settings of Manta, Wham and MELT have been described above. cn.MOPS was executed in a custom implementation on Terra for all samples, in ~200-sample batches. For each batch, the read depth (RD) per 100 bp bin across each genome was calculated and merged to form an RD metric. We composed RD matrixes across all samples at 300 bp and 1 kbp resolution, excluding any samples with a median bin coverage of zero per contig, then ran cn.MOPS with R v3.3.3 (*39*), split raw calls per sample, segregated calls into deletions (copy number < 2) and duplications (copy number > 2), merged the 300 bp and 1 kbp resolution calls per sample per CNV type using BEDTools (*10*) merge, and subtracted any N-masked bases from all CNV calls using BEDTools subtract. GATK-gCNV was implemented on Terra and applied to all samples in each batch with default parameters. Full details of this method can be found here: https://github.com/broadinstitute/GATK-gCNV-publication.

##### *Integration, quality refinement and re-genotyping of SVs*

Module 01-05 of GATK-SV has been described in Collins et al. (*35*); please refer to supplementary pages 45-50 for technical details.

##### *SV refinement*

This refinement module corrects false positive variants caused by the ambiguous alignment of short reads. The refinements were applied in two layers: per-sample level and per-SV-site level. For SVs genotyped as non-reference in each sample, multiple features were collected, including averaged read depth (RD) of each SV region, averaged RD of the 1 kbp flanking region of each SV, count of aberrant pair ends (PE) within 150 bp of each SV, count of split reads (SR) within 100 bp of each breakpoints, variant type, allele fraction across all samples, count of children that carry this SV as *de novo*, genomic location of the SV (split into simple repeats, SDs, and the rest relatively unique genomic sequences), genotype prediction and quality from previous modules, and whether they were discovered by the original algorithms. These features were integrated into a lightGBM model using these settings: *objective = "binary", metric = "auc", num_class = 1L, learning_rate = 0.1, min_data_in_leaf = 1L, min_sum_hessian_in_leaf = 1.0*. SVs in three samples, i.e., HG00514, HG00733 and NA19240, which were overlapped by PacBio SVs reported by Chaisson et al.(*3*) and validated by VaPoR (*40*) on the high-coverage PacBio sequences were selected as truth set, while SVs that neither have overlap in PacBio nor supported by VaPoR were considered as false positives. A unique score, referred to as 'boost score' (BS) in this manuscript, would be assigned to each SV by the lightGBM model. To keep the false discovery rate (FDR) of SVs under 5%, we considered SVs with BS lower than -0.404 as low quality and removed from the callset.

For each SV site, a boost model ratio (BR) is defined as the proportion of samples that passed the lightGBM model among all samples that had non-reference genotypes and used to infer quality of the SV locus. To keep the *de novo* rate of SVs under 5% in the 602 trios, which represents a combination of false positives in children and false negatives in parents, we set BR at 0.509 and labelled SVs with lower BR as low-quality loci.

It is noticed that VaPoR has decreased power when evaluating large CNVs located within SDs, so we also applied the VCF refinement method described by Collins et al. (*35*) (pages 50-51 of the supplemental methods) and included the large CNVs (>5 kbp) that passed this method into the final GATK-SV callset.

**Comparison of SVs between short read Illumina sequences and long read PacBio sequences.**

SVs detected from srWGS were compared against SVs from lrWGS with matching samples on the matched sample. SVs were considered concordant if they met these following criterias:

1.For insertions: a pair of SV loci were considered concordant if their reported insertion points was were within 100 bp from each other, and the length of their inserted sequences was within 10 times of size of each other.;

2. For deletions duplications, CNVs, inversion and complex SVs that are >5 kpb in size, 50% reciprocal overlap and match of variant types were required to be considered concordant; while for variants that are between 50 bp and 5 kbp, 10% reciprocal overlap and match of variant types were required. Complex SVs were considered concordant if overlapped by inversions, and CNVs were considered concordant if they overlap either deletions or duplications.

### 13. Bionano Genomics discovery

(***Contributors: Alex Hastie, Joyce Lee and Feyza Yilmaz***)

#### *13.1. Bionano Genomics de novo assembly and structural variant calling*

*De novo* assemblies of the samples were obtained using the Bionano's *De Novo* Assembly Pipeline (Bionano Solve v3.5) with haplotype-aware arguments (optArguments_haplotype_DLE1_saphyr_human_downSampleLongestMole.xml in http://ftp.1000genomes.ebi.ac.uk/vol1/ftp/data_collections/HGSVC2/working/20200219_Bionano_optical_map_SVs/). With the Overlap-Layout-Consensus paradigm, pairwise comparison of DNA molecules, which are at least 250 kbp in length and contribute to a coverage of 250X, was generated to create a layout overlap graph and produce initial consensus genome maps. By realigning molecules to the genome maps (alignment confidence cutoff of p-value < $10^{-12}$) and by using only the best match molecules, we applied a refinement step to label positions on the genome maps and to remove chimeric joins. Next, during an extension step, the software aligned molecules to genome maps (p-value < $10^{-12}$) and extended the maps based on the molecules aligning past the map ends. Overlapping genome maps were then merged (p-value < $10^{-16}$). These extension and merge steps were repeated five times before a final refinement was applied to "finish" all genome maps.

To identify all alleles, clusters of molecules that are aligned to genome maps with unaligned ends >30 kbp in the extension step were re-assembled to identify potential alternate alleles. To identify alternate alleles with smaller size differences from the assembled allele, we also searched for clusters of molecules that aligned to genome maps with internal alignment gaps of size <50 kbp, in which case, the genome maps were converted into two haplotype maps. To improve accuracy, large repetitive regions, such as SDs, are identified in a *de novo* fashion during the last round of merging. Maps sharing more than 140 kbp in common but diverging on both sides of the non-unique region generate splits within alignment maps. Molecule clustering and SD filtering generate more accurate diploid assemblies. We called and annotated SVs using Bionano's *De Novo* Assembly Pipeline (Bionano Solve v3.5), in which the final genome maps were aligned (p-value $<10^{-12}$) to the reference genome (GRCh38).

#### *13.2. Bionano Genomics discovery of large, complex structural variants*

Using Bionano genome maps, we identified 19,821 insertions and deletions that are at least 5 kbp in size mapping to high-confidence regions where there were no more than two aligned genome maps at breakpoints (Table S17). We clustered calls based on 80% size concordance and 80% reciprocal overlay to generate a nonredundant set of 2,017 insertion and 1,978 deletion clusters (Table S18), of which 516 insertion and 331 deletion clusters were localized to regions where we identified at least five different SVs contributed by at least seven samples. About 75% of these multi-site complex polymorphic variants overlapped SDs, and 2-4% of them overlapped gaps in the current human reference genome suggesting that these have been particularly problematic during sequence and assembly. We plotted these large complex SV sites as well as the underlying set of coordinates in GRCh38 (Fig. S24). In addition to insertions and deletions, we also identified 162 nonredundant inversions and 192 duplications across all samples, of which 53% and 58% of them overlapped SDs, respectively.

We compared Bionano Genomics SV calls to those discovered by PAV, requiring 50% concordance with respect to size and breakpoints within 1-5 kbp (Fig. S25). With such criteria, 72% of the 19,821 Bionano calls overlapped SVs detected by the phased assembly of PacBio data (Table S17). The comparison revealed 3,453 Bionano unique insertions and 2,137 Bionano unique deletions, corresponding to 1,657 clusters (Table S19). A cluster might have PacBio overlapped calls in one sample but not in other samples, and we found that there were 697 insertion clusters and 478 deletion clusters (Table S20) that had never been found in any of the PacBio assembly based callsets. In addition, 383 insertion and 247 deletion clusters were localized at SDs and overlapped genes (Table S21) .

We highlight several examples of complex structural polymorphisms corresponding to gene-rich regions that are not yet fully resolved at the sequence level within our phased assemblies. For example, we identified a 150 kbp region enriched in SDs mapping to chromosome 1 (chr1: 108.3–108.45 Mbp), which overlaps *NBPF5P* and *NBPF6*. We also identified a 75 kbp inversion in 24 out of the 30 samples, and a 74 kbp deletion was found in the alternate allele in two of the samples. The same deletion was also found in three samples that do not have the inversion allele, and another 74 kbp deletion and 130 kbp inversion were found in two different samples (Fig. S26). In addition, we identified duplications of up to 10 copies on chromosome 5 (21.1–21.7 Mbp), which overlaps with *GUSBP1* (Fig. S27). Each one of these complex regions were characterized and manually evaluated by checking single-molecule support for each sample. Critical labels, which were located in unique regions, were identified to detect the number of molecules spanning regions of interest.

Similarly, we identified 18 haplotypes at 3q29 (chr3:195.6-196Mbp), which overlaps *SDHAP2* genic region (Fig. S28, Table S22), by manual curation. Single molecules that were anchored to unique regions confirmed each haplotype (Optical mapping molecule support and haplotype segment coordinates are available at https://figshare.com/s/9dcf5f72dbd28f55f3ae). Haplotype 1 (H1) is the most common with 12 haplotypes in five super populations. GRCh38 reference assembly haplotypes is the third most common haplotype observed in all populations except for AMR. A summary of the assembly contig coverage for 3p29 can be found in Table S23.

#### 14. Strand-seq Inversion detection and genotyping

(*Contributors: Ashley Sanders, David Porubsky, Wolfram Höps, Hufsah Ashraf, Maryam Ghareghani, Tobias Marschall, Jan Korbel*)

To detect inversions using Strand-seq data, directional composite files were generated for each sample as previously described (*3*, *41*). To automate composite file generation, a merging protocol was implemented using the breakpointR 'synchronizeReadDir' function (*42*), which locates Watson-Watson (WW) and Crick-Crick (CC) regions in each chromosome and for each cell before building these into the sample-specific composite files (Fig. S29). Segmental changes in composite file orientation, suggestive of an inverted allele, were identified using breakpointR. To detect both larger and smaller strand-state changes, we used breakpointR in two settings—applying either a window size length of 5 kbp or 20 reads per bin. In both cases, we scaled an initial bin size by multiples of 2, 3, 4, 5, 10 and 20. This resulted in a redundant dataset with putative inversions detected per sample.

To construct a nonredundant set of Strand-seq inversions, we merged and filtered all detected strand-state changes (putative inversions) in multiple stages as follows: First, we cropped regions that overlap with highly identical SDs (>=98% identity) or gaps defined in GRCh38 from each inversion breakpoint. Second, we iteratively merged ranges with >=50% RO. Such collapsed ranges were then subjected to regenotyping using 'genotypeRegions' of the primatR package (*43*). Each region in each sample was assigned a genotype: 'HET' - approximately equal mixture of plus and minus reads, 'HOM' - majority of minus reads, 'REF' - majority of plus (reference) reads and 'lowReads' - less than 20 reads in a region. Ranges that genotype only as a reference ('REF') orientation or has less than 20 ('lowReads') reads across all samples were filtered out. Next, we collapsed ranges that share the same genotype across all samples and are embedded with respect to one another. Lastly, from regions that genotyped only as a 'HET' or ‘lowReads’, we retained only those that do not overlap with regions where Strand-seq inversion call was already made. The same procedure was repeated for windows defined by the readcount (20 reads per bin), which allowed adding smaller inversions missed by the larger bin size to the final Strand-seq inversion callset.

*Results for inversion detection and genotyping.* The strategies for inversion discovery described in the previous sections operate on the basis of individual samples, leaving us with the need to unify these inversion calls across samples. Additionally, we saw an opportunity to integrate information about inversion loci across samples to further improve the individual callsets.

To this end, we developed a new method, ArbiGent, which determines inversion genotype likelihoods for chosen genomic loci of 5 kbp or larger in size based on Strand-seq data. We built ArbiGent utilizing a statistical framework previously devised for discovering subclonal structural variation in cancer (*7*), which we extended to allow estimating germline SV genotype likelihoods for DNA segments of choice using strand-specific reads. In particular, on the basis of a Bayesian probability framework that models strand- and haplotype-specific read counts using Negative Binomial distributions, ArbiGent computes inversion genotype likelihoods for inversions and copy number changes. SV genotype likelihoods derived from individual cells from the same sample are concatenated by summing up log-likelihoods across cells, to result in a combined genotype likelihood estimate per sample and genomic locus of interest. The code of ArbiGent is available open source at https://github.com/friendsofstrandseq/pipeline/tree/arbitrary-segments.

Utilizing inversion discovery techniques based on Strand-seq, Bionano and aligned long reads, we found 394 genomic loci with evidence for an inversion across the 64 haplotypes studied (see “Nonredundant callsets, filtering, and properties” below). We used ArbiGent to assign genotype likelihoods to these loci across all samples and filtered them for high-quality sites, which led to the removal of 60 sites for which Arbigent did not confirm any inverted haplotype across the samples considered (‘suspected false positive’). We additionally tested the inversion genotypes for Mendelian consistency, utilizing trio-based Strand-seq data previously generated in Chaisson et al. (*3*), which resulted in another eight rejected inversion sites. Sites that were consistently predicted as complex events (6) and those for which short-read mappability was low (71) were flagged, but kept in the final callset. Accordingly, we report 316 high-quality inversion sites with genotypes, with a median of 130 detected inversion events (54 in HOM state, 54 HET and 23 complex events) comprising 24.8 Mbp of inverted DNA per diploid human genome.

*Analysis of 16p12 regions.* Detailed analysis of 16p12 regions is based only on Strand-seq inversion calls detected based on the distribution of direct and inverted Strand-seq reads in composite files as stated above (Fig. S32a). Each inversion was phased by separating Strand-seq reads per haplotype as reported before (*43*). Inversions that fail to reliably phase in any given sample (due to a low read coverage) were considered as a reference orientation. Next, we enumerated the frequency of unique combinations of inverted alleles along the 16p12 region per superpopulation. We evaluated the genetic background of each inversion by exploring SNVs that lie -/+500 kbp from each inversion breakpoint. SNVs from within the inversion and those that overlap with SDs (>=98% identity) were removed from the analysis. For each inversion, we constructed a neighbor-joining tree based on the flanking SNVs. Inversions that occurred at seemingly different genetic backgrounds (flanking SNVs) were considered as likely recurrent or toggling (Fig. S33). Last, we phased genome assembly alignments to GRCh38 of 16p12 regions have been analyzed. We report only phased contig alignments with mapping quality 60. We summarized alignment gaps in respect to GRCh38 as a coverage of all gaps (Fig. S32b).

### 15. MEI discovery and integration

**(*Contributors: Scott Devine, Nelson Chuang, Weichen Zhou, Ryan Mills, Bernardo Rodriguez-Martin, Martin Santamarina*)**

Mobile element insertions (MEIs), including long interspersed element-1 (L1), *Alu*, and SVA (SINE-VNTR-Alu) retrotransposons, comprise approximately 46% of the human genome and represent ~25% of all SVs in human genomes (*44*, *45*). They have been shown to play an important role in human development, population diversity, and genomic disease (*46*–*50*). Various strategies have been developed to identify candidate polymorphic MEIs from srWGS data (*30*, *51*), though they struggle in regions where reads can map equally to multiple alternative genomic positions. Long-read sequencing technology provides a better resolution in such regions by directly sequencing long stretches of contiguous DNA that enable the discovery of potential overlooked MEIs (*3*, *21*, *52*, *53*). By leveraging the PacBio assemblies, especially from HiFi reads, we are able to discover the largest set of non-reference MEIs with *bona fide* characteristics among diverse samples, allowing the investigations of their impacts on shaping human genome and population diversity.

We initially identified 9,448 non-reference MEIs, including 7,741 *Alu*s, 1,168 L1Hs, and 539 SVAs in the final PAV PacBio assembly-based callset (after filtering and removal redundancy) using MEIGA annotation (Fig. S34a). We also generated an integrated callset to annotate the generic INS calls in the final phased assembly SV calls (PAV) to further maximize our discovery sensitivity. Based on the integrated callset, 9,950 non-reference MEIs, including 8,110 *Alu*s, 1,248 L1Hs, 589 SVAs, and three HERV-Ks, were annotated in the final PAV PacBio assembly-based callset (Fig. S34b).

We provided three methods for calling, integrating, and annotating MEIs: MELT, PALMER, and MEIGA (see 15.2. MEI integration). Overall, MELT reported 6,935 *Alu*s, 977 L1Hs, 502 SVAs, and 32 HERV-Ks; PALMER reported 7,607 *Alu*s, 1,226 L1Hs, and 478 SVAs among diverse samples; and MEIGA reported 7,866 *Alu*s, 1,194 L1Hs, and 556 SVAs by tracing back and annotating the original pre-filter PAV callset, and further reported 7,741 *Alu*s, 1,168 L1Hs, and 539 SVAs by tracing back and annotating the final PAV callset (Fig. S34a). Afterwards, we applied an integration strategy across these different platforms to obtain a better comprehensive landscape of MEIs among the diverse populations and then to annotate PAV callset (see 15.2. MEI integration) .

#### 15.1. Callsets across different platforms

##### 15.1.1. MELT

**(*Contributors: Scott Devine and Nelson Chuang*)**

Mobile element insertions (MEIs) were discovered in the same 33 genomes that were sequenced by PacBio sequencing in this study.  High-coverage (~30-40X) Illumina WGS were obtained from the 1000GP bucket at Amazon.  MEIs (*Alu*, L1, SVA, and HERV-K elements) were discovered using the mobile element locator tool (MELT, ver. 2.1.5 (*30*, *51*)) using MELT-SPLIT, which is a stepwise version of MELT that uses discordant read pairs and split reads to perform MEI discovery across all samples in a given population. This improves the modeling at each MEI site, and improves genotyping, compared to analyzing the samples as individuals.  We only included calls in the final VCF that were classified as “PASS” and “lc” (low complexity), and all calls with alternative filters were removed.  Build GRCh38 of the reference human genome was used to perform MEI discovery.  3' transduction analysis was conducted using the 3' transduction finder in MELT (*30*).  The MELT software package, along with documentation, can be downloaded from:  https://melt.igs.umaryland.edu/.

##### 15.1.2. PALMER

***(Contributors: Weichen Zhou and Ryan Mills)***

We developed an enhanced version of PALMER (Pre-mAsking Long reads for Mobile Element inseRtion) (*53*) to detect MEIs across the long-read-sequenced genomes (https://github.com/mills-lab/PALMER). Reference-aligned BAM files from long-read technology are used as input. Known reference repetitive sequences (L1s, *Alu*s or SVAs in reference) are used to pre-mask the portions of individual reads that align to these repeats. After the pre-masking process, PALMER searches subreads against a library of mobile element sequences within the remaining unmasked sequences and identifies reads with a putative insertion sequence (including 5′ inverted L1 sequence, if available) as candidate supporting reads. PALMER opens the bins in 5′ upstream and 3′ downstream of insertion sequence for each read and then identifies candidate TSD motifs, transductions, and poly(A) tract sequence. All supporting reads are then clustered at each locus and those with a minimum number of supporting events are reported as putative insertions. To improve the accuracy of non-reference MEI sequences derived from individual subreads, which have lower per-read base-pair accuracy, we used local sequence alignments and error-correction strategies. Error correction was conducted by applying CANU to subreads that contain the PALMER MEI sequence, allowing the generation of error-corrected reads that served as inputs for local realignment using minimap2. A second pass of the PALMER pipeline was then executed using these locally aligned error-corrected reads to generate a high-confidence callset of germline non-reference MEIs. Eventually, we used CAP3 to assemble all the MEI sequences reported by the second pass of the PALMER pipeline and obtained a high-confidence consensus contig for each non-reference MEI event.

##### *15.1.3. MEIGA*

**(*Contributor: Bernardo Rodriguez-Martin and Martin Santamarina*)**

*Mapping-based MEI callset construction*

Pre-aligned BAM files from 44 genomes sequenced through PacBio CLR (N=30) and HiFi (N=14) technologies were processed with MEIGA (Mobile Element Insertion Genome Analyzer) version 0.17.0 (https://gitlab.com/ mobilegenomesgroup/architect). MEIGA calls non-reference MEIs, including *Alu*, L1, SVA and ERV-K insertions, based on the identification of two distinct types of supporting read clusters: (i) “spanning-read clusters”, composed by long reads completely spanning the insertion, so they are identified as standard insertions on the reference; and (ii) “clipped-read clusters”, composed by reads that only span one of the inserted sequence ends, so they get clipped during their alignment in the reference. Typically, short insertions are primarily supported by spanning reads, with the number of clipped reads increasing as the insertions are larger in size. Spanning and clipping clusters supporting the same insertion event are grouped based on RO into metaclusters. Then, high-quality consensus sequences for each insertion event are obtained by a read-polishing approach with Racon v1.4.3. To search for candidate MEI events, these sequences are aligned using minimap2 v2.17 against a database of transposable element sequences, containing representative sequences for all human *Alu*, L1, SVA and ERV subfamilies. Based on sequence alignment hits, candidate MEIs are called and multiple insertion features are inferred, including orientation, insertion length, 5' inversion and truncation lengths. Unaligned sequence ends are interrogated for polyadenylate (PolyA) tracts and 5' or 3' transductions. Transductions are small tracks of nonrepetitive DNA sequences mobilized through L1 or SVA retrotransposition that can be used as barcodes to trace L1 or SVA insertions to individual source SVA loci (*54–57*). Finally, candidate MEIs not supported by at least four reads, with less than 60% of their sequence resolved or without polyA signal, are filtered out to generate a high-quality set of non-reference insertions. MEIs identified along all 42 germline genomes were clustered using a breakpoint offset of ≤50 bp and the MEI supported by the highest number of reads (spanning plus clipped reads) in each cluster was selected as representative to generate a nonredundant MEI site list. In those cases where the MEI was detected both in HiFi and CLR samples, the events identified in the HiFi samples were prioritized.

*Assembly-based MEI callset construction*

Phased assembly SV calls (PAV), in the form of a multisample VCF, were processed using a custom pipeline derived from MEIGA (https://github.com/brguez/HGSVC2) to detect non-reference *Alu*, L1 and SVA insertions. First, candidate retrotransposition events were identified by searching for poly(T) and poly(A) tails at the 5' and 3' ends of assembled inserted sequences for PAV insertion calls. Poly(A/T) tails were required to be at least 10 bp in size, have a minimum purity of 80%, and be at a maximum distance of 30 bp to the insert end. Then, inserted sequences for candidates were realigned using minimap2 (*26*) v2.10 into a database of consensus L1, *Alu*, SVA and ERV sequences. Sequence alignments are chained based on complementarity in order to identify the minimum set of nonoverlapping alignments that contribute to resolve the maximum percentage of the inserted sequence. Based on the alignment chain, MEIs are called and multiple insertion features are inferred, which includes orientation, insertion length, 5' inversion and truncation lengths. Unresolved sequence 3' ends are interrogated for Poly(A/T) tracts using the same criteria described above. MEI calls harboring a single tract are classified as “solo” insertions, while events with multiple tracts are considered potential 3' transduction events with the transduced sequences in between. Candidate 3' transduced sequences together with unresolved 5' ends for each MEI are aligned into the reference genome using BWA-mem 0.7.17 (*26*, *58*) to search for 5' and 3' partnered transductions. 5' and 3' transductions calls are made if at least 75% of the target sequence aligns on the reference with a mapping quality over 30, respectively. Candidate “solo” and partnered-transductions with less than 60% of their sequence resolved are filtered out to generate a high-quality set of non-reference insertions. Subfamily for L1, *Alu* and SVA inserts was inferred using two distinct strategies. For L1 events, subfamily assignment was performed through the identification of subfamily diagnostic nucleotide positions on their 3' end (*43*, *59*). L1 integrations bearing the diagnostic “ACG” or “ACA” triplet at 5,929-5,931 position were classified as “pre-Ta” and “Ta”, respectively. Ta elements were subclassified into “Ta-0” or “Ta-1” according to diagnostic bases at 5,535 and 5,538 positions (Ta-0: G and C; Ta-1: T and G). Elements that did not display any of these diagnostic profiles could not be assigned to a particular category, and their subfamily status remained undetermined. For *Alu* and SVA, the assembled inserts were processed with RepeatMasker v4.0.7 to determine the subfamily. If multiple RepeatMasker hits were obtained, the one with the highest Smith-Waterman score was selected as representative. In addition, the coding potential for each L1 sequence was assessed in every of the six potential reading frames. Codon equivalencies followed the Standard Genetic Code currently adopted by NCBI, considering AUG, UUG and CUG as starting codons, and UAA, UAG and UGA as terminators (*60–62*). Every candidate open reading frame (ORF) identified through this strategy was compared with reference L1 ORF1 and L1 ORF2 in terms of 1) length and 2) sequence identity, before being catalogued as one of the two main components of the L1 coding system (*63*). Reference amino acid sequences encoded by L1Hs ORF1 and ORF2 proteins were obtained from Uniprot (*63*, *64*). To end, a multi-sample VCF containing MEI calls together with all collected pieces of annotation was generated as output.

#### *15.2. MEI integration and PAV annotation*

***(Contributors: Weichen Zhou and Ryan Mills)***

To gain a more comprehensive landscape of MEIs in human genomes and better understand the assembly-based method (PAV) on MEI calling, we carried out a integration analysis for MEIs across different platforms/pipelines, including: MELT (independent caller for Illumina), PALMER (independent caller for PacBio mapping-based), and MEIGA_PAV (pre-filtered PAV assembly-based callset with MEIGA annotation) .

MEI calls were pre-merged at the caller level before MEI integration. We then stratified the information of MEIs from four callers into three tiers: Tier 1, the MEI type/family, the insertion orientation/strand, the insertion site, the caller; Tier 2, the insertion length, the structures (e.g., 5' inverted sequence, 3' transduction, intact ORFs), the consensus contig; Tier 3, detailed information at every single discovery sample for each call (e.g., VNTR variation in SVAs). We first merged the Tier 1 information with the exact MEI family, insertion strand, and an overlap of insertion sites in bin sizes of ±50 bp. A CIPOS (Confidence Interval POSition) information is provided showing the difference between the original insertion sites from callers and the merged site. Callers and the number of callers were also recorded. The consensus contigs in Tier 2 are integrated by priorities as follows: Assembly HiFi, Assembly CLR, CLR with local assembled strategy. Structures/characteristics of MEIs are then identified based on the consensus contigs. Tier 3 information was not included in the merged VCF but kept in the sample-level VCFs.

In the integrated callset, there are 11,882 MEIs, including 9,516 *Alu*s, 1,646 L1Hs, 688 SVAs, and 32 HERV-Ks from three different methods. 5,499 (57.8%) *Alu*s are called by at least three callers, 1,889 (19.9%) are called by two callers, 2,128 (22.4%) are called by one single caller, and 1,215 out of 2,128 are singleton events. 653 (39.8%) L1Hs are called by three callers, 445 (27.0%) are by two callers, 548 (33.3%) are only called by one, and 308 out of 548 L1Hs are singletons. There are 337 (49.0%) SVAs called by three callers, 174 (25.3%) called by two callers, 177 (25.7%) SVAs are called by one, and 95 out of 178 SVAs are singletons (Fig. S34a).

To annotate the final PAV callset, the similar strategies (for the Tier 1 and partial Tier 2) were used to compare with the prior integrated callset. Eventually, 9,950 non-reference MEIs, including 8,110 *Alu*s, 1,248 L1Hs, 589 SVAs, and three HERV-Ks were annotated in the final PAV PacBio assembly-based callset. Considering the intersection with two independent callers (MELT and PALMER), there are 6,615 (66.5%) calls that are supported by two callers, including 5,587 *Alu*s, 671 L1Hs, and 357 SVAs; and 2,424 (24.4%) calls are supported by one caller, including 1,843 *Alu*s, 422 L1Hs, 156 SVAs, and three HERV-Ks. Overall, 90.8% of the PAV calls have evidence from at least one or more of independent callers in the prior MEI integrated callset (Fig. S34b).

#### *15.3. Characteristics of MEIs and analysis*

##### 15.3.1. Sequence-resolved full-length L1s

(**Contributors: Bernardo Rodriguez-Martin and Martin Santamarina**)

The human reference genome contains almost one million copies of the L1 element, representing ~17% of the genome (*44*). The vast majority of L1s (99%) are inactive due to mutations, internal rearrangements and truncation or inversion of their 5' ends (*44*). A limited set of L1s remain potentially retrotransposition competent (i.e., active) as they are full-length and have two intact ORFs (*65*). Given that full-length L1s (FL-L1s) are progenitors of newly acquired insertions, a major overarching aim in L1 research has been their identification and characterization. During the last decade, population-scale studies have detected thousands of novel L1 polymorphisms in distinct human populations (*30*, *35*, *51*). However, as these efforts have relied on short-read sequencing, a technology that only provides partial inserted sequences, the repertoire of competent L1s in the population remains largely unexplored. Here, we leveraged the unique opportunity provided by the high-quality L1 sequences derived from PacBio assemblies to create the largest collection of sequence-resolved FL-L1s to date and to investigate their activity potential.

The MEI callset includes 1,170 L1 non-reference insertions. Although most are extremely truncated and inverted on their 5' ends, a large proportion (28%; 329/1,170) are FL-L1s (Table S24). The large majority (78%; 257/329) of FL-L1s have both ORFs intact and, therefore, are potentially competent copies (Fig. S35). A minority of them only have ORF1 (12%; 40/329) or ORF2 (6%; 19/329) intact, while only 4% (13/329) have truncating mutations in both ORFs. Then, we looked at the ratio of substitution rates at both ORFs, denoted dN/dS, for FL-L1s (Fig. S35). Remarkably, ORF1 and ORF2 have on average dN/dS of 0.71 and 0.56, which represents moderate levels of negative selection against mutations truncating L1 machinery if compared to the dN/dS of 0.1 found in protein coding genes. This observation may suggest the existence of selective forces behind the reported high enrichment on competent copies among FL-L1s. Overall, each individual carries on average between 50 and 70 non-reference FL-L1s (Fig. S35), which is in agreement with current estimates for active source L1s (*55*, *66*). In addition, we aimed to collect all known active FL-L1s. With this purpose, we generated a database containing 198 FL-L1 copies reported to be active (Table S25) based on in-vitro retrotransposition assays (*65*), plus germline and somatic 3' transduction detection in three large collections of germline (1000GP) (*30*) and cancer genomes (PCAWG, TUBIO) (*54*, *55*). The majority of active copies (72%; 142/198) are included in our resource and, therefore, now sequence resolved. Interestingly, 19% (27/142) of them have at least one ORF disrupted, which may indicate that these copies are still operative via trans-complementation (i.e., using the machinery from another intact L1) (*67*). Subfamily classification of sequence-resolved active L1s based on diagnostic nucleotides reveals that they are enriched in the youngest Ta-1 subfamily (56%; 79/142), which is consistent with the idea that Ta-1 is the current replicating dominant subfamily (*59*). Finally, we built a phylogeny containing all sequence-resolved active L1s in our collection Fig. S36) using the consensus sequence for *Pan troglodytes*-specific L1s (L1Pt). Inferred L1 ages were consistently associated with distinct L1 features as subfamily status, allele frequency and transduction activity rate. While older sequences were usually fixed and lowly active loci belonging to the pre-Ta and Ta-0 subfamilies, younger L1s were enriched in highly polymorphic and active copies belonging to Ta-1 subfamily. The copy at 2q24.1 is a particularly interesting representative of the second subgroup. It is a rare variant (minor allele frequency [MAF] <1) that has been reported to be extremely active both in the population (*30*) as somatically in cancer genomes (*54*, *55*). Indeed, The L1 copy at 2q24.1 ranks as the seventh most active copy out of 124 in a recent Pan-cancer study (*66*). In addition, this copy displays a quite remarkable pattern of activity, previously termed as Plinian, which involves occasional massive bursts of somatic retrotransposition in cancer genomes (*66*). This behavior was hypothesised to be characteristic of very young L1 copies (*66*), what appears to be the case as this sequence, with an estimated age of ~0.3 myr, is one of the youngest in the dataset. Overall, this collection of sequence-resolved FL-L1s contributes to largely expanding the repertoire of potentially active L1s in humans and provides hints regarding the evolutionary history of these sequences.

##### 15.3.2. Phylogenetic analysis and age estimation for active sequence-resolved L1s

(**Contributor: Martin Santamarina**)

DNA sequences for 142 human-specific (L1Hs) and L1Pt [GenBank: KF661301.1] full-length L1 elements were aligned using MUSCLE (*68*) v3.8 with default number of iterations. Manual inspection of the multiple alignment was performed with Jalview (*69*) v2.11 in order to remove short upstream and downstream spurious sequences. L1 phylogenies were built using the following R packages: Ape (*70*) v5.4 and Phangorn (*70*, *71*) v2.5. Nucleotide substitution model selection was performed with phangorn::modelTest. GTR + G + I was set as the best model for tree inference according to both AIC (Akaike Information Criterion) and BIC (Bayesian Information Criterion) scores. Maximum Likelihood tree was estimated with phangorn::optim.pml function, starting from an initial distance-based tree. Tree topology was adjusted via nearest neighbor interchange (*72*). After Maximum Likelihood Optimization, a midpoint rooting strategy was followed, rendering L1Pt as the expected tree outgroup. Non-parametric bootstrap analysis with 1000 replicates was done in order to determine the node consistency along the inferred tree. For display purposes, only values informative of high support (>=80%) were labeled to the tree nodes.

The average of L1 elements could be estimated by measuring the level of sequence divergence from each subfamily consensus sequence (*73*, *74*). We used DECIPHER v2.14 to generate a consensus sequence for each L1Hs subfamily (pre-Ta, Ta1 and Ta0) from our set of active elements. For each element, we identified the number of nucleotide substitutions acquired since the ancestral state represented by this consensus. Using an L1-specific mutation rate of 0.25% per million years (*59*), and assuming constant and neutral rate of evolution, we obtained rough estimates about the age of individual L1 elements.

##### 15.3.3. A map of active source SVA loci in diverse populations

(**Contributors: Bernardo Rodriguez-Martin and Martin Santamarina**)

While recent studies have created comprehensive catalogues of active L1 sequences and have investigated their activity profiles both in the population (*30*) and in cancer tissues (*30*, *54*, *55*, *75*), little is still known regarding the repertoire of active SVA copies. Here, we mined SVA sequence-resolved insertions to search for SVA-mediated transductions and created a catalogue of source SVA loci active in the population.

We processed all 540 sequence-resolved SVA insertions identified across the 66 assembled haplotypes to search for transduced non-repetitive bits of DNA at their 5' or 3' ends. 15% (77/540) of SVA insertions correspond to transductions (Table S26). Interestingly, 5' transductions (45) were more abundant than 3' transductions (32) as opposed to L1s, which are known to primarily mediate 3' transductions (*54*, *76*). Transduction length oscillated between 30-1,789 bp with a median of 230 bp. SVA-transductions originated from 56 source SVA loci (Table S27). Source SVAs were both present (26) and absent (30) on the reference genome. Interestingly, The majority of source loci (87%, 47/54) belong to the youngest human-specific SVA-E and SVA-F subfamilies. We observed big differences in activity amongst source SVAs, finding that only 5 source SVAs accounted for 27% (21/77) of all SVA-mediated transductions detected in the HGSVC2 dataset (Fig. S37). Among these, one copy at 14q11.2 was particularly active as shown by 7 transductions. These observations resemble the activity patterns described for source L1s (*30*, *54*, *55*) and support the idea that SVA may follow a similar propagation model, with a subset of highly active copies driving the bulk SVA retrotransposition in the population. Retrotransposons can mobilize coding sequences to new genomic loci via transductions (*54*, *57*). Interestingly, we found one instance of exon shuffling where the complete exon of *HGSNAT* is mobilized to 5q31.2 by a SVA source element located in the adjacent intron in antisense orientation (Fig. S38). Finally, analysis of transduced sequences led to the discovery of two intriguing integrations at 1q21.3 and 17q25.1, respectively. Both instances consist of a full-length SVA together with a chain of three non-repetitive DNA pieces interleaved by poly(A) stretches on the SVA 3' end (Fig. S39). Each of these ambiguously aligns into a different chromosomal position. This singular sequence pattern is consistent with the existence of a chain of three consecutive SVA transductions. The first jump corresponds to the mobilization of a source SVA from 3p21.1 to 1q24.2. Then, this new copy mobilizes itself to 1q25.3, together with a 280 bp transduced sequence (Jump-2). Last, the derived copy mediated two transductions to 17q25.1 (Jump-3a) and 1q21.3 (Jump-3b), respectively. In addition, we decomposed each transduced sequence into all possible 61-mers and searched for them into a k-mer database derived from all 1000GP Phase 3 WGS. Consistent with the sequential acquisition of transduced sequences, the average number of samples where k-mers are found for each transduced sequence sharply decreases from the first (2,175), second (1,306) to the last transduction hit (1,044). The differences in sample counts may also reflect the existence of long time frames between each event, particularly pronounced between the first and second transductions.

##### 15.3.4. VNTR distributions in SVAs

***(Contributors: Weichen Zhou and Ryan Mills)***

An SVA is a composite noncoding retrotransposon, which likely uses the L1 machinery for its mobilization into human genomes (*77*). It consists of a hexameric CCCTCT repeat on the 5' end, followed by two antisense *Alu*-like fragments, a variable number of GC-rich tandem repeats (VNTR), a SINE-R sequence, a canonical polyadenylation signal AATAAA, and a poly(A) tract (*77*). Few studies have the privilege to investigate the variation of VNTR regions in SVAs before the advent of long-read sequencing technologies, given it's a high-repetitive region containing multiple copies of a 35-50 bp repeat (*78*).

To define the copy numbers of VNTRs in both reference and non-reference SVAs, at the individual level, we used the sequences directly from the assemblies of discovery samples (27 CLR samples and 11 HIFI samples). We are able to resolve their expansions (duplications and deletions) by unequal homologous recombination at individual and population level as well. The locations of reference SVAs are obtained from RepeatMasker track from UCSC, and the insertion sites of non-reference SVAs are based on PacBio assembly callset (PAV). All sequences of SVAs were obtained from consensus contig of each individual assembly, stratified into subfamilies (SVA_F, SVA_E, SVA_D, etc.) by using the diagnostic nucleotides, and then divided into different populations to conduct the population level analysis. The sample frequency of SVAs are calculated based on the number of discovery samples for each event.

We eventually obtained the assembly sequences that can be used for downstream analysis for 271 reference SVAs and 435 non-reference SVAs among the discovery samples (27 CLR samples and 11 HIFI samples). In the Fig. S40, we observed a) increase variable length of VNTRs in non-reference SVAs as compared to reference SVAs (p-value < 1x10-5, student's t-test, two-sided), and b) reference SVAs have a higher sample frequency than non-reference SVAs (89.1% vs. 17.0%, p-value < 1x10-5, student's t-test, two-sided). We also observed that in the non-reference SVAs, the youngest non-reference SVAs (SVA_F) showed the most variable VNTR copy numbers compared to the non-reference SVA_Es, than the non-reference SVA_Ds. In addition, we observed the VNTR length distributions in non-reference SVAs across five superpopulations and found that the AFR population had more length variability (the difference between maximum and minimum: 257.8 bp ± 338.2) comparing to that seen in the other superpopulations (210.0 bp ± 225.0). Our hypotheses for the different length variations of the VNTR in SVAs, between reference and non-reference are: a) SVA elements harbor copy number variations of VNTR in their interior regions that have been associated with changes in local gene expression (*79*), thus those copy numbers in VNTR tend to be conservative by time under the pressure of the purifying selection and fewer copy number variations are left in the reference (older) SVAs than those in the non-reference (younger) SVAs; b) in the VNTR region of SVAs, there is a sequence, GGGGGGTCAGCCCCCC, that can lead to a structure of imperfect palindrome which may result in VNTR deletion or its copy number variation during DNA replication (*77*, *79*). Because of the burden of nucleotide substitution by time, the reference SVA would have less possibility to have an intact segment of this palindrome sequence, which may have lower chance of the DNA replication problem caused by the palindrome and thus less copy number variations happening in the VNTR region of the reference SVAs than that in the non-reference SVAs.

##### 15.3.5. Distributions of poly(A) tract and EN cleavage site sequences

***(Contributors: Weichen Zhou and Ryan Mills)***

The length of the poly(A) tracts can represent the mobilization activities of non-reference retrotransposons (*80*–*82*). With the help of assembled sequences, we are able to take a close look at the exact poly(A) tract length without worrying about the ambiguous mapping of the short reads. Meanwhile, we are able to identify the sequence distribution of endonuclease (EN) cleavage sites for the first time in such large scale population-level samples for different subfamilies of retrotransposons (*83*–*85*), where L1 retrotransposition machinery occurs.

We divided the non-reference MEI into subfamilies: *Alu* (AluYb8, AluYa5, AluY, and AluS), L1Hs (L1Ta, L1Ta-1, L1Ta-0, and L1PreTa), and SVA (SVA_F, SVA_E, and SVA_D) (Fig. S41). We compared in a pairwise manner, the overall length of the poly(A) tracts for each subfamilies in the *Alu*, L1Hs, and SVAs. In the L1Hs set, we observed that the poly(A) length in the non-reference calls of the L1Ta subfamily are significantly longer than the ones in L1Ta-0 or L1PreTa (p-value < 0.05, student's t-test, two-sided). We made a similar observation when we compared the poly(A) tract length in the non-reference calls of the AluYb8 or AluYa5 subfamilies with those belonging to the AluS subfamily (p-value < 0.05, student's t-test, two-sided). We did not find anything significant in the non-reference SVA subfamilies. The results that younger subfamilies of non-reference *Alu*s and L1Hs have longer poly(A) tracts indicate that they would have a better chance to be retrotransposed by L1 machinery and tend to be more active in the genomes. We further investigated the EN cleavage sites of all subfamilies. By building the sequence logos for all different subfamilies among diverse populations, we observed a preference for the EN cleavage sites 5'-TTTT/AA-3', as seen in previous studies (*83*–*85*). Meanwhile, we also observed that the signals of this pattern are getting weaker as the subfamily of MEIs are getting older, indicating the degeneration of this pattern in the retrotransposition history or the slightly different EN cleavage preferences for older subfamilies.

### 16. Nonredundant callsets, filtering, and properties

***(Contributors:* Peter Audano)**

#### *16.1. Nonredundant callset merging*

*Merging independent PAV samples.* We produced PAV calls independently for 32 genomes (11 HiFi and 21 CLR), excluding replicate CLR and child samples, and applied QC steps per sample (see section 10 and Table S28). We applied the three-step method (see section 9) to merge them in a stepwise process similar to previous studies (*21*) where samples are iteratively added to the nonredundant set. To seed the merge, the first sample is taken as the nonredundant set. The next sample is intersected with the nonredundant set using the three-step approach for SVs and indels or exact matches for SNVs (see section 9.2). Variants that intersect the callset are added as annotations to the existing calls and variants that do not intersect are appended to the callset. This process is repeated until all samples have been merged.

The end product is a set of nonredundant variants (Fig. 2c) that may be supported from one or more samples. The sample and variant ID are retained so the record can be linked back to the supporting calls without ambiguity. The lead variant, which represents the whole set, is from the first sample with the variant. Since merging order matters, we added HiFi trio parents first (CHS, PUR, then YRI), then we added supplementary HiFi samples ordered by contig N50 (best assembly N50 first), and finally CLR samples ordered by contig N50.

The average callset growth per haplotype is estimated by using the number of singleton variants that would be added to the nonredundant callset by each of the 64 haplotypes if it was merged last. Compared to out-of-Africa haplotypes, African haplotypes are adding 2.21× SVs, 3.70× indels, and 2.97× SNVs (Table S29).

The average callset growth rate per sample by new homozygous variants was estimated by examining callset growth from homozygous variants in the final sample if each sample was added last. Compared to out-of-Africa haplotypes, African samples are growing the callset with new homozygous variants at a rate of 1.80× SVs, 2.40× indels, and 2.74× SNVs per sample (Table S29).

Compared to previous long-read studies (*21*, *22*, *86*–*88*), we added 30,149 insertions and 15,734 deletions that were not previously sequence resolved (Fig. S42).

***Table S28. Variant discovery from phased assemblies***

| | | | | SV | | | | Indel | | | | | |
|---|---|---|---|---|---|---|---|---|---|---|---|---|---|
| | | | | Insertions | | Deletions | | Insertions | | Deletions | | SNV | |
| Sample | Pop | Superpop | Sex | N | Mbp | N | Mbp | N | Mbp | N | Mbp | N | Ti/Tv |
| HiFi | | | | | | | | | | | | | |
| HG00512 | CHS | EAS | M | 14,055 | 7.80 | 8,937 | 5.23 | 367,796 | 1.26 | 370,030 | 1.23 | 3,620,202 | 2.06 |
| HG00513 | CHS | EAS | F | 13,990 | 7.87 | 9,073 | 5.23 | 371,578 | 1.27 | 374,452 | 1.24 | 3,654,446 | 2.05 |
| HG00731 | PUR | AMR | M | 14,009 | 7.86 | 8,867 | 5.16 | 379,989 | 1.30 | 379,972 | 1.26 | 3,693,860 | 2.06 |
| HG00732 | PUR | AMR | F | 14,314 | 8.19 | 9,089 | 5.09 | 391,334 | 1.34 | 376,235 | 1.27 | 3,746,657 | 2.06 |
| NA19238 | YRI | AFR | F | 16,238 | 8.78 | 10,655 | 5.96 | 440,260 | 1.52 | 457,660 | 1.51 | 4,496,371 | 2.06 |
| NA19239 | YRI | AFR | M | 15,889 | 8.56 | 10,665 | 5.92 | 435,745 | 1.50 | 451,879 | 1.49 | 4,466,461 | 2.06 |
| HG02818 | GWD | AFR | F | 16,072 | 8.79 | 10,794 | 6.10 | 436,495 | 1.48 | 449,078 | 1.46 | 4,469,835 | 2.06 |
| HG03125 | ESN | AFR | F | 16,355 | 8.67 | 10,775 | 5.90 | 438,853 | 1.49 | 449,021 | 1.46 | 4,470,531 | 2.06 |
| HG03486 | MSL | AFR | F | 16,263 | 8.85 | 10,915 | 6.18 | 442,090 | 1.50 | 452,407 | 1.48 | 4,512,588 | 2.06 |
| NA12878 | CEU | EUR | F | 13,954 | 7.35 | 8,931 | 5.48 | 367,945 | 1.24 | 372,590 | 1.22 | 3,643,864 | 2.06 |
| NA24385 | Ashk. | EUR | M | 14,025 | 7.73 | 9,054 | 5.19 | 372,475 | 1.26 | 373,972 | 1.24 | 3,659,919 | 2.06 |
| | | | **Mean:** | 15,015 | 8.22 | 9,796 | 5.58 | 404,051 | 1.38 | 409,754 | 1.35 | 4,039,521 | 2.06 |
| | | | **Median:** | 14,314 | 8.19 | 9,089 | 5.48 | 391,334 | 1.34 | 379,972 | 1.27 | 3,746,657 | 2.06 |
| CLR | | | | | | | | | | | | | |
| HG00096 | GBR | EUR | M | 14,232 | 8.25 | 9,068 | 5.20 | 401,627 | 1.27 | 351,303 | 1.17 | 3,610,388 | 2.06 |
| HG00171 | FIN | EUR | F | 14,436 | 8.35 | 9,264 | 5.68 | 401,057 | 1.28 | 355,985 | 1.18 | 3,639,951 | 2.06 |
| HG00864 | CDX | EAS | F | 14,524 | 8.09 | 9,371 | 5.62 | 386,596 | 1.26 | 364,378 | 1.20 | 3,662,845 | 2.06 |
| HG01114 | CLM | AMR | F | 14,468 | 8.02 | 9,346 | 5.51 | 404,492 | 1.27 | 380,236 | 1.21 | 3,672,056 | 2.06 |
| HG01505 | IBS | EUR | M | 14,441 | 8.34 | 9,077 | 5.69 | 387,770 | 1.25 | 360,609 | 1.19 | 3,626,081 | 2.06 |
| HG01596 | KHV | EAS | M | 14,329 | 8.39 | 8,873 | 5.41 | 413,216 | 1.28 | 351,974 | 1.16 | 3,597,839 | 2.06 |
| HG02011 | ACB | AFR | M | 16,197 | 9.13 | 10,722 | 6.01 | 438,444 | 1.43 | 412,011 | 1.36 | 4,329,583 | 2.06 |
| HG02492 | PJL | SAS | M | 13,993 | 8.46 | 8,994 | 5.50 | 372,637 | 1.21 | 347,792 | 1.15 | 3,565,097 | 2.06 |
| HG02587 | GWD | AFR | F | 16,824 | 9.48 | 11,033 | 6.19 | 434,588 | 1.42 | 442,539 | 1.41 | 4,431,941 | 2.06 |
| HG03009 | BEB | SAS | M | 14,045 | 8.20 | 8,930 | 5.55 | 423,231 | 1.28 | 332,469 | 1.13 | 3,577,520 | 2.06 |
| HG03065 | MSL | AFR | M | 16,365 | 9.45 | 10,844 | 6.21 | 449,936 | 1.47 | 420,840 | 1.38 | 4,438,462 | 2.07 |
| HG03371 | ESN | AFR | M | 16,457 | 9.39 | 10,740 | 6.40 | 442,339 | 1.44 | 415,725 | 1.36 | 4,385,788 | 2.06 |
| HG03683 | STU | SAS | F | 14,616 | 8.62 | 9,427 | 5.45 | 389,074 | 1.28 | 370,133 | 1.23 | 3,744,652 | 2.06 |
| HG03732 | ITU | SAS | M | 14,443 | 8.63 | 9,192 | 5.60 | 397,082 | 1.27 | 372,999 | 1.21 | 3,690,391 | 2.06 |
| NA12329 | CEU | EUR | F | 14,543 | 8.63 | 9,216 | 5.71 | 392,273 | 1.26 | 357,642 | 1.19 | 3,655,436 | 2.06 |
| NA18534 | CHB | EAS | M | 14,075 | 8.32 | 9,000 | 5.58 | 399,009 | 1.26 | 353,915 | 1.17 | 3,614,601 | 2.06 |
| NA18939 | JPT | EAS | F | 14,506 | 8.43 | 9,162 | 5.56 | 394,775 | 1.27 | 356,188 | 1.18 | 3,638,870 | 2.06 |
| NA19650 | MXL | AMR | M | 14,425 | 8.18 | 9,221 | 5.68 | 421,126 | 1.30 | 377,226 | 1.21 | 3,675,028 | 2.06 |
| NA19983 | ASW | AFR | F | 16,525 | 9.16 | 10,627 | 6.31 | 439,469 | 1.44 | 407,811 | 1.35 | 4,368,448 | 2.06 |
| NA20509 | TSI | EUR | M | 13,920 | 8.28 | 8,759 | 5.29 | 405,990 | 1.26 | 359,209 | 1.16 | 3,560,979 | 2.06 |
| NA20847 | GIH | SAS | F | 14,726 | 8.89 | 9,283 | 5.71 | 406,889 | 1.29 | 376,523 | 1.22 | 3,728,072 | 2.06 |
| | | | **Mean:** | 14,861 | 8.60 | 9,531 | 5.71 | 409,601 | 1.31 | 374,643 | 1.23 | 3,819,716 | 2.06 |
| | | | **Median:** | 14,468 | 8.43 | 9,221 | 5.62 | 404,492 | 1.28 | 364,378 | 1.20 | 3,662,845 | 2.06 |
| | | | **Mean:** | 14,914 | 8.47 | 9,622 | 5.67 | 407,693 | 1.33 | 386,713 | 1.27 | 3,895,274 | 2.06 |
| | | | **Median:** | 14,456 | 8.41 | 9,219 | 5.61 | 403,060 | 1.28 | 374,212 | 1.22 | 3,673,542 | 2.06 |

On average, we discovered 24,563 SVs, 794,406 indels, and 3.9 million SNVs per genome after initial QC on sample callsets. Merging and additional QC yielded a nonredundant set of 107,590 SVs, 2.3 million indels, and 15.8 million SNVs (Table S28).

#### *16.2. Post-merge filtering*

*Filtering nonredundant SVs.* For SVs, we required support from one or more of PBSV, Bionano, DeepVariant, PAV run with LRA alignments (PAV-LRA), LRA-based alignments with CIGAR string variant discovery (LRA Assembly), Subseq, or 61-mer support over breakpoints (details above for each of these methods). For 61-mers, we counted support if non-reference k-mers were identified at variant breakpoints and the k-mers were supported by Illumina reads. If the lead variant passed one or more of these filters, it was accepted into the final callset. By manually inspecting SVs ≥ 50 kbp (n = 96 SVs), we identified 54 assembly errors that resulted in large deletions where the largest 9 (690 kbp - 17.3 Mbp) were all false calls.

*Filtering nonredundant indels and SNVs.* For indels, we required support from two or more of DeepVariant, GATK, PAV-LRA, or 61-mer support over breakpoints (same 61-mer approach as for SVs). For SNVs, we required support from two or more of DeepVariant, Longshot, GATK, PAV-LRA, or 61-mer support over breakpoints (same 61-mer approach as for SVs).

*Assessing orthogonal callset support.* To assess support from other callers and to compare HiFi and CLR, we intersected variant calls from multiple sources for HG00733 (Fig. S43). There is a higher error rate among CLR samples, which is seen in the PAV callset and previous studies (*3*, *21*). This CLR error is reflected low validation rates by subseq for calls only seen by PBSV CLR. Outside annotated tandem repeats (UCSC track), the trends are more stratified with very little subseq support for any single caller, including variants called by PAV in both HiFi and CLR. Many of these variants will not make it through the filtering stages (Fig. S44).

#### *16.3. Callset quality estimates*

*SV variant support.* Including orthogonal callsets used for filtering, we added support from DeBreak, Inspector, and an LRA-based validation method (LRA Chaisson) to estimate merged callset FDR (Table S30). Single-method FDR estimates ranged from 35.3% (DeBreak) to 7.38% (Inspector). It is interesting to note that PAV-LRA alone estimates 25.8% FDR, which highlights the importance of the aligner for variant discovery and likely improvements yet to be made to LRA. The callset was filtered with some of these methods (excluding the ones added for QC) and required support from two or more sources, we estimate 5.27% FDR. Inspector alone predicts an FDR of 6.73%.

#### *16.4. Variant frequency distributions*

***(Contributors:* Peter Audano, Jan Korbel, Tobias Rausch)**

In previous work, we observed a linear relationship in the distribution of SV variant allele frequencies on a log scale (*51*). Before applying quality filters to the merged callset, we observed a visible increase in low allele frequency variants. Although the allele frequency was not considered while making callset filtering decisions, low-frequency variants were filtered more heavily (Fig. S45), and the distribution shifts closer to a linear pattern.

#### *16.5. Identify SV clusters - hotspot analysis*

***(Contributors:* David Porubsky)**

*Hotspot definition.* To identify SV clusters, we selected the middle position of each SV and submitted it to the ‘hotspotter’ function from the primatR package (*89*) (parameters: bw=200000, num.trial=1000). This function searches for regions of increased density of SV midpoints around the genome by using the density function to perform a KDE (kernel density estimation). A p-value was calculated by comparing the density profile of the genomic events with the density profile of a randomly subsampled set of genomic events (bootstrap support).

*Hotspot analysis.* Using the total number of SVs (≥50 bp) detected across all autosomes and chromosome X (n=105,327), we found SVs significantly (p-value < 0.001) clustered at the terminal 5 Mbp region of each chromosome with ~4-fold enrichment (Fig. S46 a). To alleviate this bias, we removed SVs that reside at the terminal 5 Mbp (remaining SVs n=73,105) of each chromosome and detected a total of 221 SV hotspots (Fig. S46 b). As expected, we found these hotpots significantly enriched for SDs (p-value < 0.001, ~6.6-fold enrichment) (Fig. S46 b left). Of these, 112 hotspots have been previously identified (*51*) while the rest are novel (n=109) (Fig. S47a,c). As expected, a number of hotspots overlaid with regions of lower mapping quality from our phased contigs against GRCh38. (Fig. S47 b). In total, we report 278 SV hotspots sites including 57 terminal hotpots (Fig 2d, Table S33) covering ~279 Mbp of the genome (Fig 2d inset, Fig. S48).

#### *16.6. HLA analysis*

***(Contributors:* David Porubsky)**

We focused specifically on the major histocompatibility complex (MHC on chr6:28,510,120-33,480,577) to assess the level of completion and variation. This region is sequence resolved (MAPQ60 contig alignments) with a median of two assembled contigs per haploid assembly covering 98.36% of the locus. The assembly with the lowest amount of coverage still resolves 96.35% of the sequence in a single assembled contig, corresponding to missing sequence of ~181 kbp. (Table S34). The high completeness of this biologically relevant region allows us to compare the level of divergence across all phased genomes (Fig. S49). At the SNV level, we observed three clusters of high sequence divergence that are also sites of previously detected SV hotspots (Fig. S50) and there is an indication of a population-specific variation (Fig. S51a, arrowhead). We further investigated the effect of this variation on exons. As expected, we found exons of HLA genes carrying more alternative alleles which resulted in an elevated number of non-synonymous amino-acid changes (Fig. S51b,c,d). At the SV levels, we found the HLA region to cluster in about seven distinct haplotypes (Fig. S52) with a relatively even distribution of the haplotypes across populations (Fig. 2e, Fig. S53).

#### *16.7 Shared variants*

***(Contributors: Peter Audano)***

We defined a set of shared variants that were discovered in all haplotypes. For this analysis, we excluded variants that were not callable in at least one haplotype (i.e., AN=64 and AF=1). In total, 1,573 SVs were shared, 1,310 (83.3%) were genotyped with 100% allele frequency in population samples by PanGenie (see section 19.2), 1,548 (98.4%) were supported by at least one call in our previous work, and 1,118 (71.1%) matched a known shared variant (*21*). Of the

2,238 shared SVs reported by (*21*), 1,118 (50.0%) were also annotated as shared in this study. These variants represent a refined set of reference errors or extreme minor alleles (*21*).

### VALIDATION

#### 17. SV validation by reads and contigs

We noted 7 chromosomes affected by a bug in QC code that dropped heterozygous calls for some chromosomes. This is fixed in our final callset, which is not yet released. Affected chromosomes are HG00513 HiFi (chr21), HG00514 HiFi (chr9), HG00733 HiFi (chr21), HG00732 CLR (chr3), HG02492 CLR (chr10), HG03009 CLR (chr10), and NA20509 CLR (chr17).

We also identified 10 misassemblies that appear to be large deletion SV calls ranging from 12.4 Mbp to 565 kbp. These caused PAV to drop calls intersecting the deletions, which lost sensitivity for some samples in these regions. We added a feature to PAV to flag and ignore known assembly errors and generated a new callset, which is also in the unreleased final callset. Identified misassemblies are chr13-46437077-DEL-17373620, chr15-72620759-DEL-2696104, chr9-83827217-DEL-1974975, chr1-222496126-DEL-1472517, chr17-30655670-DEL-1416038, chr22-23332910-DEL-1330209, chr10-45681225-DEL-1137697, chr16-29396552-DEL-841511, chr7-65062627-DEL-689786, and chr16-16198408-DEL-564551

##### *17.1. Subseq validations with raw reads*

***(Contributors:* Peter Audano)**

As a QC metric, we assessed raw-read support for SVs with the minimap2 alignments used for PBSV. For each SV, a window around the SV is defined by extending the breakpoints upstream and downstream of the SV (padding) with a larger pad for larger SVs (SVLEN <100: 20 bp, 100 <= SVLEN < 200: 25 bp, 200 <= SVLEN < 500: 40, 500 <= SVLEN < 1 kbp: 100 bp, 1 kbp <= SVLEN < 2 kbp: 200 bp, 2 kbp <= SVLEN < 4 kbp: 250 bp, SVLEN >= 4 kbp: 300 bp). We found these pad values to work well for each variant class based on expected distributions and agreement with other QC metrics, such as orthogonal caller support.

For each padded window, all reads traversing the window were identified and the length of the reads mapping to the window (distance in read space between the bases mapped to the first and last base of the window) using custom code (subseqfa, https://github.com/EichlerLab/seqtools) (Fig. S54). If there were fewer than four reads spanning the window, we did not attempt to make a validation call. If four or more reads spanned the window and at least two supported the SV call allowing up to a 50% shift in the expected size of the window, then the variant call was validated by this method. Although we have PBSV calls from the same alignments, this method allows SV calls to be fragmented into multiple insertion or deletion events relative to the SV call.

There is a clear difference between HiFi and CLR validations with HiFi showing a much cleaner pattern (Fig. S55). CLR is much more difficult to validate and we expect a higher validation error rate with CLR with more calls falsely invalidated.

#### *17.2. LRA read and contig alignment support*

(***Contributors: Mark Chaisson***)

In order to filter out calls due to misassembly or misalignment, SVs were detected from raw reads. A combination of two alignment methods was used, LRA v1.0 (https://github.com/ChaissonLab/LRA/archive/v1.0.0.tar.gz), and minimap version 2.17-r941. An aligned read was considered to be supporting the variant if an SV was detected in the read with length between 50% and 200% of the length of the assembly based call (e.g., 50% length overlap), and with breakpoints within 1 kbp. Calls are considered validated if at least four reads support the call in either dataset (Fig. S56). By including both alignment approaches, additional support is given for SV support.

As expected, the application of two alignment methods can provide support for more variants than one alignment method alone. Considering SV calls from the CLR assembly of the Puerto Rican individual, HG00733, 89.3% of calls had at least four reads supporting an SV from either method, compared to 85.1% of calls having similar support by only LRA and 86.7% of calls are supported by minimap2 alignments. An example of a call that is not supported by either alignment method is shown in Fig. S57.

The average percentage of variants detected from HiFi-based assemblies supported by either method was 92.3%, with 86.7% of variants supported by LRA and 88.6% of variants supported by minimap2. The average percentage of variants detected from CLR-based assemblies was lower: 85.0% of variants had support from at least one alignment method (80.1% LRA and 82.3% minimap2). The full per-assembly rate is shown in Fig. S58 given in Table S35 for normal, wide, and broad support. The fraction of variants that are not supported generally increases as the coverage decreases; however, this is a combination of the effect of assembly quality and number of reads available to support a call. The genome-wide SV distribution that is not supported by reads is shown in Fig. S59 and S60.

#### *17.3. Comparison of phase 1 and phase 2 callsets.*

Variant calls are made from different alignment approaches between phase 1 and phase 2. Various stringencies of relative length and proximity to SV were used to measure the number of unique calls in the phase 1 versus phase 2 callsets. The SV callsets from phase 1 generated only by local assembly (asm), as well as the multi-technology, merged, nonredundant (mnr) were compared to PAV calls made from HiFi and CLR assemblies. The phase 1 callsets were filtered to ensure at least four reads supporting the call, similar to the calculated rate for PAV calls. The counts of the calls that are compared are in Table S36.

Under the most lenient overlap, where a call of the same type is made within 1 kbp, an average of 1,279 variants are missing from the PAV CCS assembly-based callsets, and 1,274 variants are missing from the PAV CLR assembly-based callsets when compared to the asm callsets. Also, 3,618 and 3,584 calls are missing from the CCS/CLR callsets, respectively, when compared to the mnr callset (Table S37). Requiring one call to be within 25% length of the other, the number of calls that are unique to phase 1 increases such that 2,141 and 2,144 calls are missing from the CCS/CLR callsets compared to the asm callset, and 4,795/4,775 calls are missing from the CCS/CLR callsets compared to the mnr callset. The number of calls unique to phase 1 increases as the length of call increases (Table S37). This indicates a difference in the representation of calls, rather than missed calls, and so subsequent analyses could use the metric of any proximal SV (within 1 kbp) of the same type, without requiring a specific length.

We categorized each variant with the following genomic features: overlapping SDs, tandem repeats (TR), and overlapping assembled contigs (ASM) to determine if variants were more likely to be missed if they were in that genomic feature (variants will be de-facto missed if they do not overlap an assembly contig). Across all classes of comparisons, 493-837 variants were missed that overlap SDs relative to the fraction of calls made in SD in the phase 1 calls. This represents 22-44% of calls missed in phase 2, and a 2.7- to 5.8-fold enrichment relative to all missed calls. Tandem repeats, while representing a greater number of variants that are missing from PAV assembly calls (914-2,587 variants/callset), also represent the majority of calls made from long-read sequencing and are not overrepresented as a fraction of missed calls (enrichment 0.71-1.09).

The majority of variants in the phase 1 callsets that are missing from the assembly-based callsets are annotated as overlapping assemblies. Each assembly was mapped using LRA and a base was considered overlapping by an assembly if an alignment from either haplotype overlapped. Depending on the callset, between 92-96% of variants overlapped an assembly. This indicates that missing calls are due to either filtering parameters from assembly-based alignments or incomplete representation of haplotypes in the assemblies.

#### *17.4. Variant concordance by reads, contigs and assemblies (Inspector)*

(***Contributors: Zechen Chong, Yu Chen***)

To further validate SV calls with raw reads, Inspector version v1.0.1 (https://github.com/Maggi-Chen/Inspector) was applied on all CLR and CCS samples. Raw reads were aligned against all assembled and phased contigs with minimap2 version 2.15-r905. Read alignments at each SV breakpoint were used for evaluating the confidence of SV calls. A highly confident SV call should have reasonable depth (half of the sequencing depth due to haplotype-resolved assembly) and no indel or clipped alignments. Strong indel signals at the SV breakpoint often suggest the presence of assembly errors. Extremely high or low read-alignment depth suggests assembly collapse or expansion, respectively. Variants that passed both filters were classified as “PASS”, while variants with indels or clipped alignment near breakpoints were marked as “NoisyAlign”. Variants were marked as “HighDepth” or “LowDepth” when read alignment depth at SV breakpoints were higher than two times of the mean sequencing depth or <= 3, respectively (Fig. S61).

#### *17.5. Raw read variant and breakpoint concordance.*

(***Contributors: Kai Ye, Jiadong Lin and Xiaofei Yang***)

To examine the breakpoint accuracy of the SV callset, MUMMer (https://github.com/mummer4/mummer) and NucDiff (https://github.com/uio-cels/NucDiff) based raw reads realignment were applied to all CCS and CLR samples. Since insertions only have one breakpoint, we used the reported breakpoint location plus the insertion length as another breakpoint for further evaluation. All SV spanning reads were extracted from the pbmm2-aligned BAM file with SAMtools. If no reads were fetched, this call was marked as “NoRawReads”. On the contrary, PAV calls with spanning reads were realigned with MUMMer and their corresponding breakpoint type was classified by NucDiff. We first examined whether the breakpoints of each call could be supported by the realigned breakpoint. If none of the breakpoint differences between the PAV call and the realigned breakpoint are smaller than the allowed distance (allowed_dist=500), this PAV call was marked as “NoRealignSupport”. For the rest of the PAV calls, we then grouped the breakpoints from each realigned read based on their position (100 bp difference allowed) and type to make realigned calls. For a PAV call and its corresponding realigned call, if the position difference of the two breakpoints are both smaller than maximum breakpoint shift (allowed_shift), this PAV call is considered as “Precise”, otherwise it is considered as “Imprecise”. As a result, PAV calls were marked as “NoRawReads”,“NoRealignSupport”, “Precise” and “Imprecise” (Fig. S62).

For the HiFi samples, the realignment validation rate for both DEL and INS is around 75% for each sample (Fig. S63 A). We observed approximately 1.5% of the calls as “NoRawReads”, indicating the alignment issue of HiFi reads in these regions. But we found some regions could be aligned with NGMLR (Fig. S64 A). By setting the allowed_dist = 500 bp, ~9% and ~7% of the DELs and INSs, respectively, became labeled as “NoRealignSupport”, with ~80% of the calls being smaller than 100 bp (Fig. S64 B). We further annotated these SVs with RepeatMasker and TRF (Fig. S64 C), leading to ~65% calls inside the repeat regions and more than half of these calls (58%) found in STR regions (repeat unit <=7 bp). For the CLR samples, we used the same parameters (Fig. S63 C) and noticed the percentage of precise calls to be highly variable among CLR samples, whereas HiFi-based calls showed consistently better performance for both DEL and INS (~75%). One possible explanation for this observation is that the segment chaining algorithm used by MUMMer and increased sequencing errors by CLR lead to inconsistent performances for the CLR dataset.

#### *17.6. Breakpoint analysis*

**(*Contributors: Sushant Kumar and Mark Gerstein*)**

We further characterized the underlying mechanism for a subset of our SV (deletions and insertion) callsets after identifying their precise breakpoints. For the mechanism analysis, we applied the mechanism detection module of our previously published BreakSeq method. Briefly, the mechanism module of BreakSeq utilizes distinct sequence-based signatures within and around the breakpoint junction of a given SV to detect its underlying mechanism. For instance, it uses the RepeatMasker program to detect extensive coverage of tandem repeats and low-complexity regions within a given SV to classify them as a variable number of tandem repeats (VNTRs). Similarly, SVs belonging to the nonallelic homologous recombination (NAHR) class show extensive homology at their breakpoint junction (consisting of flanking sequences of a given SVs). SVs that align to known interspersed MEIs in the genome are considered transposable element insertions (TEIs). It further subclassifies TEIs based on whether a given SV is aligned to a single (STEI) or multiple (MTEI) transposable element insertion. Furthermore, the SV mechanism pipeline also identifies processed pseudogenes that originate due to TEI-associated mechanism. Finally, SVs for which BreakSeq extracts the flanking sequence, but lack any of the signatures mentioned above, were classified to the nonhomologous recombination (NHR) mechanism class.

In this work, we applied BreakSeq's mechanism pipeline in two distinct settings by varying the homology length cutoff (>=50 bp & >=200 bp) and flanking sequence length (200 bp & 1 kbp) for detecting NAHR events (Table S38 A-B). Furthermore, in the first setting (homology length >=50 bp), we grouped VNTR events with repeat length >=50 bp and NAHR events together to define a new homology-associated event class. In contrast, VNTRs (with repeat length >=50 bp and covering 50% of SV regions) constituted the newly defined homology-associated category in our second analysis set. Finally, deletions and insertions that overlap with the assembly-resolved MEIs (generated as part of the current work) were classified as TEIs. We performed these analyses on the subset of SVs, for which precise breakpoint definition was available. Overall, we applied our mechanism analysis pipeline on 97,162 SVs (33,531 DELs and 63,631 INSs). Furthermore, we also quantified mechanism distribution differences for a subset of SVs with precise breakpoints that overlapped with the short-read-based SV call set (Table S38 C-D).

Our mechanism characterization of the long-read-based SVs indicated different mechanism distribution for deletions and insertions (Fig. S65). For instance, while we observed that a large fraction of deletions and insertions belonged to the homology-associated mechanism class, the homology-associated mechanism had, on average, a relatively higher contribution to deletions (mean fraction value of 0.628) than insertions (mean fraction value of 0.512). In contrast, a somewhat smaller fraction of deletions (mean value of 0.14) and insertions (mean value of 0.18) belonged to the NHR mechanism class. We observed a distinct mechanism composition pattern for our SV dataset with a more stringent definition of NAHR events (homology length >=200 bp) (Fig. S68). While, the homology-associated and NAHR mechanism class together, on average (mean value of 0.187), had a slightly higher contribution toward describing deletions compared to NHR class (mean value of 0.172), a large fraction of insertions belonged to the NHR mechanism category (mean value of 0.266).

Additionally, we compared the underlying mechanism distribution for deletions and insertions of different lengths (Fig. S66). We classified deletion and insertions into three distinct categories; 1) length below 200 bp (lt200bp), 2) length between 200 and 1 kbp (lt1K), and 3) length above 1 kbp (ge1K). Overall, we found that SVs with length below 200 bp primarily belonged to homology-associated class (mean fraction value of 0.8). In contrast, a relatively larger fraction of SVs belonging to lt1K and ge1K categories corresponded to TEIs (mean value of 0.377) and NHR (mean value of 0.416) mechanism classes. Interestingly, we observed similar mechanism contribution pattern differences for distinct length categories of insertions and deletions (Fig. S66). However, with a more stringent definition for the NAHR mechanism category, we observed few differences in length distribution pattern for deletions and insertions (Fig. S69). Notably, we observed that a relatively higher fraction of deletions belonging to the lt1K category corresponded to NHR mechanism class (mean value of 0.42). In contrast, the NHR mechanism contributed similarly to insertions corresponding to lt200bp (mean fraction value of 0.302) and lt1K (mean fraction value of 0.305) categories.

Finally, we also quantified relative contributions of different mechanism categories for SVs overlapping with distinct functional elements in the genome (Fig. S67). Overall, we found a large fraction of functionally relevant SVs belonged to the homology-associated and NHR classes. Interestingly, we found that homology-associated mechanisms explained most of the functionally pertinent deletions and insertions. In contrast, the NHR mechanism had a more considerable contribution to the emergence of functionally relevant deletions. We also observed subtle variability in the contribution of distinct mechanisms toward deletion overlapping with different functional elements. For instance, a relatively larger fraction of deletions overlapping with DHS (mean value of 0.439) and regulatory elements belonged to NHR mechanism class (mean value of 0.431). In contrast, a relatively larger fraction of deletions overlapping with ncRNA (mean value of 0.49) and UTRs (mean value of 0.59 for 3' UTR and 0.66 for 5' UTR) belonged to the homology-associated class. Similarly, a larger fraction of insertions overlapping with intronic regions (mean value of 0.51), 5' UTRs (mean value of 0.527), and CPG (mean value of 0.53) regions belonged to the homology-associated mechanism category. Interestingly, with a stringent definition of NAHR mechanism class, we saw a completely distinct contribution distribution for deletions and insertions overlapping with different functional elements (Fig. S70). In particular, we found that a relatively large fraction of functionally relevant deletions belonged to the NHR class compared to the homology-mediated mechanism class. Deletions overlapping with DHS (mean fraction value of 0.465) and CCRE elements (mean fraction value of 0.467) primarily belonged to the NHR mechanism category. We observed a relatively higher contribution from homology-associated mechanism category toward functionally relevant insertions compared to deletions. However, there was a sharp decline in the NAHR mechanism's contribution toward explaining insertions overlapping with various functional elements in the genome.

#### 18. Analyzing non-reference k-mers using 3,202 genomes

(***Contributors: Tobias Rausch***)

Using 3,202 Illumina sequenced (30X) samples, including 2,504 unrelated samples from the 1000GP and an additional 698 related samples, we created a database of non-reference k-mers absent in the GRCh37 reference of the 1000GP (hs37d5). For each sample, we first error corrected the raw sequencing reads with Lighter v1.1.2 (*13*) using a k-mer length of 23 and then computed a compacted de Bruijn graph with BCALM 2 v2.3.0 (doi.org/10.1093/bioinformatics/btw279) using a k-mer length of 61. We then used dicey v0.1.6 from the GEAR genomics framework (doi.org/10.1186/s12864-020-6635-8) to compute k-mer hash sums and to extract non-reference k-mers for each sample by means of depleting the reference k-mers present in hs37d5. Each sample contained ~270–370 million non-reference k-mers, with African samples showing an increase of non-reference k-mers (Fig. S71).

We then aggregated all non-reference k-mers in a single database, separately for the set of 2,504 unrelated samples from the 1000GP and the additional 698 related samples.

We then utilized this resource to evaluate the assemblies and assigned each non-reference k-mer from the assembly its occurrence count in the 1000GP. We interpreted a count of zero, a so-called missing, non-reference k-mer, as a likely assembly error or as a k-mer that cannot be sequenced by Illumina due to very high or low GC content. Notably, the CLR assemblies showed, on average, 8.3% missing, non-reference k-mers compared to 3.7% for the CCS assemblies, reflecting the higher sequencing accuracy of CCS compared to CLR.

We also used this non-reference k-mer resource to assess whether assembly-based SV calls are supported or unsupported in the Illumina data using a simple approach that interrogates SV breakpoint k-mers of length 61 for being present or absent in the non-reference k-mer database of the 3,202 Illumina-sequenced genomes.

### GENOTYPING AND ASSOCIATION

#### 19. Genotyping Illumina Genomes

##### *19.1. Genotyping Paragraph*

(***Contributors: Wayne Clarke & Mike Zody***)

Using Paragraph (*90*), a sequence graph-based genotyping tool, we genotyped 107,590 SVs using a per-sample approach. Paragraph requires, as input, a) a VCF file of variants, b) a reference sequence FASTA, c) a manifest containing the path to the sample BAM file and d) additional information including the read length, average read depth, read length, and the sample sex. Paragraph outputs a genotype for each of the input variant sites, which we merged into a multi-sample VCF. Variants were evaluated for Mendelian consistency and Hardy-Weinberg equilibrium (HWE), with sites violating Mendelian consistency and sites with extreme deviations from HWE being removed, resulting in 96,145 SVs carried forward.

#### *19.2. Genotyping PanGenie and graph construction*

**(*Contributors:* Jana Ebler)**

*PanGenie.* We used our recently developed method PanGenie (*91*) to showcase the utility of our haplotype resources for improving short-read-based genotyping. This method uses a panel of known population haplotypes and sequencing reads in order to genotype a new sample. In a first step, PanGenie constructs an acyclic and directed graph which represents the genetic variation across the input panel haplotypes. Variants are represented as bubble structures and each haplotype constitutes one path through this graph. In the next step, k-mers uniquely characterizing variant alleles are determined and counted in the reads. These counts provide information about the presence or absence of variant alleles in the sample to be genotyped. However, SVs might be located in genomic regions poorly covered by unique k-mers. Therefore, our model additionally leverages the global haplotype structure provided by the input haplotype panel. This enables genotype imputation, which is especially useful in such difficult-to-access regions.

PanGenie is based on a hidden Markov model (HMM), which integrates information from k-mer counts and known haplotypes. It aims at constructing the unknown sample haplotypes such that they best explain the observed read k-mer counts and, at the same time, are mosaics of the panel haplotypes. Genotype likelihoods are computed based on the Forward-Backward algorithm. This model has been demonstrated to provide ultra-fast genotyping by bypassing the expensive read alignment step, making it well suited especially for genotyping large sets of samples. Leveraging the provided haplotype structure enables one to access regions of the genome otherwise hard to interrogate from short reads only. Since the base version of PanGenie becomes rather slow for panels with more than 15 individuals (30 haplotypes), we extended it to efficiently handle larger panels for genotyping. Per default, this extended version of PanGenie randomly divides the panel samples into groups of 15 haplotypes and computes genotype likelihoods based on an HMM, constructed only on these paths. The likelihoods obtained for all subsets in this way are later summed up and normalized in order to obtain the final genotype likelihoods from which to make a genotype prediction.

Using this approach, we genotyped all 3,202 sequenced samples in order to demonstrate the utility of our method for large, short-read-based studies. For each sample, we provided PanGenie with short-read sequencing reads (in FASTQ format) as well as a multisample VCF file containing phased variant calls from the 64 assemblies. This input VCF file is derived from the merged PAV callsets produced for SNVs, indels and SVs. At first, we removed all positions at which more than 20% of the panel haplotypes carried a missing allele. Furthermore, we kept only variants located on chromosomes 1-22 and chromosome X for genotyping. We then created a multiallelic VCF representation in which overlapping variants are combined into multiallelic positions. Some variants in the input PAV callsets were overlapping on the same haplotype (e.g., an SNV inside of a deletion). We removed such conflicts by setting the corresponding alleles to missing ("."). In this way, the VCF represents an acyclic and directed variation graph representing the panel genomes. Table S39 provides an overview of the number of variants obtained. As an output, PanGenie generates a VCF file that contains a genotype for all these variants. We merged all 3,202 sample VCFs with genotypes into a single multisample VCF, and additionally produced a biallelic version of this VCF.

We started with a pilot set consisting of 300 individuals selected from the 3,202 samples. This subset was constructed by randomly choosing 20 trios from each of the five superpopulations (AFR, AMR, EAS, EUR, SAS). We then ran PanGenie in order to genotype all 15.5 M SNVs, 1.03 M indels and 96.1 K SVs across these 300 samples. For comparison, we additionally ran Paragraph to derive genotypes for all SVs. As a QC measure, we computed allele frequencies across all 200 unrelated samples from the PanGenie and Paragraph genotypes and compared them to the allele frequencies derived from the PAV calls for all 64 assembly haplotypes. For Paragraph, we observed allele frequency correlations (Pearson correlation) of 0.61 and 0.54 for deletions and insertions. For PanGenie, these values are 0.85 and 0.86. Fig. S72 indicates that Paragraph tends to genotype variants as heterozygous and especially struggles with genotyping insertions. We concluded that PanGenie seems to be more suitable for genotyping our SV calls and proceeded with genotyping all SNVs, indels, and SVs across the 3,202 samples using PanGenie. We determined the number of heterozygous SVs for each population from these genotypes (Fig. S73) and observed higher numbers for the African populations, reflecting their increased genetic diversity. We also computed allele frequency correlations from the allele frequencies derived from the PanGenie genotypes for all 2,504 unrelated samples and the PAV calls. We observed correlations of 0.98, 0.95, and 0.85 for SNVs, indels, and SVs, respectively. However, these numbers indicate that there are variants for which PanGenie and PAV allele frequencies differ significantly. In order to filter out such potentially wrong genotyped calls, we defined a strict subset of variants based on statistics we computed from the genotypes of the 3,202 samples. We defined the five filters listed below:

- **ac0_fail**: A variant was genotyped as absent (genotype 0/0) by PanGenie in all 3,202 samples (i.e., the allele frequency is zero).
- **mendel_fail**: There are 602 trios among the 3,202 genotyped samples. For each variant, we counted the number of trios with Mendelian-consistent genotypes. Here, we only take trios with at least two different genotypes into consideration, meaning we skip trios in

which all samples were typed as 0/0, 0/1 or 1/1 respectively. A variant fails this filter if the Mendelian consistency is below 90%.

- **gq_fail**: At least 200 low-quality genotypes were reported by PanGenie.
- **nonref_fail**: All panel samples were genotyped as homozygous reference.
- **loo_fail**: In addition to genotyping the 3,202 samples, we conducted a leave-one-out experiment, in which we repeatedly take out one of the panel samples from the input and use PanGenie to genotype it based on the remaining samples in the panel. We then compare the predicted genotype to the left-out, ground-truth genotype of the sample. This enables us to compute the genotype concordance across all panel samples at each variant position. This filter fails if the genotype concordance of the panel samples is below 80%.

We applied these filters to all SNVs, indels, and SVs genotyped by PanGenie. For SVs, 16,343 out of 60,238 insertions (27%) failed the "ac0_fail" filter and were genotyped with an allele frequency of zero across all 3,202 samples. For deletions, 8,948 out of 35,862 (25%) were genotyped as homozygous reference in all samples. About 57% of these variants are rare and were carried by only a single haplotype in the input panel. Such variants are, in particular, difficult to genotype by a panel-based approach like PanGenie, especially if the k-mer counts show no strong indication for the presence of an allele in a sample. To obtain a filtered callset, we removed all variants for which at least one of the five filters failed. This leads to a rather stringent, but high-quality set of genotypes that serves as a basis for further analysis. Our filtered set contains 12,283,650 SNVs (79%), 705,893 indels (68%), and 24,107 SVs (25%).

We provide callset statistics for this strict set in Figure 5b and Figs. S75-77. Figure 5b shows the allele frequencies obtained for SVs across the PanGenie genotypes, as well as the corresponding allele frequencies in the input panel haplotypes. The allele frequencies for PanGenie were computed based on all 2,504 unrelated individuals part of the 3,202 samples. For both variant types, insertions and deletions, the allele frequencies match well with very few outliers, indicating that the genotypes are of good quality. We obtained an allele frequency correlation of 0.99 (0.98 for deletions, 0.99 for insertions). Likewise, allele frequency correlations for SNVs and indels in this filtered set are 0.99 and 0.99, respectively. We also investigated the relationship between variant length and allele frequencies across the PanGenie genotypes. The peaks in Fig. S75 showed a clear tendency of *Alu* insertions towards lower allele frequencies, while it seems to be the opposite for *Alu* deletions. We suspect that this behavior is caused by Alu insertions present in the reference genome. Figs. S76 and S77 show the relationship between Fst values of the five superpopulations (AFR, AMR, EAS, EUR, SAS) and the length of the SVs. For each superpopulation, Fst was computed between individuals that are part of it and the union of the remaining populations. We observed higher Fst values for African and East Asian populations indicating higher degrees of differentiation among these populations.

*Defining a lenient set.* For SVs, our filtered set contains only 25% of all input variants. In addition to this strict set, we also defined a larger, more lenient set of SVs using a machine-learning approach based on support vector regression. Our model is designed to assign scores close to -1 to poorly genotyped SVs and scores around 1 to those passing all our filters. For training, we used our strict SV set as "true positives". The "true negatives" were defined as all variants genotyped with an allele frequency larger than zero ("ac0_fail" did not fail) that failed at least three of the remaining filters. The regression is based on 70 features collected from the PanGenie genotypes, the leave-one-out experiments and concordances with k-mer-based presence/absence genotyping [see Section 18]. We predicted regression scores for all yet unlabeled SVs (with an allele frequency > 0). We then created more lenient callsets by adding those variants to the strict set for which the scores were above a certain threshold. We investigated three different cutoffs: -0.5, 0.0 and 0.5. Table S40 contains corresponding callset statistics. As expected, numbers improve as we increase the cutoff that we use to define a lenient callset. While allele frequency correlations are 0.86 and 0.85 for insertions and deletions in the unfiltered set, they reach levels between 0.95 and 0.98 when applying the different cutoffs. Likewise, average genotype concordances of the PanGenie genotypes with the PAV calls for the assembly samples increase, reaching levels above 94% for cutoff 0.5. While containing more than 50% more variants, statistics for the lenient set with cutoff 0.5 are close to the strict set for which we observed correlations of 0.99 for insertions and deletions, as well as genotype concordances of 96.9% and 96.1%, respectively.

### 20. RNA-seq analysis

#### *20.1. Read QC and mapping*

**(*Contributors: Arvis Sulovari, Marc Jan Bonder, Jan Korbel)***

To gain insight into the molecular impact of indels (insertions and deletions <50 bp), SVs (≥50 bp) and SNVs, we jointly analyzed two lymphoblastoid cell lines (LCLs) RNA-seq datasets, the GEUVADIS study (n=462) as well as 33 newly generated, deeply sequenced (>230M output/sample) 1000GP LCL RNA-seq samples (d1KG). To minimize batch effects, we re-analyzed the raw data.

First, low-quality reads were removed and both adapters and low-quality bases were trimmed using Trim Galore! (*92*) (v0.6.5), a wrapper around FastQC (*93*) and cutadapt (*94*). Next, trimmed and QC reads were aligned using STAR (*95*) (version: 2.7.5a), using a custom two-pass alignment. All alignments were performed using the GRCh38.p12 reference genome and NCBI Homo sapiens gene annotation (R109.20190125) annotations. The first pass was done on all samples, using the default parameters as proposed by ENCODE (c.f. STAR manual) with the exception of the two-pass alignment flag. Based on this mapping and an initial expression quantification based on featureCounts (*96*) (subread 2.0.1), we performed sample QC.

We chose to remove samples with: 1) high multimapping percentage (> 20%), 2) low % Picard usable bases (<85%), and 3) samples failing per_base_n_content (“fail”) from FastQC.

Afterwards, we performed a PCA on the edgeR (*97*) normalized initial featureCount quantification and removed outliers based on the first two principal components (per dataset). Keeping in 411 GEUVADIS samples and all of the deeply sequenced 1000GP samples. Using VerifyBamID (*98*) (v 1.1.3) we verified the links between the genotype and the RNA-seq data, using the PanGenie (*91*) SNP genotypes (Section 19) as a reference. First, we verified low FREEMIX scores for the passed QC samples (<0.1) and filtered links between genotype and expression data (CHIPMIX <0.02). This removed an additional 14 GEUVADIS samples, leaving 397 samples. In the d1KG samples, we found three straight sample swaps and three samples matching better to the genotype of another with every genotype observed once. No sample swaps were observed in the GEUVADIS cohort.

After sample QC, we obtained 430 high-quality samples, which underwent the second-pass mapping in STAR. In this second pass, we used the same settings and added the WASP output mode and then used an aggregated splice junction database from the first round of alignments. Specifically, we aggregated the junction databases of the HQ samples and included junctions that: 1) were observed in at least 40 RNA-seq runs, 2) had support of at least 45 uniquely mapping reads, and 3) had spanning reads support of at least 20 bases before and after the splice junction, In total, we included 135,356 high-quality *de novo* LCL-based splice junctions to the STAR junction database. To enable the WASP mode, which requires SNP information, we chose to use a custom (single) VCF with all SNP variants observed in the PanGenie genotyping (Section 19) with a MAF of 1% and above in the samples with RNA-sequencing data. After mapping, we filtered the reads based on the WASP filters, allowing reads without a WASP flag or the flag "vW:i:1".

##### *20.2. Expression and splicing quantification and normalization*

Based on the filtered alignments, we quantified both gene-expression levels and splicing levels. The gene-expression levels are based on featureCounts, using "-p -B -C --primary -p", to count only primary alignments, not count chimeric reads and count-only reads where both end of paired end sequencing data are mapped. The featureCount read counts were transformed to transcripts per million and normalized for library size using edgeR, before log transformation.

#### 21. eQTL and GWAS

##### *21.1. eQTL and sQTL analyses*

**(Contributors: *Marc Jan Bonder, Arvis Sulovari, Yang Li, Jan Korbel*)**

To assess the impact of SVs, indels, and SNVs on gene expression and splicing, we performed expression quantitative trait locus (eQTL) using the high-quality genotypes from PanGenie (Section 19). The eQTL and sQTL analyses were carried out using a linear mixed-model approach implemented in LIMIX (*99–101*) and employing an IBD matrix to account for population structure. The IBD matrix was calculated with PLINK (*102*) (v1.0.7), using pruned SNP variants (MAF>5%) from the PanGenie imputation. The QTL mappings were run in a mega-analysis setting, analysing the 430 expression samples (397 GEUVADIS and 33 d1KG), originating from 427 unique donors, all in one single analysis.

To select the number of expression covariates to include in the eQTL mapping, we initially performed a focused eQTL mapping analysis for chromosome 2, on SNPs with at least a MAF of 10% and within a window of 10 kbp around the genes. By varying the number of hidden factors taken along in the eQTL mapping (including 0-100 principal components [PCs] by steps of 5), we observed that using 60 PCs maximizes the fraction of genes with an eQTL (eGenes) discovered. Therefore, we selected 60 PCs to correct our eQTL map (Fig. S78).

Next, we performed the full *cis*-eQTL mapping analysis linking gene expression levels to all PanGenie-derived high-quality genotypes, within a window of 1 Mbp centered around the transcription start site of a gene and testing all variants with a MAF of 1% and higher and a Hardy-Weinberg equalibrium P $\leq 0.0001$. We simultaneously considered all variants genotyped using PanGenie (i.e., SNVs, indels and SVs). For SVs, we made sure to include all variants intersecting with the test window, in order not to miss potential SV associations. To correct for multiple testing, we used an approach related to FastQTL (*103*), first correcting for genetic variants based on genotype permutations and using a beta approximation to increase precision, followed by a Storey Q-value based correction for the number of tested genes. By mapping the permuted p-value that corresponds to gene level FDR 5%, we derived all eQTLs (for further details on this approach, see (*100*)).

In the eQTL mapping analysis we considered 23,866 genes, which were expressed (i.e., non-zero count) in at least 20% of the samples and which showed an average TPM of at least 0.01 in the 430 post-QC RNA-seq samples. Using this setup, we identified 9,387 lead *cis*-eQTLs using an FDR of 5% and 847,807 *cis*-eQTLs in total. Of the lead eQTLs, 8,640 were SNV-eQTLs (8,284 unique SNVs, 92% of the total *cis*-eQTL signals), 713 are indel-eQTLs (680 unique Indels, length ≤49 bp, 7.6% of total) and 34 are SV-eQTLs (33 unique SVs, length ≥50 bp, 0.36%) (Table S41). Interestingly, the set of lead *cis*-eQTL hits was enriched for indels (Fisher's exact test P-value = 1.92e-6, OR = 1.33 [95% CI: 1.2-1.5]) and SVs (P-value = 0.0436, OR = 1.8 [1.0 - 3.5]), and depleted for SNPs (P-value = 2.7e-7, OR = 0.74 [0.7-0.8]). All association tests were conducted on nonredundant counts of variants, using the expected proportions of SNVs, indels, and SVs from the full set of tested high-quality genotypes. The average indel-eQTL and SV-eQTL lengths were 5 bp and 2,437 bp, respectively. Additionally, eight of the SV-eQTLs (24%) were *Alu* insertions with lengths ranging from 120 bp to 327 bp.

##### *21.2. GWAS intersection and enrichment analysis*

**(Contributors: *Junjie Chen, Chong Li, Xinghua Shi*)**

We examined if PanGenie genotypes and eQTLs were associated with human phenotypes or traits, by intersecting SNV, indel, and SV eQTLs with SNPs previously reported in genome-wide association studies (GWAS). We first aggregated GWAS summary statistics and generated a union GWAS set that contains known GWAS associations (P-value <= 1.0e-6) extracted from the GWAS Catalog (*104*, *105*), Pan-UKB project and UK Biobank (UKBB (Pan-UKB team. https://pan.ukbb.broadinstitute.org. 2020.)) and PhenoScanner V2 (*106*, *107*). Since GWAS SNPs in UKBB and PhenoScanner are based on GRch37, we lifted over their coordinates to GRch38 and mapped them to corresponding rsIDs using dbSNP version 151 (*108*). In total, the union GWAS set contains 1,887,371 autosomal SNPs associated with 6,528 traits (Fig. S79).

For SNVs, we counted those variants that are in the union GWAS dataset, while for indels and SVs, we counted those indels/SVs that have at least 1 bp overlap with any GWAS SNP in the union set. We found that 1,132,074 SNVs, 2,339 indels, and 3,564 SVs overlapped or intersected with GWAS signals. For eQTLs, we found 285,568 SNV eQTLs, 735 indel eQTLs, and 470 SV eQTLs overlapped or intersected with GWAS signals (Table S36 and Fig. S80). Next, we assessed the linkage disequilibrium (LD) between any SNV/indel/SV with known GWAS SNPs within a 1 Mbp window using Plink v1.90b6.10 (*102*). We identified 537,725 SNV eQTLs, 7,664 indel eQTLs, and 133 SV eQTLs that are in high LD ($r^2$ >= 0.8) with GWAS SNPs (Table S42).

We observed that SV eQTLs are enriched for GWAS signals compared with random subsets of SVs that were fed into the eQTL analysis pipeline (P-value < 1.0e-4, Fig. S81). We performed 10,000 random permutations, with each permutation sampling 2,099 random SVs, with the same number as in the SV eQTL set, from the 146,426 SVs that were evaluated in our eQTL analysis. When generating these random SVs to match the distributions of SV eQTLs, we stratified the sampling process so that we controlled for the chromosome sources, SV types, SV sizes, and distances to Transcription Start Sites (TSS) and Transcription End Site (TES) respectively. In particular, SV sizes were matched after logarithm transformation to the base 2 and rounded down, while the distances to TSS/TES were matched after dividing the distance by 1,000 and rounding down. In each permutation, we counted the number of random SVs that overlapped with at least one GWAS SNP in the union GWAS set. We then estimated the statistical significance of the observed overlap of SV eQTL with the union GWAS set, compared with the overlaps from these random permutations.

#### *21.3. GWAS and eQTL co-localization analysis*

**(Contributors: *Junjie Chen, Chong Li, Xinghua Shi*)**

We conducted a GWAS and eQTL co-localization analysis to identify human trait associations driven by eQTLs that may indicate a molecular mechanism induced by genetic variants. We performed this analysis using a Summary-data-based Mendelian Randomization (SMR) test (*109*) based on summary statistics from our eQTL analysis and GWAS summary statistics from the union GWAS set. More specifically, the SMR approach used genetic variants as instrumental variables to calculate two-step-at-least-square estimates and test for the causative effect of the expression level of a gene on a trait without confounding from non-genetic factors. Consequently, Chi-square tests were performed to evaluate the significance of the estimated causal effect of gene expression on an associated trait, and the Benjamini-Hochberg correction (Benjamini, Yoav; Hochberg, Yosef Controlling the false discovery rate: a practical and powerful approach to multiple testing. J. Roy. Statist. Soc. Ser. B 57 (1995), no. 1, 289–300.) was then used to control the multi-test correction. Those gene-trait pairs passed the SMR test implied a potential causal link between the eQTL, gene and the associated trait.

To perform the SMR test, we first mapped SNV/indel/SV eQTLs to GWAS SNPs if the eQTLs have any overlap with GWAS SNPs regarding their chromosomal locations. For each of the 5,305 eGenes with GWAS associations, we collected the summary statistics of SNV/indel/SV eQTLs and those of GWAS SNPs that are located within a 2 Mbp window centered around the midpoint position of the tested gene. We identified 5,296 genes that are associated with 3,976 traits that passed the SMR test (5% FDR, Table S43). Among these genes, 1,178 of them have SV eQTLs and 4,494 of them have indel eQTLs. These observations indicated that SV and indel eQTLs potentially drive the changes of gene expression that lead to variations in the associated traits.

### 22. Ancestry Analysis

**(*Contributors: Rebecca Serra Mari and PingHsun Hsieh*)**

#### *22.1 Local ancestry*

*Local ancestry inference using haplotype-phased assembly, RFMix and an HMM.* We leveraged the haplotype-phased, sequence-resolved assemblies to infer local ancestry along chromosomes using RFMix (*112*) and an HMM. Ancestry information is based on haplotypes from a predetermined set of reference populations. For our analysis, a reference population panel was assembled using published phased genomes from the 1000GP (*1*). To control for potential biases due to the inclusion of admixed individuals in the reference panel, we used ADMIXTURE (*113*) and chose less admixed samples from African (LWK, MSL, GWD, YRI, and ESN; n=472), European (CEU, GBR, FIN, IBS, and TSI; n=381), East Asian (CHB, CHS, JPT; n=185), and South Asian (ITU and STU; n=186) 1000GP samples. In addition, as part of our reference panel, we also included 19 Native American samples from the Simons Genome Diversity Project (*113*, *114*) that show little European ancestry. Note that for the inference of local ancestry in Puerto Rican individuals, we also specifically explored and set the reference panel to be African, European, and Native American populations given the recent demographic history of this population (*115*). To avoid inaccurate ancestry calls, we removed SNVs in known gaps, SDs, heterochromatin, telomeric, and centromeric sequences (GRCh38) from this analysis.

In short, RFMix segments haplotypes from a sample into windows of SNVs and uses a model of random forests based on conditional random fields to determine the ancestry of each genomic window. Analysis was performed with the following flags: -G 15 -e 5 -w 0.4 -n 5 -c 0.2 -s 0.2 --rf-minimum-snps=100 --reanalyze-reference. Note that we chose a node size of 5 to reduce bias in random forests resulting from unbalanced reference panel sizes as suggested in previous studies (*115*). We also developed an HMM-based method to infer the local ancestry of each genomic region of the haplotype-resolved assemblies (Fig. S82; https://github.com/rebeccaserramari/kyoshi). Briefly, the computation of the most likely ancestral population for each genomic position is inferred through an HMM that models the reference haplotypes from a reference panel as hidden state sequences and the target haplotype as the observed sequence and computes a standard forward-backward algorithm on this HMM. At each variant position, the HMM penalizes two types of events: 1) discrepancies between target and reference haplotype and 2) switches to a different reference haplotype happening between two variant positions. As a consequence, the computed probability will be high for reference haplotypes that largely coincide with the target. The outcome of this method is a set of probabilities for underlying ancestries at each variant position on a haplotype. The probabilities of reference samples that belong to the same superpopulation are then added up to get a likelihood for every superpopulation at each position.

We restricted our downstream analysis to a set of high-confidence ancestry markers where the HMM gives evidence for a single ancestry with a probability >90%. To construct the final ancestry callsets for each individual haplotype, we first computed the concordant calls between the RFMix and HMM results. Because the HMM results provide a per-variant ancestry resolution, we re-called the ancestry of a discordant region if there are more than 10 consecutive high-confidence ancestry markers.

To explore the quality of our ancestry inference, we performed the analysis on three trio families where we have both HiFi and CLR data using the parameters described above. In general, our results show a high concordance for ancestry calls between the two datasets (>90%), indicating that the input data type has little impact on the inferred local ancestries (Fig. S83). Our analysis showed that while the ancestry distribution of European and East Asian as well as most of African samples are relatively homogenous, there are clear admixture signals in other continental population samples (Figs. S84 and S85). Consistent with the history of slave trade from Africa to North America, the two African American samples (HG02011 and NA19983) carry large segments with European ancestry (>17%). In addition, the four admixed American samples all carry long-stretch ancestry blocks from African (2-5%), American (6-13%), and European (83-89%) populations, reflecting the colonial history of these populations in America (*104*). Notably, comparing with the Strand-seq inferred recombination breakpoints in the parental chromosomes in the trios, we found that ancestry blocks on the paternal and maternal haplotypes of the child switch at the locations of the crossing-over events in the parental chromosomes (Figure 6B). This highlights the importance of these haplotype-resolved assemblies as a resource for population genetics and evolutionary analyses.

#### 22.2 Variant age estimations

We estimate the age of an SV of interest using the software Relate (*116*). In short, Relate reconstructs the local genealogy of the region of interest using a scalable computation, which guarantees the inferred genealogy exactly producing the observed data. We include SNVs from the 500 kbp sequences flanking individual candidate SVs and remove/mask sites that have missing genotypes. In addition to the SV of interest, to approximate the age of the SV of interest, we also select a focal SNV that is in complete LD ($r^2$=1) to estimate variant age. The mutations are then mapped onto the branches of the resulting local tree using Relate to estimate mutation age.

#### 22.3 Ancestral state determination from primate assemblies

We categorize SVs as ancestral if they were matched to calls in two nonhuman primates (NHPs) (9,829) (*117*, *118*), non-ancestral if they were not seen in both (72,641), and unknown if they could not be confidently assigned (25,120) because they were uncallable (14,865) or differed (10,255) with the latter, dominated by tandem repeats (91%). The allele frequency differed significantly with an average of 0.40 for ancestral and 0.12 for non-ancestral SVs ($p <$ 1e-15, t-test). These polymorphic ancestral variants may be a useful resource for studying recurrence. We observe a significant difference between the numbers of insertions and deletions between ancestral and non-ancestral states with more ancestral deletions and non-ancestral insertions ($p <$ 1e-15, Fisher's exact test (FET)). Reference collapses are known to enrich for fixed insertions (*21*), and we find SV insertions are more likely to be fixed (AF = 1 by PanGenie) than deletions (23% vs 4.6%, $p = 5\times10^{-165}$, FET), but only 0.88% insertions and 0.33% deletions are fixed, which is significantly lower compared to ancestral ($p <$ 1e-15, FET). In total, 1,301 SVs are homozygous for the alternative allele in all 3,202 samples and also exhibit AF=1 in the assembly-based call set, indicating cases where GRCh38 contains errors or includes alleles with very low AF, with the expected bias toward insertions where the reference assembly likely collapsed (1,152 INS, 149 DEL). This is an increase from our previous work where we reported 507 such events (*21*).

#### 22.4 Population Stratification and Population Branch Statistics

To identify variants stratified by population, we computed Fst values for each superpopulation (superpopulation vs. all other samples) and each sample (sample vs. all other samples). We assigned a variant to the population with the maximum Fst value if the value was at least 0.2 and greater than 0.1 from all other populations (Table S43). Similarly, we assigned the remaining variants to superpopulations with the maximum Fst value if the value was at least 0.2 and 0.1 higher than all other superpopulations (Fig. 6c).

To identify variants in regions subject to selection, we computed population branch statistics (PBS) (*119*) for combinations of populations for all SVs within 5 kbp of a gene. For each variant, we found the maximum PBS for each population using all possible combinations of an ingroup (same superpopulation) and an outgroup (different superpopulation) (Table S44). We focused specifically on the top 10 hits per superpopulation (Fig 6d). As expected, there are distinct differences among population groups attributable to bottlenecks and expansions in their recent evolutionary history (Fig S86).

### 23. Functional Annotations

**(*Contributors: Peter Audano*)**

#### *23.1. Functional variant annotations*

Variant calls were intersected with RefSeq annotations (retrieved from UCSC on 2020-06-29) and counted the CDS, UTR, and intron sequence intersected by each variant. Variant calls were also intersected with *cis*-regulatory element (cCRE) annotations against GRCh38 from ENCODE (*120*) (Table S45). We categorized SVs as intersecting promoters (PLS), proximal and distal enhancer-like signatures (pELS and dELS), and CTCF (CTCF-only).

#### *23.2 Triplet repeat expansions*

We ran Tandem Repeats Finder (TRF) (*121*) on SV sequences and searched for perfect repeat copies by comparing the repeat motif to the SV sequence and required that perfect repeats span at least 95% of the SV. We identified 286 repeat expansions and contractions by sample (382 alleles by haplotype) within these 106 sites (Tables S46 and S47).

2. J. M. Zook, D. Catoe, J. McDaniel, L. Vang, N. Spies, A. Sidow, Z. Weng, Y. Liu, C. E. Mason, N. Alexander, E. Henaff, A. B. R. McIntyre, D. Chandramohan, F. Chen, E. Jaeger, A. Moshrefi, K. Pham, W. Stedman, T. Liang, M. Saghbini, Z. Dzakula, A. Hastie, H. Cao, G. Deikus, E. Schadt, R. Sebra, A. Bashir, R. M. Truty, C. C. Chang, N. Gulbahce, K. Zhao, S. Ghosh, F. Hyland, Y. Fu, M. Chaisson, C. Xiao, J. Trow, S. T. Sherry, A. W. Zaranek, M.

Ball, J. Bobe, P. Estep, G. M. Church, P. Marks, S. Kyriazopoulou-Panagiotopoulou, G. X. Y. Zheng, M. Schnall-Levin, H. S. Ordonez, P. A. Mudivarti, K. Giorda, Y. Sheng, K. B. Rypdal, M. Salit, Extensive sequencing of seven human genomes to characterize benchmark reference materials. *Scientific Data*. **3**, 1–26 (2016).

3. M. J. P. Chaisson, A. D. Sanders, X. Zhao, A. Malhotra, D. Porubsky, T. Rausch, E. J. Gardner, O. L. Rodriguez, L. Guo, R. L. Collins, X. Fan, J. Wen, R. E. Handsaker, S. Fairley, Z. N. Kronenberg, X. Kong, F. Hormozdiari, D. Lee, A. M. Wenger, A. R. Hastie, D. Antaki, T. Anantharaman, P. A. Audano, H. Brand, S. Cantsilieris, H. Cao, E. Cerveira, C. Chen, X. Chen, C.-S. Chin, Z. Chong, N. T. Chuang, C. C. Lambert, D. M. Church, L. Clarke, A. Farrell, J. Flores, T. Galeev, D. U. Gorkin, M. Gujral, V. Guryev, W. H. Heaton, J. Korlach, S. Kumar, J. Y. Kwon, E. T. Lam, J. E. Lee, J. Lee, W.-P. Lee, S. P. Lee, S. Li, P. Marks, K. Viaud-Martinez, S. Meiers, K. M. Munson, F. C. P. Navarro, B. J. Nelson, C. Nodzak, A. Noor, S. Kyriazopoulou-Panagiotopoulou, A. W. C. Pang, Y. Qiu, G. Rosanio, M. Ryan, A. Stütz, D. C. J. Spierings, A. Ward, A. E. Welch, M. Xiao, W. Xu, C. Zhang, Q. Zhu, X. Zheng-Bradley, E. Lowy, S. Yakneen, S. McCarroll, G. Jun, L. Ding, C. L. Koh, B. Ren, P. Flicek, K. Chen, M. B. Gerstein, P.-Y. Kwok, P. M. Lansdorp, G. T. Marth, J. Sebat, X. Shi, A. Bashir, K. Ye, S. E. Devine, M. E. Talkowski, R. E. Mills, T. Marschall, J. O. Korbel, E. E. Eichler, C. Lee, Multi-platform discovery of haplotype-resolved structural variation in human genomes. *Nat. Commun.* **10**, 1784 (2019).

10. A. R. Quinlan, I. M. Hall, BEDTools: a flexible suite of utilities for comparing genomic

features. *Bioinformatics*. **26**, 841–842 (2010).

18. V. A. Schneider, T. Graves-Lindsay, K. Howe, N. Bouk, H.-C. Chen, P. A. Kitts, T. D. Murphy, K. D. Pruitt, F. Thibaud-Nissen, D. Albracht, R. S. Fulton, M. Kremitzki, V. Magrini, C. Markovic, S. McGrath, K. M. Steinberg, K. Auger, W. Chow, J. Collins, G. Harden, T. Hubbard, S. Pelan, J. T. Simpson, G. Threadgold, J. Torrance, J. M. Wood, L. Clarke, S. Koren, M. Boitano, P. Peluso, H. Li, C.-S. Chin, A. M. Phillippy, R. Durbin, R. K. Wilson, P. Flicek, E. E. Eichler, D. M. Church, Evaluation of GRCh38 and de novo haploid genome assemblies demonstrates the enduring quality of the reference assembly. *Genome Res.* **27**, 849–864 (2017).

19. 1000 Genomes Project Consortium, G. R. Abecasis, D. Altshuler, A. Auton, L. D. Brooks, R. M. Durbin, R. A. Gibbs, M. E. Hurles, G. A. McVean, A map of human genome variation from population-scale sequencing. *Nature*. **467**, 1061–1073 (2010).

20. N. C. Sheffield, C. Bock, LOLA: enrichment analysis for genomic region sets and regulatory elements in R and Bioconductor. *Bioinformatics*. **32**, 587–589 (2016).

21. P. A. Audano, A. Sulovari, T. A. Graves-Lindsay, S. Cantsilieris, M. Sorensen, A. E. Welch, M. L. Dougherty, B. J. Nelson, A. Shah, S. K. Dutcher, W. C. Warren, V. Magrini, S. D. McGrath, Y. I. Li, R. K. Wilson, E. E. Eichler, Characterizing the Major Structural Variant Alleles of the Human Genome. *Cell*. **176**, 1–13 (2019).

22. J. Huddleston, M. J. P. Chaisson, K. M. Steinberg, W. Warren, K. Hoekzema, D. Gordon, T. A. Graves-Lindsay, K. M. Munson, Z. N. Kronenberg, L. Vives, P. Peluso, M. Boitano, C.-S. Chin, J. Korlach, R. K. Wilson, E. E. Eichler, Discovery and genotyping of structural variation from long-read haploid genome sequence data. *Genome Res.* **27**, 677–685

(2017).

35. R. L. Collins, H. Brand, K. J. Karczewski, X. Zhao, J. Alföldi, L. C. Francioli, A. V. Khera, C. Lowther, L. D. Gauthier, H. Wang, N. A. Watts, M. Solomonson, A. O'Donnell-Luria, A. Baumann, R. Munshi, M. Walker, C. W. Whelan, Y. Huang, T. Brookings, T. Sharpe, M. R. Stone, E. Valkanas, J. Fu, G. Tiao, K. M. Laricchia, V. Ruano-Rubio, C. Stevens, N. Gupta, C. Cusick, L. Margolin, Genome Aggregation Database Production Team, Genome Aggregation Database Consortium, K. D. Taylor, H. J. Lin, S. S. Rich, W. S. Post, Y.-D. I. Chen, J. I. Rotter, C. Nusbaum, A. Philippakis, E. Lander, S. Gabriel, B. M. Neale, S. Kathiresan, M. J. Daly, E. Banks, D. G. MacArthur, M. E. Talkowski, A structural variation reference for medical and population genetics. *Nature*. **581**, 444–451 (2020).

36. G. Ke, Q. Meng, T. Finley, T. Wang, W. Chen, W. Ma, Q. Ye, T.-Y. Liu, in *Advances in Neural Information Processing Systems 30*, I. Guyon, U. V. Luxburg, S. Bengio, H. Wallach, R. Fergus, S. Vishwanathan, R. Garnett, Eds. (Curran Associates, Inc., 2017; http://papers.nips.cc/paper/6907-lightgbm-a-highly-efficient-gradient-boosting-decision-tree.pdf), pp. 3146–3154.

37. R. L. Collins, H. Brand, K. J. Karczewski, X. Zhao, J. Alföldi, A. V. Khera, L. C. Francioli, L. D. Gauthier, H. Wang, N. A. Watts, M. Solomonson, A. O'Donnell-Luria, A. Baumann, R. Munshi, C. Lowther, M. Walker, C. Whelan, Y. Huang, T. Brookings, T. Sharpe, M. R. Stone, E. Valkanas, J. Fu, G. Tiao, K. M. Laricchia, C. Stevens, N. Gupta, L. Margolin, The Genome Aggregation Database (gnomAD) Production Team, The gnomAD Consortium, J. A. Spertus, K. D. Taylor, H. J. Lin, S. S. Rich, W. Post, Y.-D. I. Chen, J. I. Rotter, C. Nusbaum, A. Philippakis, E. Lander, S. Gabriel, B. M. Neale, S. Kathiresan, M. J. Daly, E. Banks, D. G. MacArthur, M. E. Talkowski, An open resource of structural variation for medical and population genetics. *bioRxiv* (2019), p. 578674.

44. E. S. Lander, L. M. Linton, B. Birren, C. Nusbaum, M. C. Zody, J. Baldwin, K. Devon, K. Dewar, M. Doyle, W. FitzHugh, R. Funke, D. Gage, K. Harris, A. Heaford, J. Howland, L. Kann, J. Lehoczky, R. LeVine, P. McEwan, K. McKernan, J. Meldrim, J. P. Mesirov, C. Miranda, W. Morris, J. Naylor, C. Raymond, M. Rosetti, R. Santos, A. Sheridan, C. Sougnez, Y. Stange-Thomann, N. Stojanovic, A. Subramanian, D. Wyman, J. Rogers, J. Sulston, R. Ainscough, S. Beck, D. Bentley, J. Burton, C. Clee, N. Carter, A. Coulson, R. Deadman, P. Deloukas, A. Dunham, I. Dunham, R. Durbin, L. French, D. Grafham, S. Gregory, T. Hubbard, S. Humphray, A. Hunt, M. Jones, C. Lloyd, A. McMurray, L. Matthews, S. Mercer, S. Milne, J. C. Mullikin, A. Mungall, R. Plumb, M. Ross, R. Shownkeen, S. Sims, R. H. Waterston, R. K. Wilson, L. W. Hillier, J. D. McPherson, M. A. Marra, E. R. Mardis, L. A. Fulton, A. T. Chinwalla, K. H. Pepin, W. R. Gish, S. L. Chissoe, M. C. Wendl, K. D. Delehaunty, T. L. Miner, A. Delehaunty, J. B. Kramer, L. L. Cook, R. S. Fulton, D. L. Johnson, P. J. Minx, S. W. Clifton, T. Hawkins, E. Branscomb, P. Predki, P. Richardson, S. Wenning, T. Slezak, N. Doggett, J. F. Cheng, A. Olsen, S. Lucas, C. Elkin, E. Uberbacher, M. Frazier, R. A. Gibbs, D. M. Muzny, S. E. Scherer, J. B. Bouck, E. J. Sodergren, K. C. Worley, C. M. Rives, J. H. Gorrell, M. L. Metzker, S. L. Naylor, R. S. Kucherlapati, D. L. Nelson, G. M. Weinstock, Y. Sakaki, A. Fujiyama, M. Hattori, T. Yada, A. Toyoda, T. Itoh, C. Kawagoe, H. Watanabe, Y. Totoki, T. Taylor, J. Weissenbach, R. Heilig, W. Saurin, F. Artiguenave, P. Brottier, T. Bruls, E. Pelletier, C. Robert, P. Wincker, D. R. Smith, L. Doucette-Stamm, M. Rubenfield, K. Weinstock, H. M. Lee, J. Dubois, A. Rosenthal, M. Platzer, G. Nyakatura, S. Taudien, A. Rump, H. Yang, J. Yu, J. Wang, G. Huang, J. Gu, L. Hood, L. Rowen, A. Madan, S. Qin, R. W. Davis, N. A. Federspiel, A. P. Abola, M. J. Proctor, R. M. Myers, J. Schmutz, M. Dickson, J. Grimwood, D. R. Cox, M. V. Olson, R. Kaul, C. Raymond, N. Shimizu, K. Kawasaki, S. Minoshima, G. A. Evans, M. Athanasiou, R. Schultz, B. A. Roe, F. Chen, H. Pan, J. Ramser, H. Lehrach, R. Reinhardt, W. R. McCombie, M. de la Bastide, N. Dedhia, H. Blöcker, K. Hornischer, G. Nordsiek, R. Agarwala, L. Aravind, J. A. Bailey, A. Bateman, S. Batzoglou, E. Birney, P. Bork, D. G. Brown, C. B. Burge, L. Cerutti, H. C. Chen, D. Church, M. Clamp, R. R. Copley, T. Doerks, S. R. Eddy, E. E. Eichler, T. S. Furey, J. Galagan, J. G. Gilbert, C. Harmon, Y. Hayashizaki, D. Haussler, H. Hermjakob, K. Hokamp, W. Jang, L. S. Johnson, T. A. Jones, S. Kasif, A. Kaspryzk, S. Kennedy, W. J. Kent, P. Kitts, E. V. Koonin, I. Korf, D. Kulp, D. Lancet, T. M. Lowe, A. McLysaght, T. Mikkelsen, J. V. Moran, N. Mulder, V. J. Pollara, C. P. Ponting, G. Schuler, J. Schultz, G. Slater, A. F. Smit, E. Stupka, J. Szustakowki, D. Thierry-Mieg, J. Thierry-Mieg, L. Wagner, J. Wallis, R. Wheeler, A. Williams, Y. I. Wolf, K. H. Wolfe, S. P. Yang, R. F. Yeh, F. Collins, M. S. Guyer, J. Peterson, A. Felsenfeld, K. A. Wetterstrand, A. Patrinos, M. J. Morgan, P. de Jong, J. J. Catanese, K. Osoegawa, H. Shizuya, S. Choi, Y. J. Chen, J. Szustakowki, International Human Genome Sequencing Consortium, Initial sequencing and analysis of the human genome. *Nature*. **409**, 860–921 (2001).

51. P. H. Sudmant, T. Rausch, E. J. Gardner, R. E. Handsaker, A. Abyzov, J. Huddleston, Y. Zhang, K. Ye, G. Jun, M. H.-Y. Fritz, M. K. Konkel, A. Malhotra, A. M. Stütz, X. Shi, F. P. Casale, J. Chen, F. Hormozdiari, G. Dayama, K. Chen, M. Malig, M. J. P. Chaisson, K. Walter, S. Meiers, S. Kashin, E. Garrison, A. Auton, H. Y. K. Lam, X. J. Mu, C. Alkan, D. Antaki, T. Bae, E. Cerveira, P. Chines, Z. Chong, L. Clarke, E. Dal, L. Ding, S. Emery, X. Fan, M. Gujral, F. Kahveci, J. M. Kidd, Y. Kong, E.-W. Lameijer, S. McCarthy, P. Flicek, R. A. Gibbs, G. Marth, C. E. Mason, A. Menelaou, D. M. Muzny, B. J. Nelson, A. Noor, N. F. Parrish, M. Pendleton, A. Quitadamo, B. Raeder, E. E. Schadt, M. Romanovitch, A. Schlattl, R. Sebra, A. A. Shabalin, A. Untergasser, J. A. Walker, M. Wang, F. Yu, C. Zhang, J. Zhang, X. Zheng-Bradley, W. Zhou, T. Zichner, J. Sebat, M. A. Batzer, S. A. McCarroll, 1000 Genomes Project Consortium, R. E. Mills, M. B. Gerstein, A. Bashir, O. Stegle, S. E. Devine, C. Lee, E. E. Eichler, J. O. Korbel, An integrated map of structural variation in 2,504 human genomes. *Nature*. **526**, 75–81 (2015).

52. J. M. Zook, N. F. Hansen, N. D. Olson, L. Chapman, J. C. Mullikin, C. Xiao, S. Sherry, S. Koren, A. M. Phillippy, P. C. Boutros, S. M. E. Sahraeian, V. Huang, A. Rouette, N. Alexander, C. E. Mason, I. Hajirasouliha, C. Ricketts, J. Lee, R. Tearle, I. T. Fiddes, A. M. Barrio, J. Wala, A. Carroll, N. Ghaffari, O. L. Rodriguez, A. Bashir, S. Jackman, J. J. Farrell, A. M. Wenger, C. Alkan, A. Soylev, M. C. Schatz, S. Garg, G. Church, T. Marschall, K. Chen, X. Fan, A. C. English, J. A. Rosenfeld, W. Zhou, R. E. Mills, J. M. Sage, J. R. Davis, M. D. Kaiser, J. S. Oliver, A. P. Catalano, M. J. P. Chaisson, N. Spies, F. J. Sedlazeck, M. Salit, A robust benchmark for detection of germline large deletions and insertions. *Nat. Biotechnol.* (2020), doi:10.1038/s41587-020-0538-8.

53. W. Zhou, S. B. Emery, D. A. Flasch, Y. Wang, K. Y. Kwan, J. M. Kidd, J. V. Moran, R. E. Mills, Identification and characterization of occult human-specific LINE-1 insertions using long-read sequencing technology. *Nucleic Acids Res.* **48**, 1146–1163 (2020).

54. J. M. C. Tubio, Y. Li, Y. S. Ju, I. Martincorena, S. L. Cooke, M. Tojo, G. Gundem, C. P. Pipinikas, J. Zamora, K. Raine, A. Menzies, P. Roman-Garcia, A. Fullam, M. Gerstung, A. Shlien, P. S. Tarpey, E. Papaemmanuil, S. Knappskog, P. Van Loo, M. Ramakrishna, H. R. Davies, J. Marshall, D. C. Wedge, J. W. Teague, A. P. Butler, S. Nik-Zainal, L. Alexandrov, S. Behjati, L. R. Yates, N. Bolli, L. Mudie, C. Hardy, S. Martin, S. McLaren, S. O'Meara, E. Anderson, M. Maddison, S. Gamble, C. Foster, A. Y. Warren, H. Whitaker, D. Brewer, R. Eeles, C. Cooper, D. Neal, A. G. Lynch, T. Visakorpi, W. B. Isaacs, L. V. Veer, C. Caldas, C. Desmedt, C. Sotiriou, S. Aparicio, J. A. Foekens, J. E. Eyfjörd, S. R. Lakhani, G. Thomas, O. Myklebost, P. N. Span, A.-L. Børresen-Dale, A. L. Richardson, M. Van de Vijver, A. Vincent-Salomon, G. G. Van den Eynden, A. M. Flanagan, P. A. Futreal, S. M. Janes, G. S. Bova, M. R. Stratton, U. McDermott, P. J. Campbell, ICGC Breast Cancer

Group, ICGC Bone Cancer Group, ICGC Prostate Cancer Group, Mobile DNA in cancer. Extensive transduction of nonrepetitive DNA mediated by L1 retrotransposition in cancer genomes. *Science*. **345**, 1251343 (2014).

55. B. Rodriguez-Martin, E. G. Alvarez, A. Baez-Ortega, J. Zamora, F. Supek, J. Demeulemeester, M. Santamarina, Y. S. Ju, J. Temes, D. Garcia-Souto, H. Detering, Y. Li, J. Rodriguez-Castro, A. Dueso-Barroso, A. L. Bruzos, S. C. Dentro, M. G. Blanco, G. Contino, D. Ardeljan, M. Tojo, N. D. Roberts, S. Zumalave, P. A. W. Edwards, J. Weischenfeldt, M. Puiggròs, Z. Chong, K. Chen, E. A. Lee, J. A. Wala, K. Raine, A. Butler, S. M. Waszak, F. C. P. Navarro, S. E. Schumacher, J. Monlong, F. Maura, N. Bolli, G. Bourque, M. Gerstein, P. J. Park, D. C. Wedge, R. Beroukhim, D. Torrents, J. O. Korbel, I. Martincorena, R. C. Fitzgerald, P. Van Loo, H. H. Kazazian, K. H. Burns, PCAWG Structural Variation Working Group, P. J. Campbell, J. M. C. Tubio, PCAWG Consortium, Pan-cancer analysis of whole genomes identifies driver rearrangements promoted by LINE-1 retrotransposition. *Nat. Genet.* **52**, 306–319 (2020).

66. ICGC/TCGA Pan-Cancer Analysis of Whole Genomes Consortium, Pan-cancer analysis of

whole genomes. *Nature*. **578**, 82–93 (2020).

81. A. M. Roy-Engel, A.-H. Salem, O. O. Oyeniran, L. Deininger, D. J. Hedges, G. E. Kilroy, M. A. Batzer, P. L. Deininger, Active Alu element "A-tails": size does matter. *Genome Res.* **12**,

1333–1344 (2002).

82. E. A. Bennett, H. Keller, R. E. Mills, S. Schmidt, J. V. Moran, O. Weichenrieder, S. E. Devine, Active Alu retrotransposons in the human genome. *Genome Res.* **18**, 1875–1883 (2008).

83. D. A. Flasch, Á. Macia, L. Sánchez, M. Ljungman, S. R. Heras, J. L. García-Pérez, T. E. Wilson, J. V. Moran, Genome-wide de novo L1 Retrotransposition Connects Endonuclease Activity with Replication. *Cell*. **177**, 837–851.e28 (2019).

84. J. Jurka, Sequence patterns indicate an enzymatic involvement in integration of mammalian retroposons. *Proc. Natl. Acad. Sci. U. S. A.* **94**, 1872–1877 (1997).

85. Q. Feng, J. V. Moran, H. H. Kazazian Jr, J. D. Boeke, Human L1 retrotransposon encodes a conserved endonuclease required for retrotransposition. *Cell*. **87**, 905–916 (1996).

86. M. J. P. Chaisson, A. D. Sanders, X. Zhao, A. Malhotra, D. Porubsky, T. Rausch, E. J. Gardner, O. Rodriguez, L. Guo, R. L. Collins, X. Fan, J. Wen, R. E. Handsaker, S. Fairley, Z. N. Kronenberg, X. Kong, F. Hormozdiari, D. Lee, A. M. Wenger, A. Hastie, D. Antaki, P. Audano, H. Brand, S. Cantsilieris, H. Cao, E. Cerveira, C. Chen, X. Chen, C.-S. Chin, Z. Chong, N. T. Chuang, D. M. Church, L. Clarke, A. Farrell, J. Flores, T. Galeev, G. David, M. Gujral, V. Guryev, W. Haynes-Heaton, J. Korlach, S. Kumar, J. Y. Kwon, J. E. Lee, J. Lee, W.-P. Lee, S. P. Lee, P. Marks, K. Valud-Martinez, S. Meiers, K. M. Munson, F. Navarro, B. J. Nelson, C. Nodzak, A. Noor, S. Kyriazopoulou-Panagiotopoulou, A. Pang, Y. Qiu, G. Rosanio, M. Ryan, A. Stutz, D. C. J. Spierings, A. Ward, A. E. Welsch, M. Xiao, W. Xu, C. Zhang, Q. Zhu, X. Zheng-Bradley, G. Jun, L. Ding, C. . L. Koh, B. Ren, P. Flicek, K. Chen, M. B. Gerstein, P.-Y. Kwok, P. M. Lansdorp, G. Marth, J. Sebat, X. Shi, A. Bashir, K. Ye, S. E. Devine, M. Talkowski, R. E. Mills, T. Marschall, J. Korbel, E. E. Eichler, C. Lee, Multi-platform discovery of haplotype-resolved structural variation in human genomes. *bioRxiv* (2017), p. 193144.

87. L. Shi, Y. Guo, C. Dong, J. Huddleston, H. Yang, X. Han, A. Fu, Q. Li, N. Li, S. Gong, K. E. Lintner, Q. Ding, Z. Wang, J. Hu, D. Wang, F. Wang, L. Wang, G. J. Lyon, Y. Guan, Y. Shen, O. V. Evgrafov, J. A. Knowles, F. Thibaud-Nissen, V. Schneider, C.-Y. Yu, L. Zhou, E. E. Eichler, K.-F. So, K. Wang, Long-read sequencing and de novo assembly of a Chinese genome. *Nat. Commun.* **7**, 12065 (2016).

88. J.-S. Seo, A. Rhie, J. Kim, S. Lee, M.-H. Sohn, C.-U. Kim, A. Hastie, H. Cao, J.-Y. Yun, J. Kim, J. Kuk, G. H. Park, J. Kim, H. Ryu, J. Kim, M. Roh, J. Baek, M. W. Hunkapiller, J. Korlach, J.-Y. Shin, C. Kim, De novo assembly and phasing of a Korean human genome. *Nature*. **538**, 243–247 (2016).

89. B. Bakker, A. Taudt, M. E. Belderbos, D. Porubsky, D. C. J. Spierings, T. V. de Jong, N. Halsema, H. G. Kazemier, K. Hoekstra-Wakker, A. Bradley, E. S. J. M. de Bont, A. van den Berg, V. Guryev, P. M. Lansdorp, M. Colomé-Tatché, F. Foijer, Single-cell sequencing reveals karyotype heterogeneity in murine and human malignancies. *Genome Biol.* **17**, 115 (2016).

90. S. Chen, P. Krusche, E. Dolzhenko, R. M. Sherman, R. Petrovski, F. Schlesinger, M. Kirsche, D. R. Bentley, M. C. Schatz, F. J. Sedlazeck, M. A. Eberle, Paragraph: a

graph-based structural variant genotyper for short-read sequence data. *Genome Biol.* **20**, 291 (2019).

91. J. Ebler, W. E. Clarke, T. Rausch, P. A. Audano, T. Houwaart, J. Korbel, E. E. Eichler, M. C. Zody, A. T. Dilthey, T. Marschall, Pangenome-based genome inference. *Cold Spring Harbor Laboratory* (2020), p. 2020.11.11.378133.

92. F. Krueger, Trim Galore: a wrapper tool around Cutadapt and FastQC to consistently apply quality and adapter trimming to FastQ files, with some extra functionality for MspI-digested RRBS-type (Reduced Representation Bisufite-Seq) libraries. *URL http://www. bioinformatics. babraham. ac. uk/projects/trim_galore/. (Date of access: 28/04/2016)* (2012).

93. S. Andrews, Others, FastQC: a quality control tool for high throughput sequence data (2010).

101. B. A. Mirauta, D. D. Seaton, D. Bensaddek, A. Brenes, M. J. Bonder, H. Kilpinen, HipSci Consortium, C. A. Agu, A. Alderton, P. Danecek, R. Denton, R. Durbin, D. J. Gaffney, A. Goncalves, R. Halai, S. Harper, C. M. Kirton, A. Kolb-Kokocinski, A. Leha, S. A. McCarthy, Y. Memari, M. Patel, E. Birney, F. P. Casale, L. Clarke, P. W. Harrison, H. Kilpinen, I. Streeter, D. Denovi, O. Stegle, A. I. Lamond, R. Meleckyte, N. Moens, F. M. Watt, W. H. Ouwehand, P. Beales, O. Stegle, A. I. Lamond, Population-scale proteome variation in

human induced pluripotent stem cells. *Elife*. **9** (2020), doi:10.7554/eLife.57390.

102. S. Purcell, B. Neale, K. Todd-Brown, L. Thomas, M. A. R. Ferreira, D. Bender, J. Maller, P. Sklar, P. I. W. de Bakker, M. J. Daly, P. C. Sham, PLINK: a tool set for whole-genome association and population-based linkage analyses. *Am. J. Hum. Genet.* **81**, 559–575 (2007).

103. H. Ongen, A. Buil, A. A. Brown, E. T. Dermitzakis, O. Delaneau, Fast and efficient QTL mapper for thousands of molecular phenotypes. *Bioinformatics*. **32**, 1479–1485 (2016).

104. R. A. Mathias, M. A. Taub, C. R. Gignoux, W. Fu, S. Musharoff, T. D. O'Connor, C. Vergara, D. G. Torgerson, M. Pino-Yanes, S. S. Shringarpure, L. Huang, N. Rafaels, M. P. Boorgula, H. R. Johnston, V. E. Ortega, A. M. Levin, W. Song, R. Torres, B. Padhukasahasram, C. Eng, D.-A. Mejia-Mejia, T. Ferguson, Z. S. Qin, A. F. Scott, M. Yazdanbakhsh, J. G. Wilson, J. Marrugo, L. A. Lange, R. Kumar, P. C. Avila, L. K. Williams, H. Watson, L. B. Ware, C. Olopade, O. Olopade, R. Oliveira, C. Ober, D. L. Nicolae, D. Meyers, A. Mayorga, J. Knight-Madden, T. Hartert, N. N. Hansel, M. G. Foreman, J. G. Ford, M. U. Faruque, G. M. Dunston, L. Caraballo, E. G. Burchard, E. Bleecker, M. I. Araujo, E. F. Herrera-Paz, K. Gietzen, W. E. Grus, M. Bamshad, C. D. Bustamante, E. E. Kenny, R. D. Hernandez, T. H. Beaty, I. Ruczinski, J. Akey, CAAPA, K. C. Barnes, A continuum of admixture in the Western Hemisphere revealed by the African Diaspora genome. *Nat. Commun.* **7**, 12522 (2016).

110. J. O'Connell, D. Gurdasani, O. Delaneau, N. Pirastu, S. Ulivi, M. Cocca, M. Traglia, J. Huang, J. E. Huffman, I. Rudan, R. McQuillan, R. M. Fraser, H. Campbell, O. Polasek, G. Asiki, K. Ekoru, C. Hayward, A. F. Wright, V. Vitart, P. Navarro, J.-F. Zagury, J. F. Wilson, D. Toniolo, P. Gasparini, N. Soranzo, M. S. Sandhu, J. Marchini, A general approach for haplotype phasing across the full spectrum of relatedness. *PLoS Genet.* **10**, e1004234

(2014).

111. B. N. Howie, P. Donnelly, J. Marchini, A flexible and accurate genotype imputation method for the next generation of genome-wide association studies. *PLoS Genet.* **5**, e1000529 (2009).

112. B. K. Maples, S. Gravel, E. E. Kenny, C. D. Bustamante, RFMix: a discriminative modeling approach for rapid and robust local-ancestry inference. *Am. J. Hum. Genet.* **93**, 278–288 (2013).

113. D. H. Alexander, J. Novembre, K. Lange, Fast model-based estimation of ancestry in unrelated individuals. *Genome Res.* **19**, 1655–1664 (2009).

114. S. Mallick, H. Li, M. Lipson, I. Mathieson, M. Gymrek, F. Racimo, M. Zhao, N. Chennagiri, S. Nordenfelt, A. Tandon, P. Skoglund, I. Lazaridis, S. Sankararaman, Q. Fu, N. Rohland, G. Renaud, Y. Erlich, T. Willems, C. Gallo, J. P. Spence, Y. S. Song, G. Poletti, F. Balloux, G. van Driem, P. de Knijff, I. G. Romero, A. R. Jha, D. M. Behar, C. M. Bravi, C. Capelli, T. Hervig, A. Moreno-Estrada, O. L. Posukh, E. Balanovska, O. Balanovsky, S. Karachanak-Yankova, H. Sahakyan, D. Toncheva, L. Yepiskoposyan, C. Tyler-Smith, Y. Xue, M. S. Abdullah, A. Ruiz-Linares, C. M. Beall, A. Di Rienzo, C. Jeong, E. B. Starikovskaya, E. Metspalu, J. Parik, R. Villems, B. M. Henn, U. Hodoglugil, R. Mahley, A. Sajantila, G. Stamatoyannopoulos, J. T. S. Wee, R. Khusainova, E. Khusnutdinova, S. Litvinov, G. Ayodo, D. Comas, M. F. Hammer, T. Kivisild, W. Klitz, C. A. Winkler, D. Labuda, M. Bamshad, L. B. Jorde, S. A. Tishkoff, W. S. Watkins, M. Metspalu, S. Dryomov, R. Sukernik, L. Singh, K. Thangaraj, S. Pääbo, J. Kelso, N. Patterson, D. Reich, The Simons Genome Diversity Project: 300 genomes from 142 diverse populations. *Nature*. **538**, 201–206 (2016).

115. A. R. Martin, C. R. Gignoux, R. K. Walters, G. L. Wojcik, B. M. Neale, S. Gravel, M. J. Daly, C. D. Bustamante, E. E. Kenny, Human Demographic History Impacts Genetic Risk Prediction across Diverse Populations. *Am. J. Hum. Genet.* **107**, 788–789 (2020).

116. L. Speidel, M. Forest, S. Shi, S. R. Myers, A method for genome-wide genealogy estimation for thousands of samples. *Nat. Genet.* **51**, 1321–1329 (2019).

117. D. Gordon, J. Huddleston, M. J. P. Chaisson, C. M. Hill, Z. N. Kronenberg, K. M. Munson, M. Malig, A. Raja, I. Fiddes, L. W. Hillier, C. Dunn, C. Baker, J. Armstrong, M. Diekhans, B. Paten, J. Shendure, R. K. Wilson, D. Haussler, C.-S. Chin, E. E. Eichler, Long-read sequence assembly of the gorilla genome. *Science*. **352**, aae0344–aae0344 (2016).

118. Z. N. Kronenberg, I. T. Fiddes, D. Gordon, S. Murali, S. Cantsilieris, O. S. Meyerson, J. G. Underwood, B. J. Nelson, M. J. P. Chaisson, M. L. Dougherty, K. M. Munson, A. R. Hastie, M. Diekhans, F. Hormozdiari, N. Lorusso, K. Hoekzema, R. Qiu, K. Clark, A. Raja, A. M. E. Welch, M. Sorensen, C. Baker, R. S. Fulton, J. Armstrong, T. A. Graves-Lindsay, A. M. Denli, E. R. Hoppe, P. H. Hsieh, C. M. Hill, A. W. C. Pang, J. Lee, E. T. Lam, S. K. Dutcher, F. H. Gage, W. C. Warren, J. Shendure, D. Haussler, V. A. Schneider, H. Cao, M. Ventura, R. K. Wilson, B. Paten, A. Pollen, E. E. Eichler, High-resolution comparative analysis of great ape genomes. *Science*. **360** (2018), doi:10.1126/science.aar6343.

119. X. Yi, Y. Liang, E. Huerta-Sanchez, X. Jin, Z. X. P. Cuo, J. E. Pool, X. Xu, H. Jiang, N. Vinckenbosch, T. S. Korneliussen, H. Zheng, T. Liu, W. He, K. Li, R. Luo, X. Nie, H. Wu, M. Zhao, H. Cao, J. Zou, Y. Shan, S. Li, Q. Yang, Asan, P. Ni, G. Tian, J. Xu, X. Liu, T. Jiang, R. Wu, G. Zhou, M. Tang, J. Qin, T. Wang, S. Feng, G. Li, Huasang, J. Luosang, W. Wang, F. Chen, Y. Wang, X. Zheng, Z. Li, Z. Bianba, G. Yang, X. Wang, S. Tang, G. Gao, Y. Chen, Z. Luo, L. Gusang, Z. Cao, Q. Zhang, W. Ouyang, X. Ren, H. Liang, H. Zheng, Y. Huang, J. Li, L. Bolund, K. Kristiansen, Y. Li, Y. Zhang, X. Zhang, R. Li, S. Li, H. Yang, R. Nielsen, J. Wang, J. Wang, Sequencing of 50 human exomes reveals adaptation to high altitude. *Science*. **329**, 75–78 (2010).

120. ENCODE Project Consortium, J. E. Moore, M. J. Purcaro, H. E. Pratt, C. B. Epstein, N. Shoresh, J. Adrian, T. Kawli, C. A. Davis, A. Dobin, R. Kaul, J. Halow, E. L. Van Nostrand, P. Freese, D. U. Gorkin, Y. Shen, Y. He, M. Mackiewicz, F. Pauli-Behn, B. A. Williams, A. Mortazavi, C. A. Keller, X.-O. Zhang, S. I. Elhajjajy, J. Huey, D. E. Dickel, V. Snetkova, X. Wei, X. Wang, J. C. Rivera-Mulia, J. Rozowsky, J. Zhang, S. B. Chhetri, J. Zhang, A. Victorsen, K. P. White, A. Visel, G. W. Yeo, C. B. Burge, E. Lécuyer, D. M. Gilbert, J. Dekker, J. Rinn, E. M. Mendenhall, J. R. Ecker, M. Kellis, R. J. Klein, W. S. Noble, A. Kundaje, R. Guigó, P. J. Farnham, J. M. Cherry, R. M. Myers, B. Ren, B. R. Graveley, M. B. Gerstein, L. A. Pennacchio, M. P. Snyder, B. E. Bernstein, B. Wold, R. C. Hardison, T. R. Gingeras, J. A. Stamatoyannopoulos, Z. Weng, Expanded encyclopaedias of DNA elements in the human and mouse genomes. *Nature*. **583**, 699–710 (2020).

121. G. Benson, Tandem repeats finder: a program to analyze DNA sequences. *Nucleic Acids Res.* **27**, 573–580 (1999).
