## Supplemental Figures for "*De novo* assembly of 64 haplotype-resolved human genomes of diverse ancestry and integrated analysis of structural variation"

**Table of Contents**


Fig. S1. Data flowchart **6**

Fig. S2. Summary of Strand-seq libraries used in this study **7**

Fig. S3. Haploid assembly contig N50 as a function of library insert size **8**

Fig. S4. Phasing accuracy for child HG00514 (HiFi) **9**

Fig. S5. Phasing accuracy for child HG00514 (CLR) **9**

Fig. S6. Haploid assembly contig coverage relative to GRCh38 **10**

Fig. S7. Haploid assembly contig coverage in GRCh38 centromeres **11**

Fig. S8. Misassemblies detected via Bionano hybrid scaffolding **11**

Fig. S9. Alignments of unscaffolded contig sequence **12**

Fig. S10. Heterozygous SNVs per sample **13**

Fig. S11. Haploid assembly k-mer counts **14**

Fig. S12. Haploid assembly contig coverage in GRCh38 issue regions **15**

Fig. S13. Ensembl Regulatory Build regions in haploid assemblies **16**

Fig. S14. Phased assembly contig coverage **17**

Fig. S15. Variant merging strategy **18**

Fig. S16. PAV alignment trimming **19**

Fig. S17. PAV flags inversion sites **20**

Fig. S18. PAV flags inversion sites **21**

Fig. S19. DeBreak variant discovery **22**

Fig. S20. Overview of GATK-SV pipeline **23**

Fig S21.Overview of the short-read Illumina integration callset on the 34 genomes with matched PacBio sequences **24**

Fig S22. Length distribution of SVs from HGSV, gnomAD and CCDG **25**

Fig S23. Distribution of normalized sequencing depth of CNVs **28**

Fig. S24. Ideogram showing Bionano calls (≥5 kbp) and clusters **29**

Fig. S25. Bionano and PAV intersection for SVs greater than 5 kbp **30**

Fig. S26. Population distribution of SVs identified on chromosome 1 **31**

Fig. S27. Population distribution of SVs identified on chromosome 5 (21.1–21.7 Mbp) **32**

Fig. S28. Full configuration of 3q29 region (195.4–196.1 Mbp) **33**

Fig. S29. Composite files summary (n=32) **34**

Fig. S30. Inversion callset summary (n=316) **35**

Fig. S31. Inversions flanked by SDs **36**

Fig. S32. Phased assembly alignments in 16p12 region **37**

Fig. S33. Genetic background of inverted haplotypes at the 16p12 chromosome region **39**

Fig. S34. UpSet plots for the integrated callset and PAV annotation in all HGSVC2 samples **40**

Fig. S35. Collection of sequence-resolved full-length L1s **41**

Fig. S36. Complete phylogeny for all active sequence-resolved Full-Length (FL)-L1s **42**

Fig. S37. Catalogue of source SVA active in HGSVC2 dataset **43**

Fig. S38. Exon shuffling by intronic SVA source element **44**

Fig. S39. Multi-transduction event disentangled via assemblies **44**

Fig. S40. Distributions of VNTR length in reference and polymorphic SVAs in each individual discovery sample **45**

Fig. S41. Box plots of poly(A) tract and sequence logos of EN cleavage sites for MEI subfamilies **46**

Fig. S42. New sequence-resolved SVs **46**

Fig. S43. PAV concordance among callers for HG00733 **47**

Fig. S44. PAV concordance among callers for HG00733 outside tandem repeats **47**

Fig. S45. Excess singleton rate is reduced by callset filtering **49**

Fig. S46. SV hotspot detection and enrichment for chromosome ends and SDs **50**

Fig. S47. Raw data supporting detected SV hotspots **52**

Fig. S48. Summary of detected SV hotspots per chromosome **53**

Fig. S49. Evolutionary distances inside and outside of the HLA region **54**

Fig. S50. Count of alternative alleles per haplotype for the HLA region **55**

Fig. S51. Summary of genetic variability over the HLA region **56**

Fig. S52. Definition of distinct HLA haplotypes based on SV clustering **57**

Fig. S53. Summary of HLA haplotypes with a similar SV distribution **58**

Fig. S54. Subseq illustration **58**

Fig. S55. Subseq region lengths deviate with SV length with less variation from HiFi **60**

Fig. S56. SV support from raw reads **61**

Fig. S57. Example of SV calls with and without support **62**

Fig. S58. Fraction of unsupported SVs per genome **63**

Fig. S59. Genomic distribution of unsupported variants detected by HiFi assemblies **65**

Fig. S60. Genomic distribution of unsupported variants detected by CLR assemblies **66**

Fig. S61. Examples of Inspector Quality Control **67**

Fig. S62. Workflow of SV QC **68**

Fig. S63. The overall QC results of HiFi and CLR samples **69**

Fig. S64. Examples of QC labels **70**

Fig. S65. Distribution of insertions and deletions belonging to different mechanism categories **71**

Fig. S66. Distribution of insertions and deletions of different length groups belonging to different mechanism categories **72**

Fig. S67. Distribution of insertions and deletions overlapping with distinct functional elements belonging to different mechanism categories **73**

Fig. S68. Distribution of insertions and deletions belonging to different mechanism categories (based on homology length ≥200 bp) **74**

Fig. S69. Distribution of insertions and deletions of different length groups belonging to different mechanism categories (based on homology length ≥200 bp) **75**

Fig. S70. Distribution of insertions and deletions overlapping with distinct functional elements belonging to different mechanism categories based on homology length ≥200 bp **76**

Fig. S71. Non-reference k-mer density distribution (violin plot) by population for 2,504 unrelated samples **76**

Fig. S72. Comparison of PanGenie and Paragraph allele frequencies on the pilot set **77**

Fig. S73. Number of heterozygous SVs per population **78**

Fig. S74. Allele frequencies of SVs in the lenient callset **79**

Fig. S75. PanGenie allele frequency vs. SV length **79**

Fig. S76. Fst vs. SV length for all superpopulations (deletions) **80**

Fig. S77. Fst vs. SV length for all superpopulations (insertions) **81**

Fig. S78. Optimization of the number of principal components (PCs) to correct for when mapping eQTLs **82**

Fig. S79. Venn diagram of known GWAS associations extracted from the GWAS catalog, UK Biobank and PhenoScanner (p-value < 1E-6) **83**

Fig. S80. Numbers of genotyped variants (in blue) and eQTLs (in orange) of various types that overlap with known GWAS SNPs **84**

Fig. S81. Enrichment of SV eQTLs for overlapping with GWAS SNPs **85**

Fig. S82. Hidden Markov model for local ancestry inference using haplotype-resolved assemblies **86**

Fig. S83. Concordant ancestry calls between HiFi and CLR data for the YRI trio **87**

Fig. S84. Length distributions of inferred genome-wide ancestry tracts for all haplotype-phased assemblies **88**

Fig. S85. Inferred local ancestry blocks across chromosome 1 for haplotype-phased assemblies **89**

Fig. S86. Top population branch statistics (PBS) hits per superpopulation **90**

Fig. S87. Structure of a 4.0 kbp insertion in the first LCT exon **91**

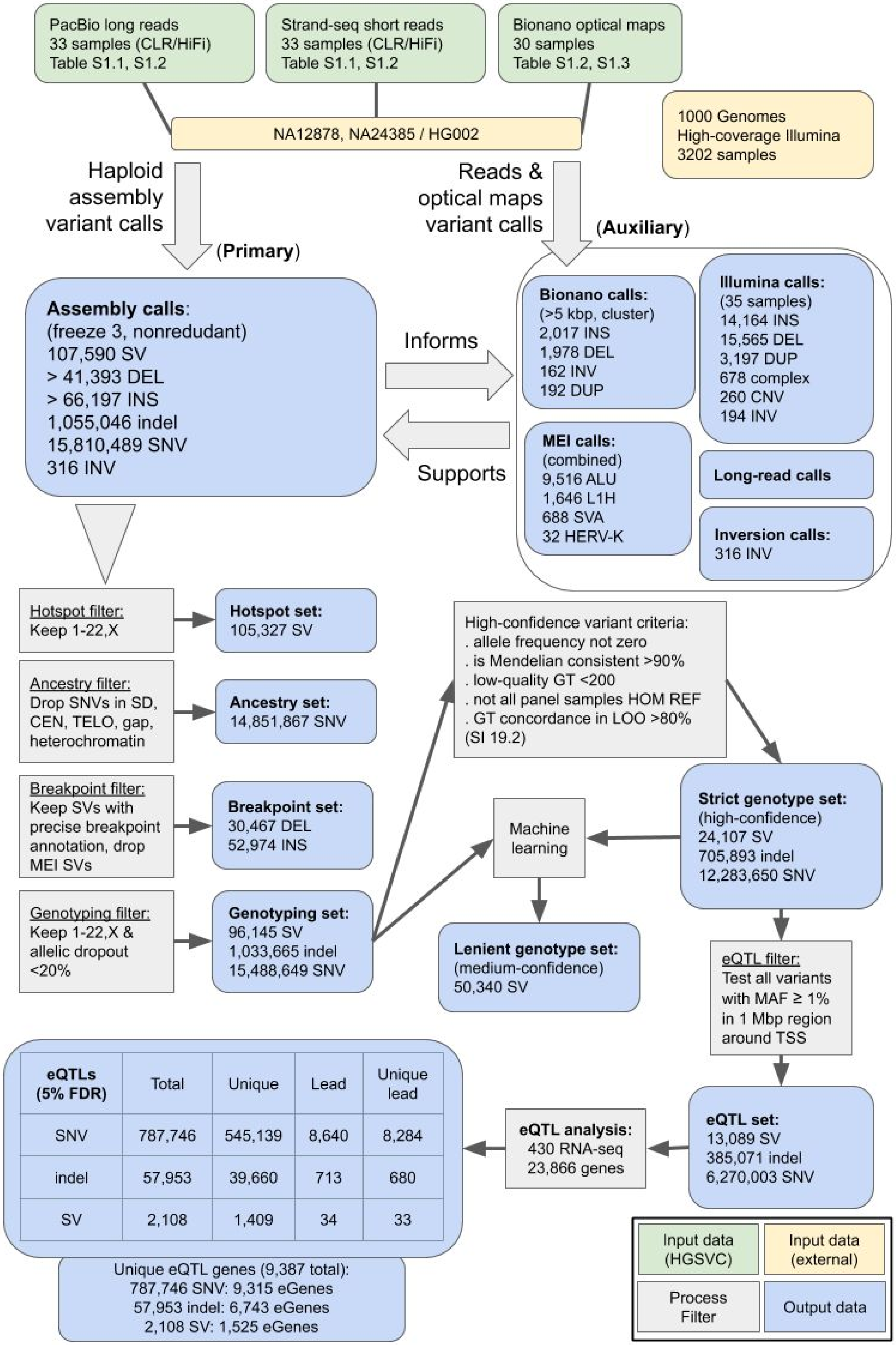


| eQTLs (5% FDR) | Total | Unique | Lead | Unique lead |
|---|---|---|---|---|
| SNV | 787,746 | 545,139 | 8,640 | 8,284 |
| indel | 57,953 | 39,660 | 713 | 680 |
| SV | 2,108 | 1,409 | 34 | 33 |

**Fig. S1. Data flowchart**

Schematic of the data flow in our study focusing on variant call sets (blue boxes) and necessary processing and filtering steps (gray boxes) to derive reduced callsets for the various analyses downstream of the haploid assembly variant calling (blue box “Assembly calls”). Green boxes: input datasets created as part of this study; yellow boxes: external datasets.

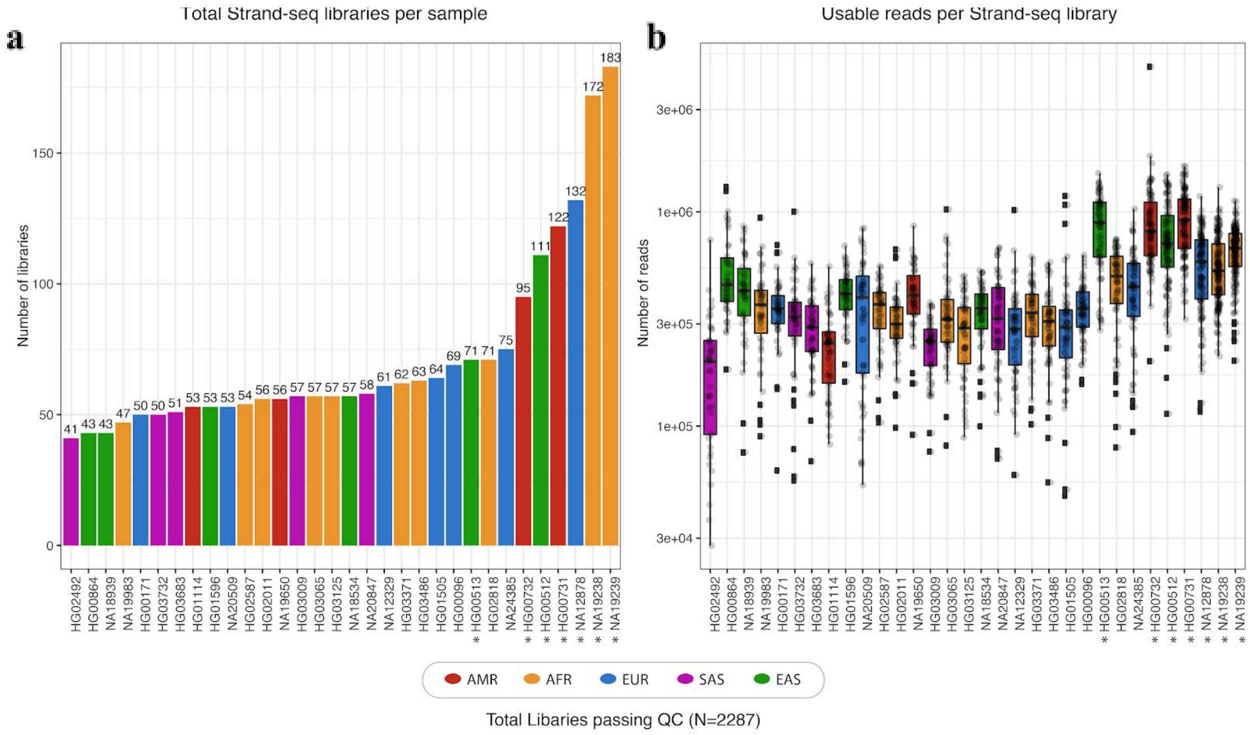


**Fig. S2. Summary of Strand-seq libraries used in this study**

**a**) Bars indicate the total number of high-quality Strand-seq libraries (N=2,287) selected for analysis, per sample. **b**) Box plots show the range of usable sequencing fragments per high-quality library. Graphs are colored by superpopulation; asterisks mark published datasets included in this analysis.

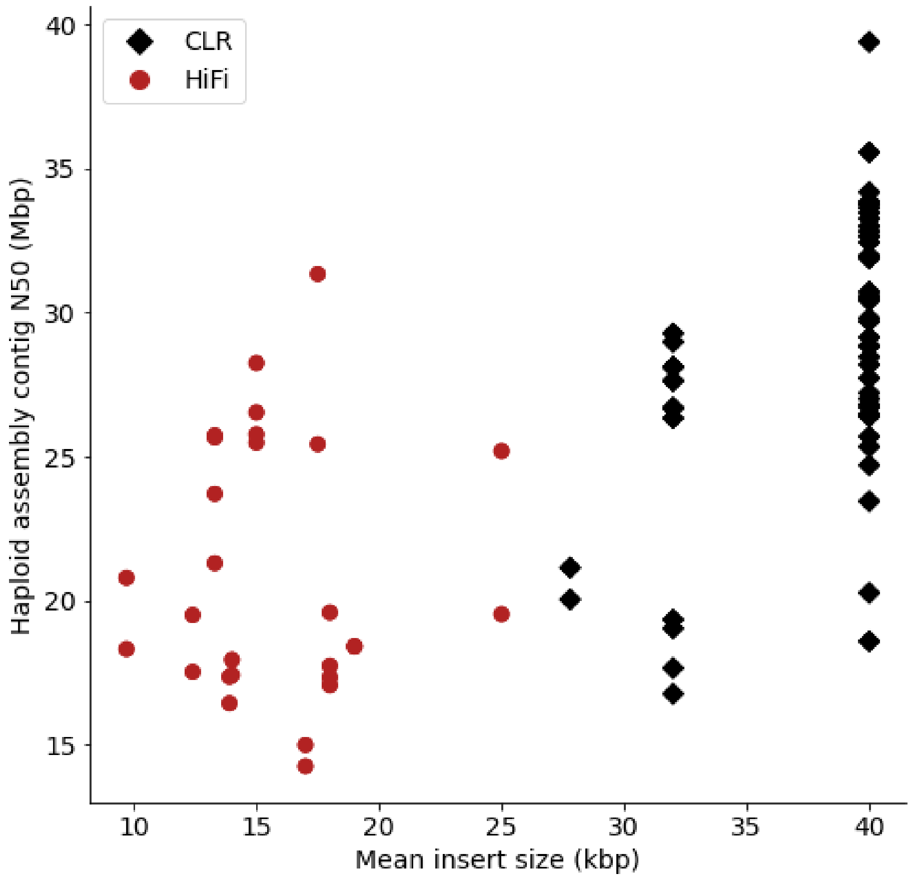


**Fig. S3. Haploid assembly contig N50 as a function of library insert size**

Relation between long-read sequencing library insert size averaged over all SMRT cells per sample (x-axis) and haploid assembly contig N50 (y-axis) plotted for CLR (black diamonds) and HiFi (red circles) samples.

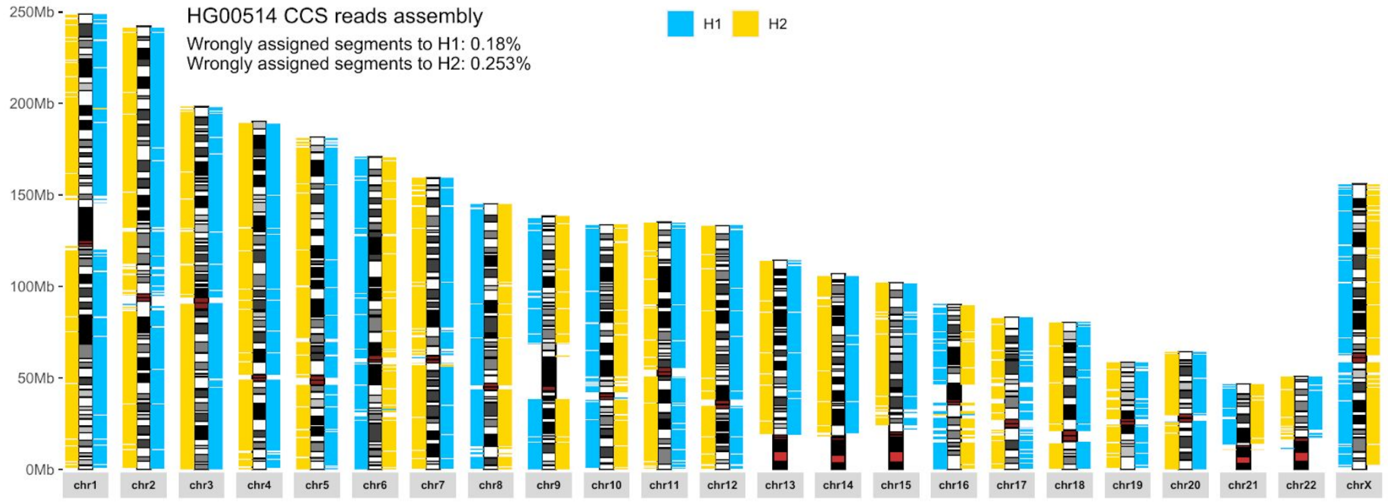


**Fig. S4. Phasing accuracy for child HG00514 (HiFi)**

The phased HiFi assembly for sample HG00514 is divided into 1 Mbp blocks sequence, which are aligned to GRCh38 and colored by parental haplotype (H1 blue, H2 yellow) based on a trio-phased set of reference SNVs for this individual. The fraction of blocks assigned to the wrong haplotype is indicated at the top.

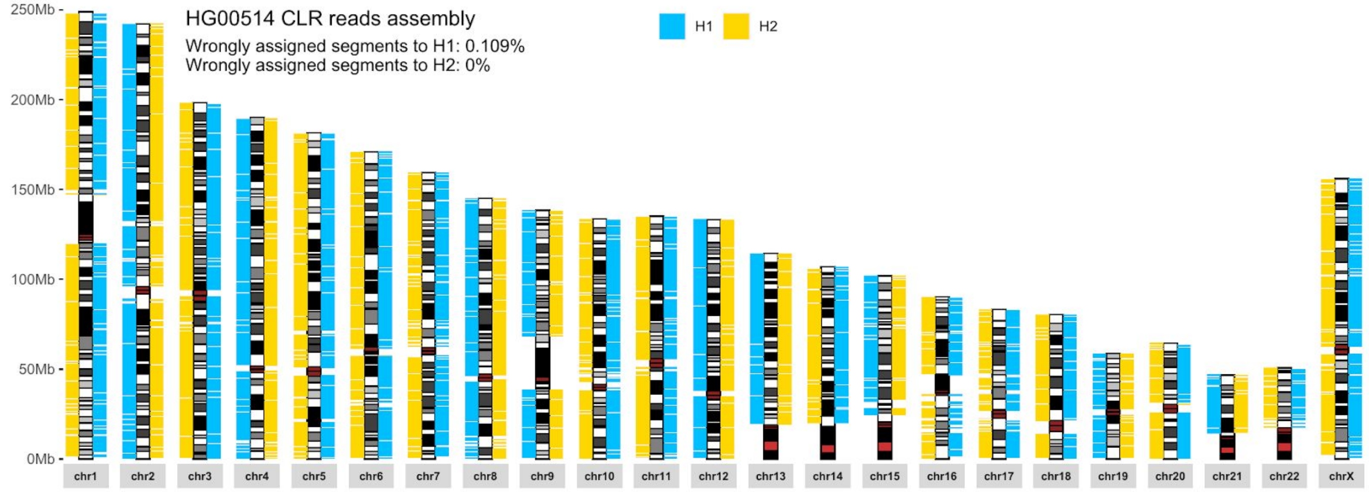


**Fig. S5. Phasing accuracy for child HG00514 (CLR)**

The phased CLR assembly for sample HG00514 is divided into 1 Mbp blocks, which are aligned to GRCh38 and colored by parental haplotype (H1 blue, H2 yellow) based on a trio-phased set of reference SNVs for this individual. The fraction of blocks assigned to the wrong haplotype is indicated at the top.

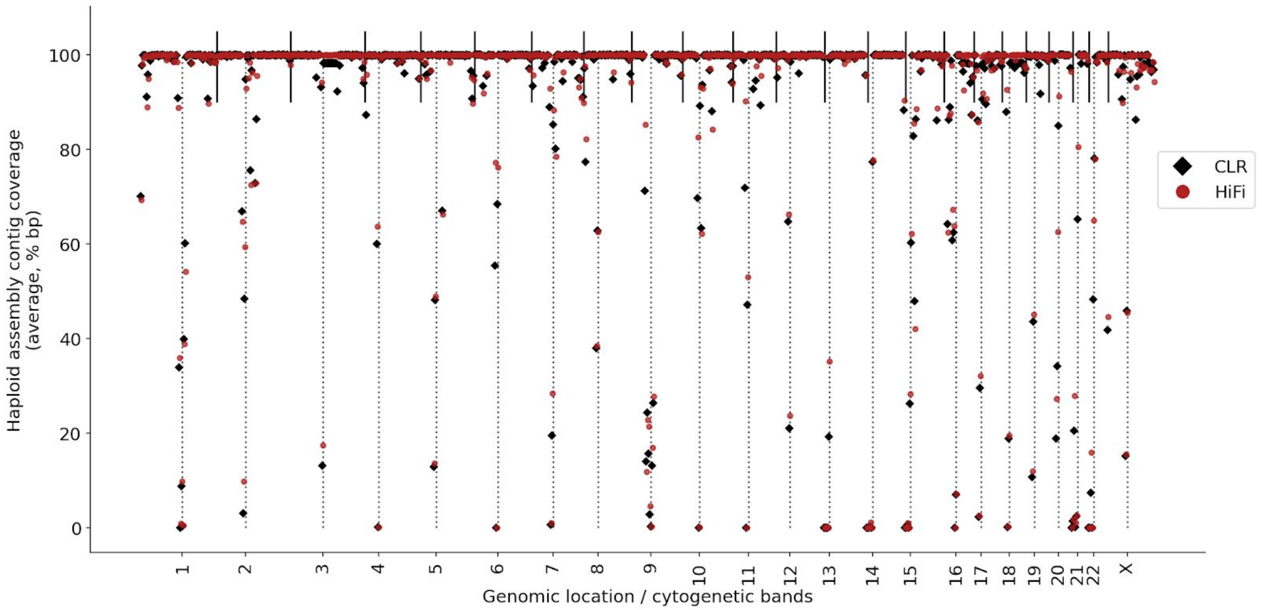


**Fig. S6. Haploid assembly contig coverage relative to GRCh38**

Cytogenetic bands are plotted with fixed width along the x-axis for chromosomes 1-22 and X. The contig coverage (y-axis, % covered bp in region) is depicted as an average over all CLR (black diamonds) and HiFi (red circles) haplotype assemblies. Centromere locations are indicated as gray dotted lines, and chromosome boundaries as black vertical lines at the top.

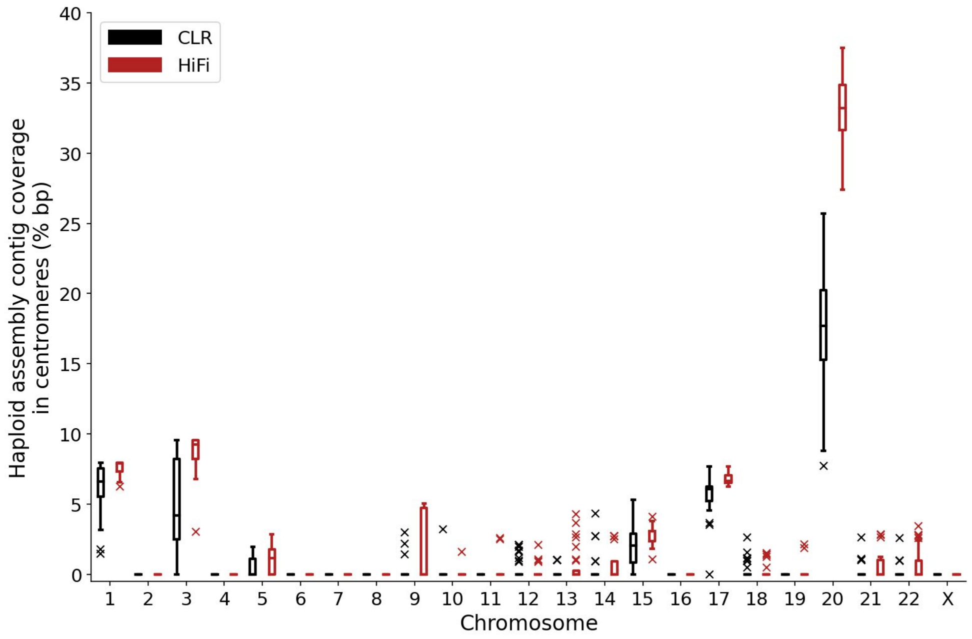

**Fig. S7. Haploid assembly contig coverage in GRCh38 centromeres**

Variation in haploid assembly contig coverage in centromeres for chromosomes 1-22 and X (x-axis) is depicted separately for CLR (black) and HiFi (red) assemblies. The symbol “x” marks outliers outside of 1.5 X interquartile range.

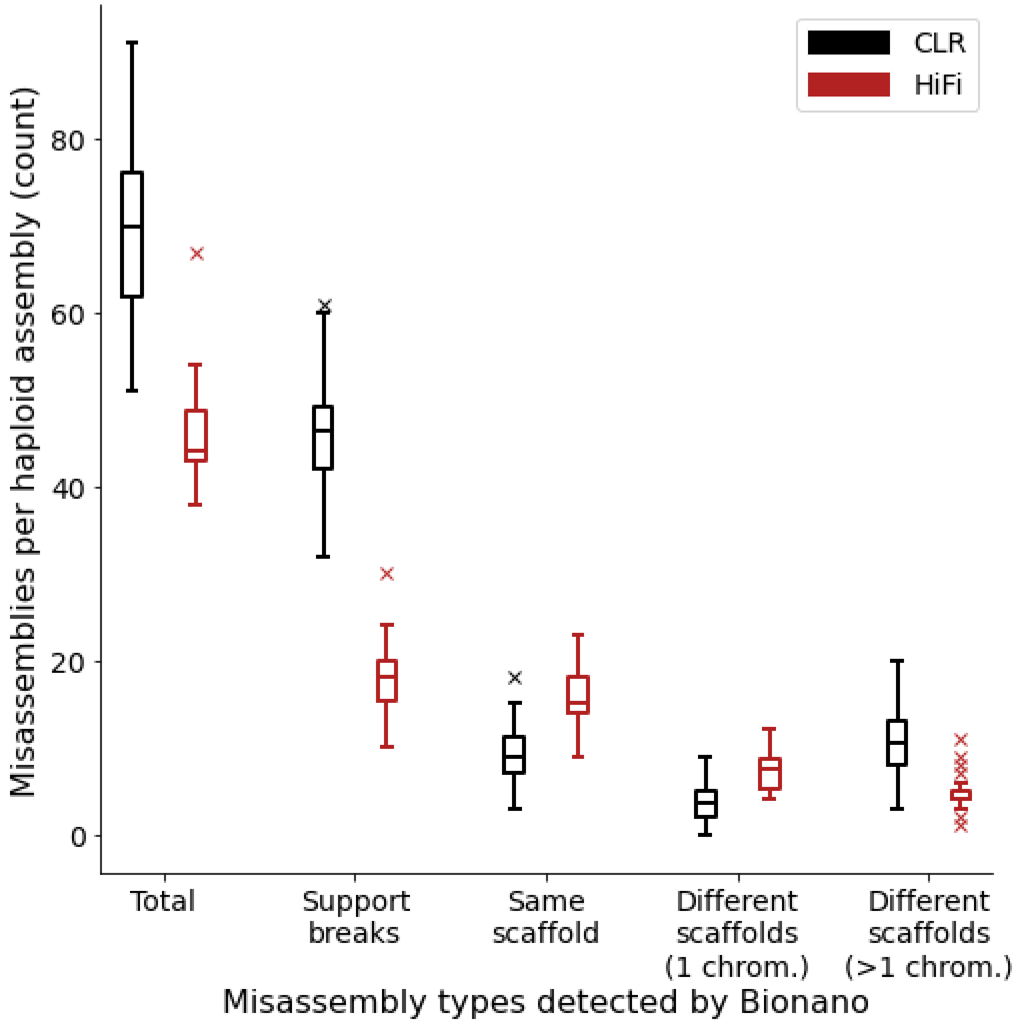


**Fig. S8. Misassemblies detected via Bionano hybrid scaffolding**

Contig-level phased assemblies were combined with Bionano optical maps to create scaffolded hybrid assemblies. Misassemblies detected per haploid assembly (x-axis, “Total”) were characterized based on the type of misassembly (x-axis, left to right). Part of contig has no Bionano support; contig is fragmented within scaffold; contig is fragmented between different scaffolds on the same chromosome; contig is fragmented between different scaffolds on different chromosomes (“chimerism”).

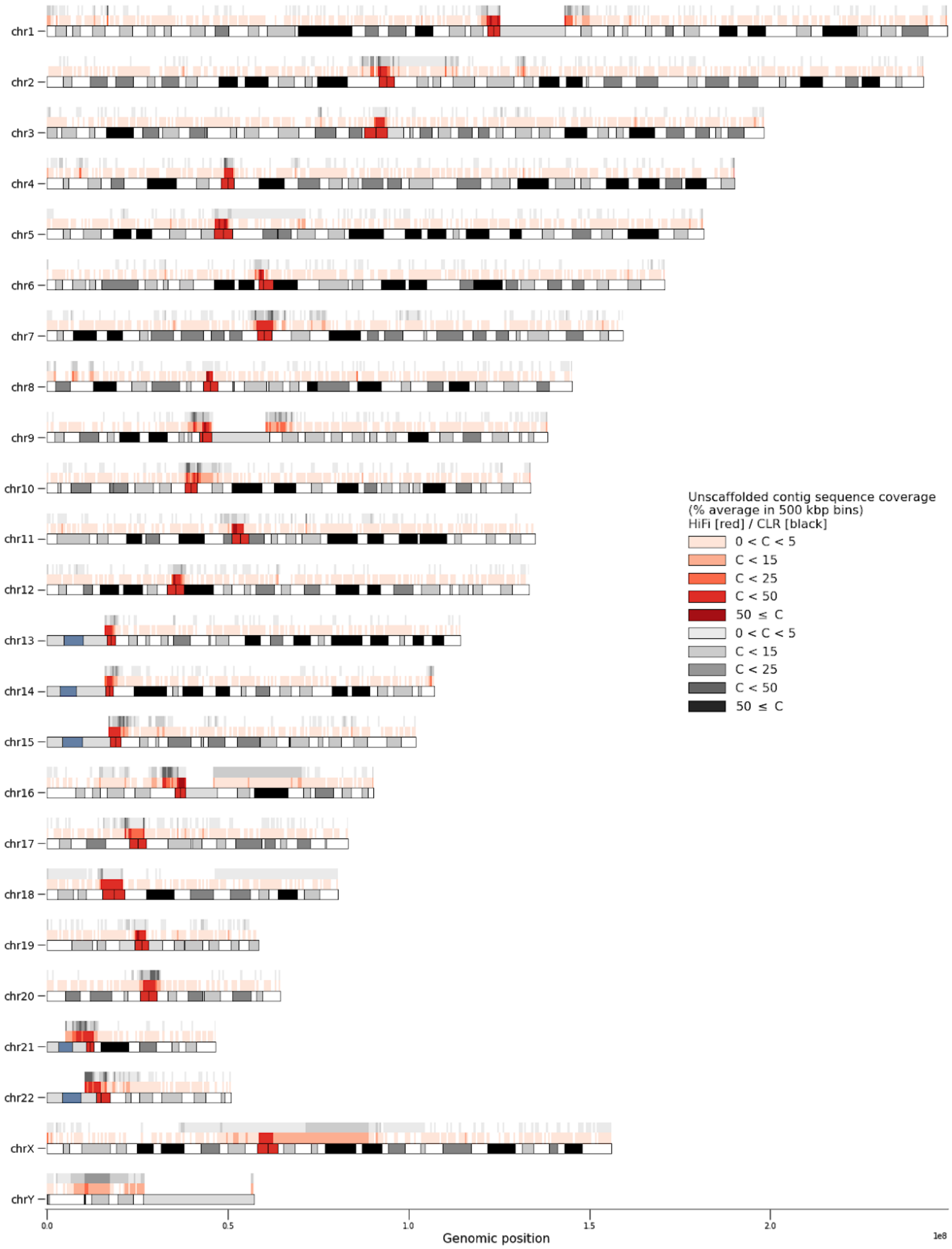


**Fig. S9. Alignments of unscaffolded contig sequence**

Unscaffolded contig sequence was split into 500 bp reads and aligned to GRCh38. Read alignment coverage was aggregated in bins of 500 kbp and are plotted as an average for all HiFi (red track) and all CLR (black track) assemblies.

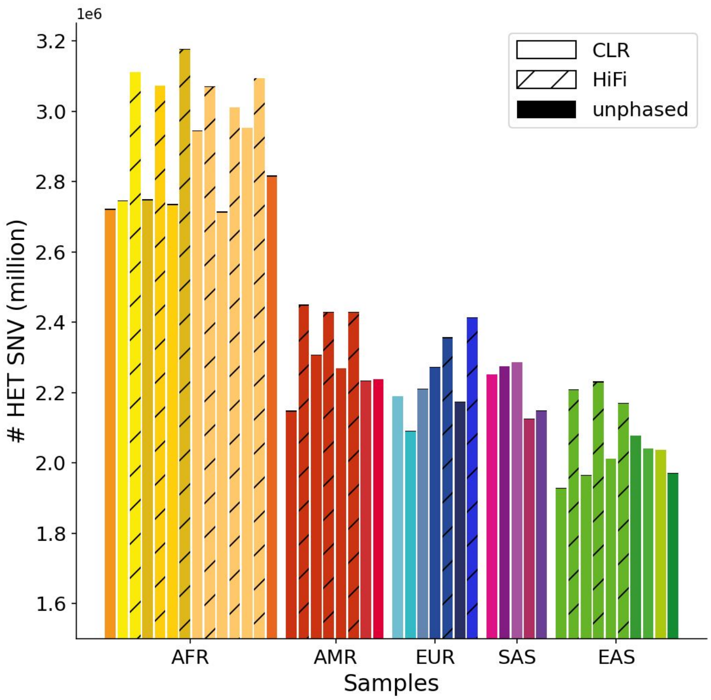


**Fig. S10. Heterozygous SNVs per sample**

Samples are grouped by superpopulation along the x-axis (for color code, see Figure 1f). Bar height (y-axis, in millions) depicts the number of heterozygous SNVs per sample, as called in the diploid assembly pipeline to obtain local phase information. The unphased fraction of variants is indicated as the top black part of each bar. HiFi samples are represented by hatched bars.

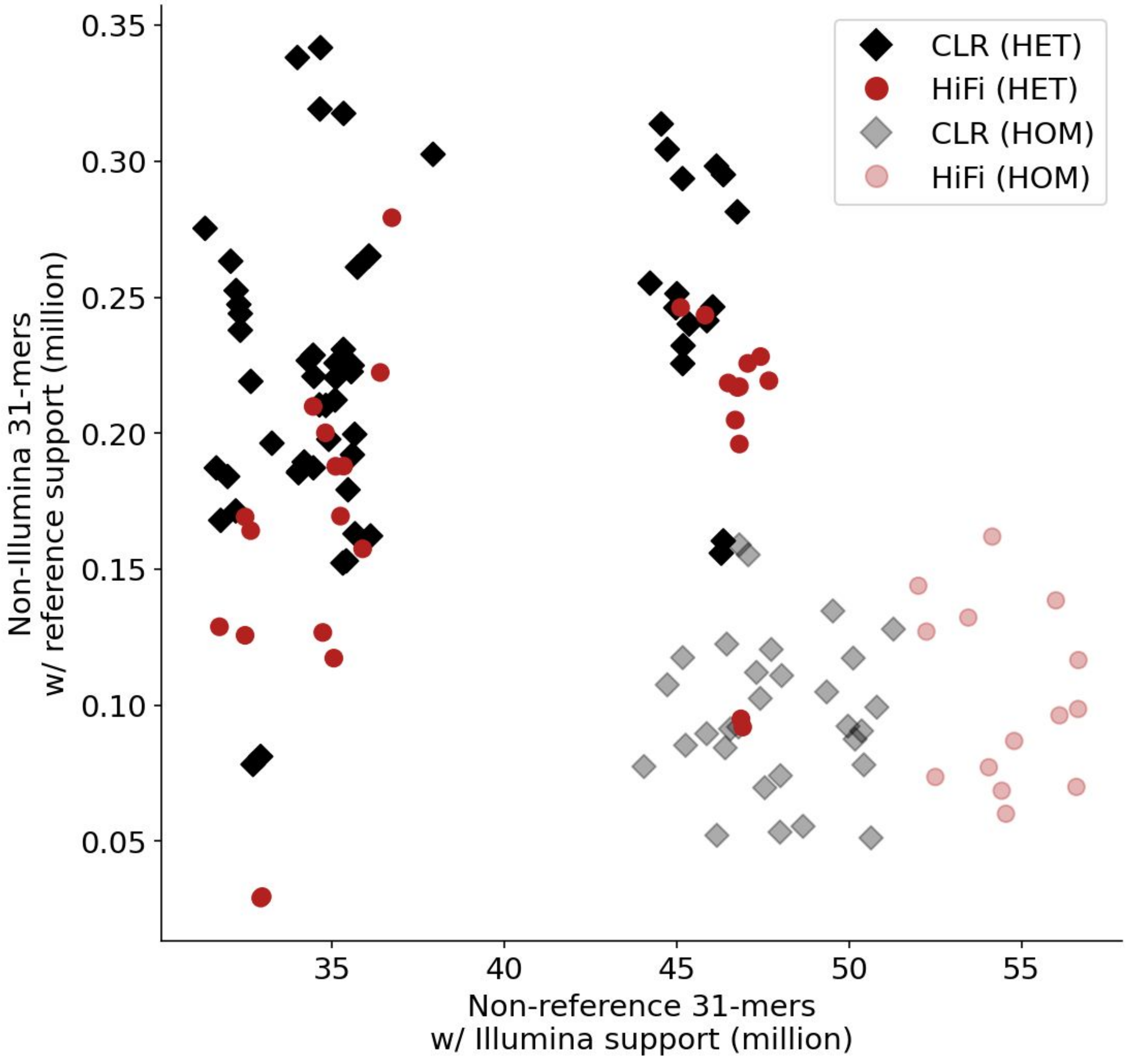


**Fig. S11. Haploid assembly k-mer counts**

Individual haplotype assemblies are depicted as black diamonds (CLR) or red circles (HiFi). 31-mer counts supported by Illumina data but not found in the GRCh38 reference are plotted on the x-axis (in million), and 31-mer counts supported by the GRCh38 reference and not found in Illumina data on the y-axis. The fraction of homozygous k-mers found in both assembled haplotypes is depicted as semi-transparent points.

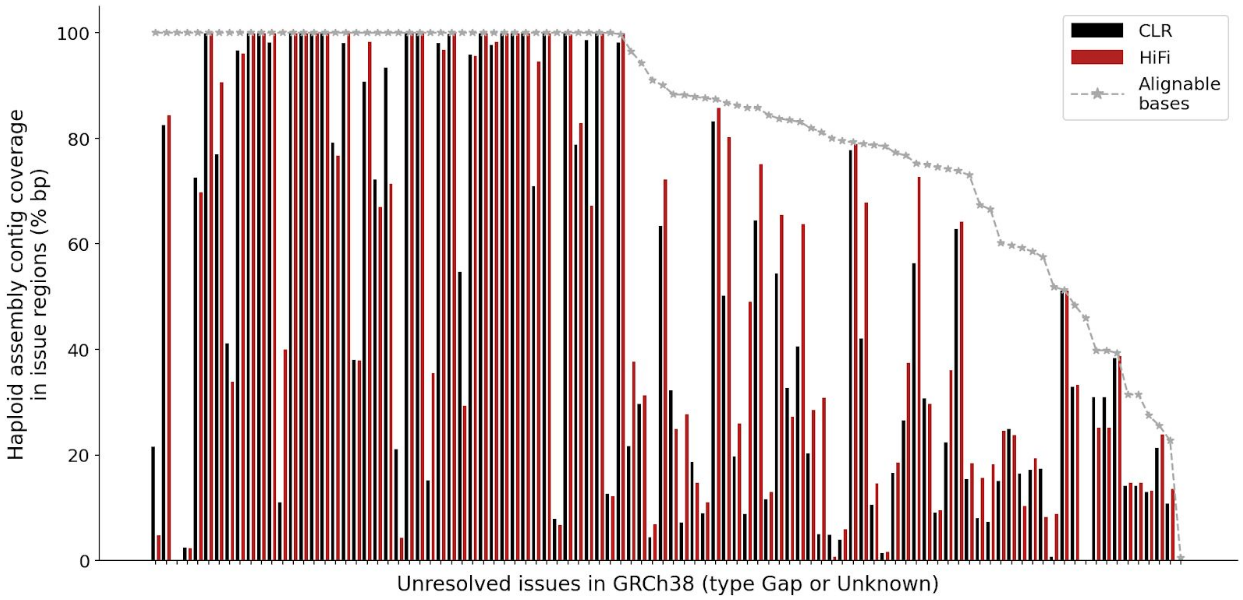


**Fig. S12. Haploid assembly contig coverage in GRCh38 issue regions**

Unresolved issues in GRCh38 of type “Gap” or “Unknown” are arranged in descending order by percent alignable bases (ACGT, but not N) on the x-axis. The maximal attainable contig coverage per issue region is depicted with the gray star line at the top. Bar height indicates average haploid contig coverage (y-axis, % covered bp in region) for all CLR (black) and HiFi (red) assemblies.

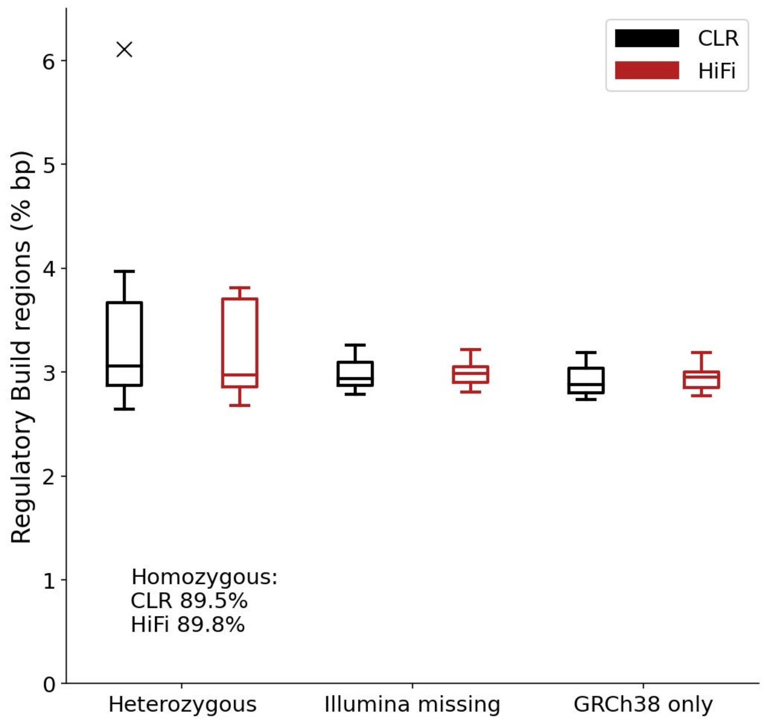


**Fig. S13. Ensembl Regulatory Build regions in haploid assemblies**

Regulatory regions are classified for each phased assembly as homozygous (CLR/HiFi average, text only for layout reasons), heterozygous (left), missing in Illumina short reads but present in at least one haplotype (middle), or only detected in the GRCh38 reference (right). Box plots illustrate the amount of detectable regulatory regions as percent-annotated base pairs over all Regulatory Build categories and separately for CLR (black) and HiFi (red) assemblies.

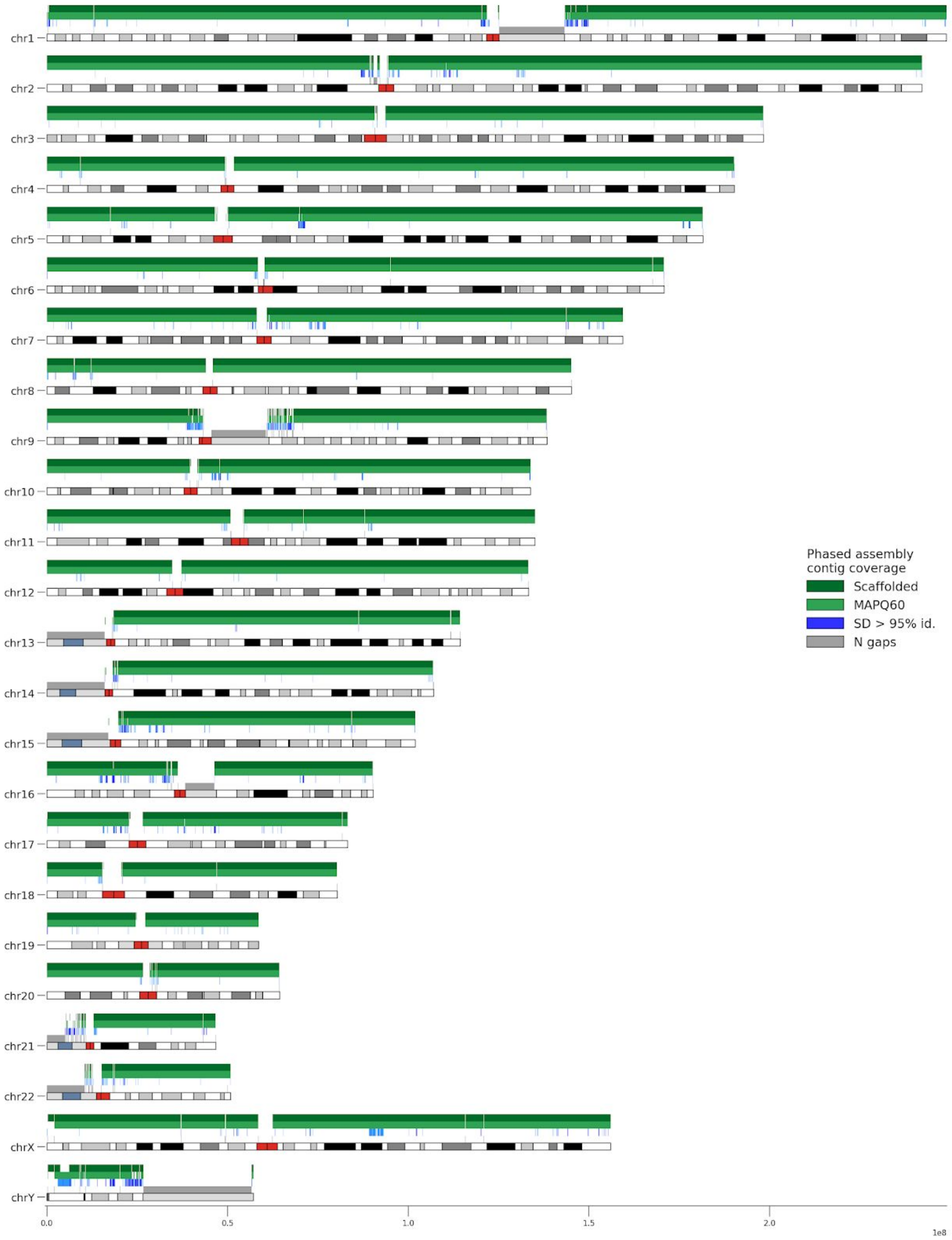


**Fig. S14. Phased assembly contig coverage**

GRCh38 regions covered with phased assembly contig alignments were defined based on Bionano hybrid scaffolded assemblies (dark green), or based on MAPQ60 threshold alignments (green). Segmental duplications (SDs) with >95% identity are shown in shades of blue (steps >98% and >99% identity), and N gaps in the GRCh38 reference are indicated in gray.

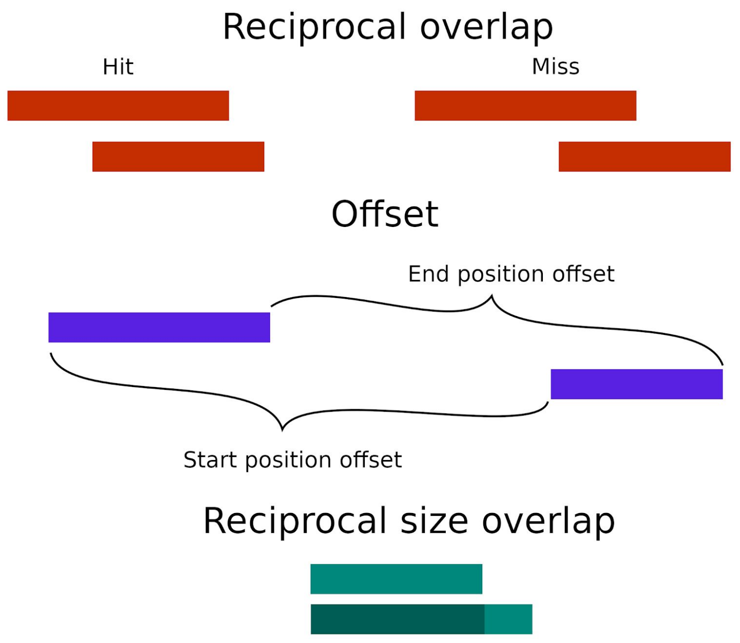


**Fig. S15. Variant merging strategy**

50% reciprocal overlap (red) often misses similar-sized events if they are shifted by a small distance, especially for smaller events. By considering a distance-size overlap, smaller variants within a given proximity (blue) and size overlap (green) can be intersected.

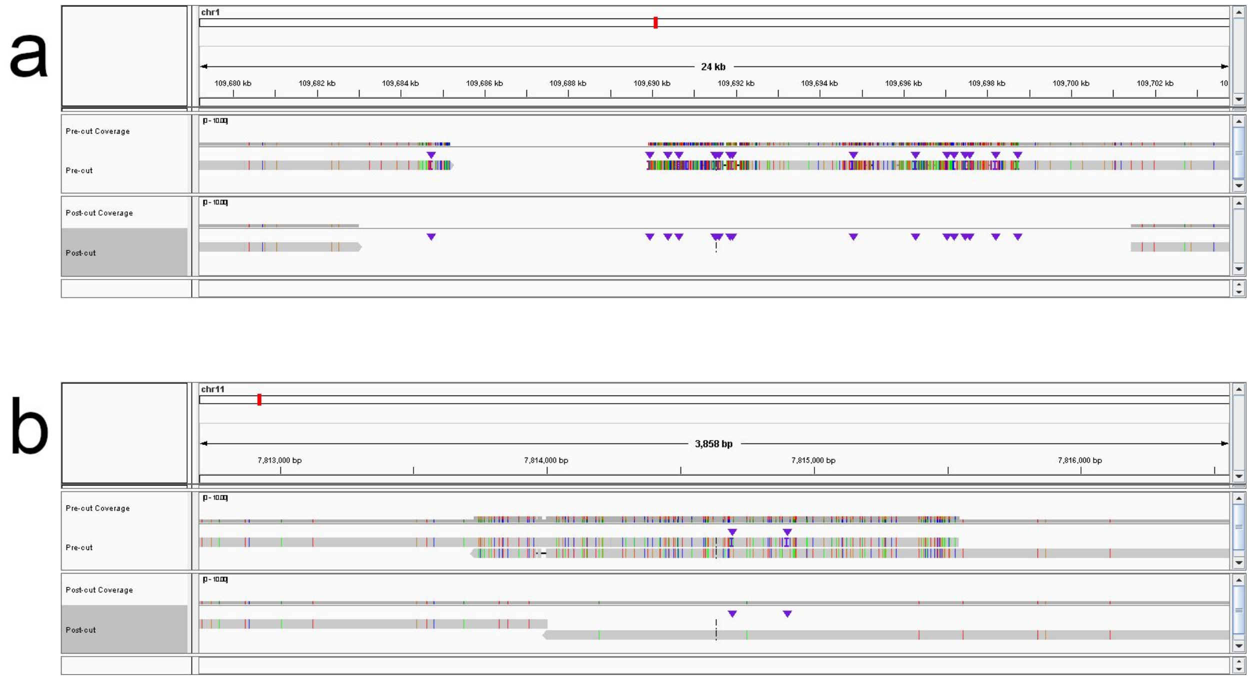


**Fig. S16. PAV alignment trimming**

a) A contig alignment was fragmented into two records around an SV deletion (18.4 kbp). This deletion was flanked by an SD repeat, and the single repeat copy in the contig was aligned to each reference copy (top track). Alignment trimming to remove multiply-mapped contig bases trimmed back the contig map removing as many non-sequence-match CIGAR operations (op codes I, D, and X) as possible (bottom track). b), A contig alignment was fragmented around an SV insertion (10.1 kbp tandem duplication). The contig has an extra copy of the duplication, and both were aligned to the same reference copy (top track). PAV trimming removed multiply-mapped reference bases leaving unaligned bases (the insertion) placed at the most likely breakpoint (bottom track).

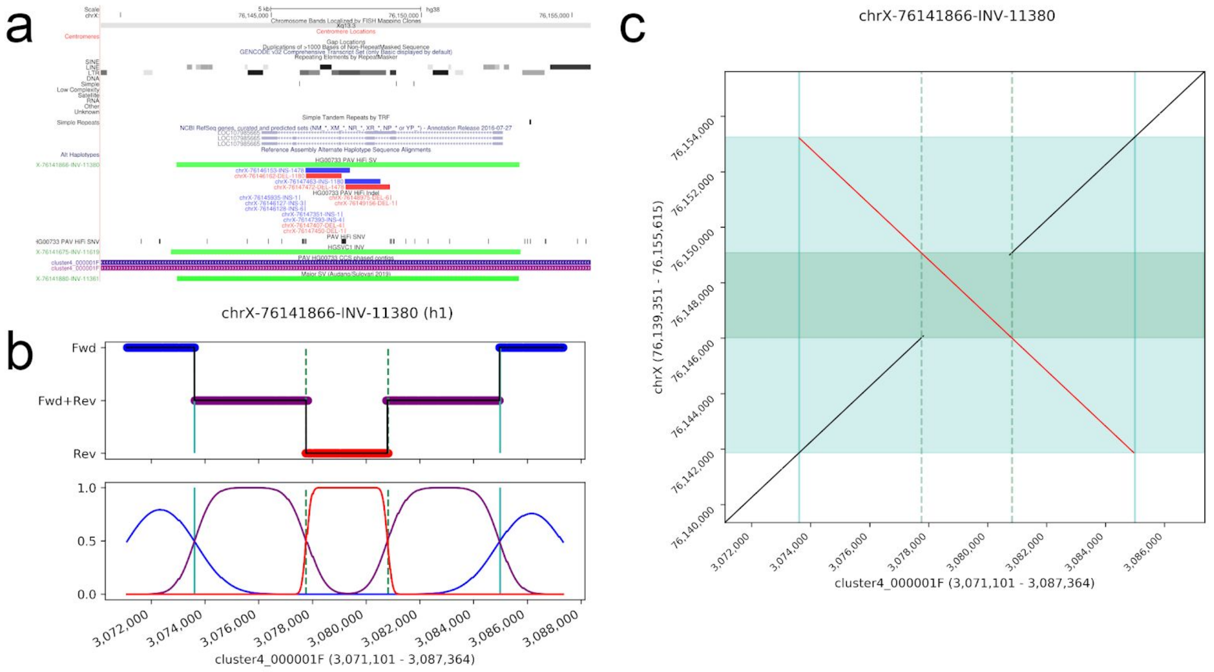


**Fig. S17. PAV flags inversion sites**

a) PAV flagged a potential inversion site by identifying matched SV insertions and deletions of a similar size. In this region, the contig alignments (pink and purple bars) were not broken by the inversion. Clusters of indels and SNVs can also be seen near the center of the inversion. These events were used to seed an inversion search. b) The k-mer density plot showing reference-oriented k-mers (blue), k-mers in reference and reverse orientation (purple), and k-mers strictly in reverse orientation (red). Top panel is the k-mers, and bottom panel is the scaled density plot for each orientation class. This inversion is flanked by a large repeat, which can be seen as large stretches of k-mers in both orientations, and the boundaries become the outer breakpoint (fwd to fwd-rev) and inner breakpoint (fwd-rev to rev) where the true inversion breakpoint is somewhere between these. c) The dotplot of the resolved inversion. The inner strictly inverted region between inner breakpoints is shown in dark green (reference) and solid lines (contig), and the inverted duplications are shown as light green regions (reference) and the area between the solid and dashed lines (contig). The inversion call itself is reported using the outer breakpoints, and the inner breakpoints are carried as an annotation.

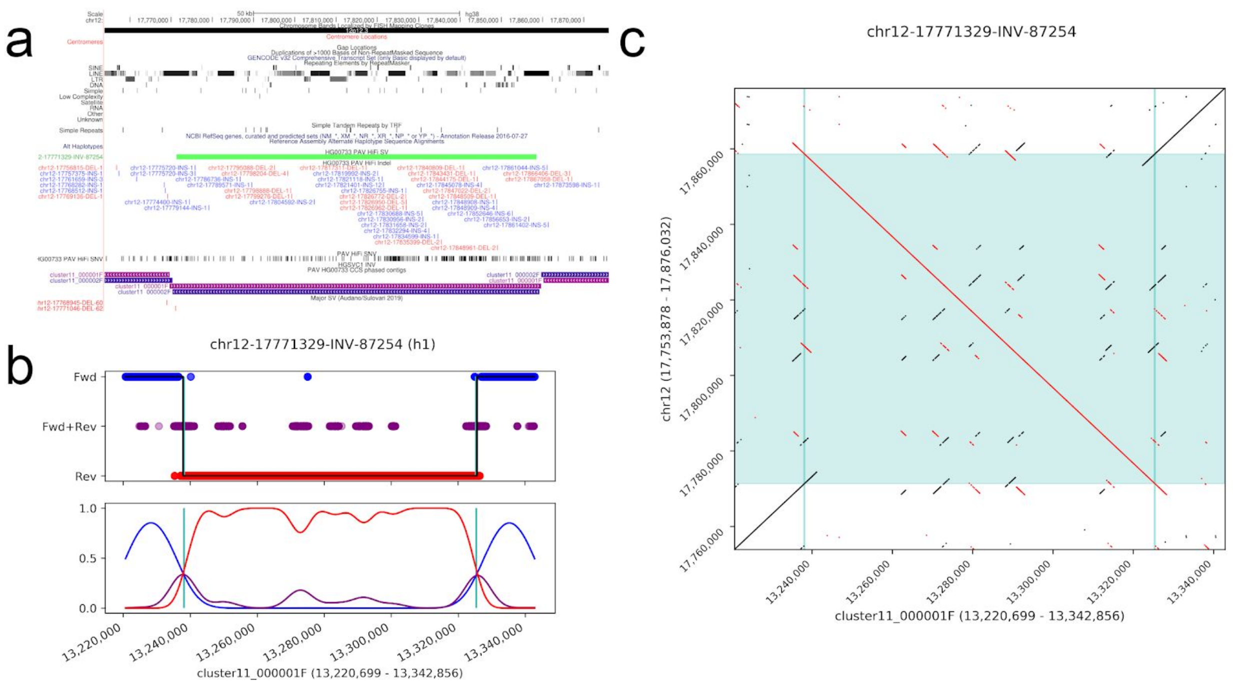


**Fig. S18. PAV flags inversion sites**

a) Contig alignments (h1 pink, h2 purple) truncated at inversion breakpoints exhibiting prototypic alignment structure over an inversion with orientation reversed inside the inversion. b) The k-mer density plot showing reference-oriented k-mers (blue), k-mers in reference and reverse orientation (purple), and k-mers strictly in reverse orientation (red). Top panel is the k-mers, and bottom panel is the scaled density plot for each orientation class. This inversion has no clear inner breakpoint. c) The dotplot with the contig (horizontal) and reference (vertical) with inversion flanks in reference orientation.

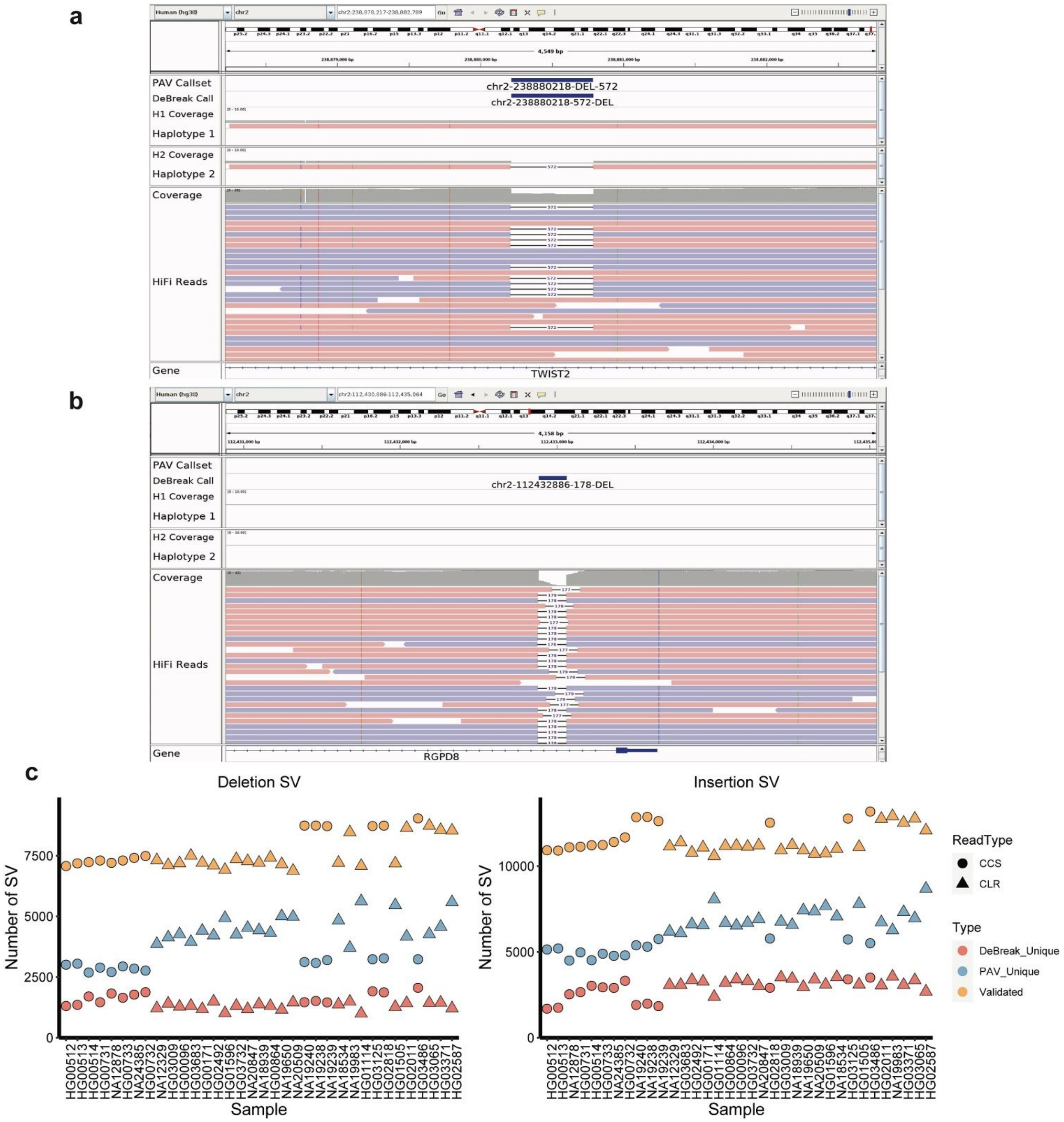


#### Fig. S19. DeBreak variant discovery

a) An example of a deletion detected by both DeBreak and PAV in HG00733 HiFi sample; b) An example of a deletion detected by DeBreak only. No assembly contigs were aligned to this region; c) Number of SVs detected by DeBreak only, PAV only, and by both callers for deletion (left) and insertion (right) SVs in all samples.

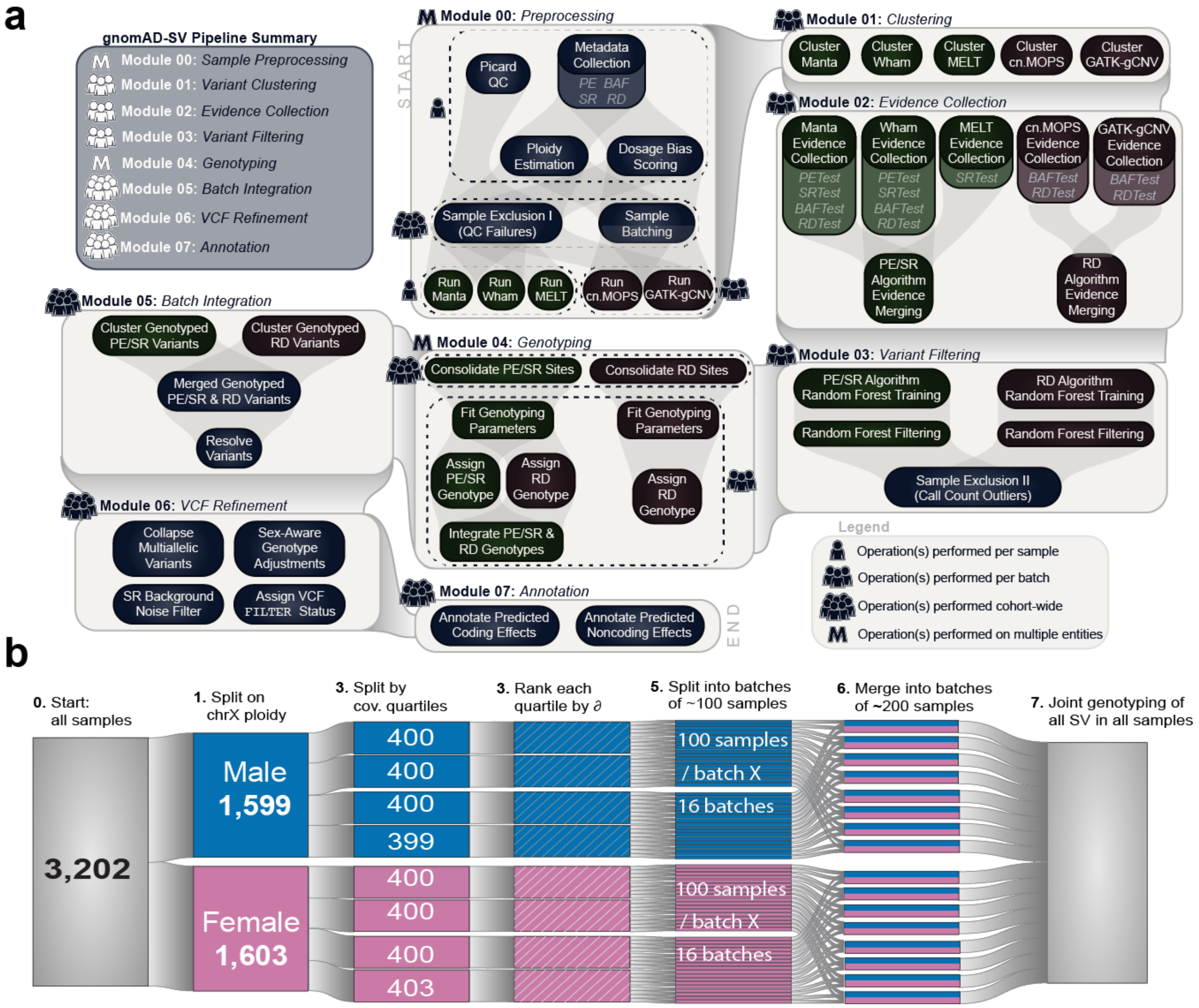


**Fig. S20. Overview of GATK-SV pipeline**

a) An overview of the GATK-SV pipeline is summarized here, and details of the method were described in Collins et al. (Collins et al. 2020) as outlined in detail in Methods. The GATK-SV discovery pipeline contains seven sequential modules (light beige boxes). The sequence of modules is listed in the top left panel and is also indicated by connections between light beige boxes. Each module contains multiple sub-modules (smaller, dark boxes) that operate on the per-sample (N=1), per-batch (N~200; see Fig S21 for a description of sample batching scheme), or cohort-wide (N=3,202) level. This pipeline has been made available as a series of publicly accessible methods on FireCloud/Terra to permit cloud-based analyses of SVs across WGS studies; b) Batching strategy applied in GATK-SV. The 3,202 samples were grouped into 200 sample batches for efficient processing of the GATK-SV pipeline. Details of the strategy are described in the supplementary methods (Supp Material 12.2.2.1).

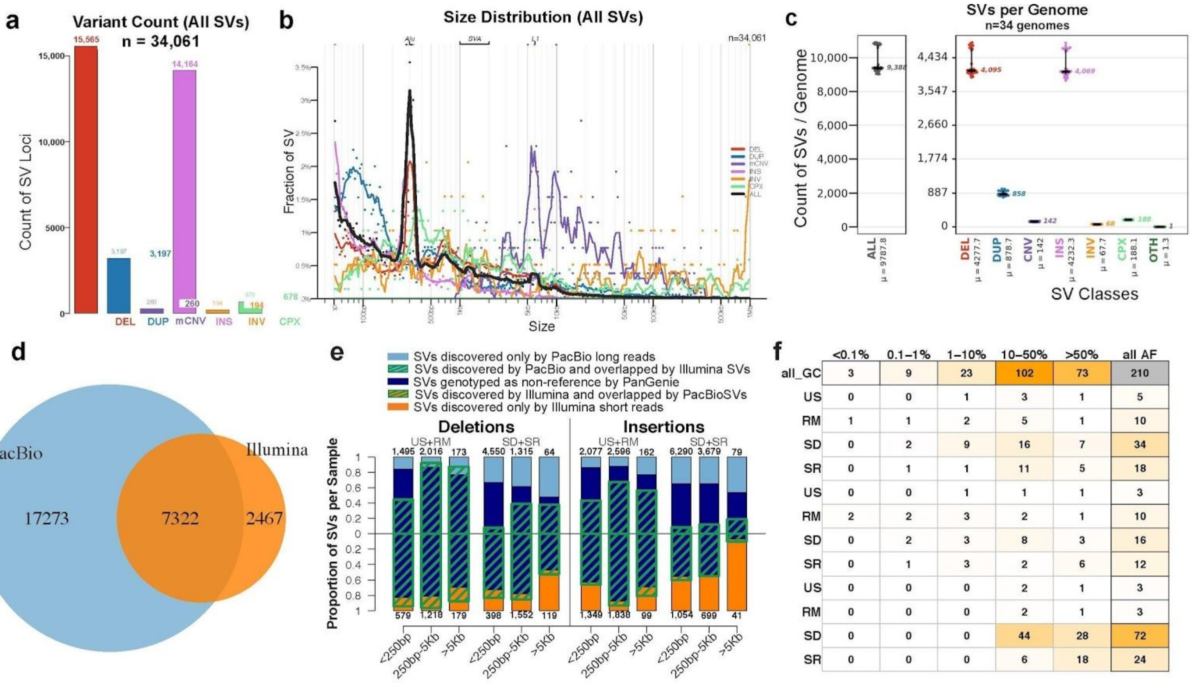


#### Fig S21.Overview of the short-read Illumina integration callset on the 34 genomes with matched PacBio sequences

a) Count of SV loci by variant type in the Illumina integration callset; b) Size distribution of SVs; c) Count of SVs per sample by variant type; d) Comparison of SV counts per sample detected by PacBio and Illumina sequences; e) Concordance of SVs between Illumina, PacBio and PanGenie genotypes by genomic location and SV sizes. US - unique sequences, RM - repeat mashed regions, SD - segmental duplications, SR - simple repeats; f) Distribution of large CNVs (>5Kb) specifically discovered by Illumina across genomic locations and ranges of allele frequencies. Abbreviations of genomic context are the same as e).

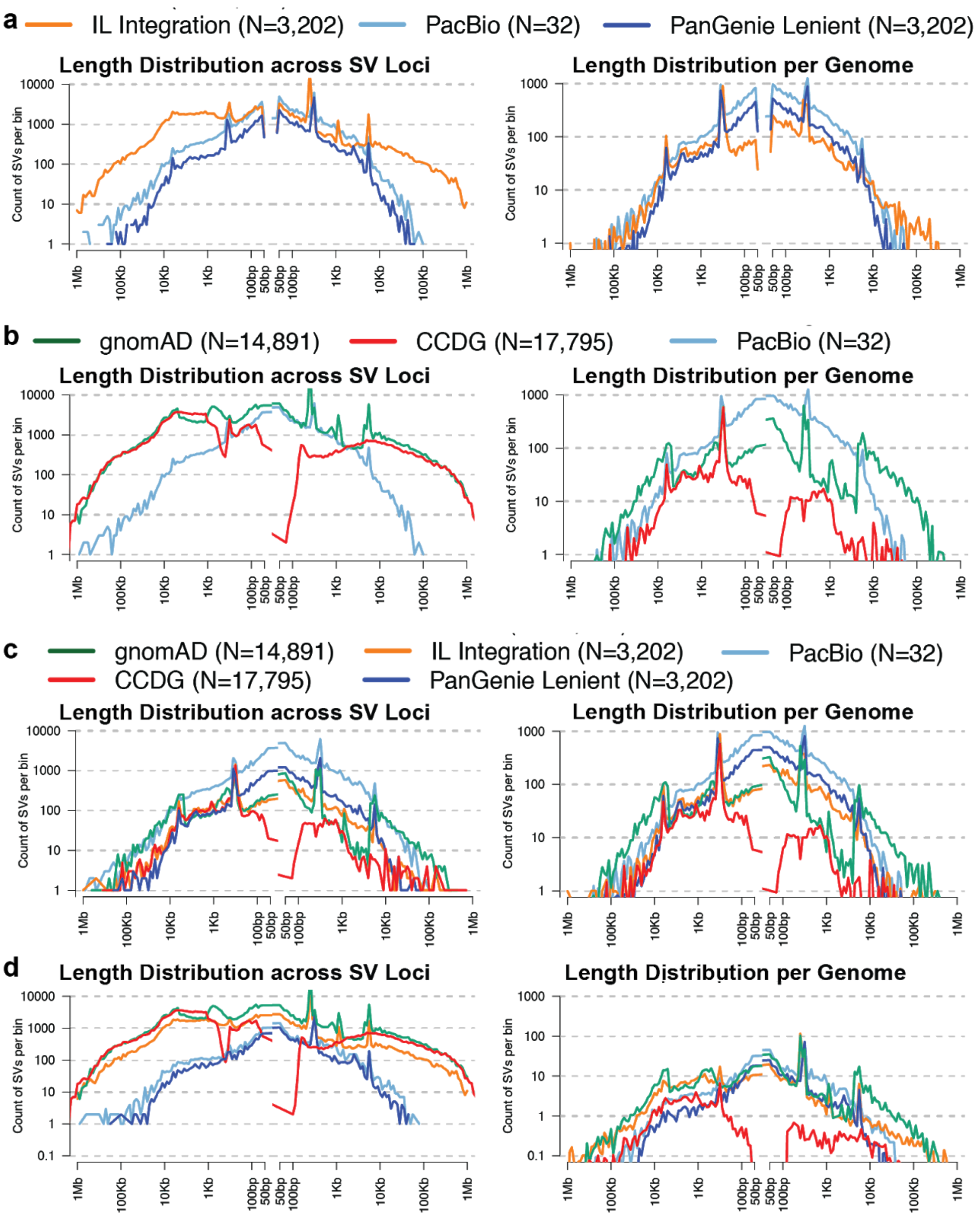


**Fig S22. Length distribution of SVs from HGSV, gnomAD and CCDG**

a) Length distribution of SVs from the HGSV Illumina integration callset (orange), PacBio callset (light blue) and PanGenie callset (blue). Left panel shows the overall distribution of all SV loci, and the right panel shows the averaged distribution per genome. b) Length distribution of SVs from the HGSV PacBio callset (light blue), gnomAD (green) and CCDG (red). Left panel shows the overall distribution of all SV loci, and the right panel shows the averaged distribution per genome. c) Length distribution of common SVs that are of 5% or higher allele frequencies from the HGSV Illumina integration (orange), PacBio (light blue), PanGenie (blue), gnomAD (green) and CCDG (red). Left panel shows the overall distribution of all SV loci, and the right panel shows the averaged distribution per genome. d) Length distribution of rare SVs that are of 5% or lower allele frequencies from the HGSV Illumina integration (orange), PacBio (light blue), PanGenie (blue), gnomAD (green) and CCDG (red). Left panel shows the overall distribution of all SV loci, and the right panel shows the averaged distribution per genome.

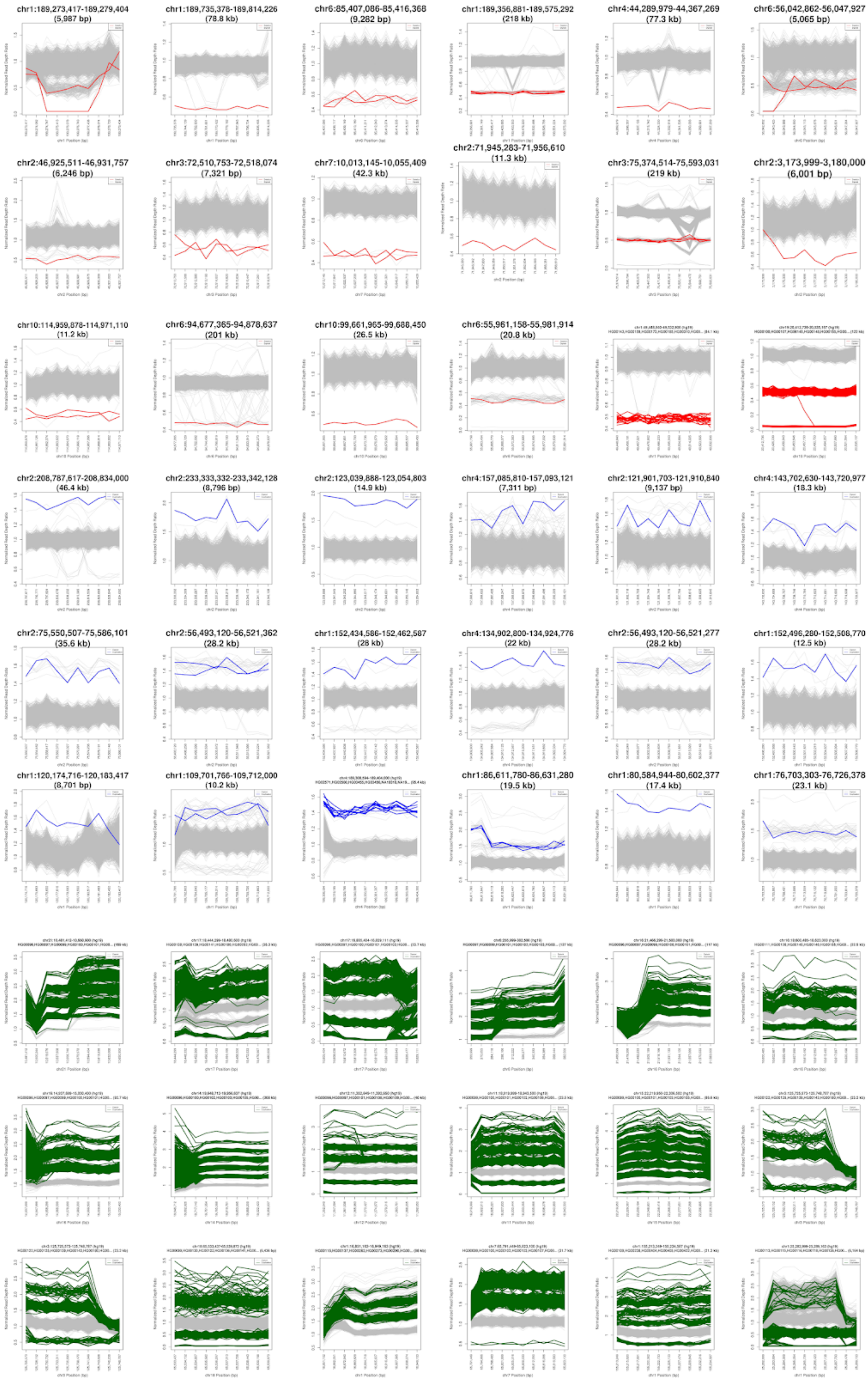
chr1:189,273,417-189,279,404
(5,987 bp)
chr1:189,735,378-189,814,226
(78.8 kb)
chr6:85,407,086-85,416,368
(9,282 bp)
chr1:189,356,881-189,575,292
(218 kb)
chr4:44,289,979-44,367,269
(77.3 kb)
chr6:56,042,862-56,047,927
(5,065 bp)
chr2:46,925,511-46,931,757
(6,246 bp)
chr3:72,510,753-72,518,074
(7,321 bp)
chr7:10,013,145-10,055,409
(42.3 kb)
chr2:71,945,283-71,956,610
(11.3 kb)
chr3:75,374,514-75,593,031
(219 kb)
chr2:3,173,999-3,180,000
(6,001 bp)
chr10:114,959,878-114,971,110
(11.2 kb)
chr6:94,677,365-94,878,637
(201 kb)
chr10:99,661,965-99,688,450
(26.5 kb)
chr6:55,961,158-55,981,914
(20.8 kb)
chr2:208,787,617-208,834,000
(46.4 kb)
chr2:233,333,332-233,342,128
(8,796 bp)
chr2:123,039,888-123,054,803
(14.9 kb)
chr4:157,085,810-157,093,121
(7,311 bp)
chr2:121,901,703-121,910,840
(9,137 bp)
chr4:143,702,630-143,720,977
(18.3 kb)
chr2:75,550,507-75,586,101
(35.6 kb)
chr2:56,493,120-56,521,362
(28.2 kb)
chr1:152,434,586-152,462,587
(28 kb)
chr4:134,902,800-134,924,776
(22 kb)
chr2:56,493,120-56,521,277
(28.2 kb)
chr1:152,496,280-152,508,770
(12.5 kb)
chr1:120,174,716-120,183,417
(8,701 bp)
chr1:109,701,766-109,712,000
(10.2 kb)
chr1:86,611,780-86,631,280
(19.5 kb)
chr1:80,584,944-80,602,377
(17.4 kb)
chr1:76,703,303-76,726,378
(23.1 kb)

**Fig S23. Distribution of normalized sequencing depth of CNVs**

A randomly selected subset of CNVs that are over 5 kbp in size, discovered by short-read Illumina sequences but were missed by long-read PacBio sequences are shown in this plot. Y-axis represents the normalized sequencing depth of each sample, with 1 representing copy number 2, 0.5 representing copy number 1, etc. Samples with deletions, duplications, and multi-allelic CNVs were shown in red, blue, and green colors, respectively.

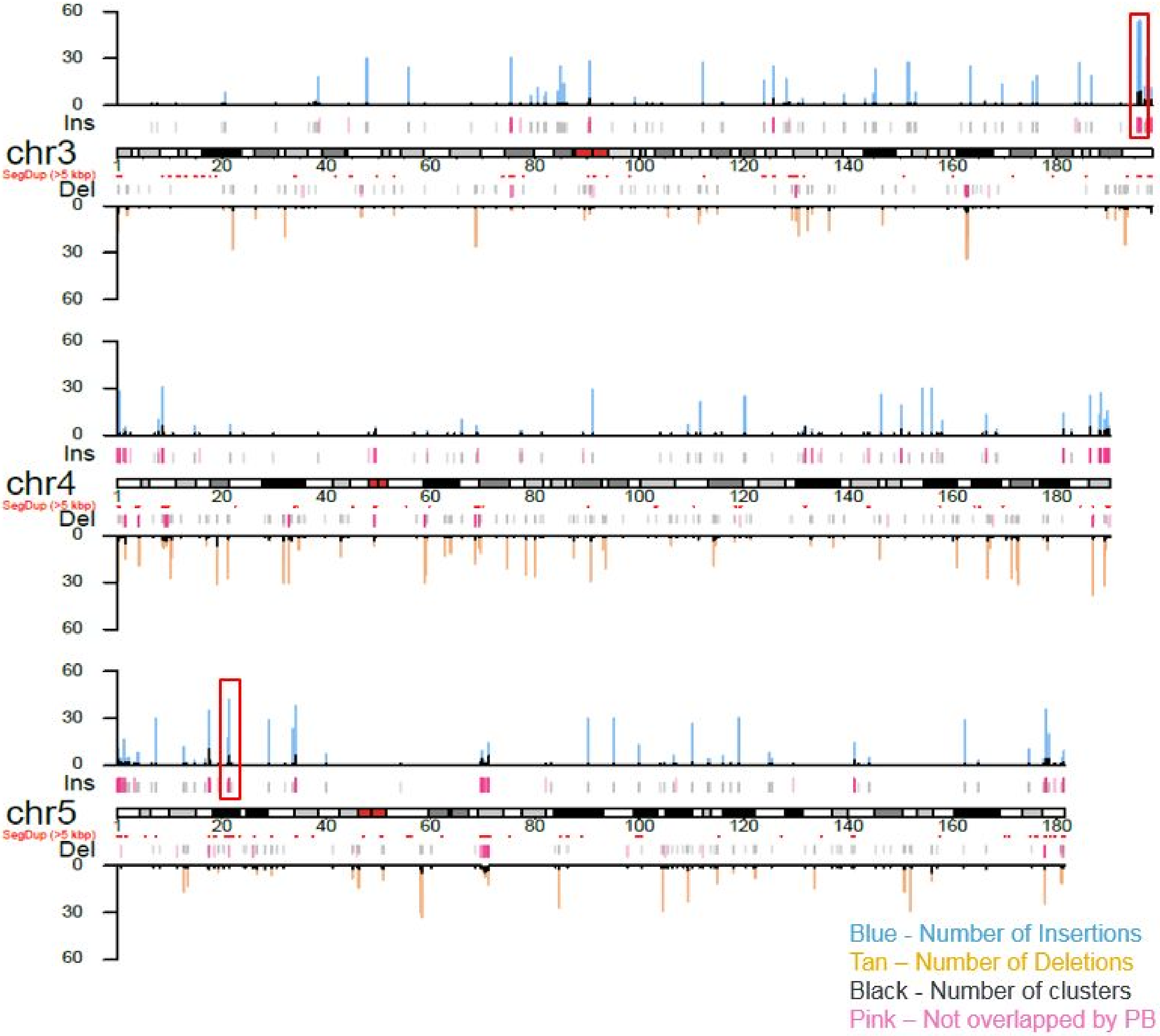


**Fig. S24. Ideogram showing Bionano calls (≥5 kbp) and clusters**

Large complex SV sites are regions with a high number of SV calls and with at least five clusters. Boxed in red on Chr3 and Chr5 are examples of these complex polymorphic regions with population configurations shown in Fig S27 (chr5) and Fig S28 (chr3). Please refer to the separate PDF version of this figure for the full-size image showing the complete ideogram.

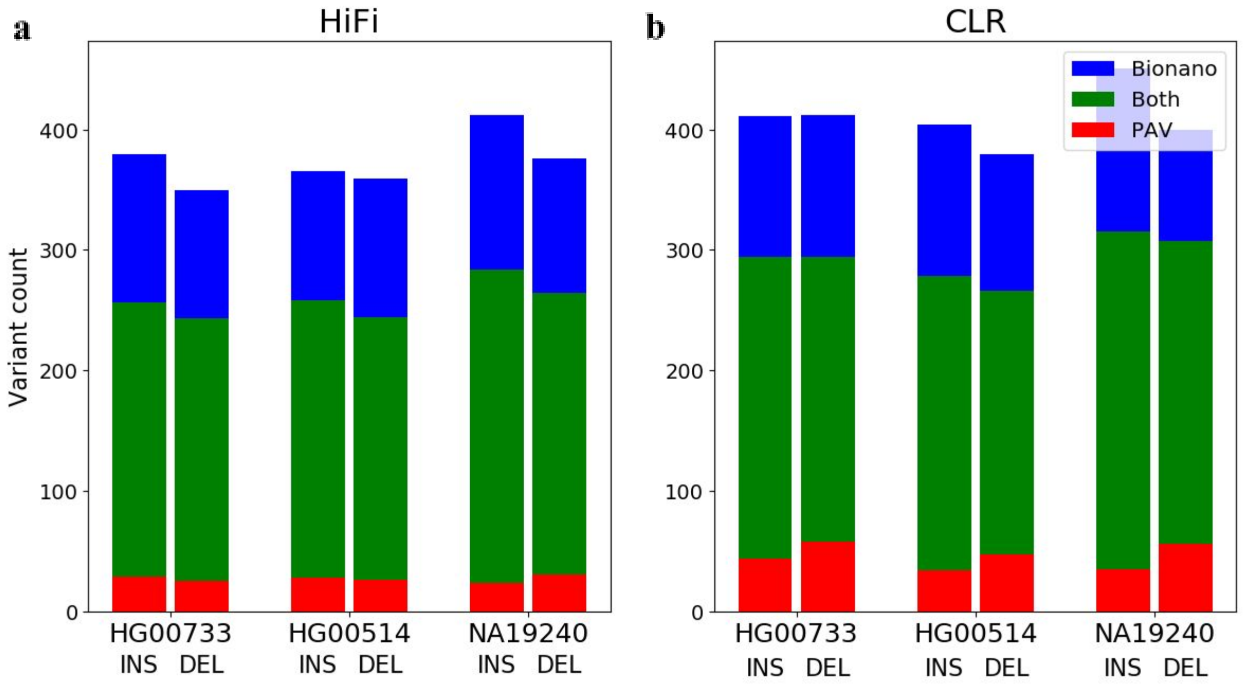


**Fig. S25. Bionano and PAV intersection for SVs greater than 5 kbp**

Each bar is a Venn diagram showing PAV only (red), PAV and Bionano (green), and Bionano only (blue) for SVs 5 kbp and greater in child samples for HiFi (**a**) and CLR (**b**).

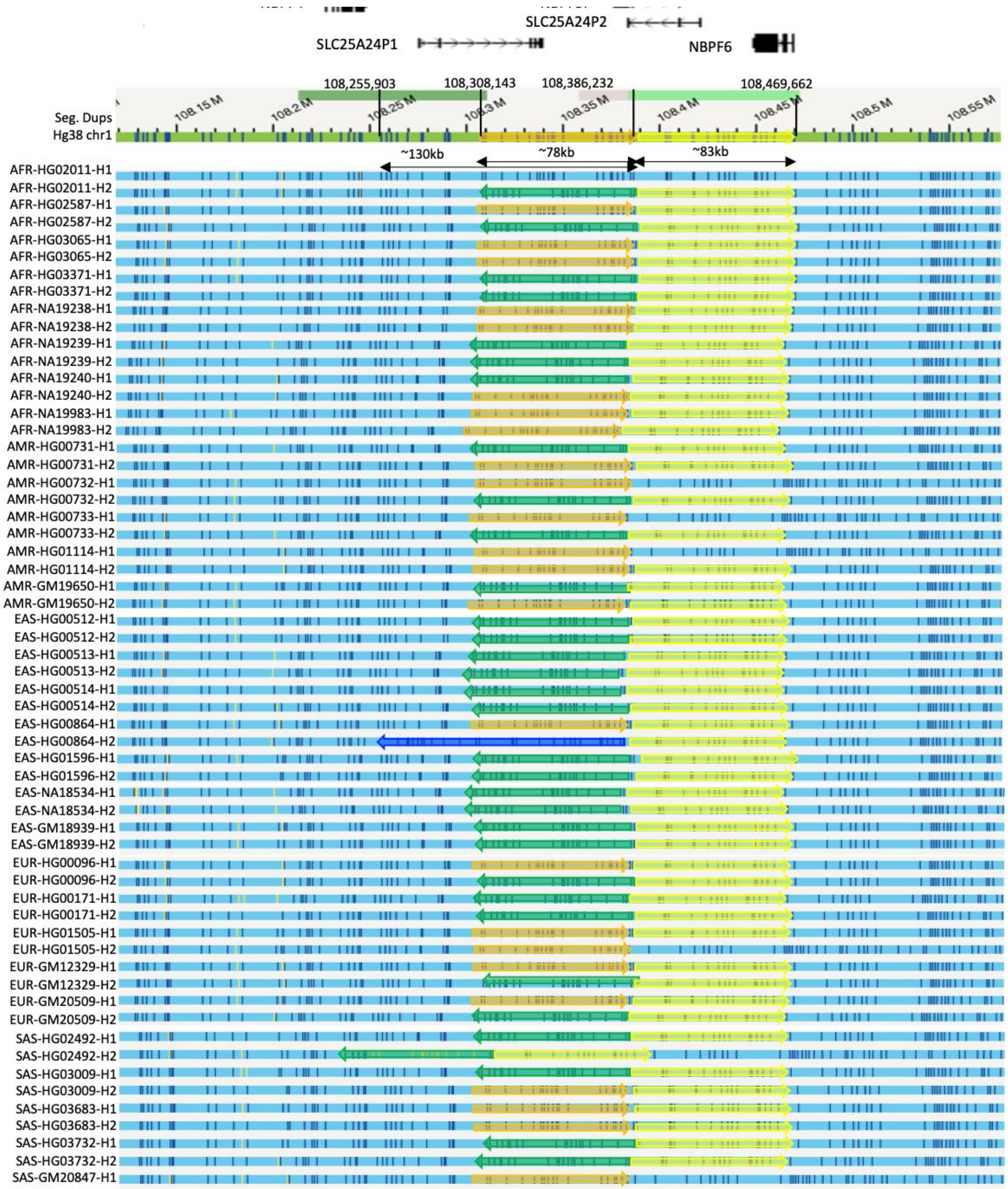


**Fig. S26. Population distribution of SVs identified on chromosome 1**

Gene annotation (GENCODE v34) at the upper panel, and SDs between 108.2–108.3 Mbp and 108.35–108.5 Mbp. Green horizontal line represents GRCh38 reference assembly and blue horizontal lines represent sample contigs. Orange and yellow colored arrows represent the loci of inversions and deletions; green and blue arrows represent two different types of inversions.

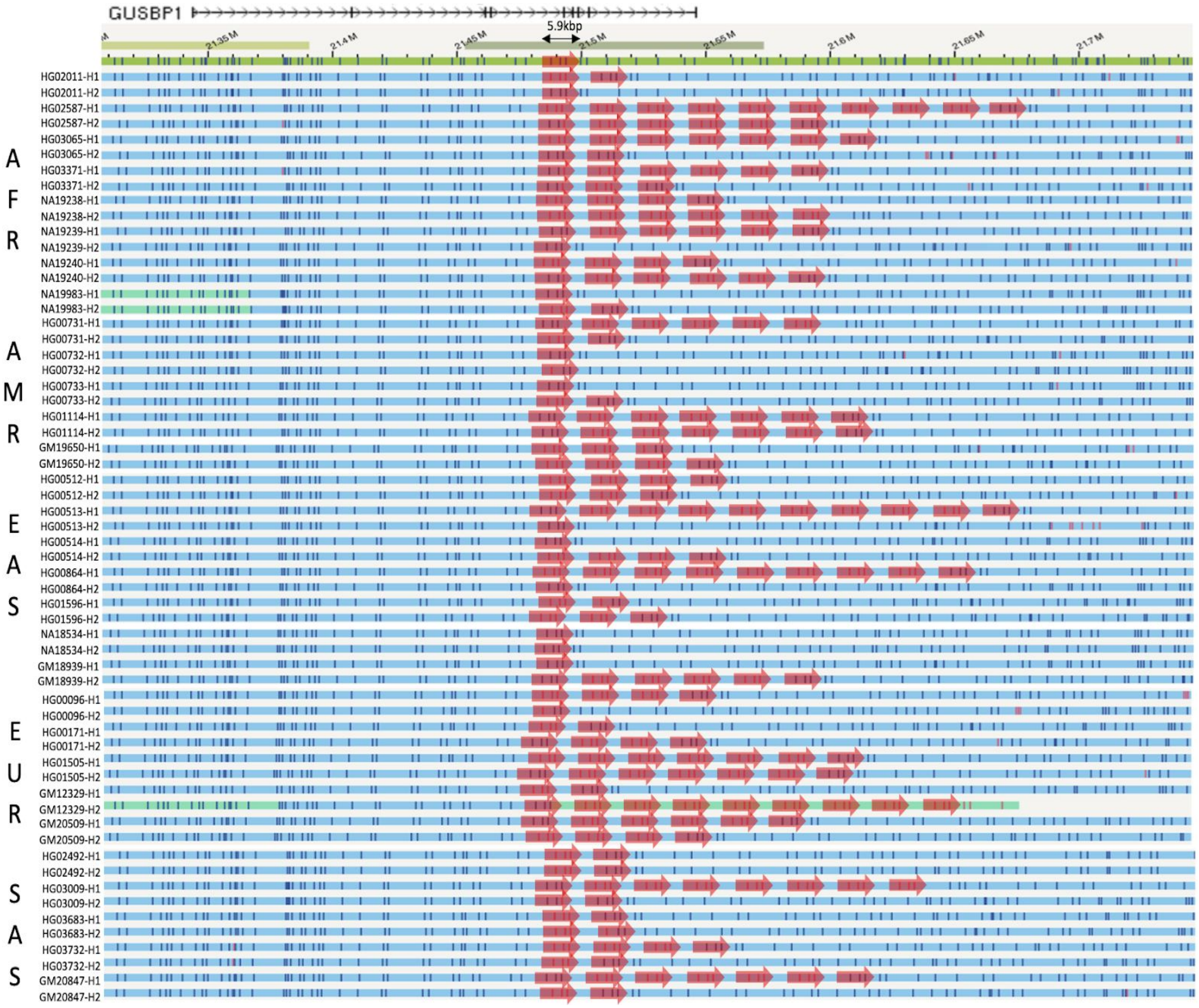


**Fig. S27. Population distribution of SVs identified on chromosome 5 (21.1–21.7 Mbp)**

Gene annotation (GENCODE v34) at the upper panel, and SDs in light and dark green. Green horizontal line represents GRCh38 reference assembly and blue horizontal lines represent sample contigs. Red colored arrows represent copy number gains.

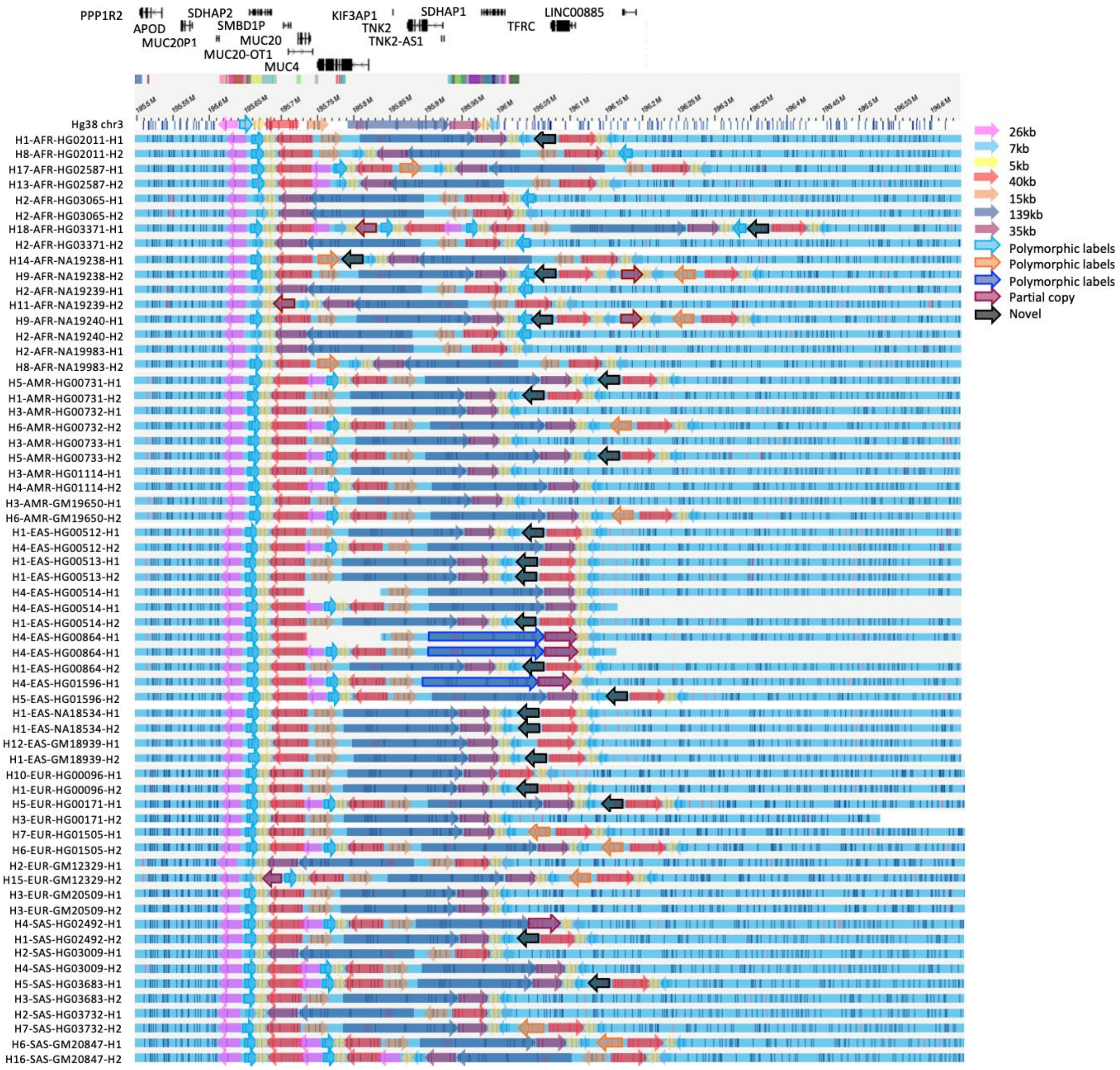


**Fig. S28. Full configuration of 3q29 region (195.4–196.1 Mbp)**

Gene annotation (GENCODE v34) at the upper panel, green horizontal line represents GRCh38 reference assembly, and blue horizontal lines represent sample contigs. SDs are represented as colored blocks above GRCh38. Colored arrows represent different types of configurations in the 3q29 region in each sample.

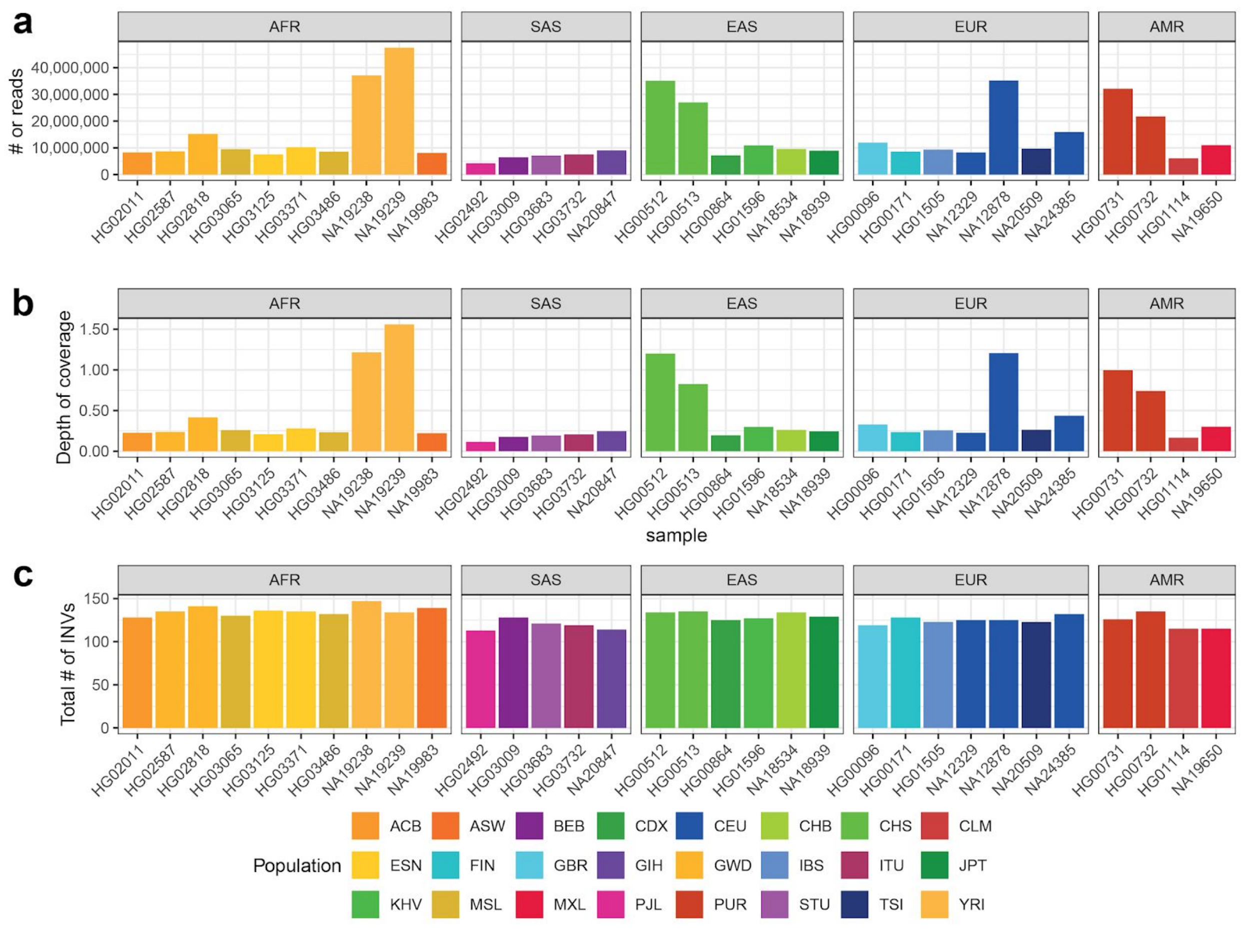


**Fig. S29. Composite files summary (n=32)**

Each Strand-seq composite file was created by concatenating reads across all informative Strand-seq libraries and homologs as previously described (*2*, *37*). a) Shows the number of reads in each sample and population-specific (AFR - African, SAS - South Asian, EAS - East Asian, EUR - European, AMR - American; Supplementary Table S33) composite file; b) A barplot showing a depth of coverage of each composite file. Strand-seq data (NA19238, NA19239, HG00512, HG00513, NA12878, HG00731, HG00732) from previous studies sequenced at higher depth; c) A barplot showing the total number of inversions detected by Strand-seq only per sample (n=32) showing that Strand-seq inversion calling is not skewed towards samples with a higher genome coverage.

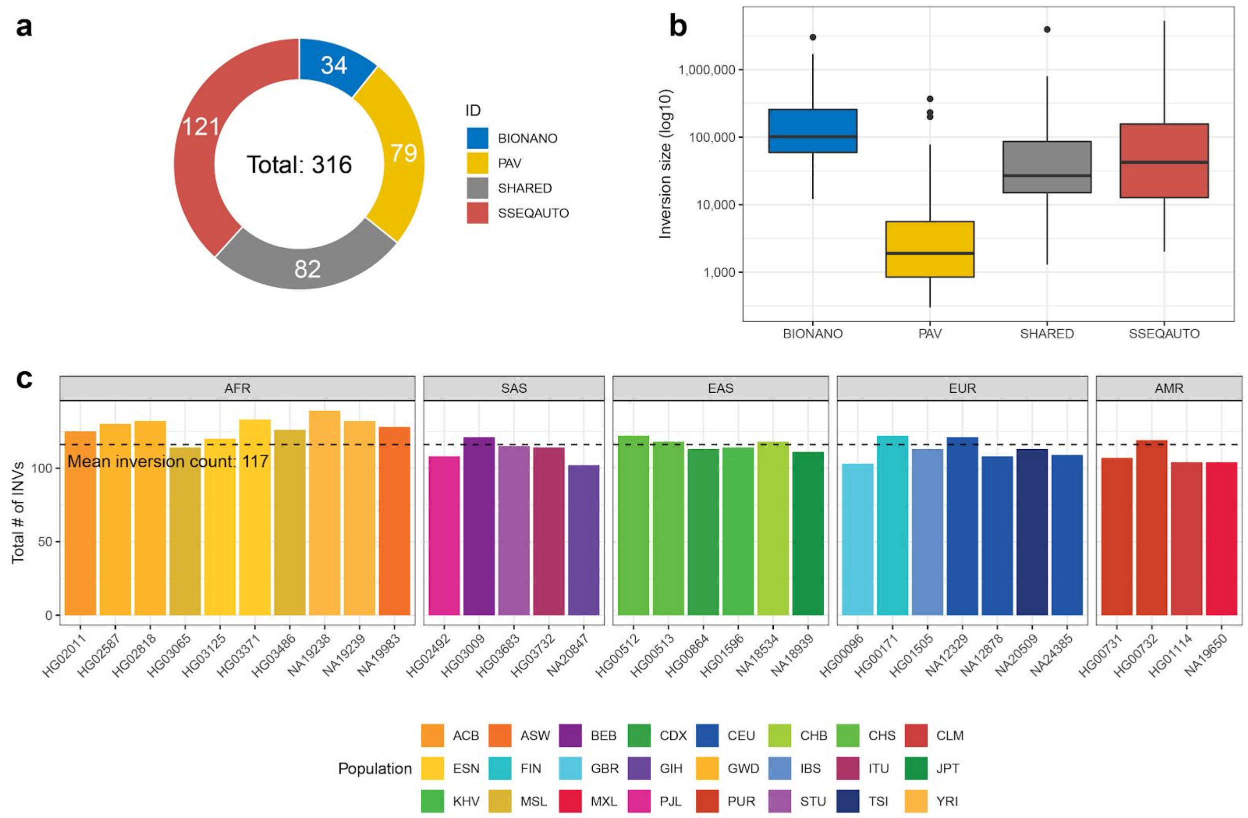


**Fig. S30. Inversion callset summary (n=316)**

a) A donut plot showing the total number of inversions called exclusively by a certain technology ('PAV' - phased assembly inversion caller, 'BIONANO' - Bionano optical maps, and 'SSEQAUTO' - automated Strand-seq inversion calls) along with inversions called by at least two independent technologies ('SHARED') (Number of Strand-seq inversion calls per category.); b) Size distribution of inversions calls per above-mentioned category; c) A barplot showing the total number of inversions per sample. Dashed horizontal line shows a median inversion count across all samples (n=32).

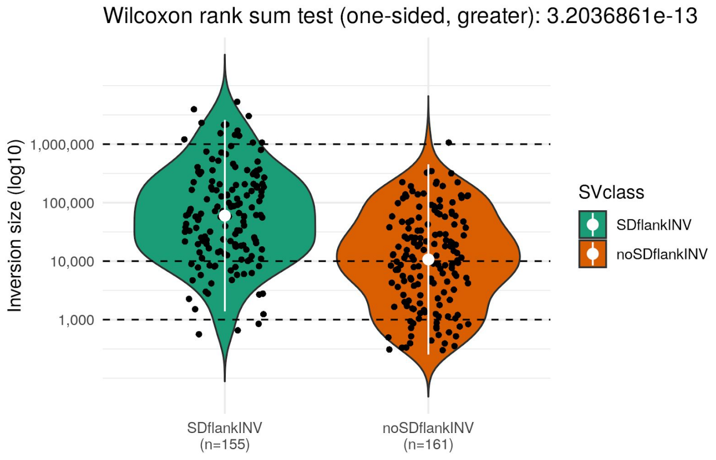


**Fig. S31. Inversions flanked by SDs**

Size distribution of inversions flanked by SDs (SDflankINV, n=155) and inversions not flanked by SDs (noSDflankINV, n=161). White dot shows the mean of each distribution along with IQR range.

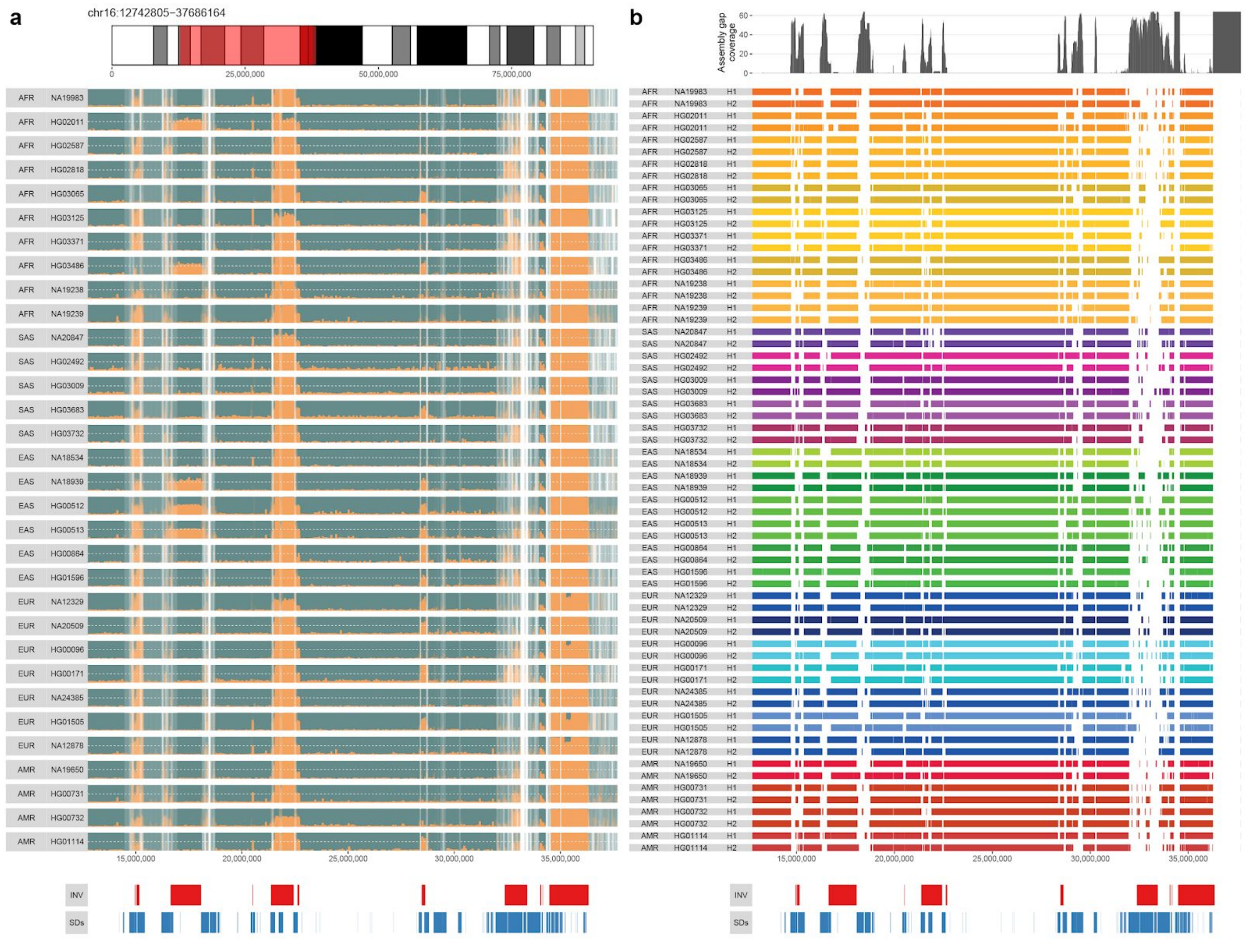


**Fig. S32. Phased assembly alignments in 16p12 region**

a) Top panel: Shows a region on chromosome 16 presented in detail in other figure panels. Middle panel: Shows binned (binsize: 10 kbp, stepsize: 5 kbp) ratio of Crick (positive strand, ‘+’, teal) and Watson (negative strand, ‘-’, orange) reads for each sample-specific composite file in a given region; b) Top panel: Gap coverage of phased assemblies aligned to GRCh38. Here, only regions where assembled contigs align with mapping quality ≤60 are summarized. Middle panel: Each row represents sample-specific haplotype-resolved assembly (H1 - haplotype1, H2 - haplotype2) aligned to GRCh38. Regions where assembled contigs do not align with mapping quality ≥60 are visible as white gaps between colored rectangles specific for each sample and haplotype; a) & b) Bottom panels: Red rectangles highlight inverted regions detected by Strand-seq while blue rectangles show the positions of SDs in GRCh38.

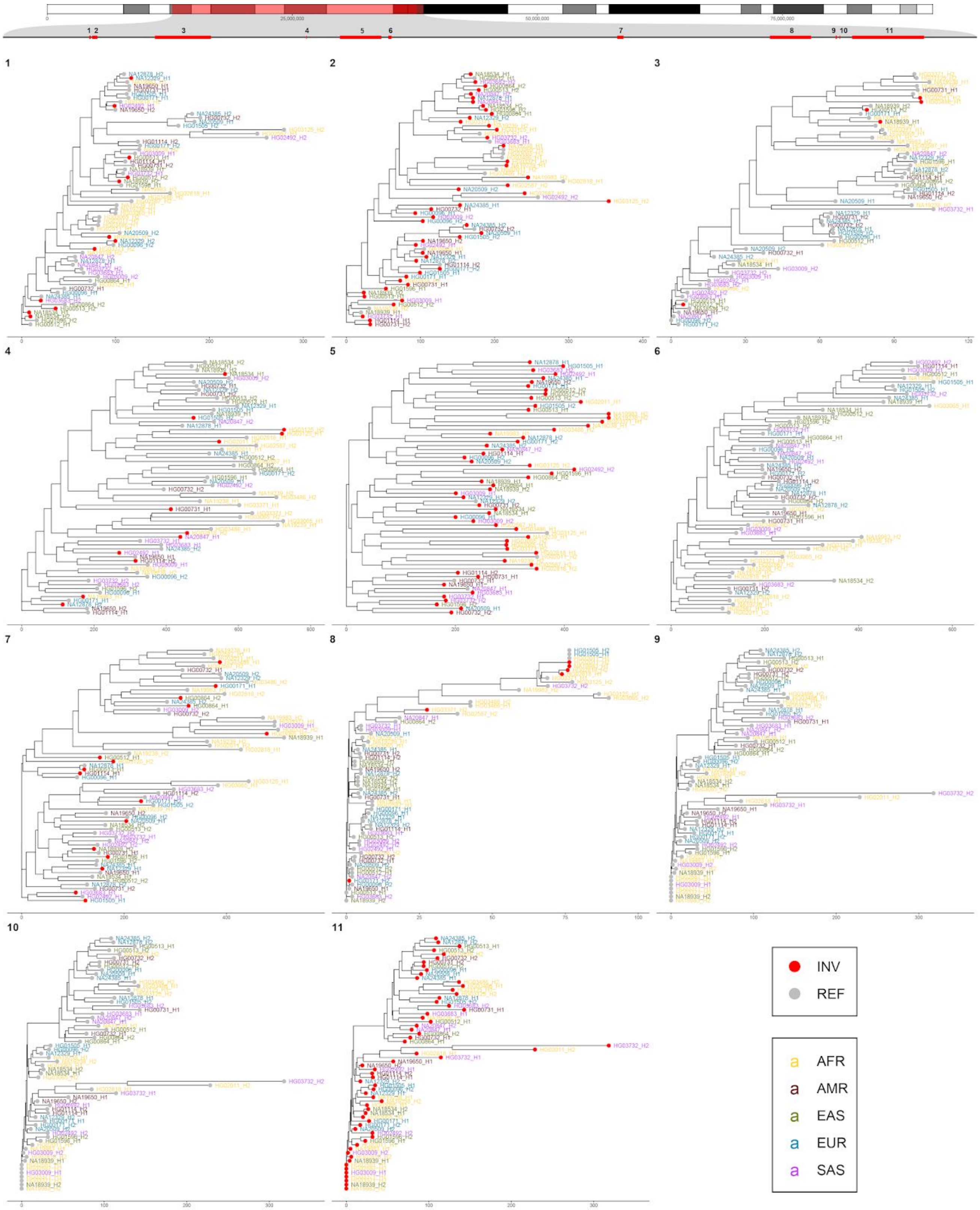
1 2 3 4 5 6 7 8 9 10 11
INV
REF
AFR
AMR
EAS
EUR
SAS

**Fig. S33. Genetic background of inverted haplotypes at the 16p12 chromosome region**

Top panel: A chromosome 16 ideogram with region of interest highlighted by red transparent rectangle. Below inverted regions are shown as red rectangles with numbers pointing to a neighbor-joining tree constructed based on phased SNVs from inversion flanking region (+/-500 kbp). SNVs from within the inverted region and those overlapping SDs (≥98% identity) were not considered for the tree construction. Haplotypes flanking inverted regions are highlighted by a red dot and those with a reference orientation (or unknown phase) are shown in gray. Each sample and haplotype label is colored based on the superpopulation of origin. Given the fact that this analysis does not take into account meiotic recombination, we suggest that inversions 1, 3, 4, 7 and 8 show recurrent traits as they seem to occur at different genetic backgrounds.

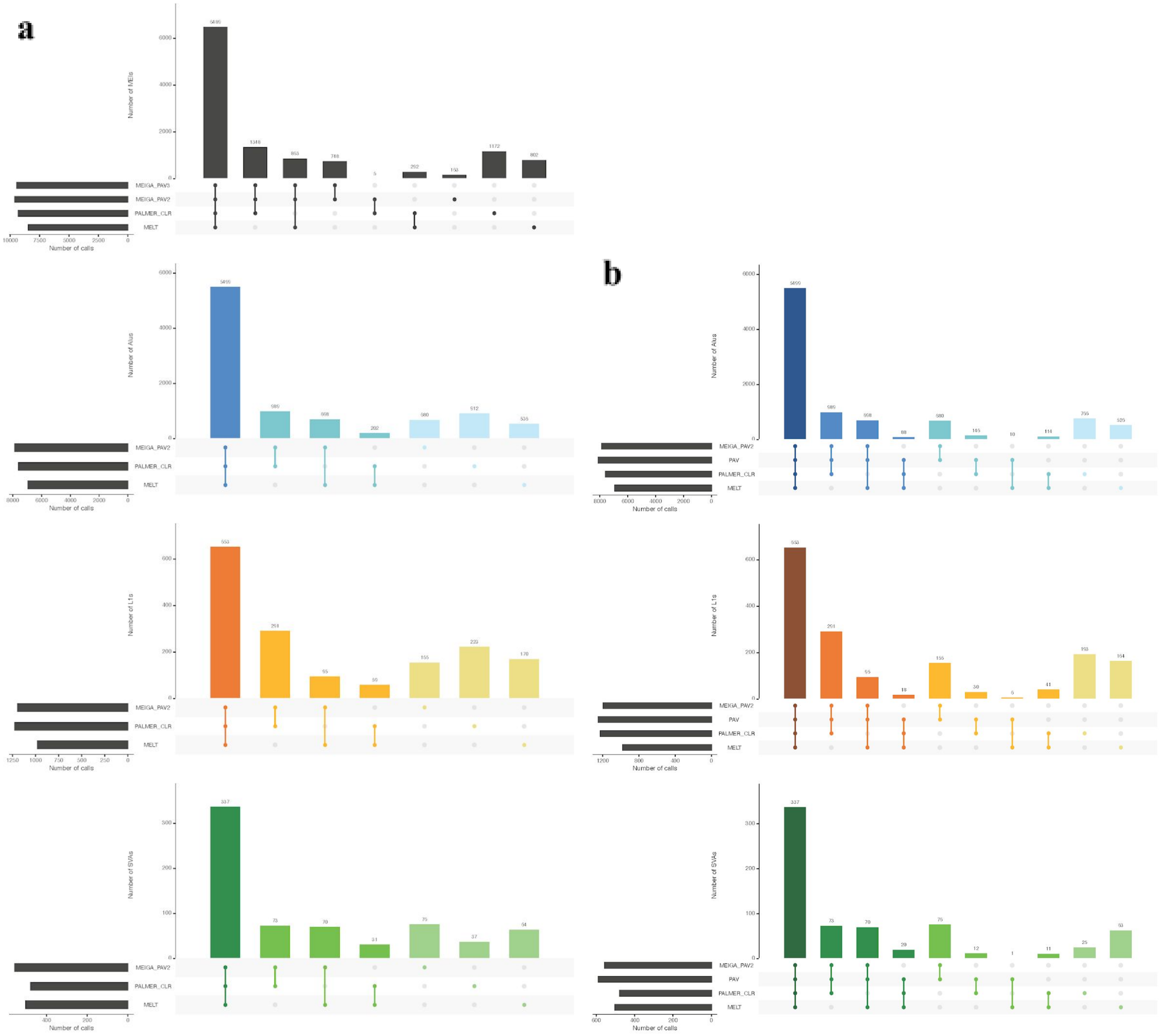


**Fig. S34. UpSet plots for the integrated callset and PAV annotation in all HGSVC2 samples**

**a**) the integration strategy was conducted on three categories of MEIs, *Alu* (blue), L1Hs (orange), and SVA (green), separately. The overall MEI intersection was shown by the top panel with black bars. Three calling methods are included: MELT (independent caller for Illumina), PALMER (PALMER_CLR, independent caller for PacBio mapping-based), and MEIGA_PAV (MEIGA_PAV2: the pre-filtered PAV assembly-based callset with MEIGA annotation; MEIGA_PAV3: the final PAV assembly-based callset with MEIGA annotation)

**b**) four callsets are included: MELT (independent caller for Illumina), PALMER (PALMER_CLR, independent caller for PacBio mapping-based), MEIGA_PAV (MEIGA_PAV2: the pre-filtered PAV assembly-based callset with MEIGA annotation), and PAV (the final PAV assembly-based callset).

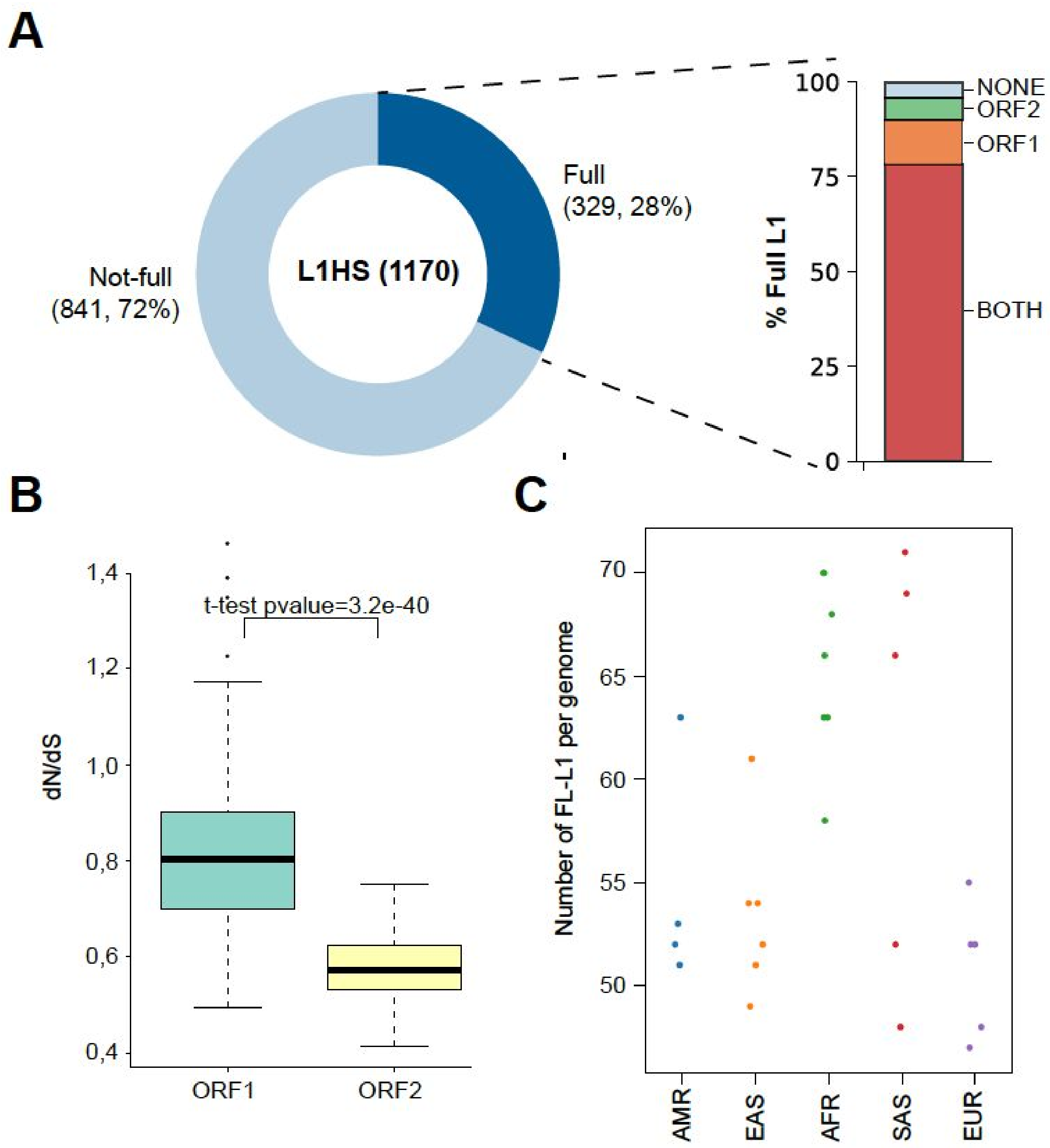


**Fig. S35. Collection of sequence-resolved full-length L1s**

a) Left, donut chart showing the fraction of full-length L1s (FL-L1s) among all germline L1 insertions identified on the HGSVC2 dataset. Right, ORF status for full-length L1s displayed as red (intact ORFs), orange (truncated ORF1), and green (truncated ORF2) stacked bars; b) dN/dS ratios for ORF1 and ORF2 in FL-L1s. *P* values indicate significance from a two-tailed Mann–Whitney U-test; c) Number of non-reference FL-L1s per donor. Donors stratified according to their population ancestry.

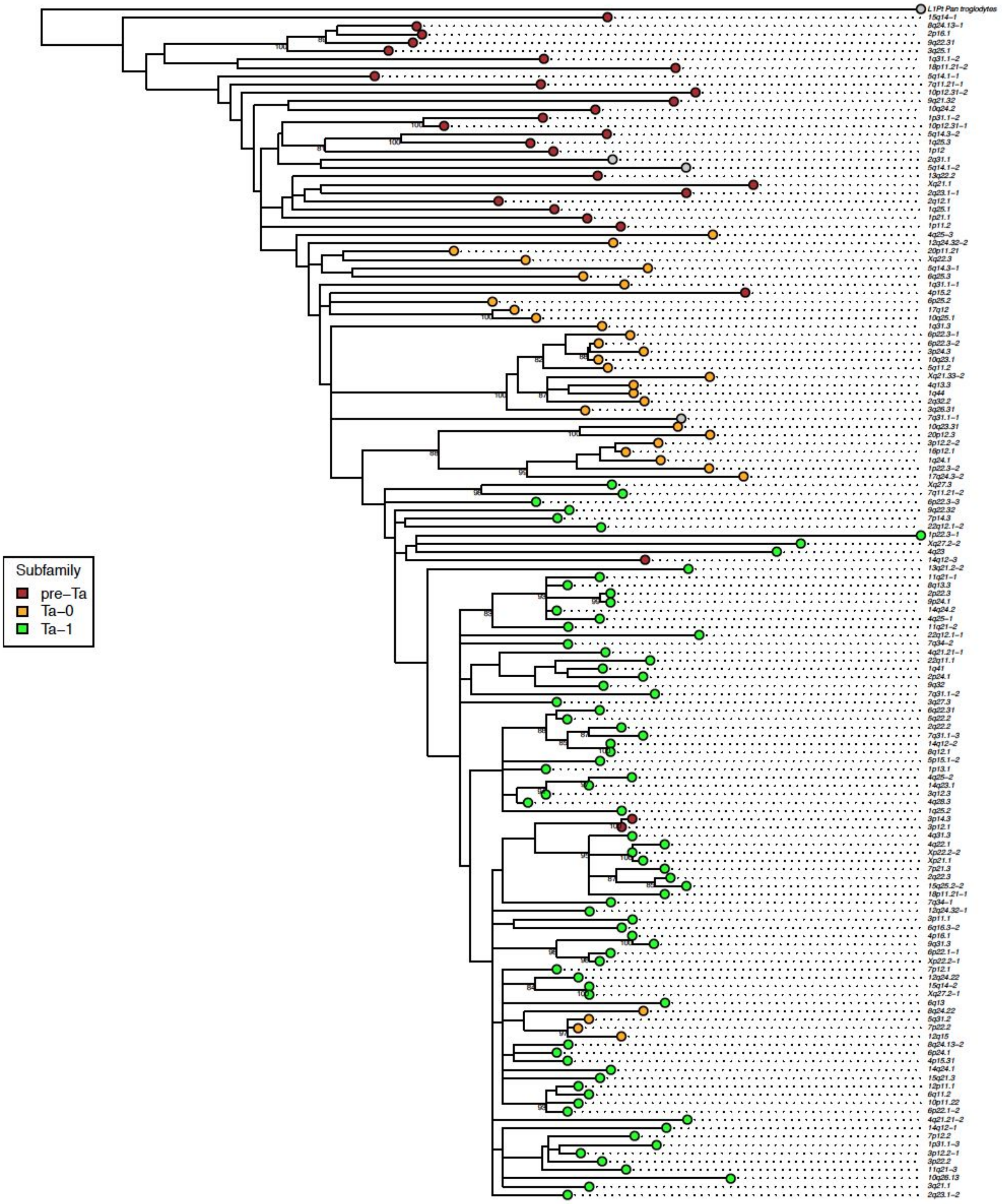


**Fig. S36. Complete phylogeny for all active sequence-resolved Full-Length (FL)-L1s**

Sequences annotated by subfamily designation. L1 *Pan troglodytes* (L1Pt) included as an outgroup. Bootstrap support values >80% indicated.

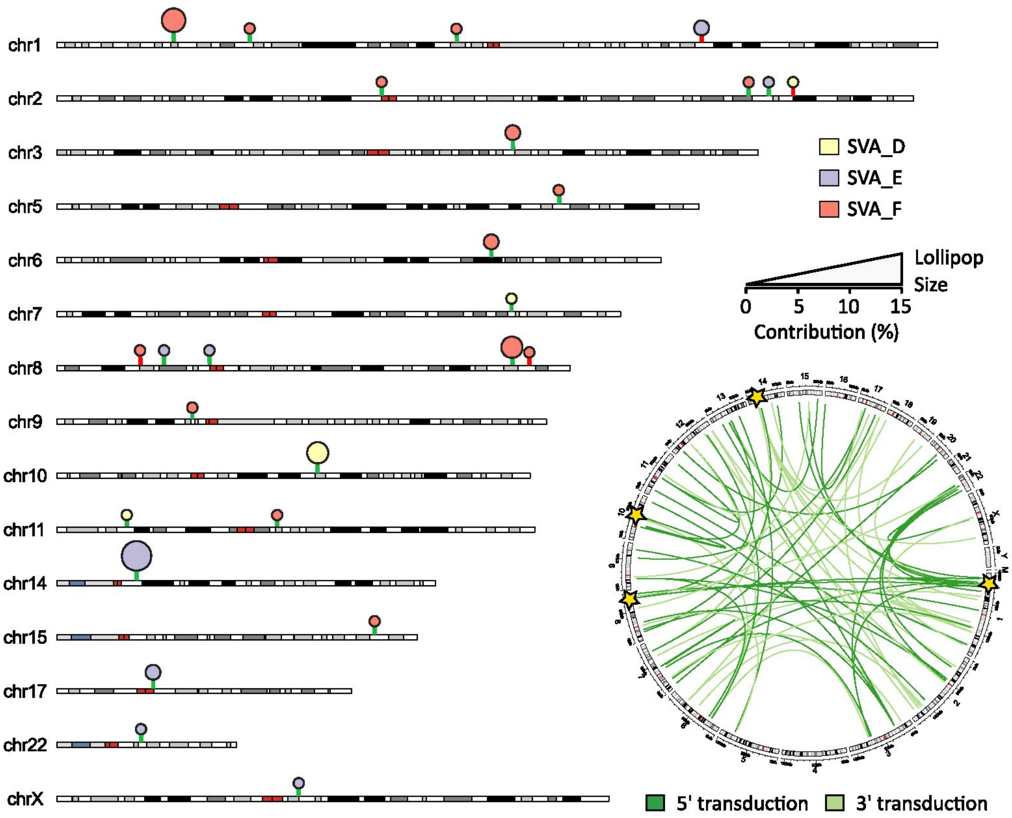


**Fig. S37. Catalogue of source SVA active in HGSVC2 dataset**

Left, chromosomal map with germline SVA source SVA elements as lollipops. Each lollipop is color-coded according to the source SVA subfamily, while lollipop sticks are colored in green and red to indicate if the element is a reference or non-reference insertion, respectively. The contribution of each source locus (expressed as a percentage) to the total number of germline transductions identified is represented as a gradient of lollipop size, with top contributing elements exhibiting larger sizes. Right, circos plot showing all SVA-mediated transductions identified at HGSVC2 dataset: 5' transductions are colored in dark green while 3' transductions in light green. Particularly active source SVAs are highlighted with gold stars.

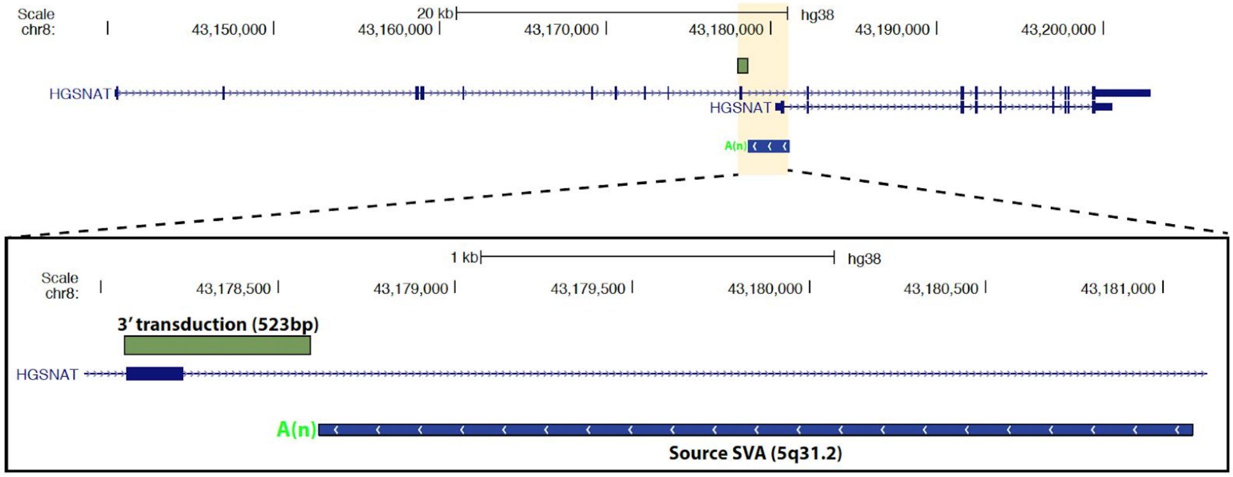


#### Fig. S38. Exon shuffling by intronic SVA source element

Partnered SVA-mediated 3' transduction mobilizes complete exon from *HGSNAT* to another genomic location at chr5:138736711.

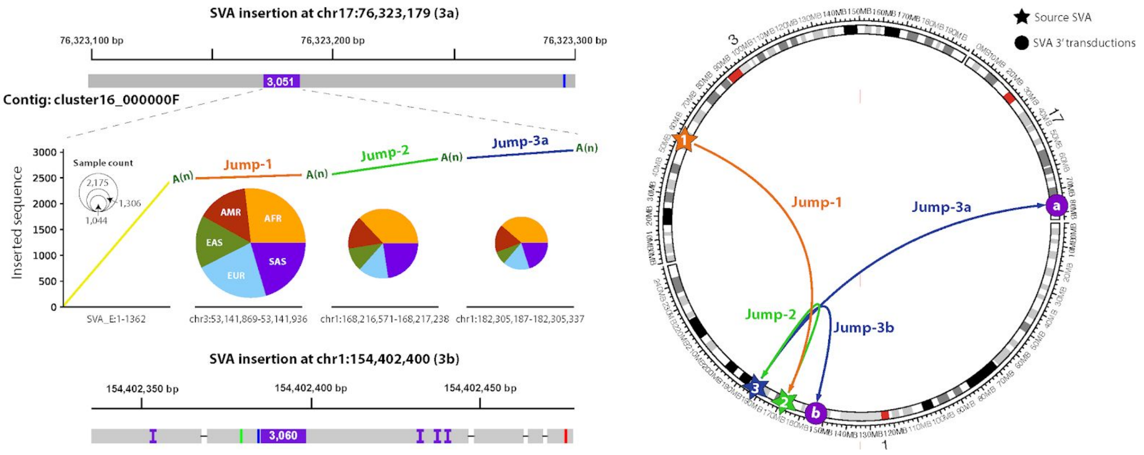


#### Fig. S39. Multi-transduction event disentangled via assemblies

Left, alignment to GRCh38 of contigs spanning two nested SVA-mediated transductions on 17q (Jump-3a) and 1p (Jump-3b). The configuration for the sequence-resolved nested transduction at 17q is depicted under the alignment as a line plot. Alignment of the insert over the SVA_E consensus and three genomic loci at 3p, 1p (chr1:168216956) and 1p (chr1:182305187) are colored in yellow, orange, green, and blue, respectively. Poly(A) stretches between each transduction event are shown in dark green. Population distribution together with the average number of 1KG samples where k-mers derived from each transduced sequence are displayed as pie charts of different sizes. Right, circos plot showing the predicted sequence of consecutive SVA transduction events leading to the generation of 17q (Jump-3a) and 1p (Jump-3b) insertions.

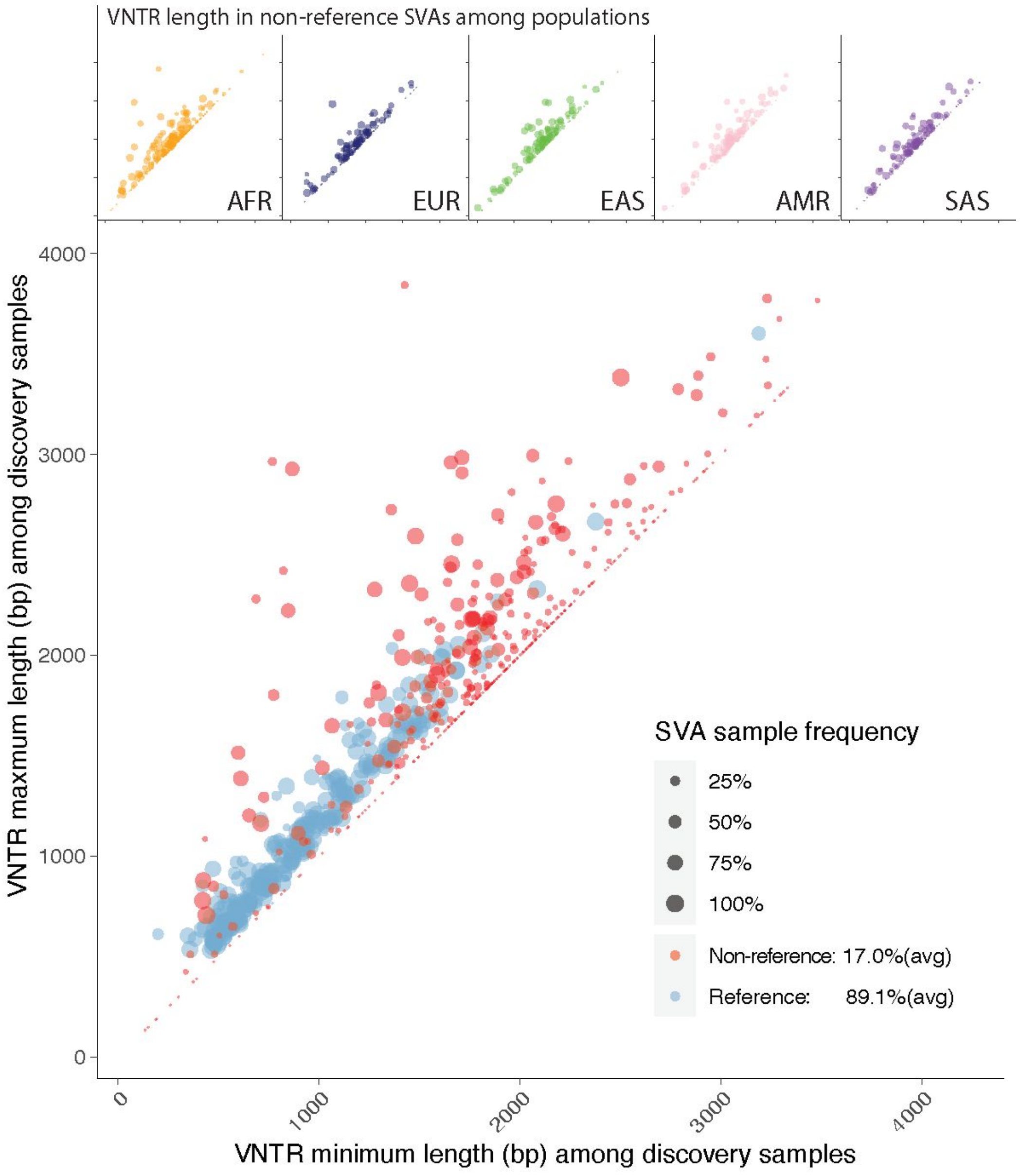


**Fig. S40. Distributions of VNTR length in reference and polymorphic SVAs in each individual discovery sample**

The main panel shows the minimum length (x-axis) versus maximum (y-axis) of VNTR regions in each SVA event (reference one as blue dot and polymorphic one as red dot). The size of dot represents the sample frequency of SVAs among discovery samples in HGSVC. The upper panel shows the separate distributions of VNTR regions in SVAs for five superpopulations: AFR (African, yellow), EUR (European, dark blue), EAS (East Asian, green), AMR (Amerindian, pink), and SAS (Southeast Asian, purple).

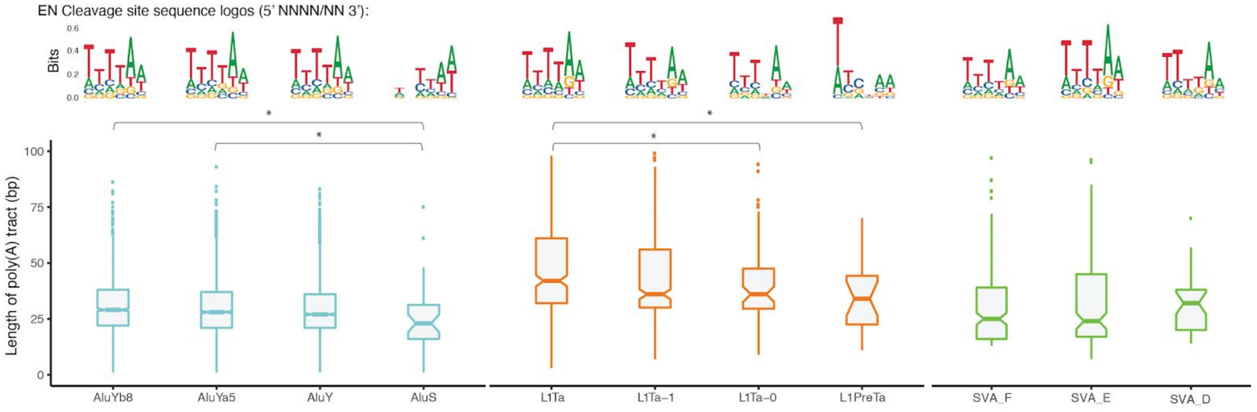


**Fig. S41. Box plots of poly(A) tract and sequence logos of EN cleavage sites for MEI subfamilies**

The lower panel shows the length distributions for the polymorphic MEI families and their main subfamilies in the human populations: Alu (AluYb8, AluYa5, AluY, and AluS, blue), L1Hs (L1Ta, L1Ta-1, L1Ta-0, and L1PreTa, orange), and SVA (SVA_F, SVA_E, and SVA_D, green). The significant difference for poly(A) tract length between the subfamilies is indicated by an asterisk (p-value < 0.05). The upper panel shows the EN cleavage site sequence logos for the corresponding subfamilies in the x-axis.

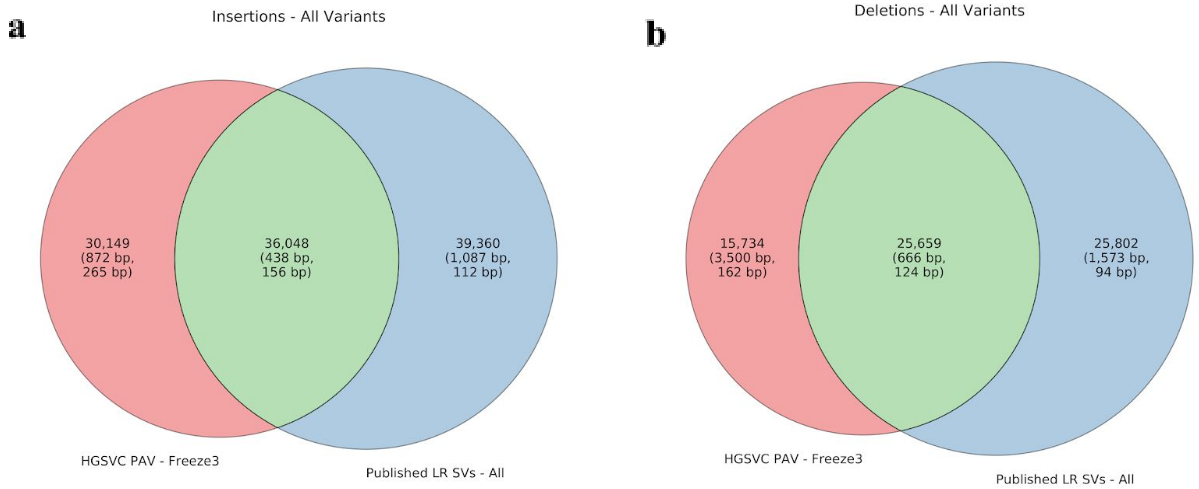


**Fig. S42. New sequence-resolved SVs**

Merged SVs in red are compared to SVs published in five other long-read studies in blue.

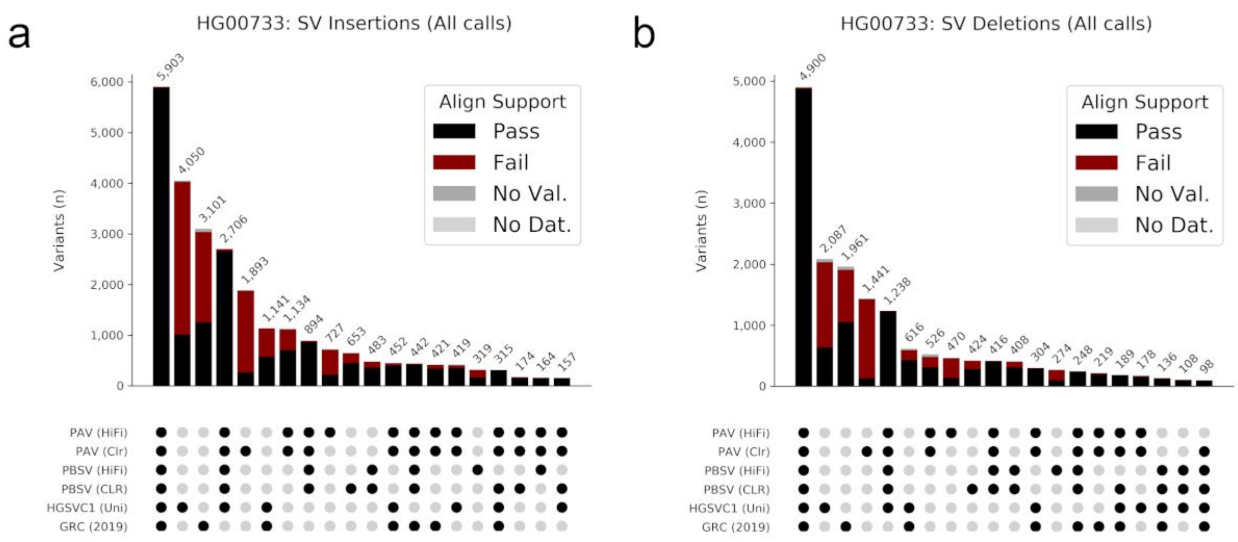


**Fig. S43. PAV concordance among callers for HG00733**

Calls from multiple sources for HG00733 were merged with the three-step approach. Shown are PAV with minimap2 alignments (PAV), PBSV, Chaisson 2019 unified callset (Chaisson et al. 2019), and Audano 2019 (Audano et al. 2019) for (a) SV insertions and (b) SV deletions. Bars are colored by subseq support. Variants supported by more than one caller also tend to have more subseq support. PAV calls from CLR contain more apparent false positives.

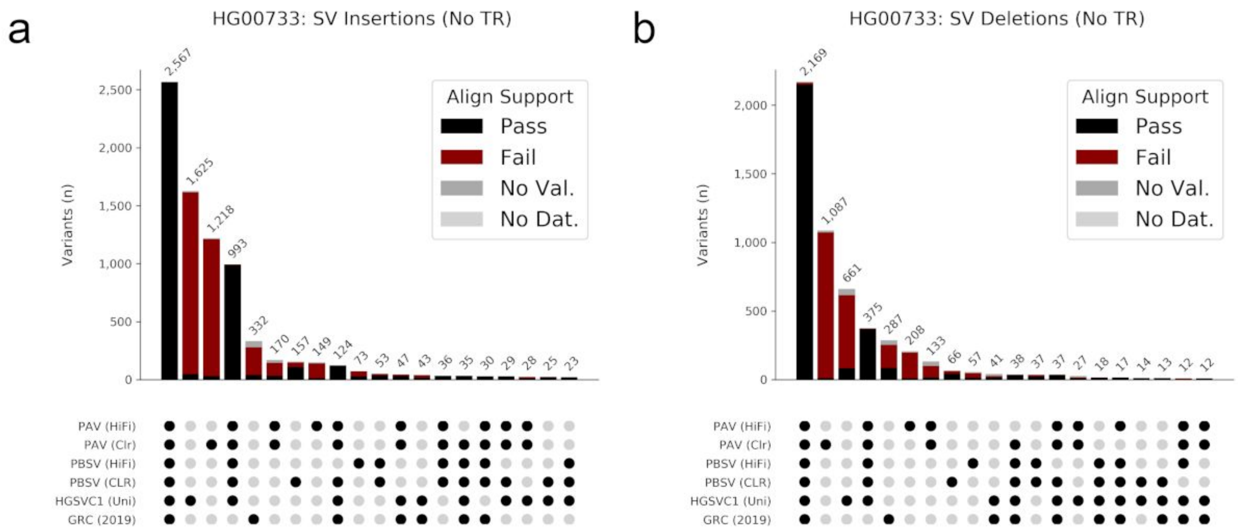


**Fig. S44. PAV concordance among callers for HG00733 outside tandem repeats**

Calls outside annotated tandem repeats from multiple sources for HG00733 were merged with the three-step approach. Shown are PAV with minimap2 alignments (PAV), PBSV, Chaisson

2019 unified callset (Chaisson et al. 2019) , and Audano 2019 (Audano et al. 2019) for (a) SV insertions and (b) SV deletions. Bars are colored by subseq support.

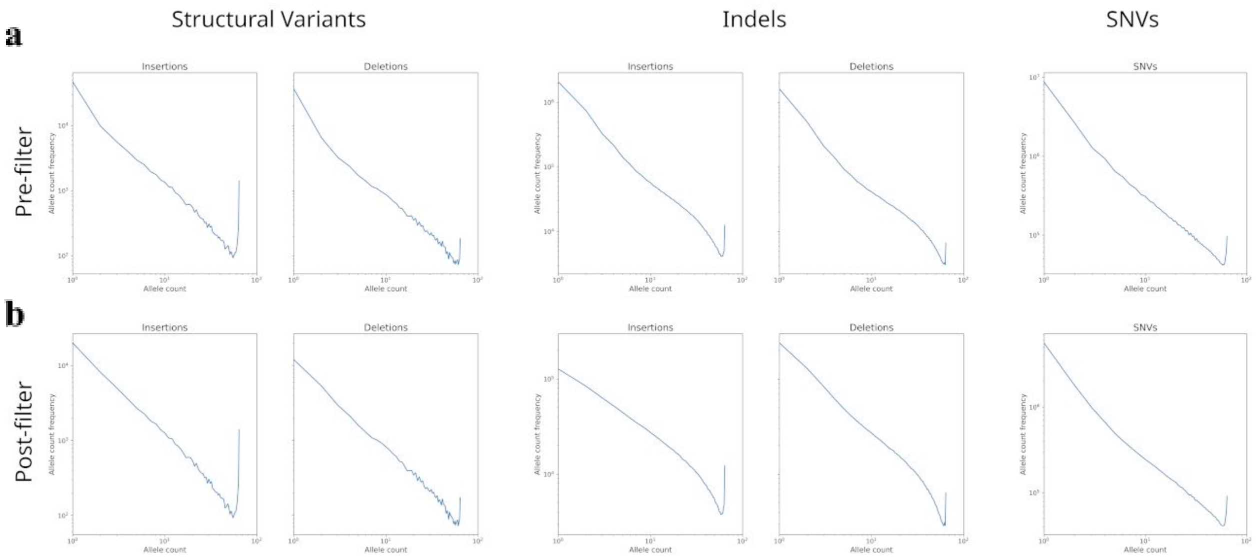


**Fig. S45. Excess singleton rate is reduced by callset filtering**

When allele counts (horizontal axis) are plotted against the frequency for each count (vertical axis), the distribution should be approximately linear. Before filtering (**a**), an increase in singletons was observed, which was corrected by applying “+1” (SVs) and “+2” (indels and SNVs) filters to the callset (**b**).

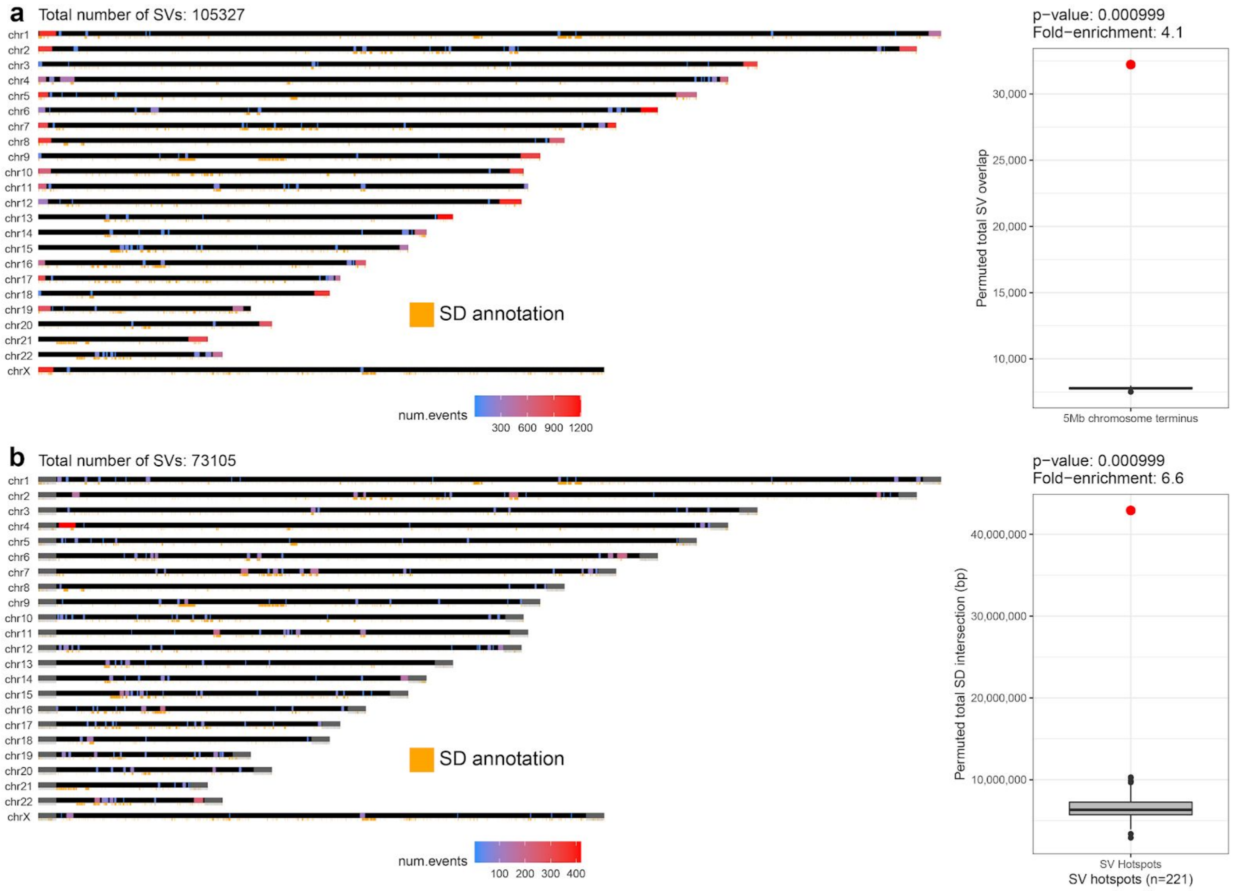


**Fig. S46. SV hotspot detection and enrichment for chromosome ends and SDs**

a) An ideogram showing detected SV hotspots using all SVs (≥50 bp). The total number of SVs in each detected hotspot is shown by a scale going from blue to red. Positions of SDs are added at the bottom of each chromosome ideogram as an orange rectangle. Left: Enrichment analysis of SVs with respect to the last 5 Mbp of each chromosome end. The red dot represents the observed number of SVs at the 5 Mbp terminus while the box plot shows the distribution of SV counts at the 5 Mbp terminus, after 1000 random shuffling of SVs (Methods).

b) An ideogram showing detected SV hotspots using after filtering SV at the 5 Mbp terminus of each chromosome (highlighted by a gray rectangle at the end of each chromosome). The total number of SVs in each detected hotspot is shown by a scale going from blue to red. Positions of SDs are added at the bottom of each chromosome ideogram as an orange rectangle. Left: Enrichment analysis of SVs in respect to SD content. The red dot represents the observed number of SD bases intersecting with detected hotspots while the box plot shows the distribution of SD overlap after 1000 random shuffling of detected hotspots (Methods).

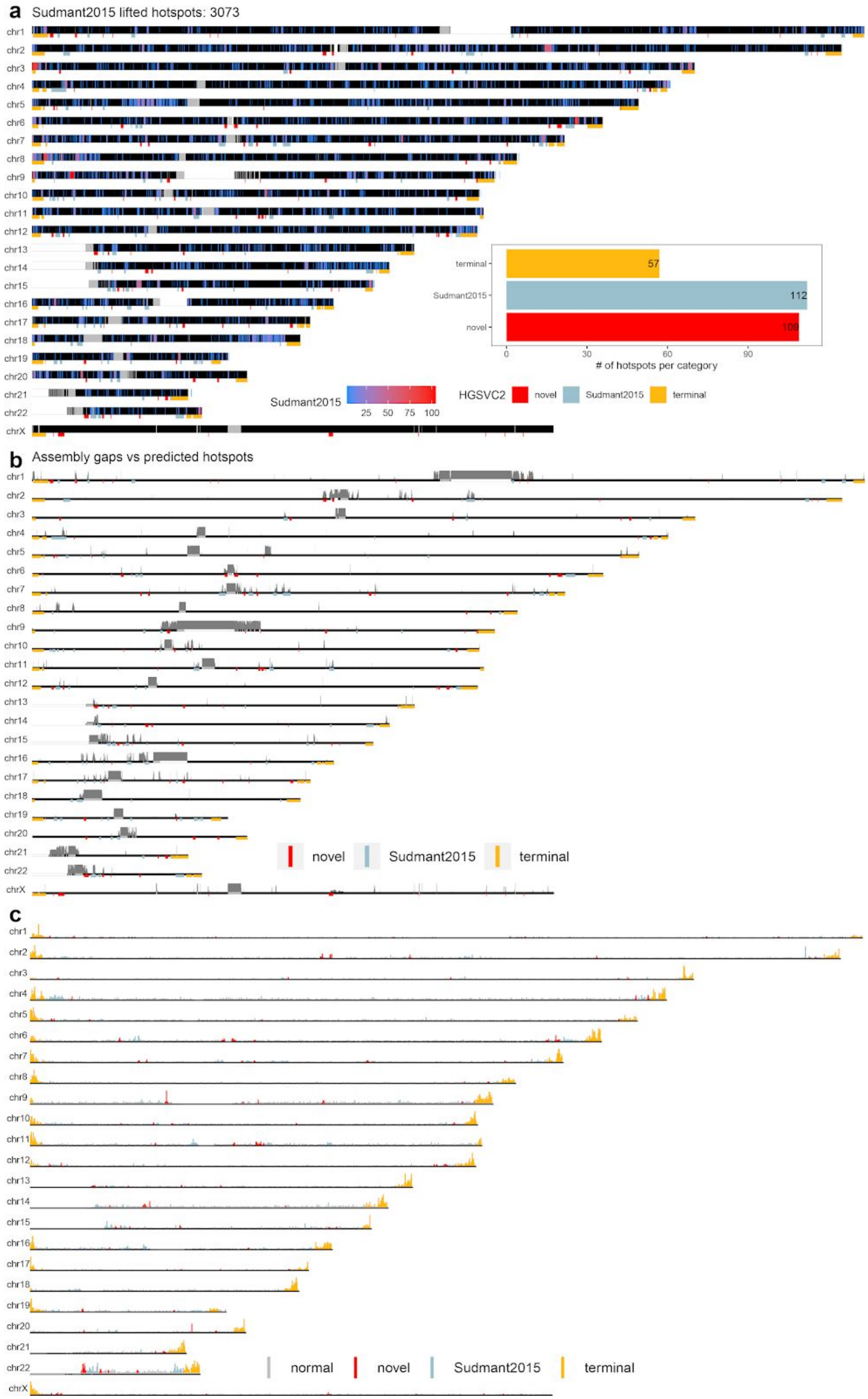
a
Sudmant2015 lifted hotspots: 3073
terminal
Sudmant2015
novel
57
112
109
0
30
60
90
### of hotspots per category
Sudmant2015
25 50 75 100
HGSVC2
novel
Sudmant2015
terminal
b
Assembly gaps vs predicted hotspots
novel
Sudmant2015
terminal
c
normal
novel
Sudmant2015
terminal

**Fig. S47. Raw data supporting detected SV hotspots**

a) Genome-wide distribution of SV hotspots detected by Sudmant et al. (*46*) (blue->red heatmap). SV hotspots are categorized into three groups: 'novel' - unique for this study, 'Sudmant2015' - overlapping with previous study, and 'terminal' - residing in the last 5 Mbp of each chromosome end. Inset: Shows a total count of hotpots in each previously defined category.
b) Gray bar along each ideogram shows assembly gaps, across all 64 assembled phased genomes, as regions where contigs map with <60 mapping quality. Underneath each ideogram, we plot a position of each hotspot as a rectangle colored by previously defined hotspots category.
c) An ideogram showing the binned counts (200 kbp bin) of SVs along each chromosome. Each bin is colored based on the overlap with the previously defined hotspot category. Bins not overlapping any hotspot are shown in gray ('normal').

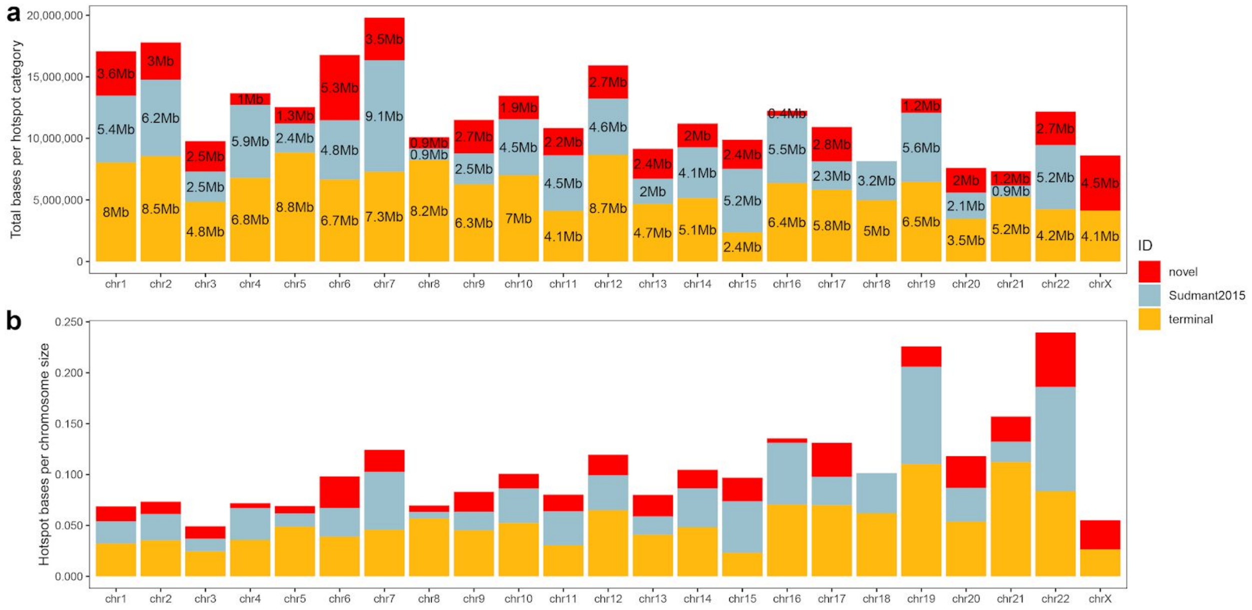


**Fig. S48. Summary of detected SV hotspots per chromosome**

a) A total size of detected hotspots per category ('novel', 'Sudmant2015' and 'terminal') and per chromosome (x-axis).

b) Number of hotspot bases per category and per chromosome normalized by the chromosome size.

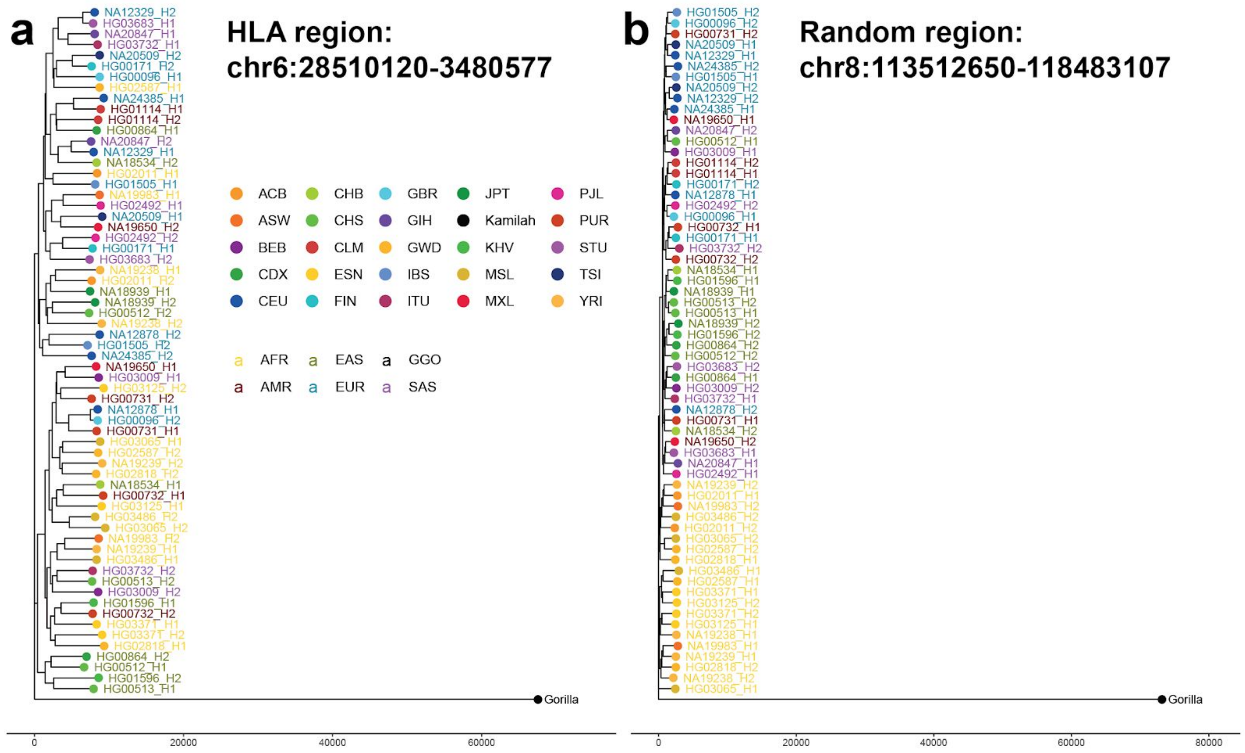


**Fig. S49. Evolutionary distances inside and outside of the HLA region**

a) A neighbor-joining evolutionary tree based on distances calculated between all phased assemblies over the HLA region (chr6:28510120-33480577) and b) Randomly chosen region (chr8:113512650-118483107). Each dot is colored based on the 1KG population identifier and color of the label defines superpopulation. We used a gorilla (GGO) squashed assembly (Kamilah individual) as an outgroup.

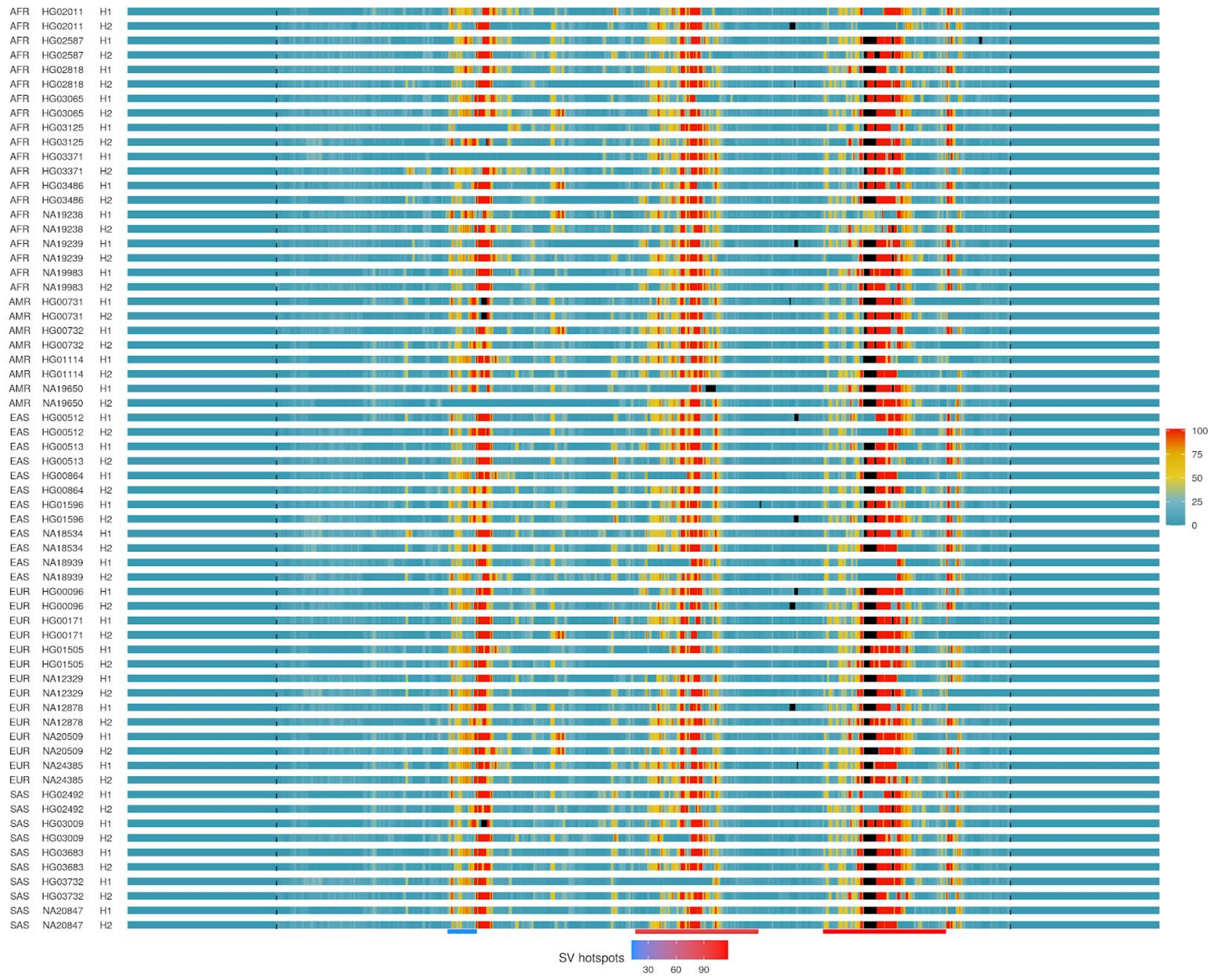


**Fig. S50. Count of alternative alleles per haplotype for the HLA region**

Each row represents binned counts (binsize – 10 kbp, stepsize – 1 kbp) of alternative alleles in each assembled haplotype with respect to GRCh38 as a reference. The number of alternative alleles in each bin are reflected in a color scheme going from blue to red (see legend). The HLA region (chr6:28510120-33480577) is highlighted by dashed lines. Regions of less reliable contig mapping with respect to GRCh38 are highlighted by black color. At the bottom of this heatmap, we show previously detected SV hotspots that overlap with the HLA region. The number of SVs in each hotspot is reflected by the color scheme going from blue to red.

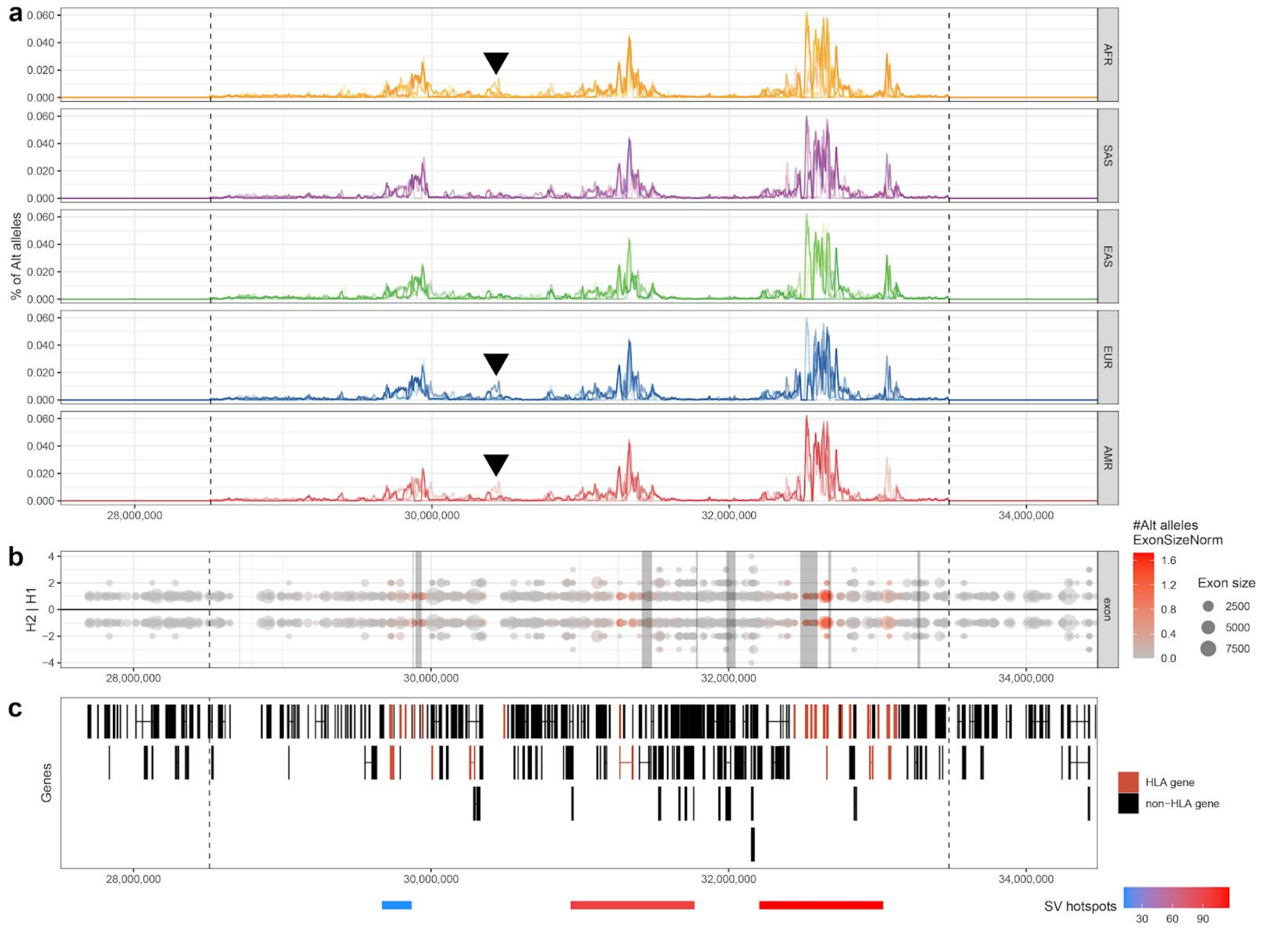


**Fig. S51. Summary of genetic variability over the HLA region**

a) Binned (binsize – 10 kbp, stepsize – 1 kbp) percentage of alternative alleles per haplotype. Haplotypes from each superpopulation (AFR - African, SAS - Southeast Asian, EAS - East Asian, EUR - European, AMR - Amerindian) are grouped together and plotted as a single line. Transparency (alpha) level serves to highlight differences within and across superpopulations. Black arrowheads point to the extent of haplotype diversity unique to AFR, EUR and AMR populations.

b) Number of alternative alleles per exon overlapping the HLA region. The size of each exon is reflected by the size of each dot. The number of alternative alleles per exon is highlighted by a color scheme going from gray to red.

c) A gene model obtained from R package 'TxDb.Hsapiens.UCSC.hg38.knownGene'. The position of each exon is plotted as a black or red rectangle for non-HLA and HLA genes, respectively. At the bottom, we show previously detected SV hotspots that overlap with the HLA region. The number of SVs in each hotspot is reflected by a color scheme going from blue to red.

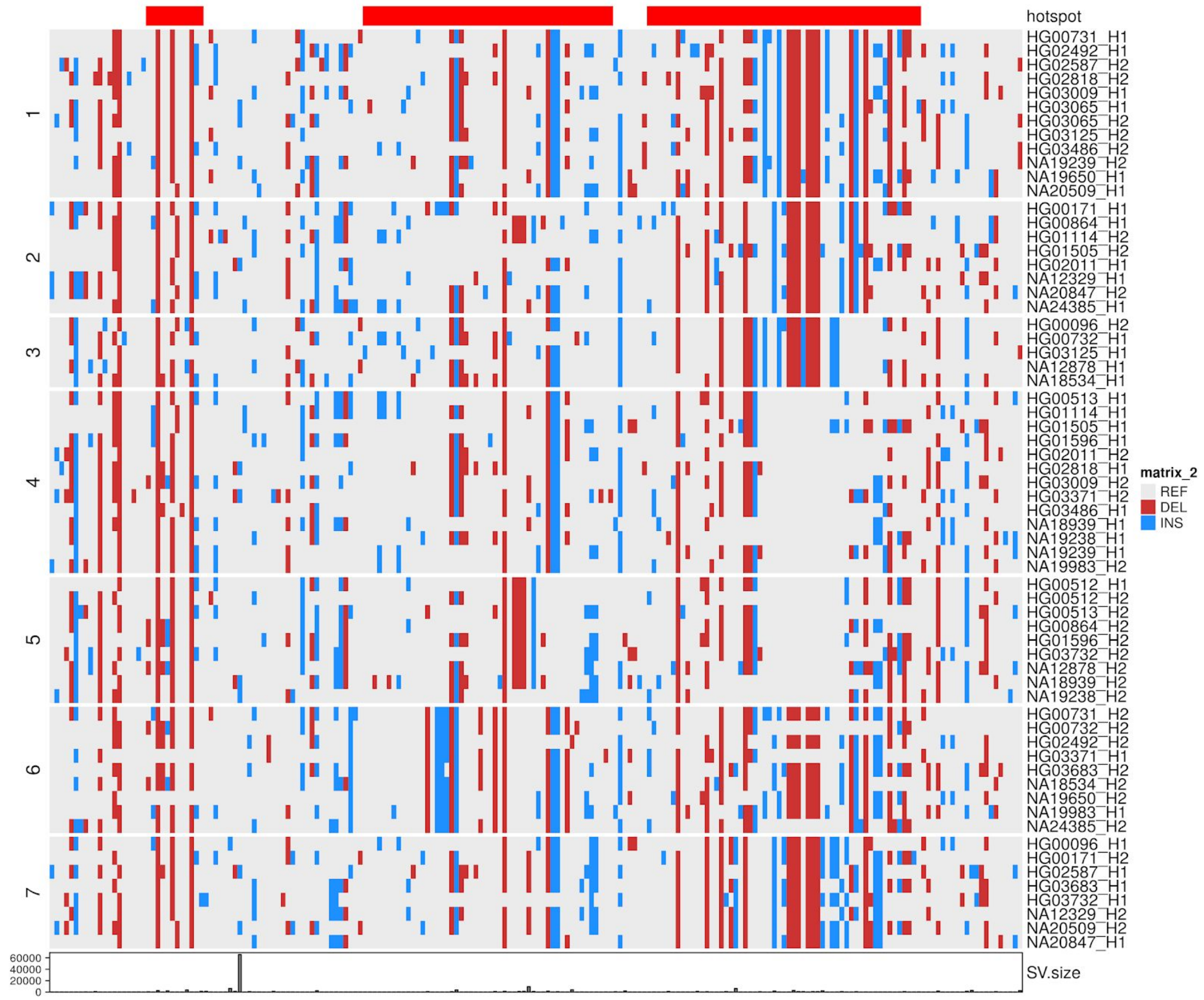


**Fig. S52. Definition of distinct HLA haplotypes based on SV clustering**

A heatmap of clustered SV that defines seven distinct haplotypes where each column represents an SV and each row represents a unique haplotype (n=64) assembled over the HLA region. Haplotype clusters have been defined by k-means clustering after removal of SVs from regions where assemblies map with lower confidence to GRCh38. Final number of clusters was decided based on manual curation and are highlighted by separation of row by white space and the cluster number (n=7). At the top of the heatmap, we point to the SVs that overlap with predicted SV hotspots. At the bottom of the heatmap, we show the size of each SV as a barplot.

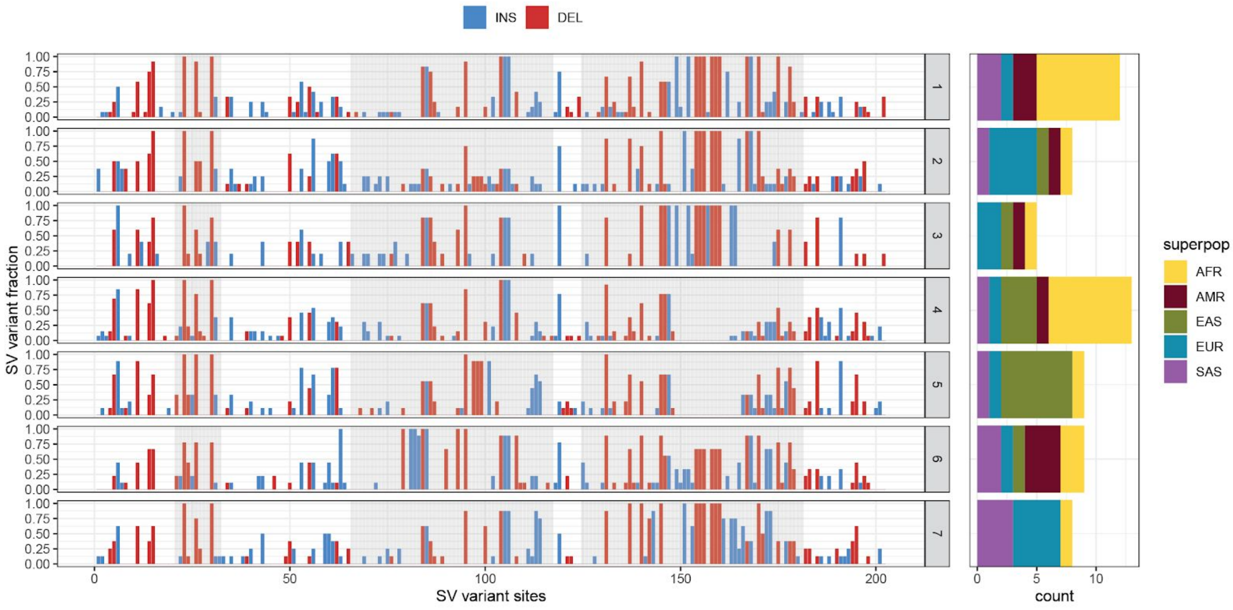


**Fig. S53. Summary of HLA haplotypes with a similar SV distribution**

Left plot shows fraction of SVs per site for each cluster defined by k-means clustering in Fig. S52. Height of each bar represents frequency of a given variant and the given position across all SV haplotypes assigned to each cluster (n=7). Regions of previously defined SV hotspots are highlighted by vertical gray bars over each SV that overlaps with a hotspot region. Barplot on the right counts the superpopulation of origin for each haplotype in each cluster.

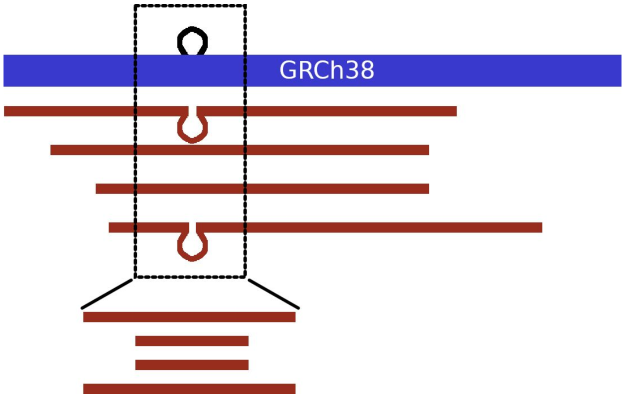


**Fig. S54. Subseq illustration**

Reads (red) aligned to the reference (blue) with an SV insertion (loop). A region surrounding the breakpoints is chosen (dashed box) with larger regions for larger SVs. All reads traversing that region (i.e. touching both ends) are extracted, and the length of the read in that region is determined by parsing the CIGAR string. In this example, regions extracted from two reads (top and middle) support a heterozygous insertion, and two reads (middle two) do not support it.

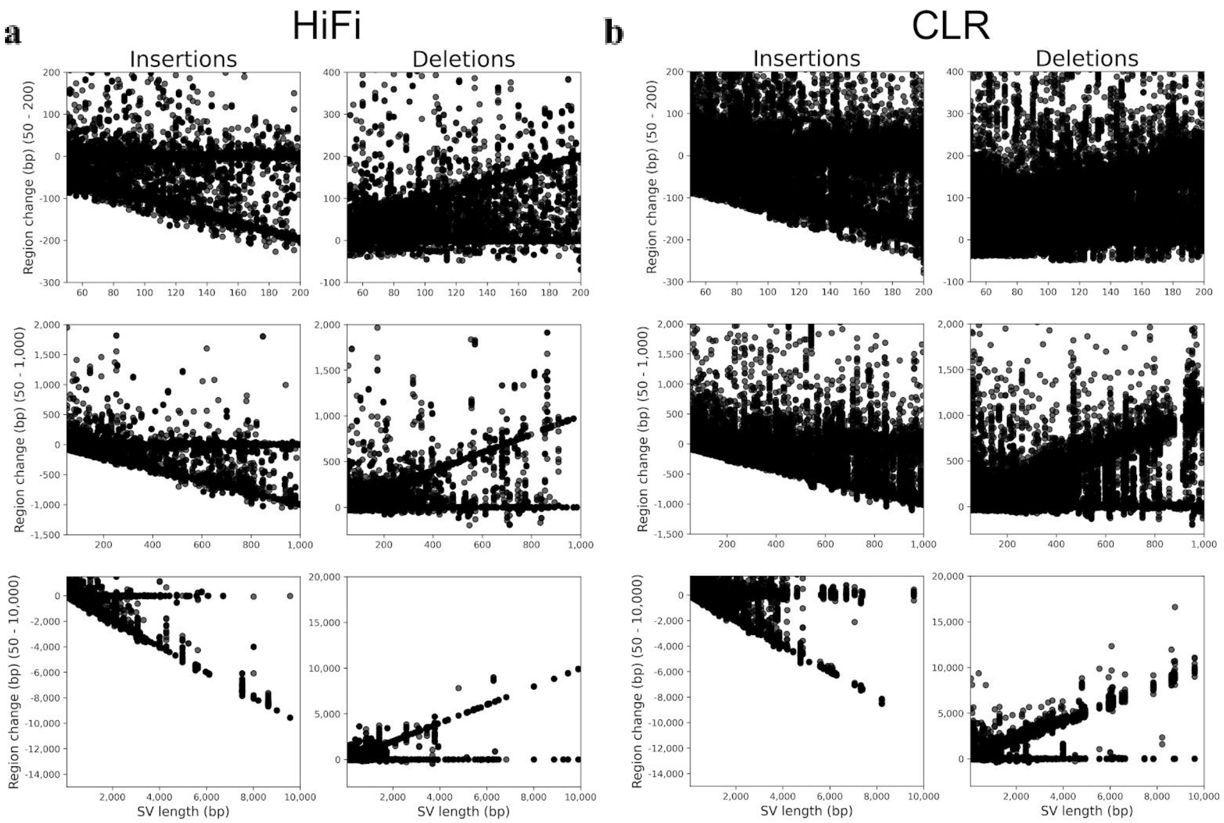


**Fig. S55. Subseq region lengths deviate with SV length with less variation from HiFi**

A region around each SV was extracted from raw read alignments. For any read spanning the region, the size of the region within the read was found and the expected deviation was computed assuming the SV was present. If the SV is supported, then the expected deviation must be within 50% of the SV length (±0.5 * SVLEN). We ran validations for HiFi (**a**) and CLR (**b**) for all samples (HG00733 shown). Each point in the figure is one read alignment with the SV length (x-axis) and the size deviation around the SV for a single read (y-axis). Reads supporting an SV call cluster along y=0. Reads not supporting the variant cluster along -SVLEN (insertions, alignment is shorter than expected if SV was present) and +SVLEN (deletions, alignment is longer than expected if SV was present). Figure shows SVs 50-200 bp (top), 50-1,000 bp (middle), and 50-10,000 bp (bottom). Each panel shows a random sample of 2,000 SV calls.

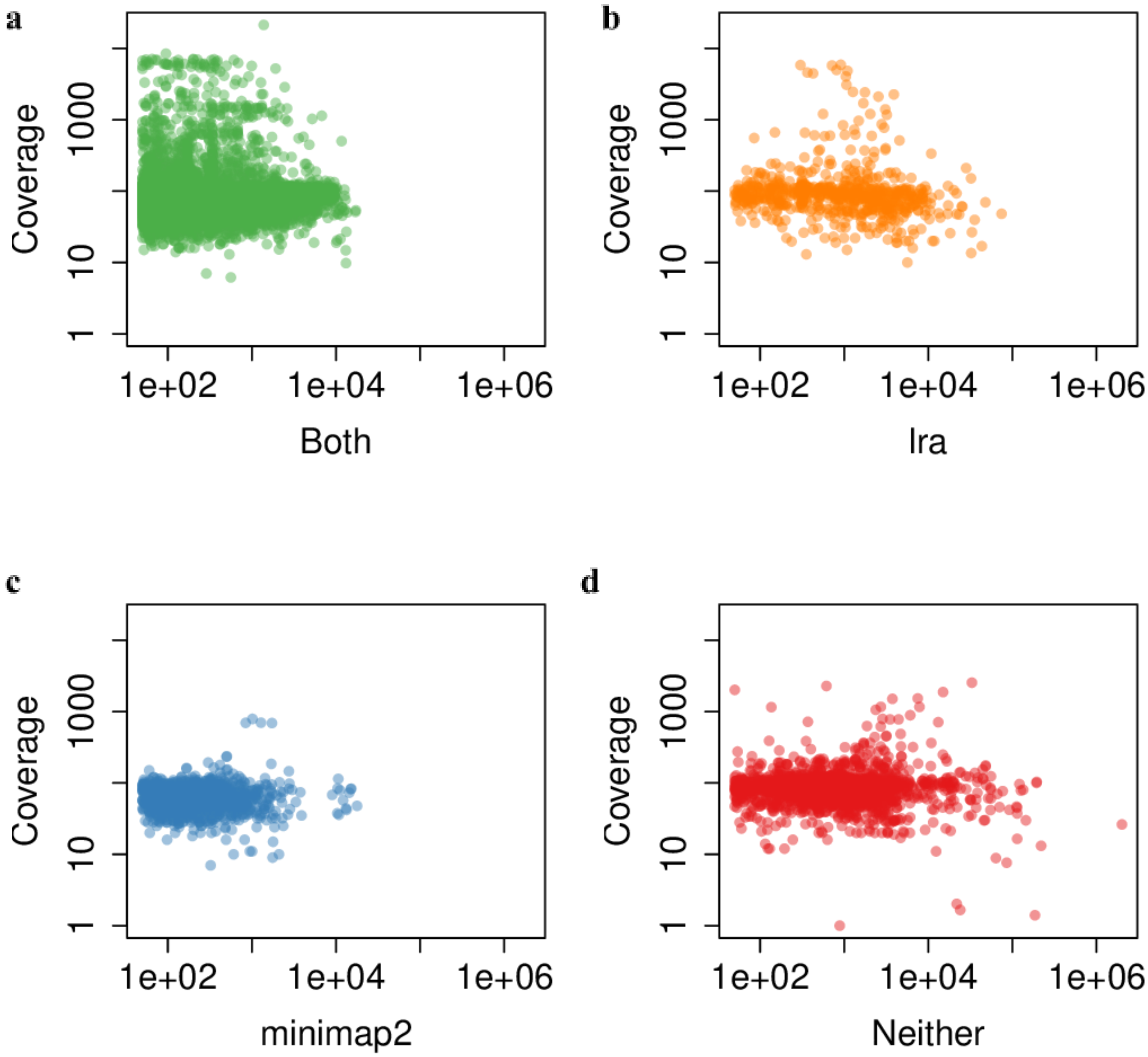


**Fig. S56. SV support from raw reads**

SV support from alignment of reads by LRA and minimap2, shown by length of read versus read support for SV calls made from a CLR assembly of the HG00733 genome. **a**) calls with both LRA and minimap2 support. **b**) calls with only LRA support. **c**) calls with only minimap2 support. **d**) calls supported by neither method.

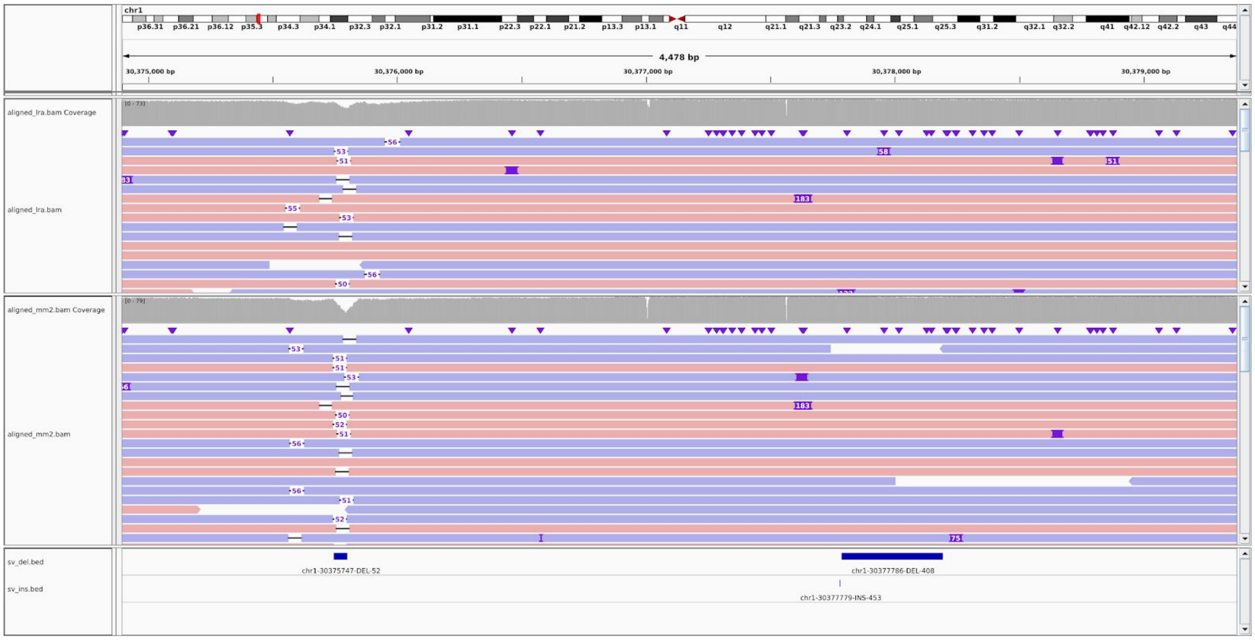


**Fig. S57. Example of SV calls with and without support**

Two deletion variant calls in HG00733 are shown in the region chr1:30,374,895-30,379,391 (bottom track). The left side call is a 52 base-pair deletion that has 66 alignments by minimap2, and 57 alignments by LRA supporting the call. For clarity, HiFi alignments are shown rather than CLR. The right side call is a 408 base-pair deletion that has no alignments from either LRA nor minimap2 supporting the call.

**a**

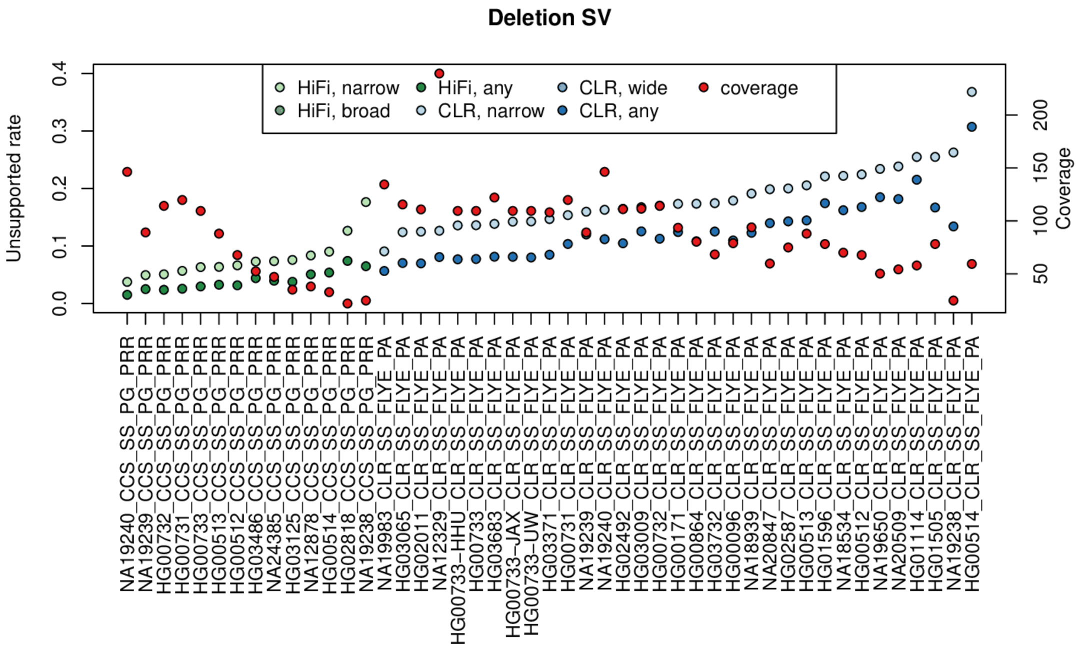


**b**

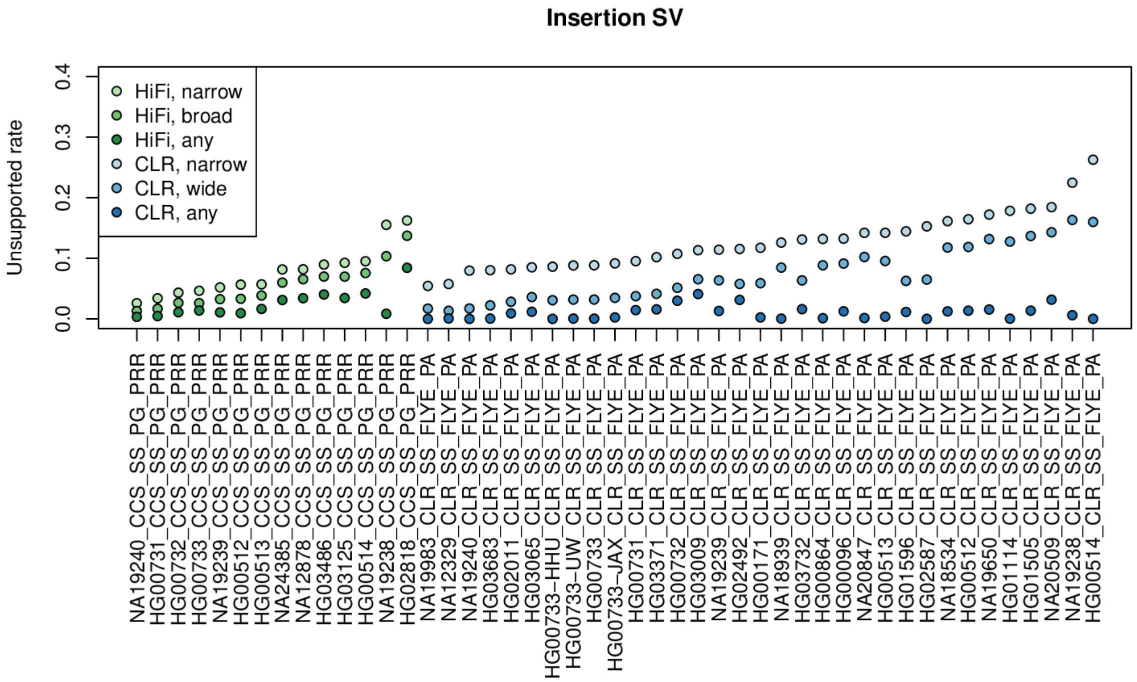


**Fig. S58. Fraction of unsupported SVs per genome**

**a**). Fraction of unsupported SVs for CCS (green) and CLR (blue) assemblies. Three levels of stringency for reads supporting an SV are calculated: narrow, wide, and any. Narrow support is defined as an SV in a read of the same type within 1 kbp and of length within 50% of the length of the assembly-based SV. Wide support has the same size constraint but allows for 10 kbp distance of SV breakpoint. Any support is within 10 kbp, and any SV length (>50 bases). The latter category is not a reasonable approach to detect read-based support for SV assembly but represents the upper bound for which variants may be supported by reads. Red points indicate the coverage of reads aligned to the reference. The coverage correlates negatively with supported reads (HiFi $r^2$ = -0.739, p=0.002; CLR $r^2$=-0.682, p=1.24E-5); however, this is likely a combined effect of assembly quality and lower number of reads that may support a variant.

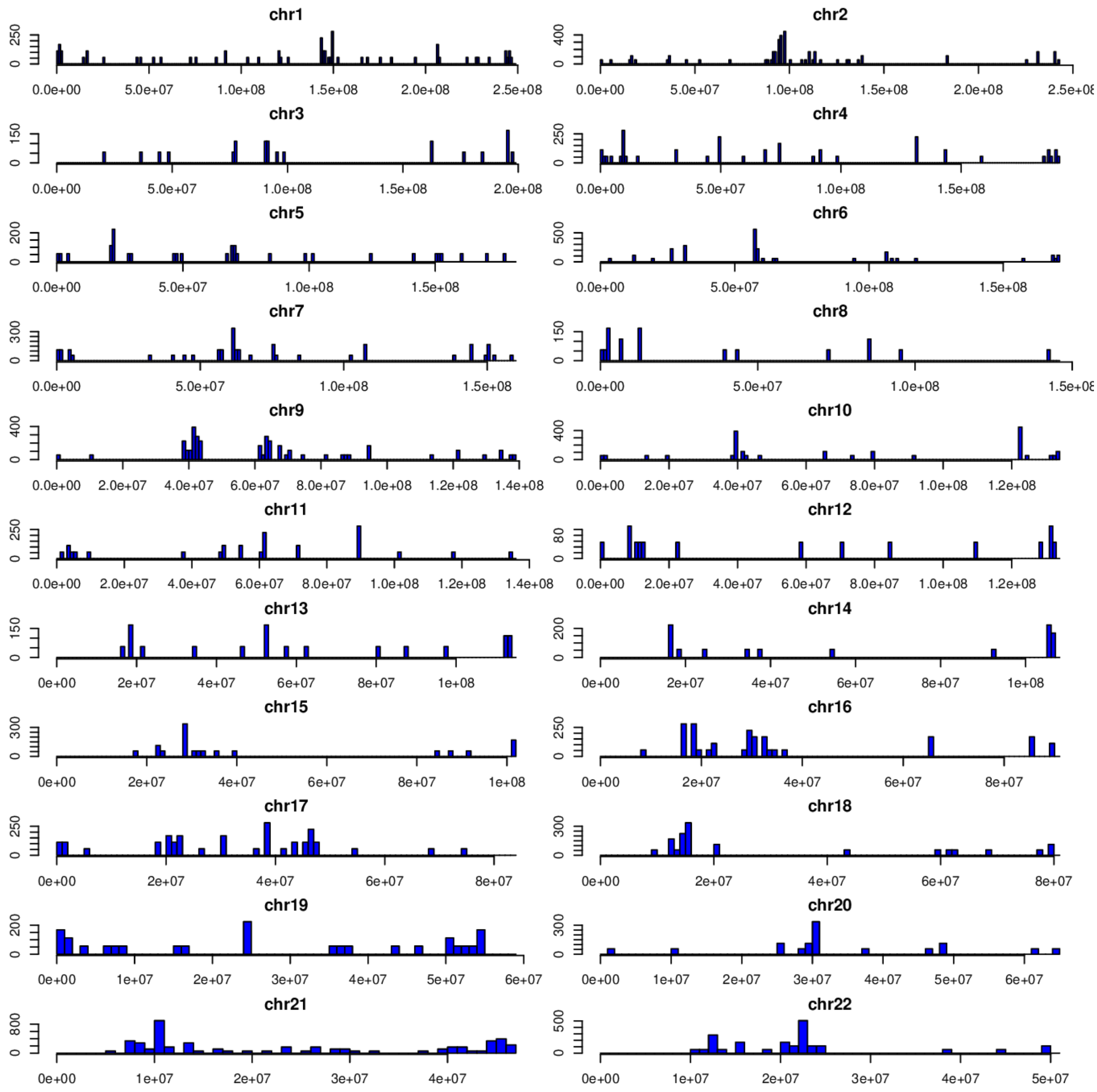


**Fig. S59. Genomic distribution of unsupported variants detected by HiFi assemblies**

Counts are aggregated over all assemblies and variant types for variants produced by PAV. Each bin represents the number of unsupported SVs per 1 Mbp per genome.

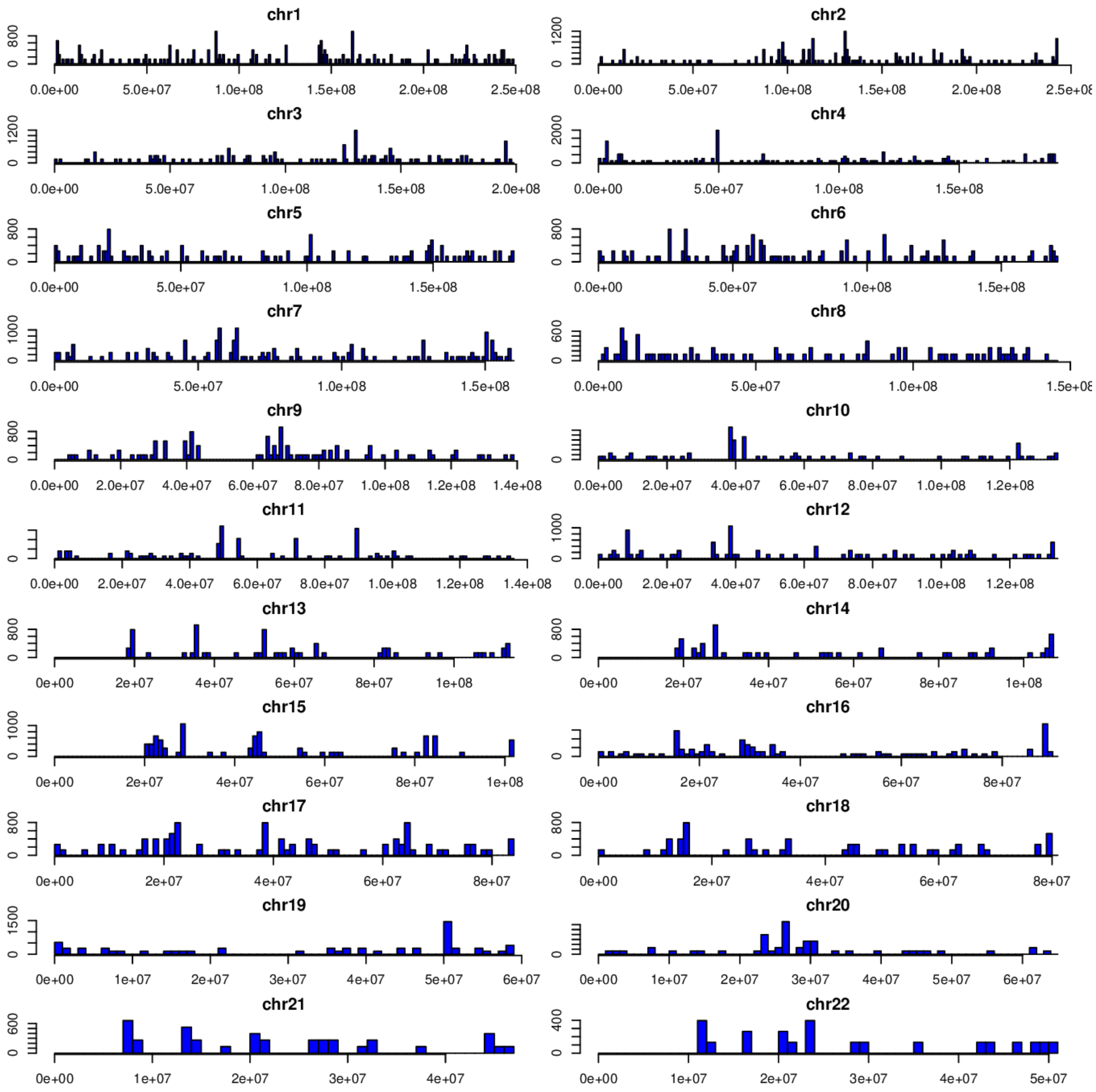


**Fig. S60. Genomic distribution of unsupported variants detected by CLR assemblies**

Counts are aggregated over all assemblies and variant types for variants produced by PAV. Each bin represents the number of unsupported SVs per 1 Mbp per genome.

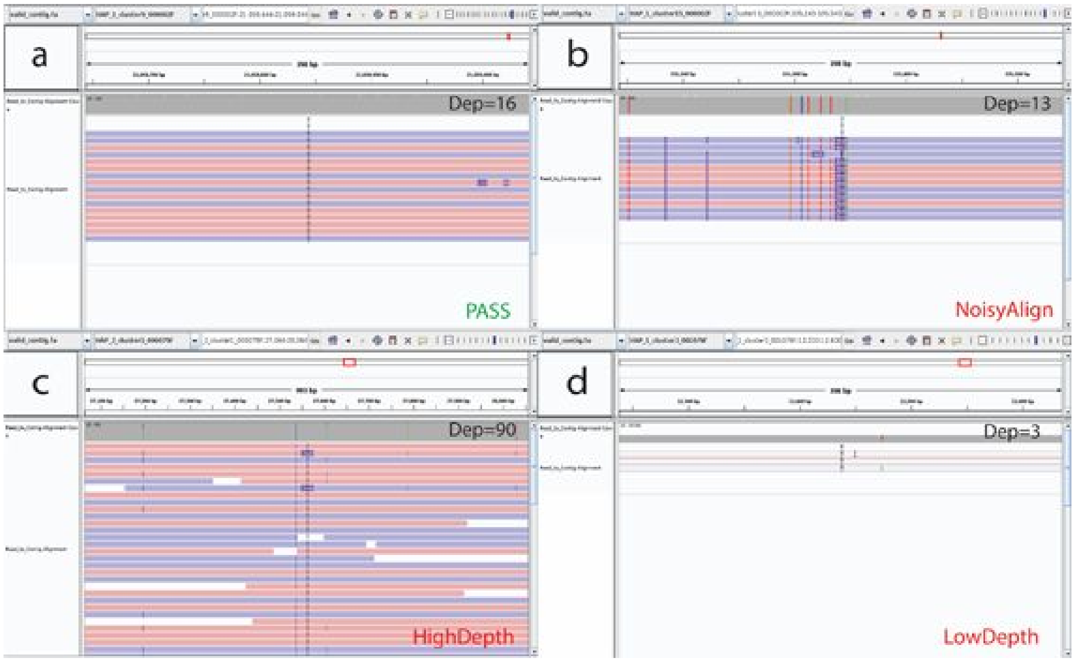


**Fig. S61. Examples of Inspector Quality Control**

Examples of read_to_contig alignment patterns for four types of SVs in HG00733 (HiFi) with mean alignment depth at ~16X. a) A “PASS” SV call with no indel or clipped alignment at its breakpoint; b) A “NoisyAlign” deletion call with 468 bp INS signals and several mismatches in read alignment; c) A “HighDepth” SV call with alignment depth of 90X; d) A “LowDepth” SV call with alignment depth of 3X at its breakpoint.

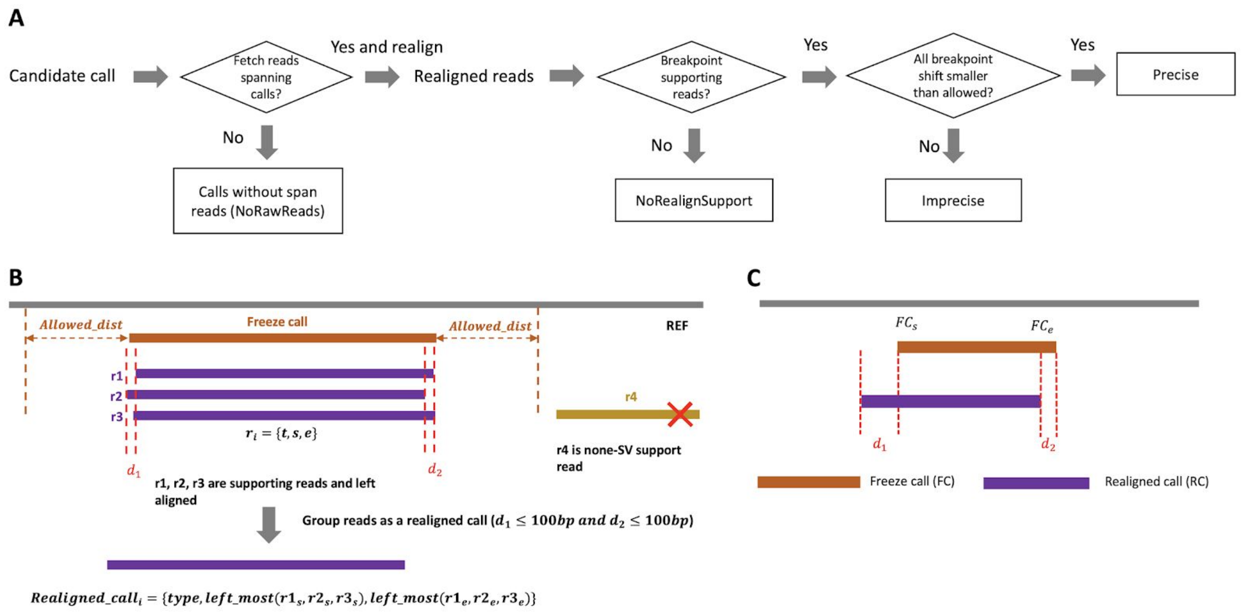


**Fig. S62. Workflow of SV QC**

a) The general workflow of SV QC, classifying the raw PAV calls into “NoRawReads”, “NoRealignSupport”, “Imprecise” and “Precise”; b) Creating realigned calls through MUMMer-based realignment. If realigned breakpoints from all SV spanning reads are outside of the allowed_dist (default is 500 bp), this SV is marked as “NoRealignSupport”; c) The diagram shows the process of labeling “Precise” and “Imprecise” calls. For a "Precise" PAV call, the breakpoint position difference between realigned call and this PAV call (d1 and d2) should be smaller than the maximum allowed breakpoint shift threshold. Otherwise, this PAV call is labeled as "Imprecise".

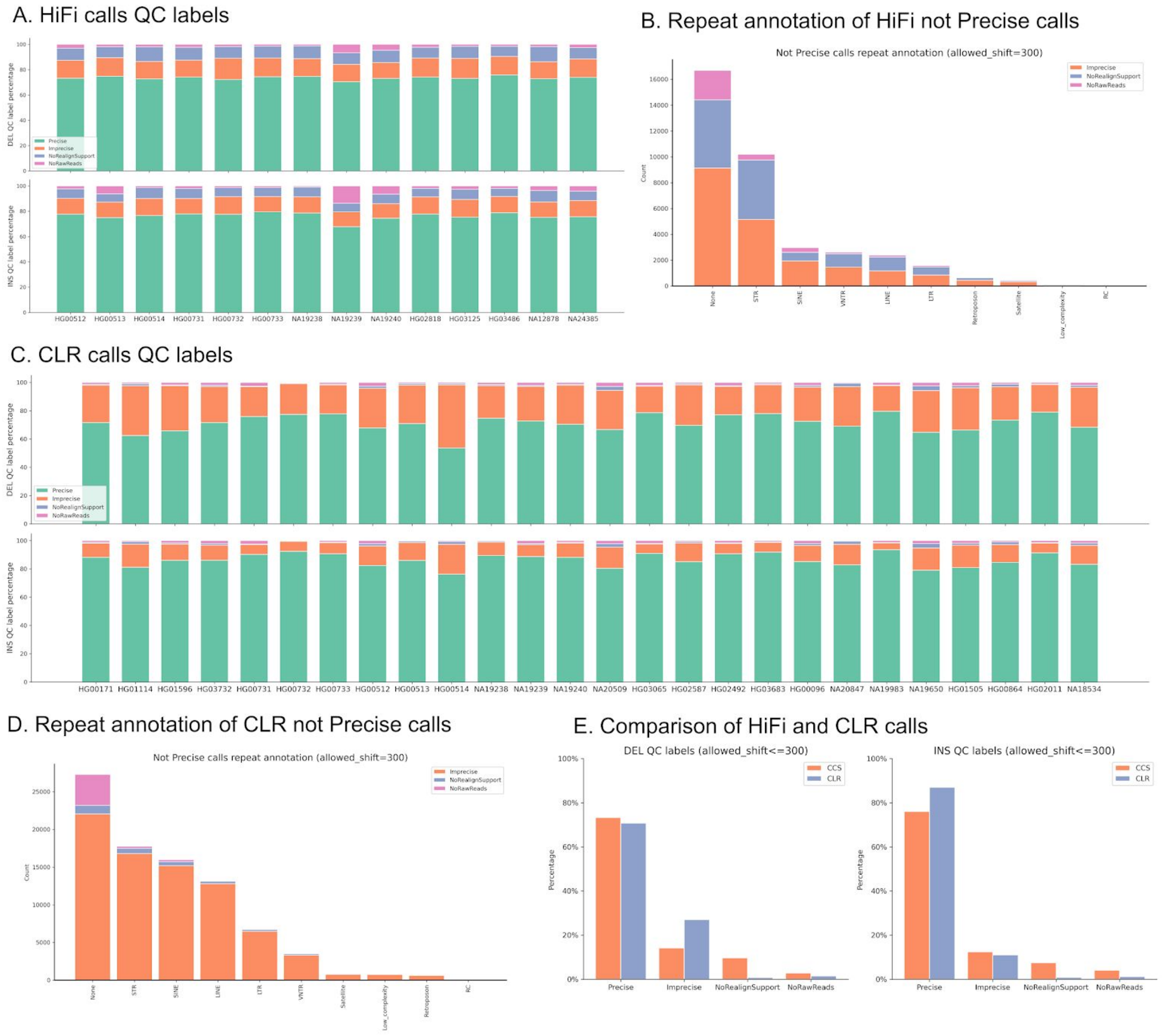


**Fig. S63. The overall QC results of HiFi and CLR samples**

a-b) QC labels of HiFi samples and their repeat annotation; c-d) QC labels of CLR samples and their repeat annotation; e) The comparison of HiFi samples and their replicated CLR samples.

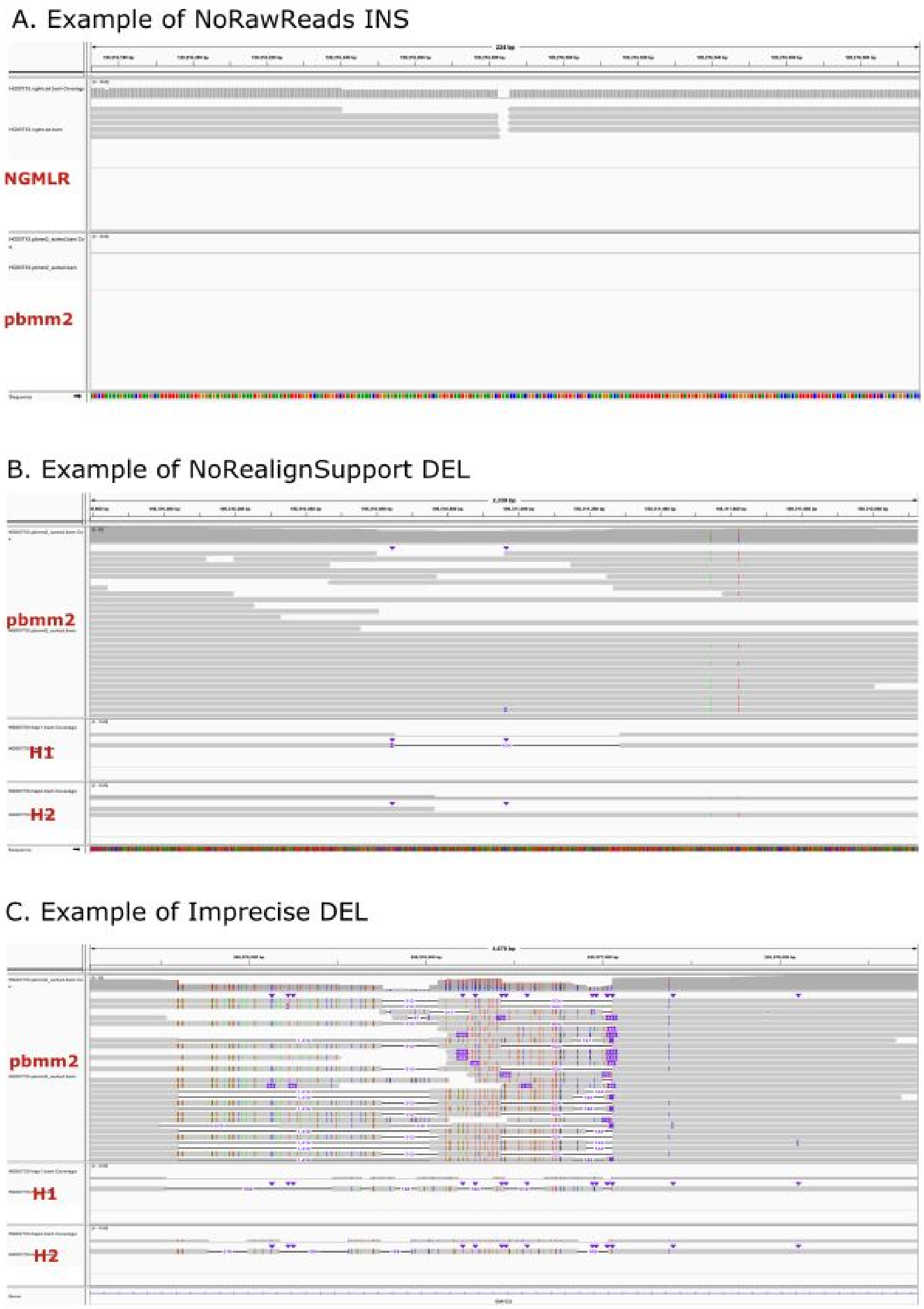


**Fig. S64. Examples of QC labels**

a) Example of “NoRawReads” INS call. Note that different aligners may have different results on the label; b-c) Examples of “NoRealignSupport” and “Imprecise” calls from sample HG00733.

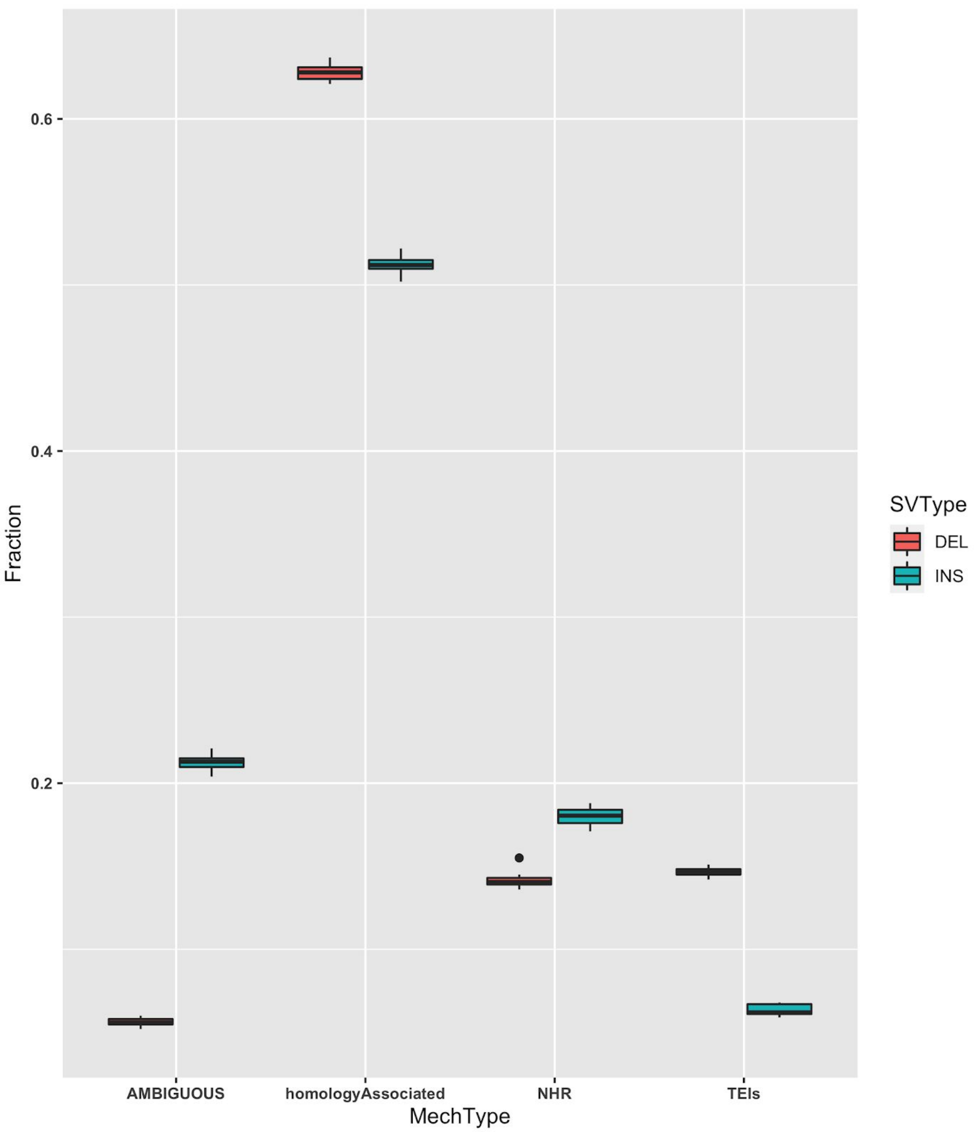


**Fig. S65. Distribution of insertions and deletions belonging to different mechanism categories**

Each data point in the box plot corresponds to the fraction of total SVs in a sample, which belong to a particular mechanism category. We performed this analysis separately for deletions (red box plot) and insertions (blue box plot).

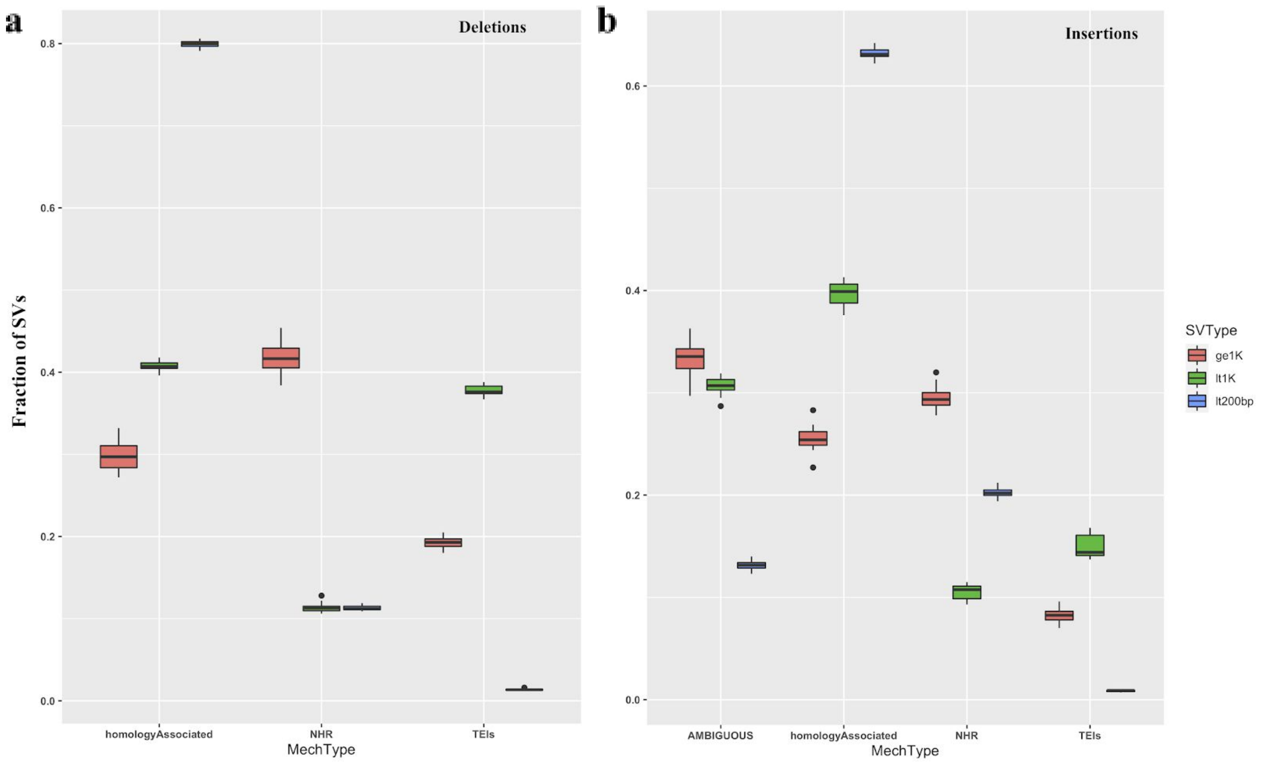


**Fig. S66. Distribution of insertions and deletions of different length groups belonging to different mechanism categories**

Each data point in the box plot corresponds to the fraction of total SVs in a sample, which belong to a particular mechanism category. We performed this analysis separately for deletions (**a**) and insertions (**b**). For each SV type, we divided SVs into three classes based on their length: i) length < 200 bp (blue), ii) 200 bp ≤ length < 1000 bp, and iii) length < 1000 bp.

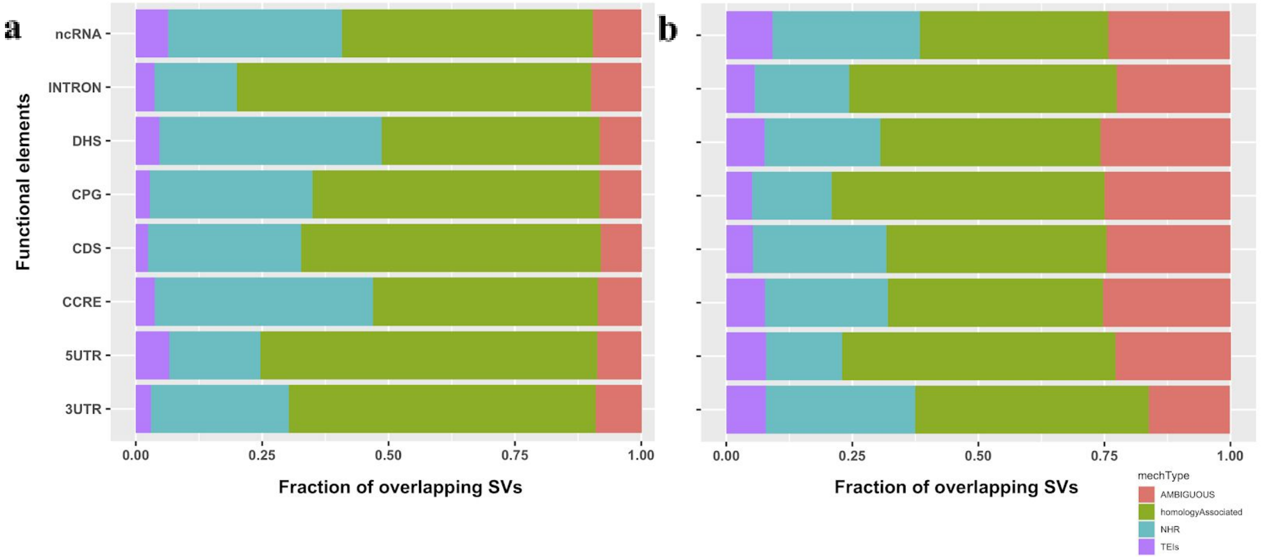


**Fig. S67. Distribution of insertions and deletions overlapping with distinct functional elements belonging to different mechanism categories**

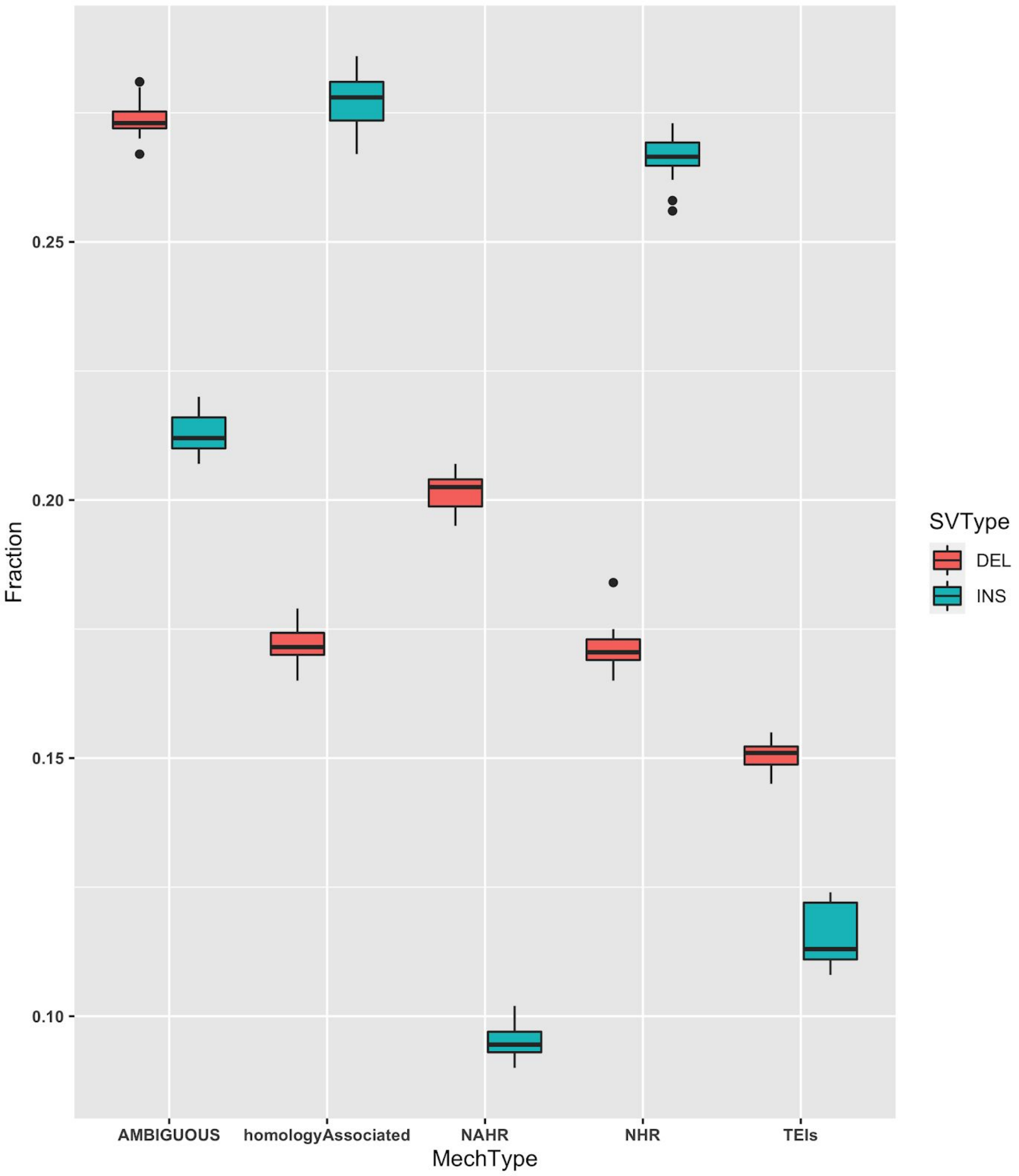


**Fig. S68. Distribution of insertions and deletions belonging to different mechanism categories (based on homology length ≥200 bp)**

Each data point in the box plot corresponds to the fraction of total SVs in a sample, which belong to a particular mechanism category. We performed this analysis separately for deletions (red box plot) and insertions (blue box plot).

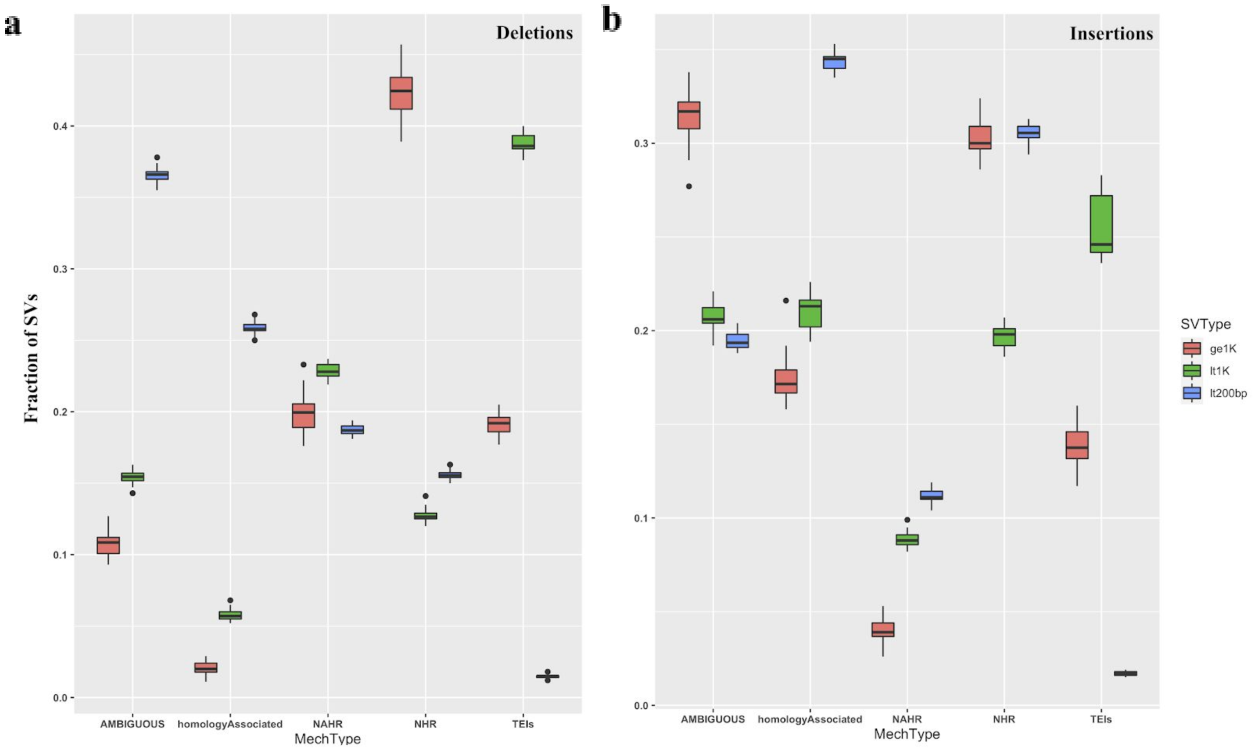


**Fig. S69. Distribution of insertions and deletions of different length groups belonging to different mechanism categories (based on homology length ≥200 bp)**

Each data point in the box plot corresponds to the fraction of total SVs in a sample, which belong to a particular mechanism category. We performed this analysis separately for deletions (**a**) and insertions (**b**). For each SV type, we divided SVs into three classes based on their length: i) length < 200 bp (blue), ii) 200 bp ≤ length < 1000 bp, and iii) length < 1000 bp.

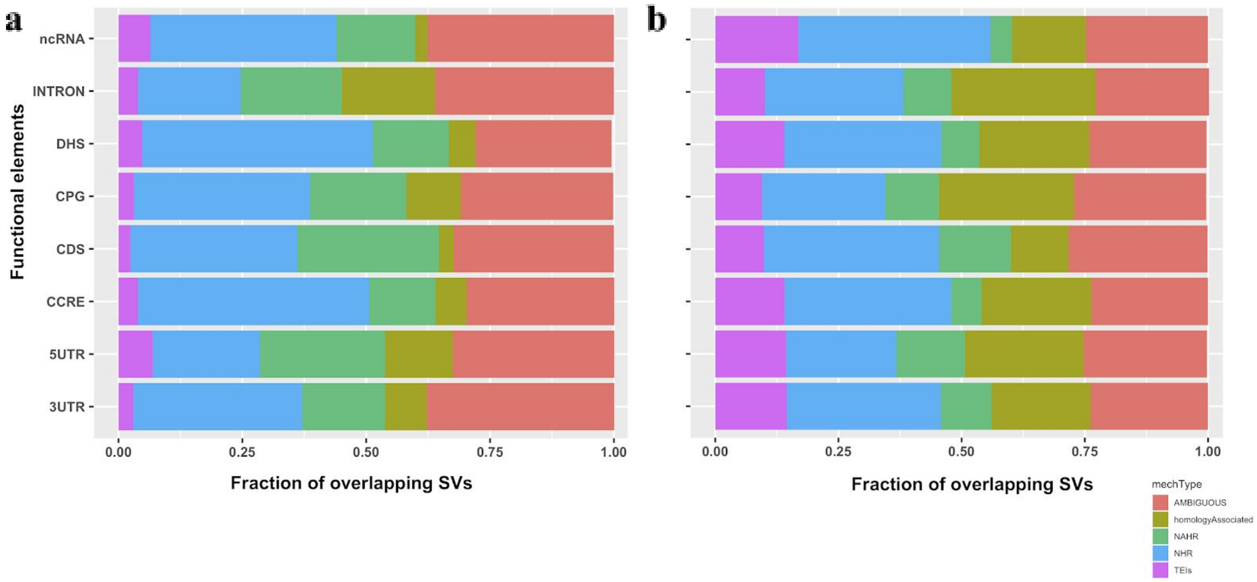


**Fig. S70. Distribution of insertions and deletions overlapping with distinct functional elements belonging to different mechanism categories based on homology length ≥200 bp**

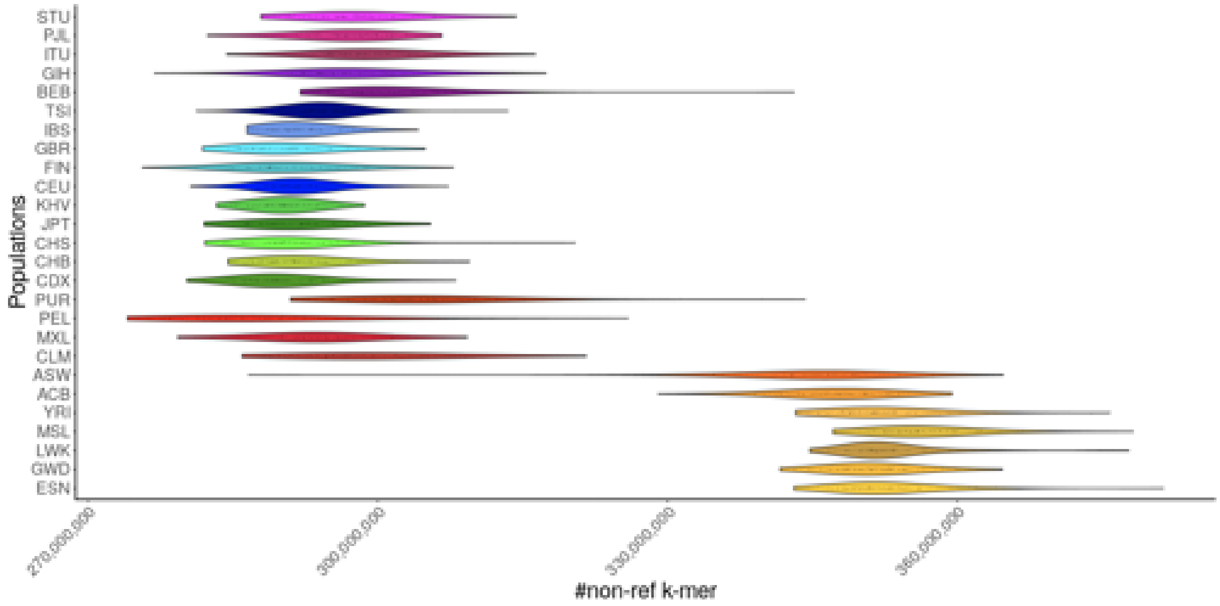


**Fig. S71. Non-reference k-mer density distribution (violin plot) by population for 2,504 unrelated samples**

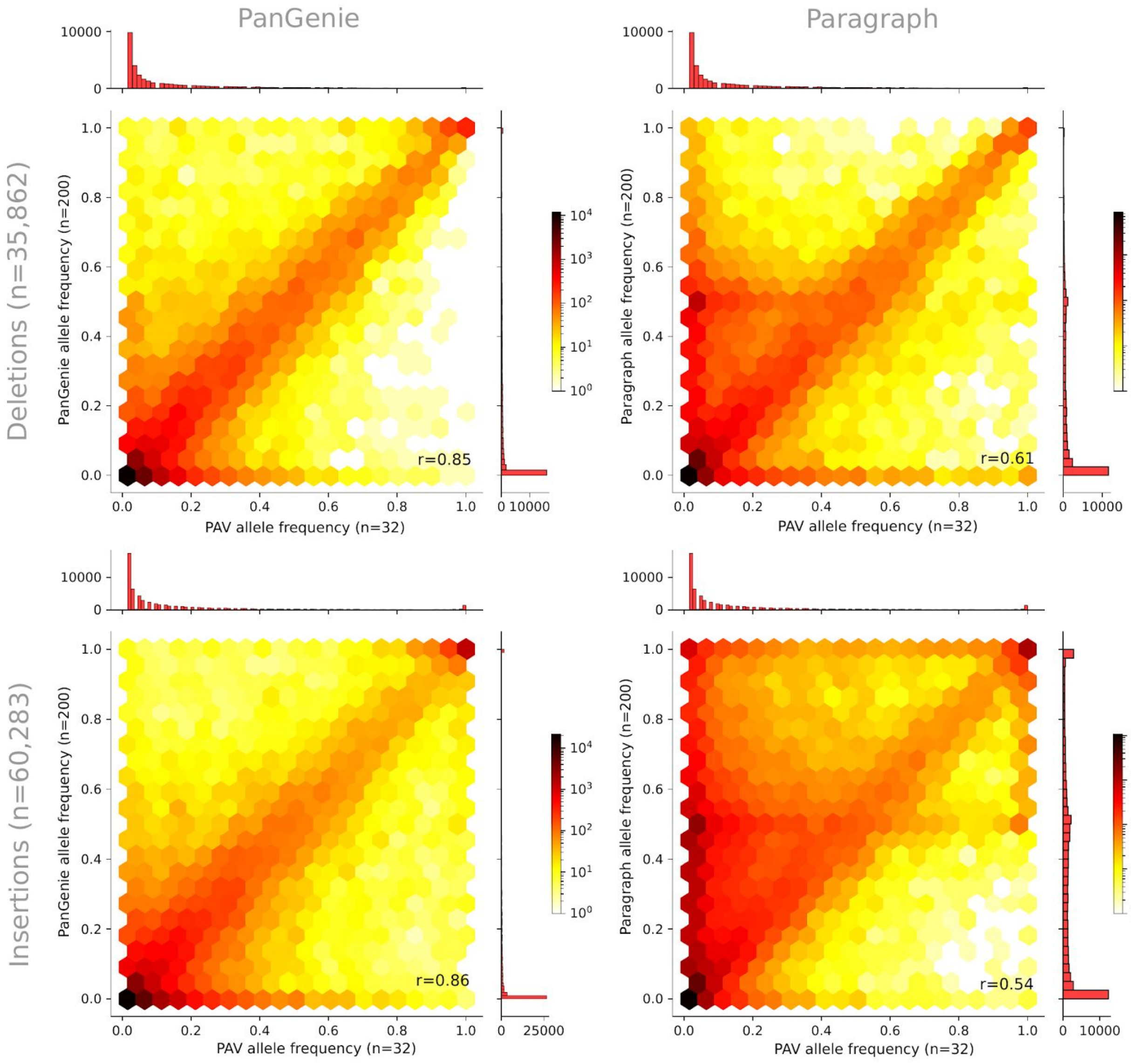


**Fig. S72. Comparison of PanGenie and Paragraph allele frequencies on the pilot set**

Paragraph and PanGenie were run on a subset of 100 trios (300 samples) in order to derive genotypes for all SVs (n=96,145). Allele frequencies were computed based on the genotypes of both methods for all 200 unrelated samples. PAV allele frequencies were computed based on all 64 assembly haplotypes.

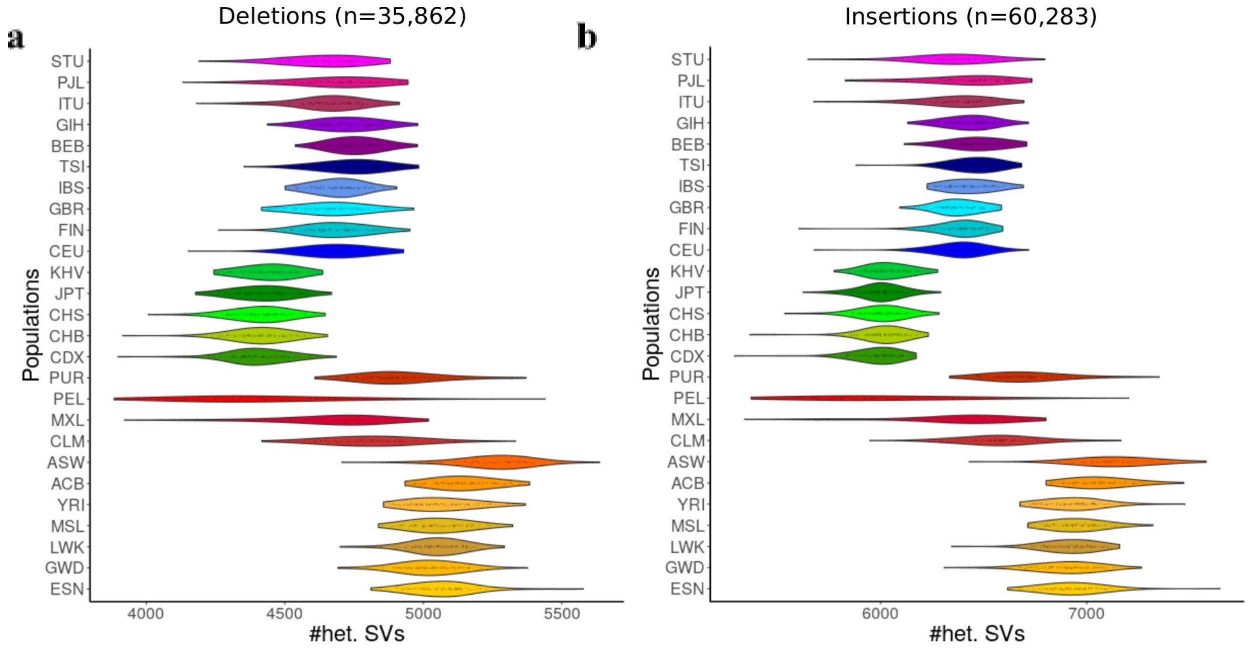


**Fig. S73. Number of heterozygous SVs per population**

Shown are the number of SVs typed as heterozygous by PanGenie for the different populations. The plots are based on the unfiltered callset containing all 96,145 SVs (35,862 deletions and 60,283 insertions).

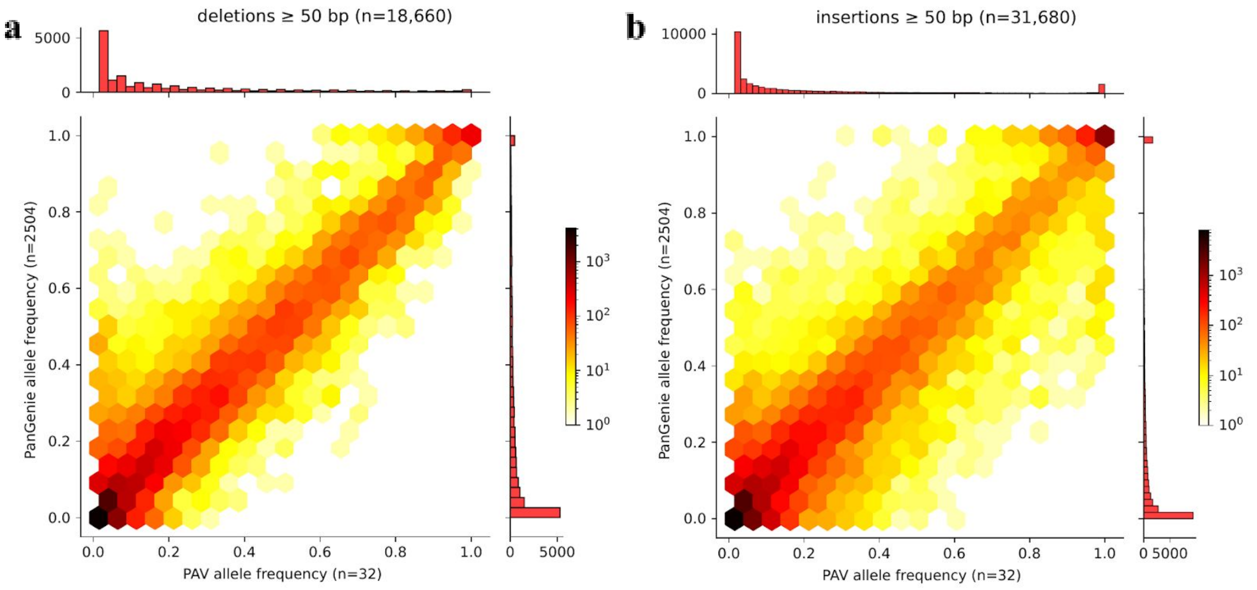


**Fig. S74. Allele frequencies of SVs in the lenient callset**

For PanGenie, allele frequencies were computed based on the genotypes of all 2,504 unrelated samples. The PAV allele frequencies were computed based on all 64 assemblies. Only SVs (≥50 bp) contained in our lenient callset (cutoff -0.5, n=50,340) were considered.

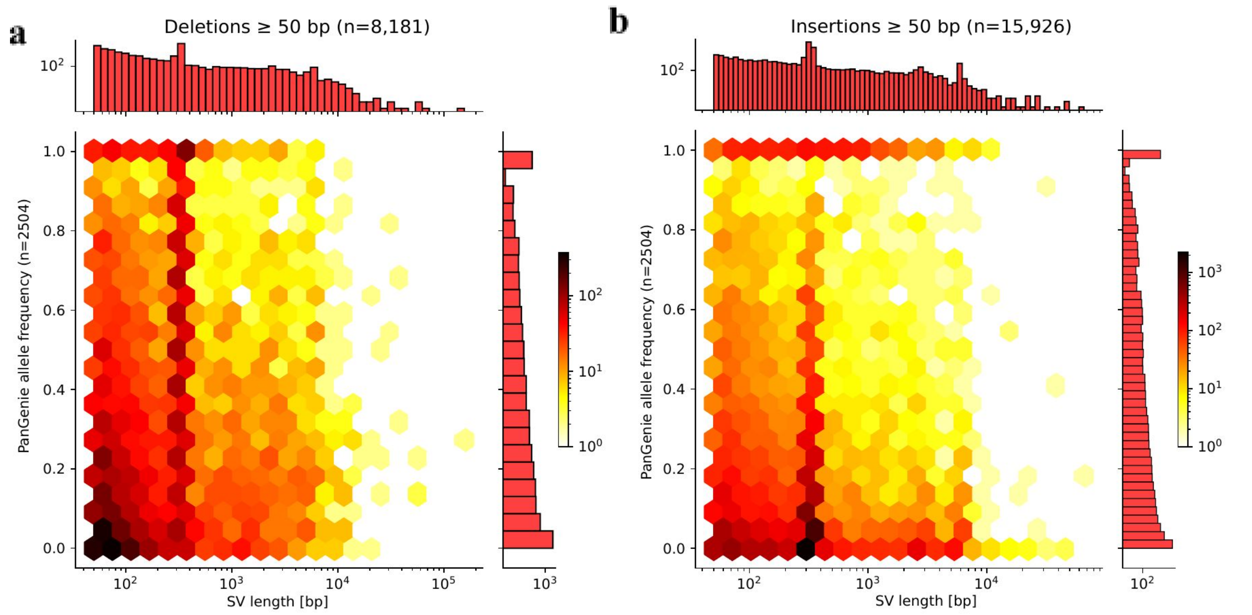


**Fig. S75. PanGenie allele frequency vs. SV length**

PanGenie allele frequencies were computed based on the genotypes of all 2,504 unrelated samples. Only SVs contained in the filtered set (n=24,107) are considered.

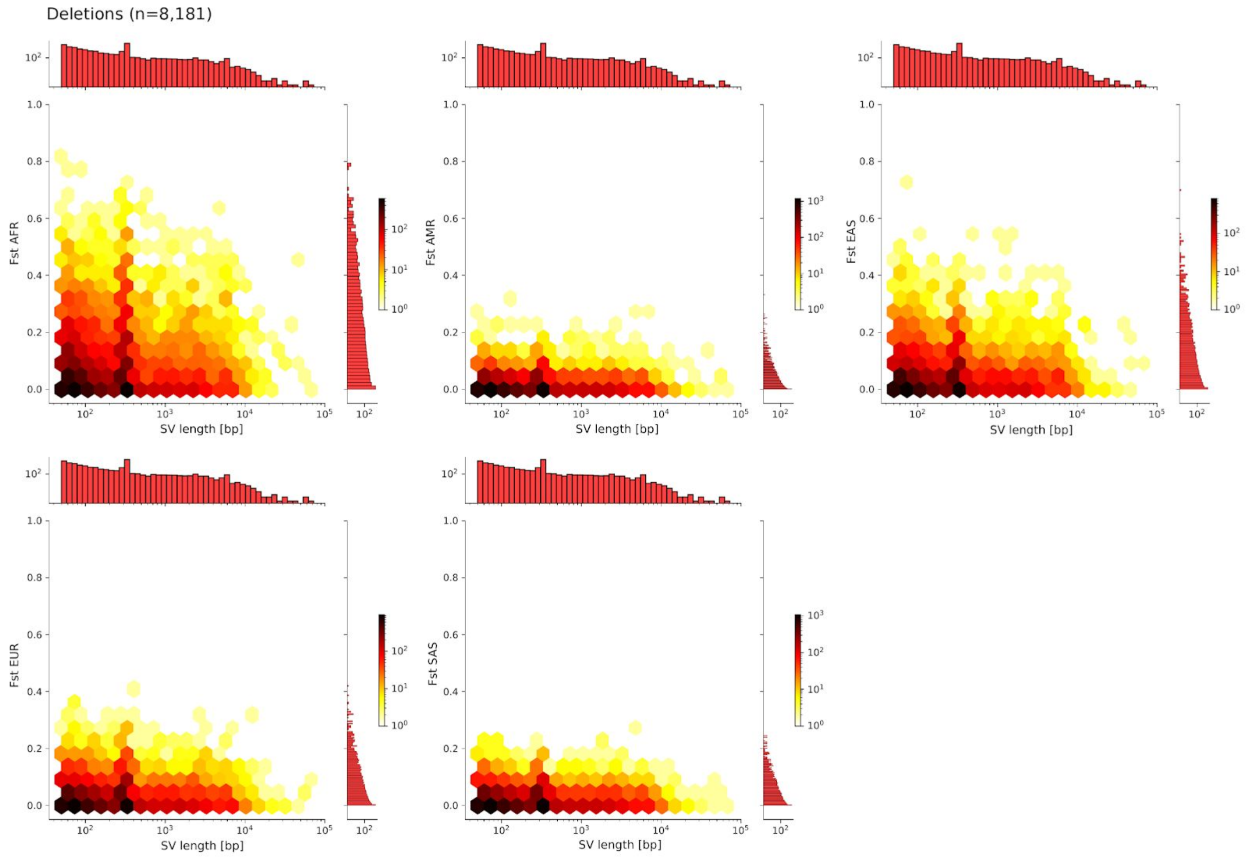


**Fig. S76. Fst vs. SV length for all superpopulations (deletions)**

For each superpopulation, Fst values were computed by comparing to the union of the remaining populations. The plots are based on the filtered PanGenie calls containing 8,181 deletions.

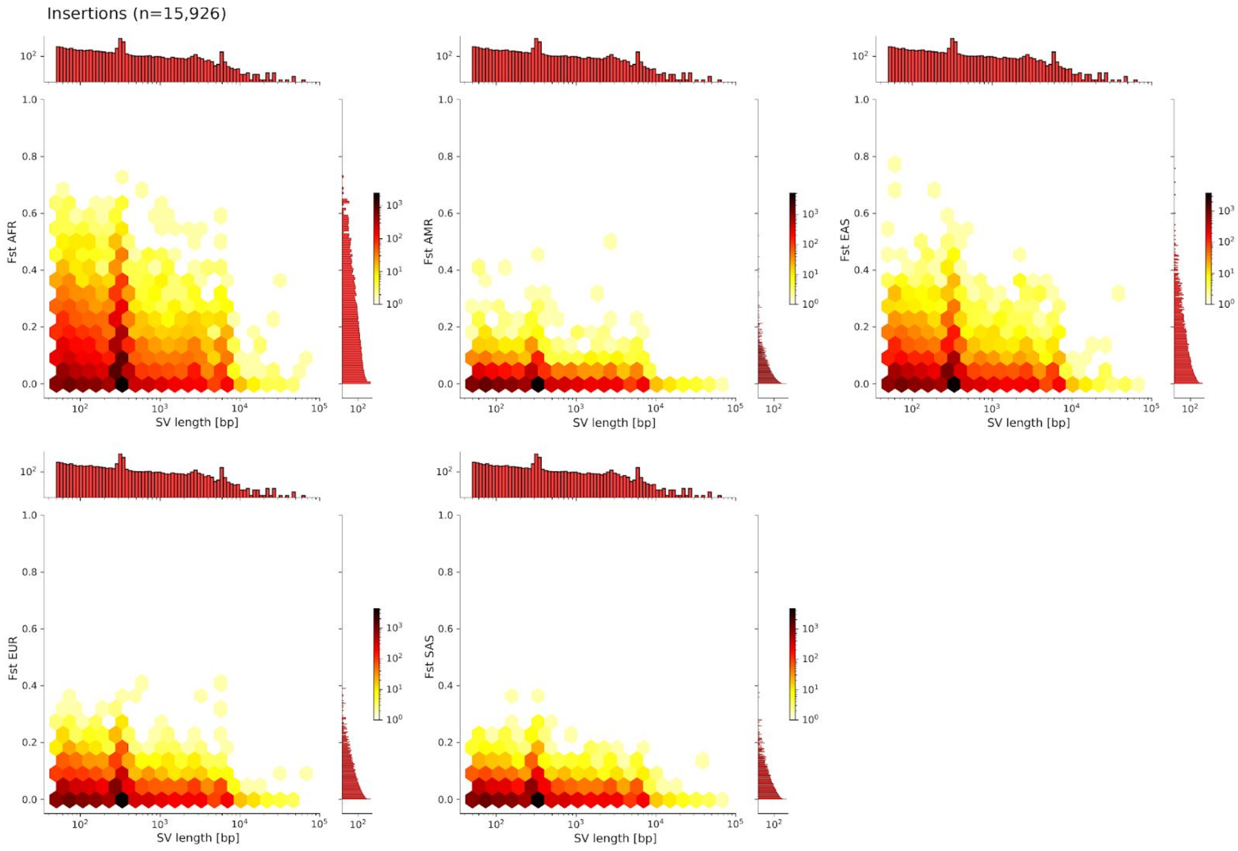


**Fig. S77. Fst vs. SV length for all superpopulations (insertions)**

For each superpopulation, Fst values were computed by comparing to the union of the remaining populations. The plots are based on the filtered PanGenie calls containing 15,926 insertions.

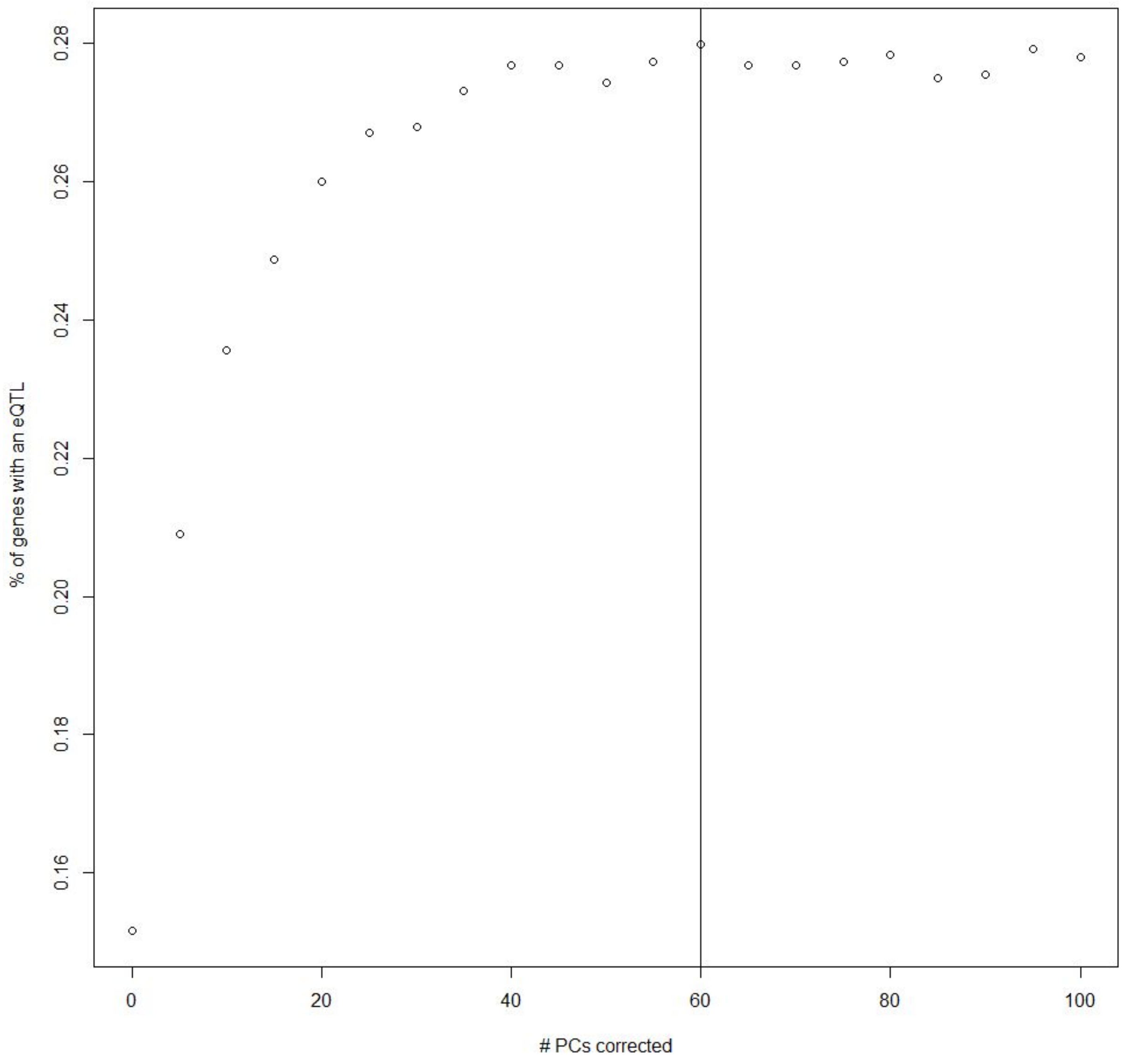


**Fig. S78. Optimization of the number of principal components (PCs) to correct for when mapping eQTLs**

The relation between the fraction of identified eGenes, the genes whose expression levels are associated with variation at a particular genetic variant, on chromosome 2 versus the number of PCs used to correct the expression data. The vertical bar (60) indicates the number of PCs used as fixed effect covariates in the eQTL mapping.

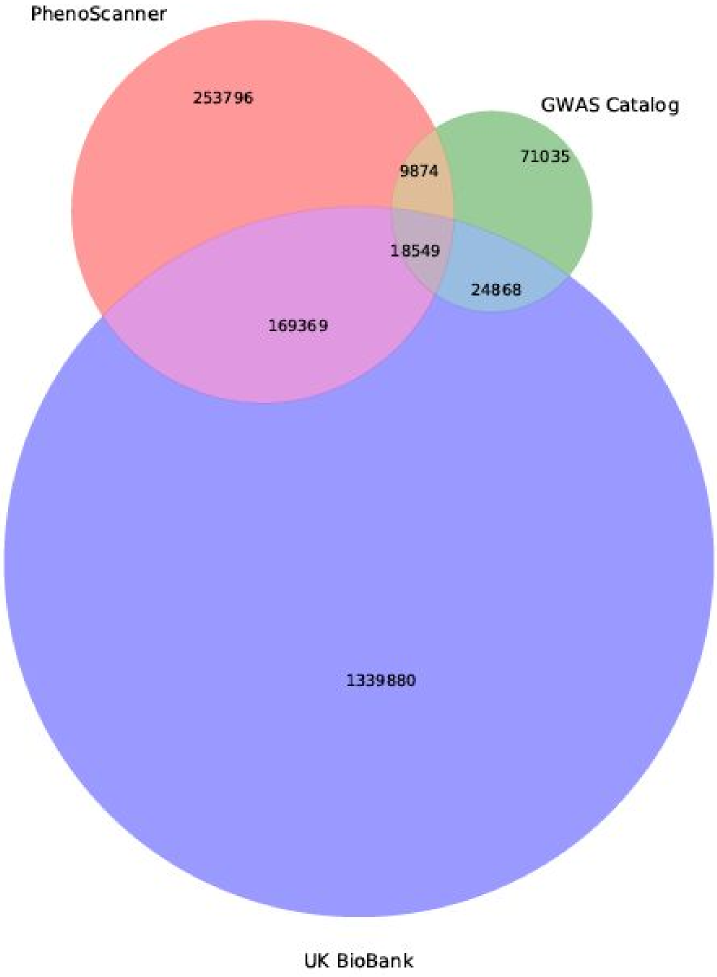


**Fig. S79. Venn diagram of known GWAS associations extracted from the GWAS catalog, UK Biobank and PhenoScanner (p-value < 1E-6)**

GWAS associations reported with p-value <= 1E-6 are kept in this study**.** SNP positions in UK Biobank and PhenoScanner are lifted over to GRCh38.

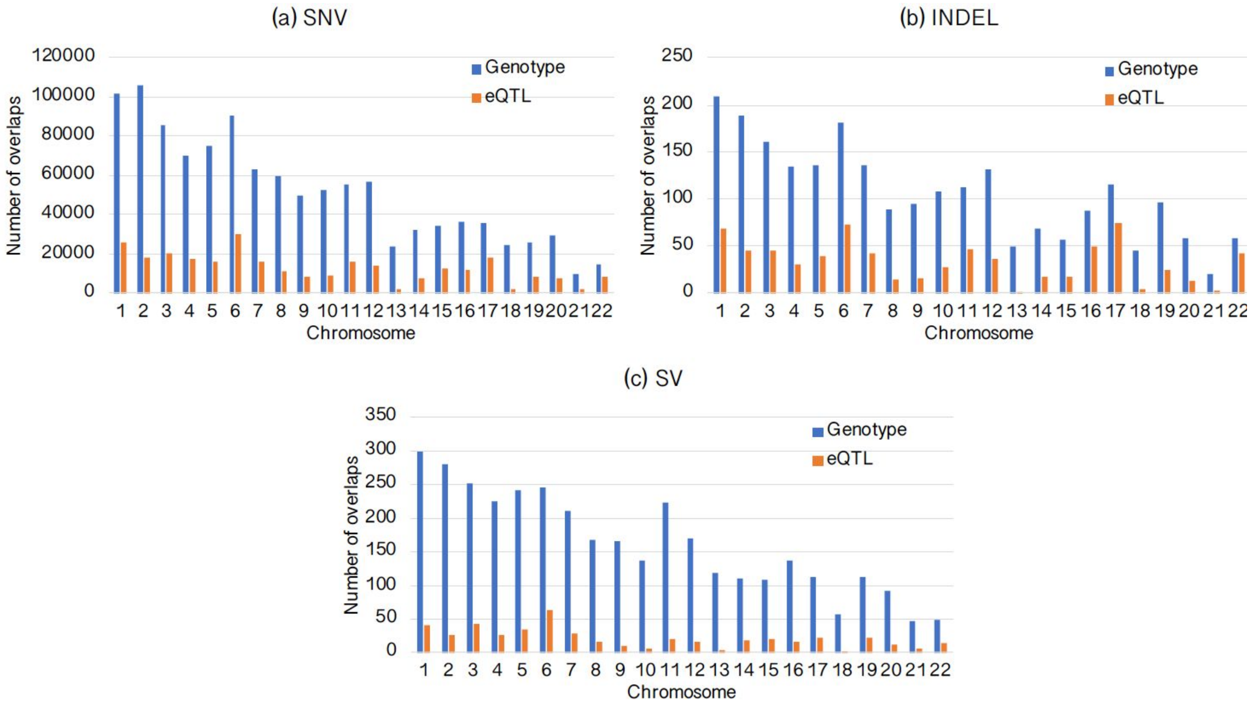


**Fig. S80. Numbers of genotyped variants (in blue) and eQTLs (in orange) of various types that overlap with known GWAS SNPs**

(a) SNVs with positions matching GWAS SNPs; (b) Indels that are at least 1bp overlap with GWAS SNPs; (c) SVs that are at least 1bp overlap with GWAS SNPs. X axis represents chromosomes and y axis represents the number of overlaps.

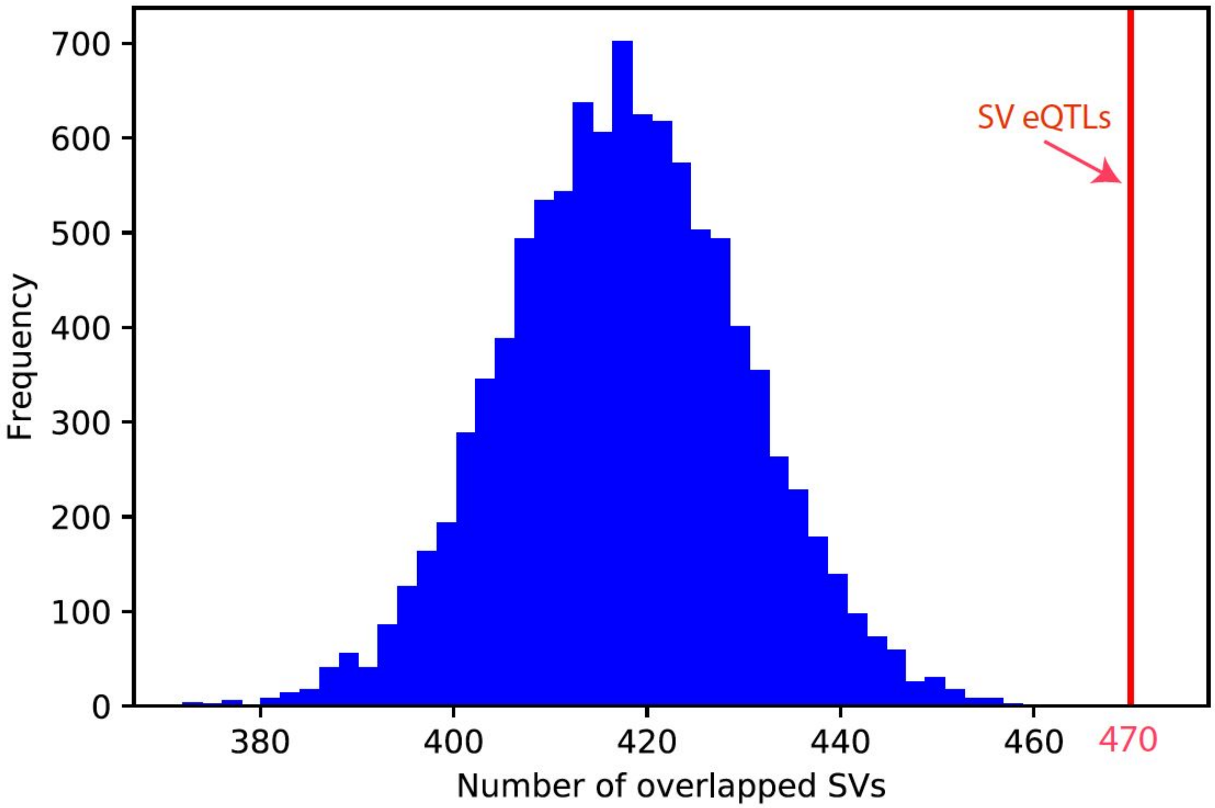


**Fig. S81. Enrichment of SV eQTLs for overlapping with GWAS SNPs**

X axis represents the number of SVs overlapped with GWAS SNPs, and y axis represents the frequency. The vertical line in red denotes the number (470) of SV eQTLs overlapped with GWAS SNPs. The enrichment was conducted by comparing the observed overlap of SV eQTLs with those random SV sets in 10,000 random permutations. In each permutation, SV regions were randomly selected from genotyped SVs that are input to the eQTL analysis pipeline, with the same distribution as SV eQTLs in terms of chromosome source, SV length, variant type, distance to transcription start site and distance to transcription end site of genes.

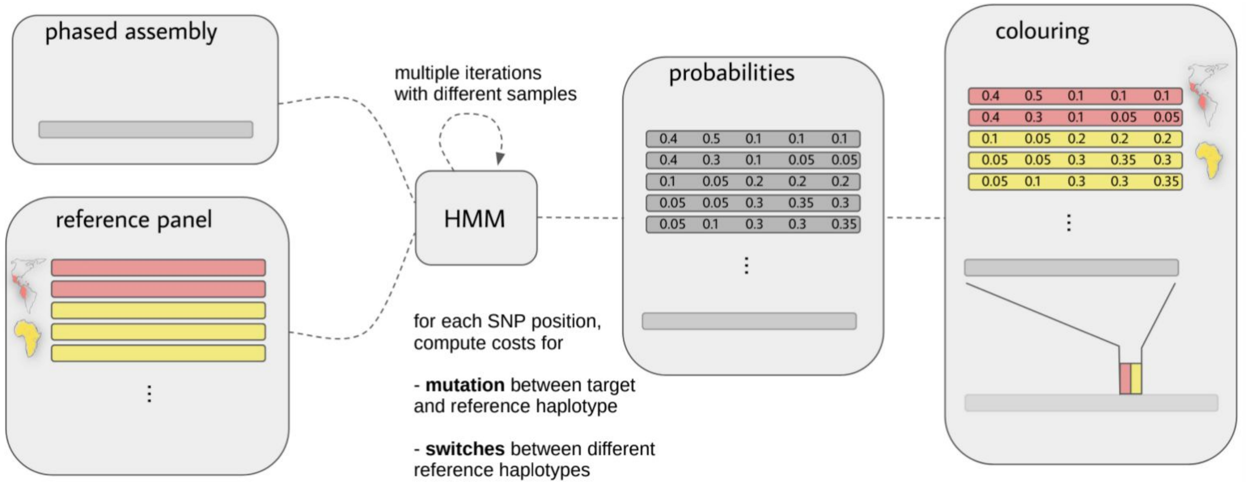


**Fig. S82. Hidden Markov model for local ancestry inference using haplotype-resolved assemblies**

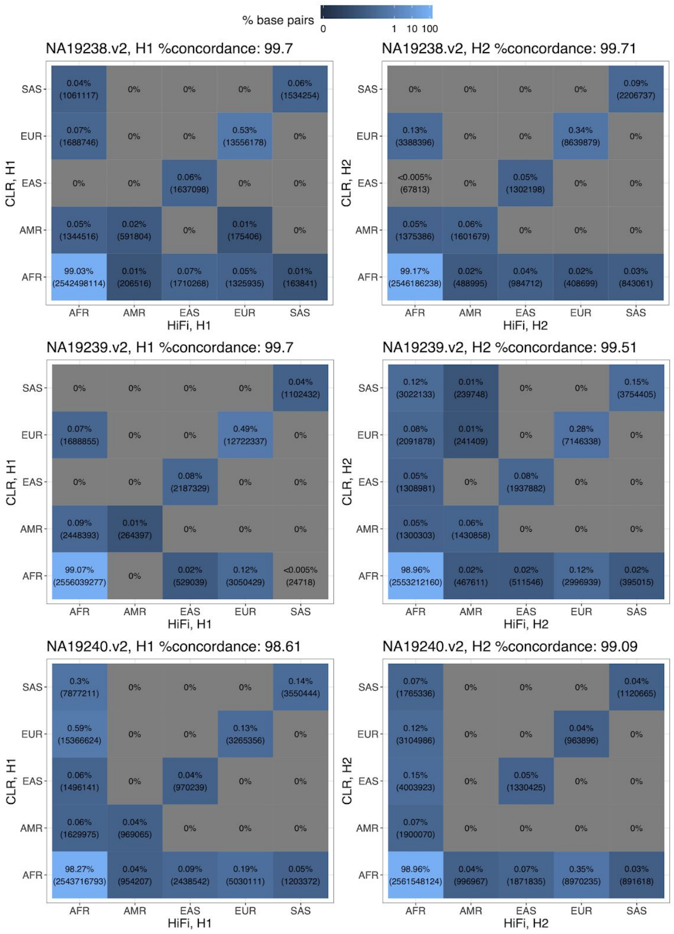


**Fig. S83. Concordant ancestry calls between HiFi and CLR data for the YRI trio**

In each row, the heatmaps indicate the fractions of concordant (diagonal entries) and discordant (off-diagonal entries) ancestry calls between the HiFi and CLR haplotypes from an individual genome. The numbers inside parentheses are the total number of base pairs in individual ancestry call combinations between the two datasets.

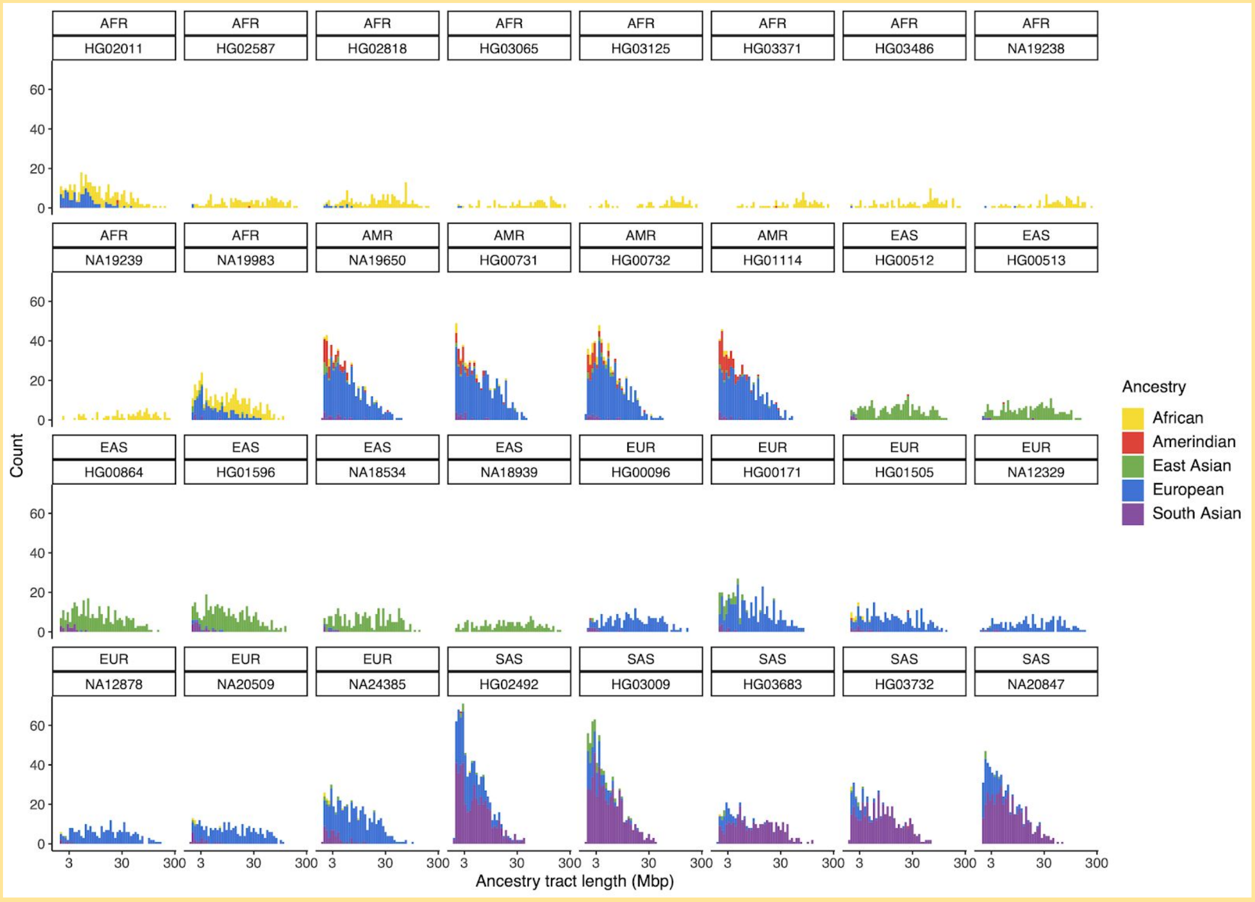


**Fig. S84. Length distributions of inferred genome-wide ancestry tracts for all haplotype-phased assemblies**

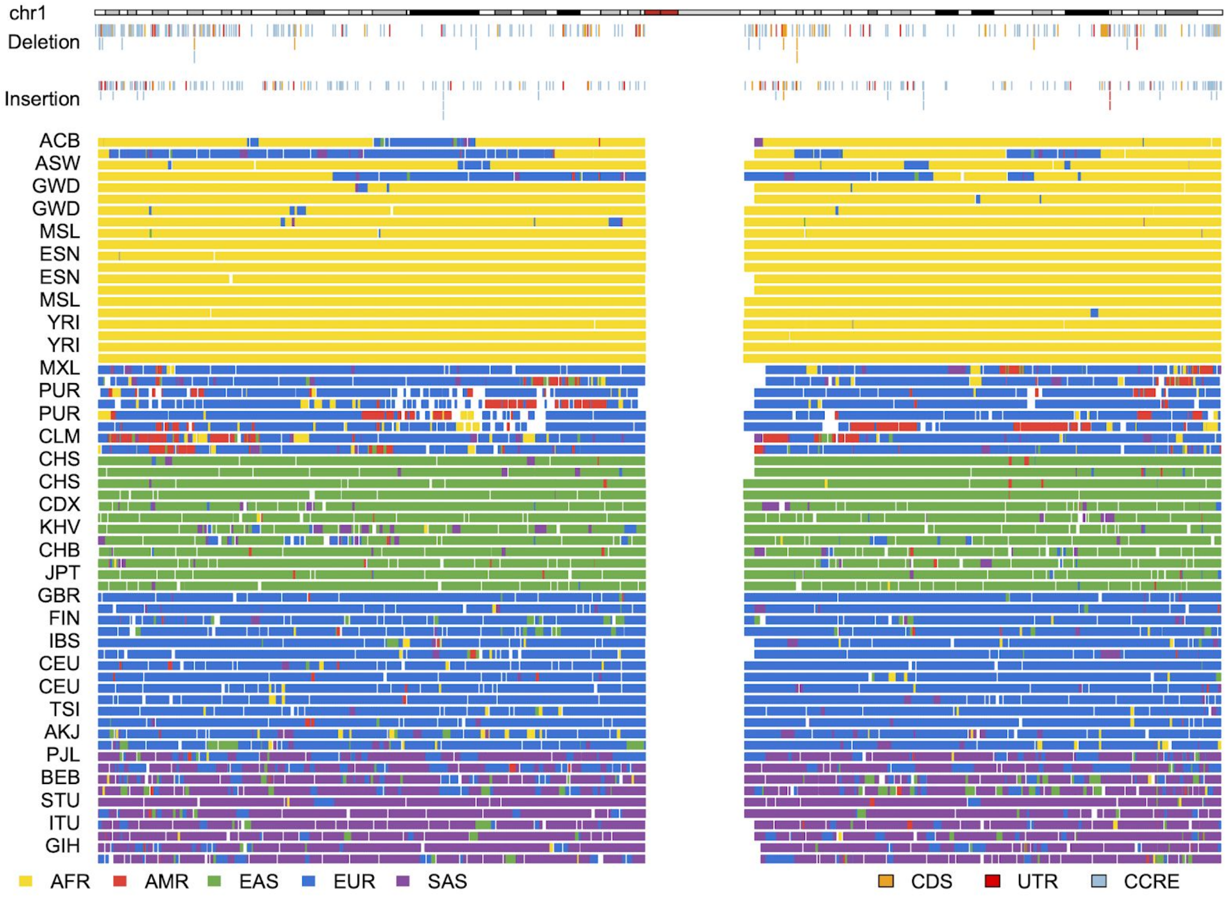


**Fig. S85. Inferred local ancestry blocks across chromosome 1 for haplotype-phased assemblies**

Ancestry calls are determined using the procedure described in the Methods. White spaces are discordant calls, known gaps, centromeres, and/or SDs. Deletion and insertion SVs that overlap with coding sequences (CDS), UTRs, and putative promoter sequences (CCRE) are annotated above.

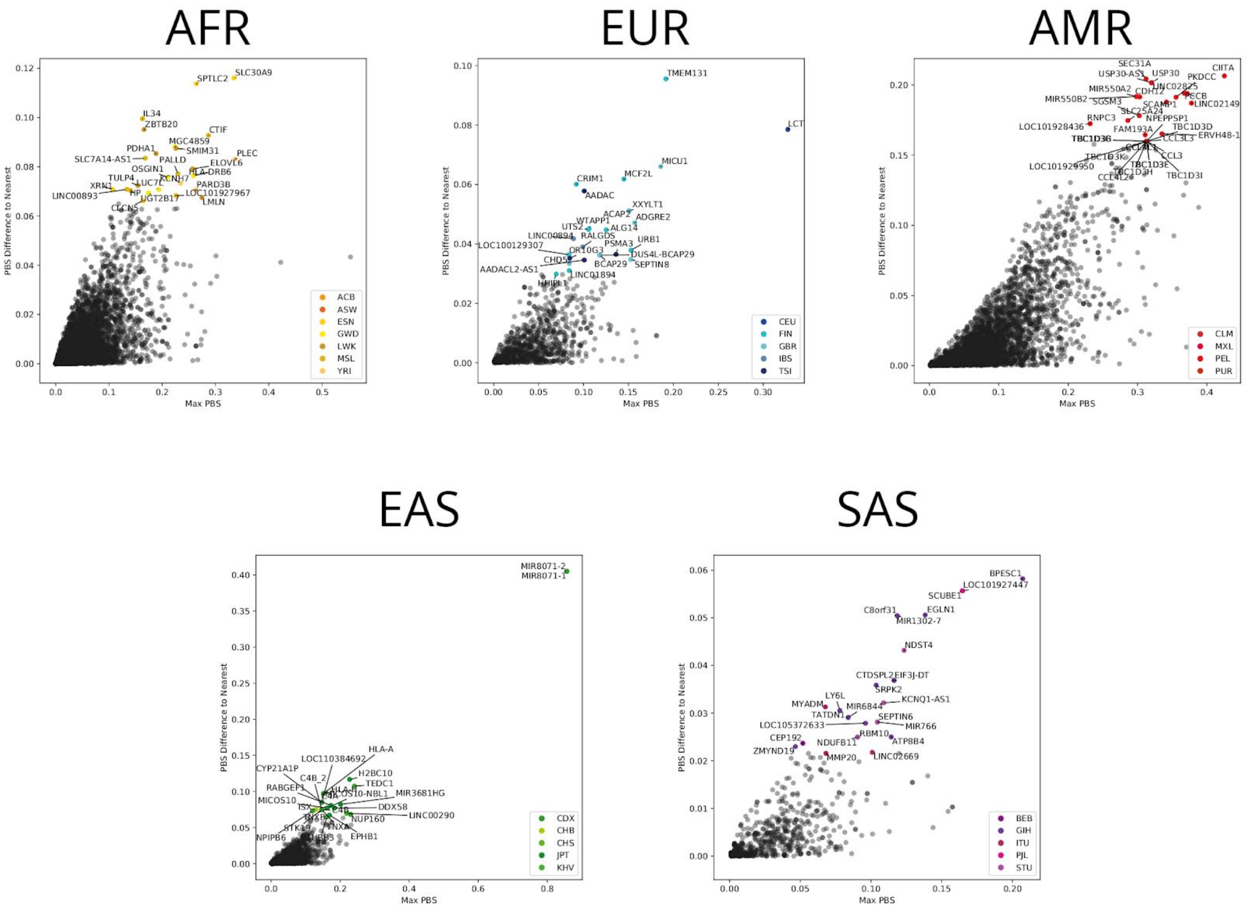


**Fig. S86. Top population branch statistics (PBS) hits per superpopulation**

For each gene, the maximum PBS value (horizontal axis) and the distance to the next highest PBS in another population (vertical axis) was determined. Each panel shows the PBS distribution for top hits in one superpopulation.

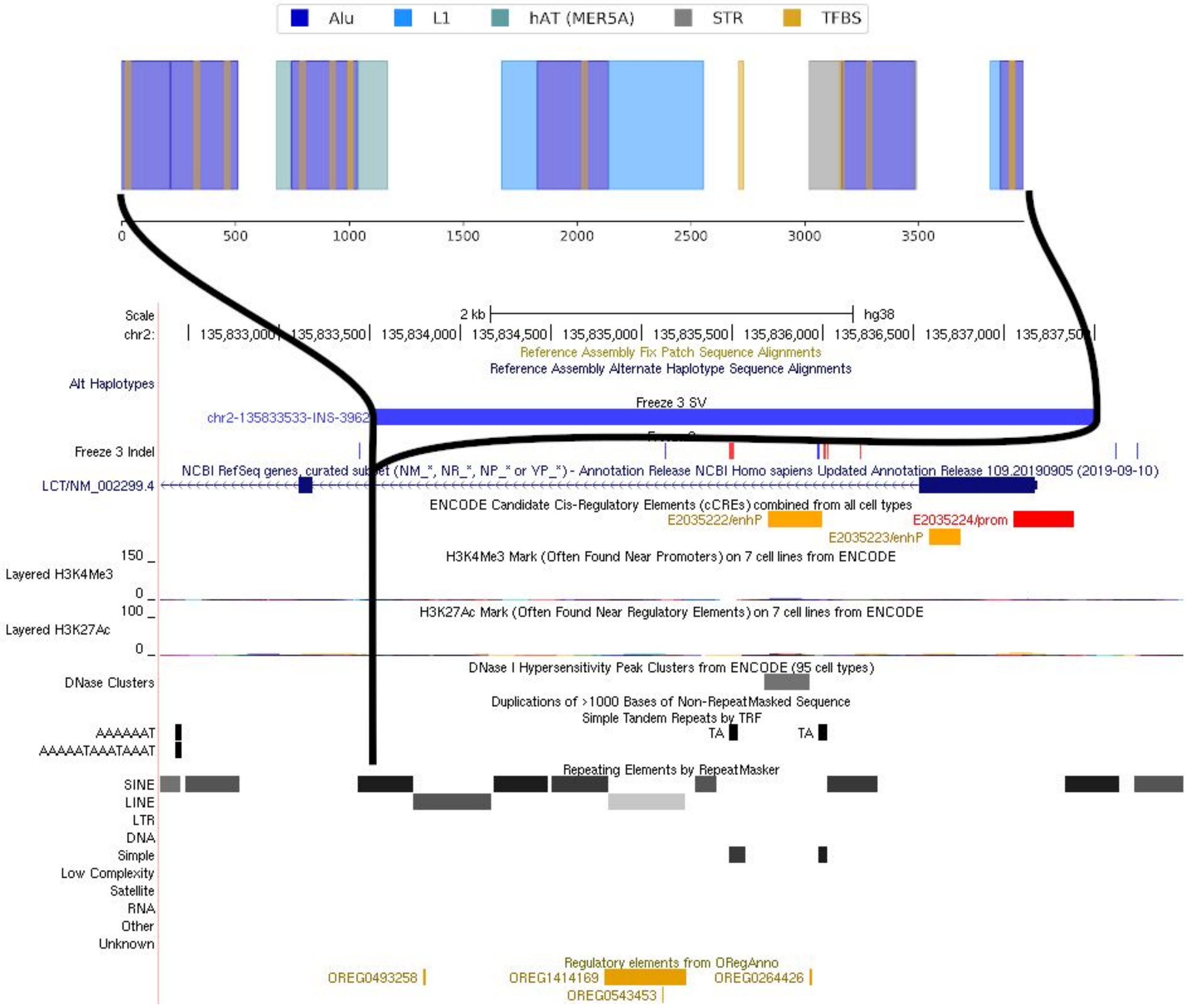


**Fig. S87. Structure of a 4.0 kbp insertion in the first *LCT* exon**

A 3,962 bp insertion in the first exon of *LCT* contains ancestral sequence with strong evidence for an AluY-mediated deletion captured as the reference allele. The insertion site is in a reference AluY, and 312 bp of AluY sequence (dark blue) is split between both ends of the inserted sequence. MEME predicts 11 transcription factor binding sites (yellow).
